# Cross-species functional modules link proteostasis to human normal aging

**DOI:** 10.1101/287128

**Authors:** Andrea Komljenovic, Hao Li, Vincenzo Sorrentino, Zoltán Kutalik, Johan Auwerx, Marc Robinson-Rechavi

## Abstract

The evolutionarily conserved nature of the few well-known anti-aging interventions that affect lifespan, such as caloric restriction, suggests that aging-related research in model organisms is directly relevant to human aging. Since human lifespan is a complex trait, a systems-level approach will contribute to a more comprehensive understanding of the underlying aging landscape. Here, we integrate evolutionary and functional information of normal aging across human and model organisms at three levels: gene-level, process-level, and network-level. We identify evolutionarily conserved modules of normal aging across diverse taxa, and importantly, we show that proteostasis involvement is conserved in healthy aging. Additionally, we find that mechanisms related to protein quality control network are enriched in 22 age-related genome-wide association studies (GWAS) and are associated to caloric restriction. These results demonstrate that a systems-level approach, combined with evolutionary conservation, allows the detection of candidate aging genes and pathways relevant to human normal aging.

**Highlights:**

- Normal aging is evolutionarily conserved at the module level.
- Core pathways in healthy aging are related to mechanisms of protein quality network
- The evolutionarily conserved pathways of healthy aging react to caloric restriction.
- Our integrative approach identifies evolutionarily conserved functional modules and showed enrichment in several age-related GWAS studies.

## Introduction

Aging is a process that affects all living organisms and results in a progressive decline in life function and a gain in vulnerability to death (Jones *et al*., 2014). In humans, aging is the main risk factor in a wide spectrum of diseases. The recent increase in human healthspan, also called ‘normal’, ‘disease-free’, or ‘healthy’ aging, is mostly due to improved medical care and sanitation (Greene, 2001; Rappuoli *et al*., 2011).

Major strides have been made in understanding the main molecular pathways underpinning the aging phenotype, leading to the definition of a number of “hallmarks” of aging, that may be common between species (López-Otín *et al*., 2013). Mitochondrial dysfunction and loss of proteostasis are two such conserved hallmarks of aging. Indeed, many comparative studies have shown mitochondrial dysfunction as a common feature of aging across species. Shared gene signatures in aging of *D. melanogaster* and *C. elegans* are linked to mitochondrial oxidative respiration, and similar results are observed in primates, including humans (McCarroll *et al*., 2004; de Magalhães, Curado and Church, 2009; Alexey A. Fushan *et al*., 2015). Collapse of proteostasis is another hallmark of aging that was shown to be important not only in short-lived species, but also in long-lived ones (Tian, Seluanov and Gorbunova, 2017). Loss of proteostasis is related to major human pathologies, such as Alzheimer’s and Parkinson’s disease, offering an opportunity to detect conserved candidate genes important in those age-related diseases (Labbadia and Morimoto, 2015; Sorrentino *et al*., 2017). The proteostasis network consists of three major mechanisms: protein synthesis, autophagy and the proteasome complex (Kaushik and Cuervo, 2015). Recent studies on the long-lived naked mole rat showed maintenance of proteasome activity throughout life (Rodriguez *et al*., 2012). Perturbations of components of the proteostasis network have already been observed in other species, such as mice (Pyo *et al*., 2013). Notably, caloric restriction, defined as a reduction of regular caloric intake by 20-40%, extends lifespan and delays the onset of age-related diseases in many species (Lee *et al*., 2006; Selman and Hempenstall, 2012; Bass *et al*., 2015; Mattison *et al*., 2017), in part through effects on mitochondria and proteostatic networks.

Although significant efforts have been made to uncover the identity of genes and pathways that affect lifespan, it is unclear to what extent the functional information of aging obtained from model organisms can contribute to human aging. Focusing on the process of aging in healthy individuals should improve the discovery of pathways important in natural aging. In addition, systems-level analysis of large datasets has emerged as an important tool for identifying relevant molecular mechanisms, as single gene-based methods are not sufficient to elucidate complex processes such as aging. The integration of various data types contributes to identify pathways and marker genes associated with specific phenotypes (Baumgart *et al*., 2016; Hasin, Seldin and Lusis, 2017). Notably, co-expression network analyses can help to elucidate the underlying mechanisms of various complex traits (Xue *et al*., 2007; van Dam *et al*., 2017).

To incorporate evolutionary and functional age-related information, we integrated transcriptome profiles of four animal species from young and old adults: *H. sapiens, M. musculus, D. melanogaster* and *C. elegans*. As a source of gene expression, we used human data from the large-scale Genotype-Tissue expression (GTEx) project (Mele *et al*., 2015), together with aging transcriptomes of model organisms. We identified the functional levels of conserved genetic modifiers important during normal aging, and related them to caloric restriction experiments and enrichments in age-related genome-wide association studies (GWAS). We used gene families as evolutionary information across distant species in a two-step approach to observe age-related conserved mechanisms. Our results show the contribution of age-related mechanisms from model organisms to human normal aging, with notably a demonstration of the conserved role of proteostasis in normal aging and in the reaction to dietary restriction.

## Results

### Data-driven integrative evolutionary approach to healthy aging

We used a three steps-approach to integrate transcriptomes across distant species (human and model organisms) and to identify evolutionarily conserved mechanisms in normal aging (Figure 1A). In the first step, we performed differential expression analysis between young and old samples in two tissues, skeletal muscle and hippocampus, from humans (*Homo sapiens*) and mice (*Mus musculus*), and in whole body for the fly (*Drosophila melanogaster*) and the worm (*Caenorhabditis elegans*). We also used transcriptome datasets related to caloric restriction in these species for validation. In the second step, we obtained 3232 orthologous sets of genes, ‘orthogroups’, across those four species (see Methods). Each orthogroup (OG) is defined as the set of the orthologous and paralogous genes that descended from a single ancestral gene in the last common ancestor to those four species (*H. sapiens, M. musculus, D. melanogaster, C. elegans*) and an outgroup species (*Amphimedon queenslandica*). Each orthogroup can contain a different number of genes, and was treated as a single functional meta-gene common to four species. We corrected for the orthogroup sizes by applying Bonferroni correction on the gene p-values from differential expression analysis within the orthogroup. Then, we selected a representative gene per species within orthogroups. We took the minimum Bonferroni adjusted p-value of a species-specific age-related gene from differential expression analysis. This allowed us to build ‘age-related homologous quadruplets’ (see Details in Figure S1A). The four p-values within each quadruplet were then summarized into a single p-value per quadruplet, by using Fisher’s combined test. We obtained 2511 gene quadruplets in skeletal muscle, 2800 in hippocampus, and 1971 in caloric restriction experiments (Table S4). We characterized their biological relevance by functional enrichment. In the third and final step, those quadruplets of age-related genes were used to build a co-expression network per species (Figure S1B). These networks were then integrated together using order statistics into one cross-species age-related network. We performed community search algorithm on this network to obtain age-related and evolutionarily conserved modules. The modules were then tested for functional enrichment and for enrichment in GWAS hits.

**Figure 1.**
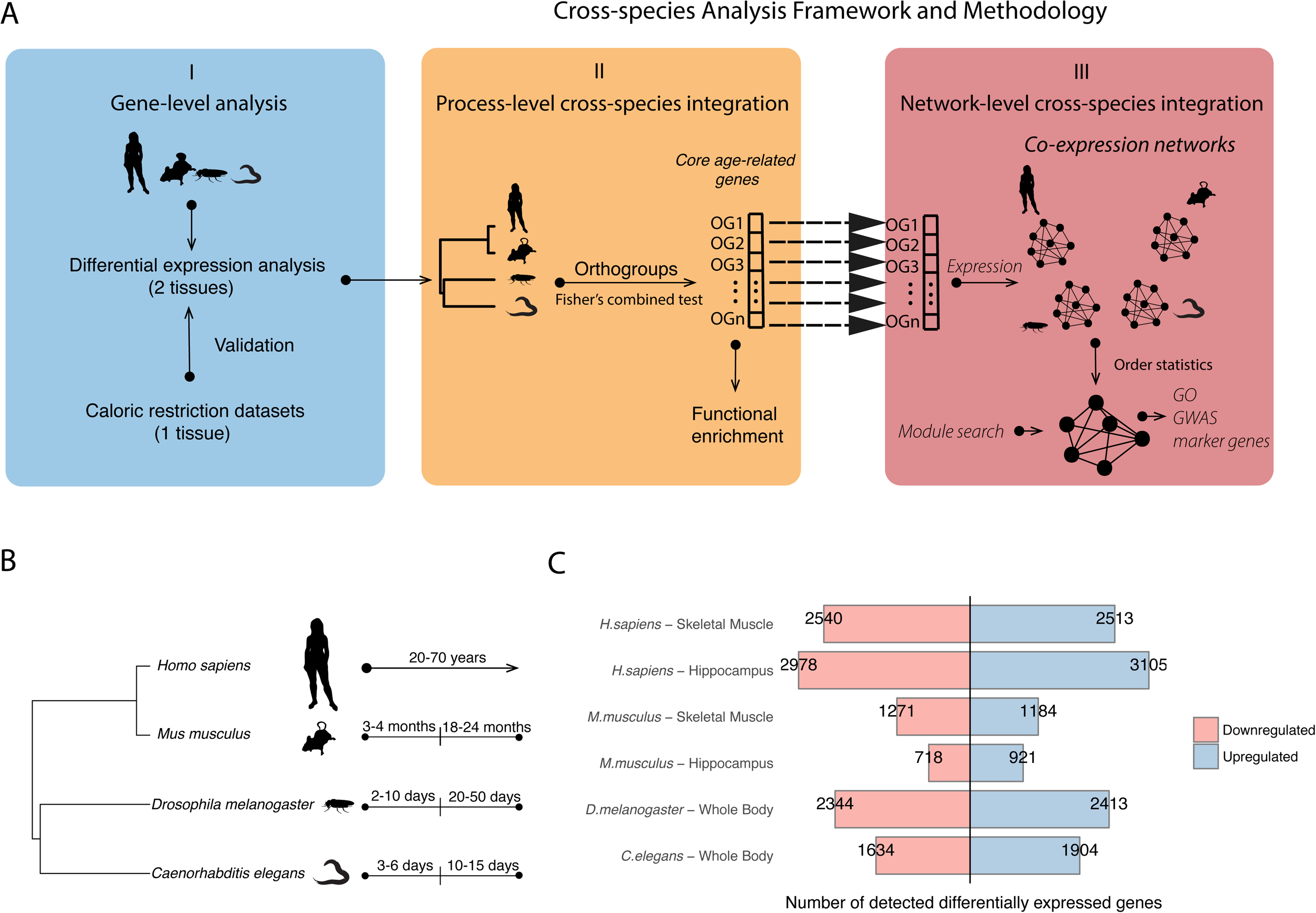
Study design and differential gene expression analysis. (A) An overview of the integration process based on transcriptomes across the species. (I) Analysis starts at the single-gene level by performing differential expression analysis per species between young and old adults (all samples in case of GTEx human data), and determining the orthogroups across species. (II) The orthogroups (OG) are summarized to single genes that represent age-associated conserved genes. (III) The same genes are then used to build the co-expression networks per species and being integrated in the final cross-species network. (See Methods, Figure S1A and S1B) (B) The species used in the study with their phylogenetic relations and the alignment of their ages categories. (C) Barplots representing the numbers of significantly differentially expressed age-related genes (FDR < 0.1) in old healthy individuals in each dataset used. Blue (resp. red) bars represent genes significantly up-(resp. down-) regulated in old adults.

### Age-related gene expression patterns in four spcies

To study normal aging, we restricted ourselves to transcriptomic studies with at least one young adult and one old adult time-point, adult being defined as after sexual maturity (Figure 1B). Transcriptomes had to come from control samples (model organism datasets) or relatively healthy individuals (GTEx dataset). We defined young and old adults across species as follows: young: 3-4 months for *M. musculus*, 2-10 days for *D. melanogaster*, 3-6 days for *C. elegans*; old: 18-24 months for *M. musculus*, 20-50 days for *D. melanogaster*, 10-15 days for *C. elegans*. For the GTEx data, samples from all adults (20-70 years old) were taken into account in a linear model to detect differentially expressed genes. In human and mouse, we focused on two tissues, skeletal muscle and hippocampus, because they are known to be profoundly affected by aging. During aging, skeletal muscle is affected by sarcopenia (Marzetti and Leeuwenburgh, 2006). Changes in hippocampus function have a significant impact on the memory performances in elderly people (Driscoll *et al*., 2003). Thus both tissues are susceptible to aging-related diseases. For human, we used transcriptomes of 361 samples from skeletal muscle tissue and 81 samples from hippocampus from GTEx V6p. For the other species we used diverse publicly available transcriptomic datasets (Table S1). The sample sizes for model organisms were variable, from 3 to 6 replicates per time-point. In order to compare samples between young and old age groups, we fitted linear regression models for each dataset. In addition, in the GTEx dataset we controlled for covariates and hidden confounding factors to identify genes whose expression is correlated or anti-correlated with chronological age, taking into account all samples (see Methods).

We observed uneven distributions of up- and down-regulated genes with aging across different species and datasets (Figure 1C, Table S2), suggesting variable responses to aging and different power of datasets. The human hippocampus shows substantially more age-related gene expression change than skeletal muscle (6083 *vs*. 5053 differentially expressed genes, FDR < 0.1). However, mouse hippocampus shows less gene expression change than skeletal muscle (1639 *vs*. 2455 differentially expressed genes, FDR < 0.1). These differences are due in part to the smaller sample size of the mouse skeletal muscle study. We limited our analysis to genes that were expressed in at least one age group, leading to detection of 15-40 % of genes that exhibits age-related gene expression changes. Of note, these changes are often very small, typically less than 1.05 fold in humans and less than 2-fold in animal models.

It has been previously reported that there is a small overlap of differentially expressed genes among aging studies (de Magalhães, Curado and Church, 2009; Yang *et al*., 2015). To make results easily comparable across species, the young and old adults of one species should correspond to young and old adults of another species (Flurkey, M. Currer and Harrison, 2007). Our clustering shows good consistency across age groups of samples between species, based on one-to-one orthologous genes with significant age variation (FDR < 0.05) (Figure 2A, Figure S2). Yet there is a low overlap of one-to-one orthologous genes with significant expression change in aging (Table S3). This observation is in line with two studies showing that the overlap between individual genes associated with aging did not reach the level of significance (Smith *et al*., 2008; Alexey A. Fushan *et al*., 2015). To go beyond this observation, we correlated log-transformed fold change (old/young; or log of α age-related regression coefficient in human) between human and model organisms. We observed weak pairwise correlations (Figure S3) when comparing single genes. This indicates that most transcriptional changes on the gene level are species-specific, and that there is little evolutionary conservation to be found at this level.

**Figure 2.**
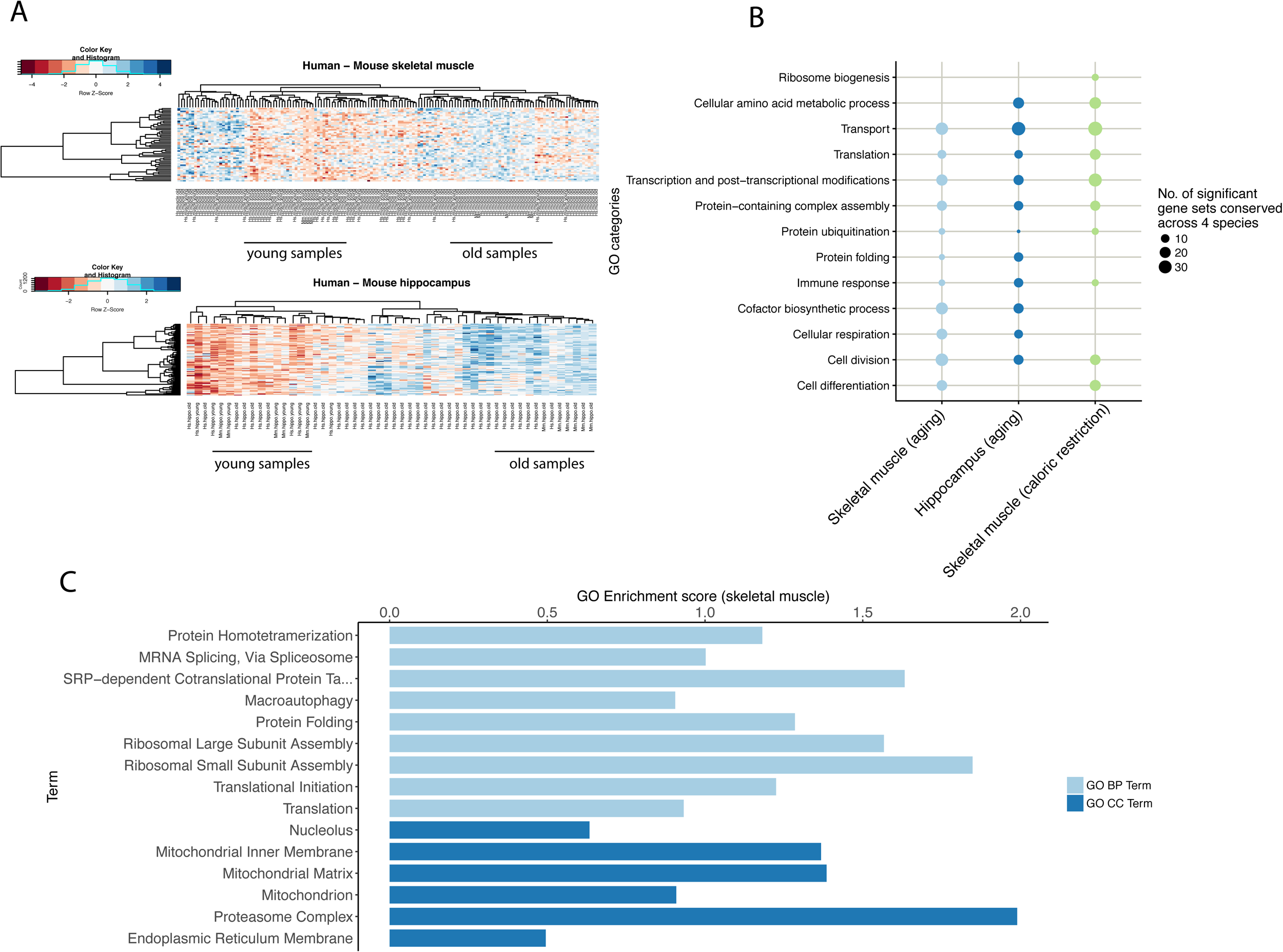
Functional enrichment analysis of integrated age-associated conserved genes. (A) Clustering of the age-related samples between human (20-30y; 61-70y) and mouse. The heatmaps show good concordance between the young and old samples between species based on the 1-1 orthologous genes that are differentially expressed. (B) Bubble plot showing the number of GO categories with conserved change of expression in aging between species. The analysis only includes categorized GO terms that are significant (FDR < 0.05) and unique to the homologous quadruplets enrichment. (C) GO enrichment of genes involved in processes related to proteostasis based on cellular component (CC) and biological process (BP). Lengths of bars represent GO log2-transformed enrichment scores.

### Cross-species integration at the process-level reveals proteostasis-linked age-related mechanisms

To assess the age-related gene expression changes on a functional level in healthy individuals per species, we performed gene set enrichment analysis (GSEA) (Subramanian *et al*., 2005) using gene ontology (GO) annotations (Gene Ontology Consortium *et al*., 2000; The Gene Ontology Consortium, 2017). We then selected significant GO terms (FDR < 0.20) that we grouped into broader categories.

All species showed a general pattern of down-regulation of metabolic processes, such as mitochondrial translation (GO:0032543) in human GTEx skeletal muscle tissue, nucleotide metabolic process (GO:0009117) in mouse muscle tissue, cellular respiration (GO:0045333) in fly whole body, and oxoacid metabolic process (GO:0043436) in worm whole body (Figure S4). The pattern of metabolic down-regulation was stronger in muscle for both human and mouse. The processes that were down-regulated in hippocampus were related to behavior (GO:0007610), cognition (GO:0050980) and neurotransmitter secretion (GO:0007269) in human, and to synaptic signaling (GO:0099536) and axonogenesis (GO:0007409) in mouse. This confirms that there is a tissue-specific signal in normal aging. Due to small samples size of the mouse skeletal muscle dataset, we were able to detect only down-regulated metabolic processes. In addition to metabolism, we observe strong immune systems response to aging, such as regulation of cytokine production (GO:0001817) in human hippocampus or leukocyte-mediated immunity (GO:0002443) in mouse hippocampus. These results are consistent with known links between metabolism, immunity and aging (Lanna *et al*., 2017).

We aggregated processes on the functional level across four species using evolutionary information to observe common age-related mechanisms rather than tissue-specific mechanisms. We integrated differential expression analysis from each species, as described above. We obtained 2010 genes in skeletal muscle / whole body, 2075 genes in hippocampus / whole body, and 1962 genes in caloric restriction experiments (Fisher combined tests, FDR < 0.10) (Table S4). We examined their biological relevance using Gene Ontology enrichment analysis (GEA) based on human annotation (Figure 2B). We did not take into account whether the processes that are shared across species are regulated in the same direction, but rather whether they are consistently perturbed during aging.

We obtained 100 significant GO terms (FDR < 0.05) related to biological processes, and aggregated them into broader GO categories. While our species-specific analysis mostly shows tissue-specific pathways, we found that terms with an evolutionarily conserved relation to normal aging are strongly enriched for processes involved in proteostasis, or protein homeostasis. The proteostasis-linked processes are more conserved than expected by chance (Figure S6). The other conserved processes are related to transport, translation, transcription and post-transcriptional modifications, and protein ubiquitination (Figure 2B, Table S5). We also confirmed previously known evolutionarily conserved age-related pathways, such as cellular respiration and immune response. Integrating caloric restriction datasets across the four species showed enrichments in similar processes (Figure 2B).

While most of the shared processes have been previously linked to aging, we focused on proteostasis and related processes. To characterize in more detail the specificity of proteostasis-linked processes, we investigated their enrichment strength in the large human GTEx dataset (Figure 2C). Since proteostasis perturbation is detected both through the GO domains of cellular localization and of biological process, we investigated these two domains, and obtained similar enrichments for both skeletal muscle (Figure 2C), and in hippocampus (Figure S6, Table S6). The most enriched cellular component terms in skeletal muscle were related to proteasome complex (GO:0000502, enrichment score: 1.99) and to mitochondrial matrix (GO:0005759, enrichment score: 1.38). We also observed strong enrichment of ribosomal large (GO:0000027, enrichment score: 1.57) and small subunit (GO:0000028, enrichment score: 1.84), of protein homotetramerization (GO:0051289, enrichment score: 1.18), and of GO biological processes that are part of the protein quality control network. Overall, the translation and proteasome complexes appear to be the parts of the protein quality control network whose involvement in aging is both evolutionarily conserved across different species, and significantly enriched in human healthy aging. Interestingly, we also detect the mRNA splicing pathway as a part of the conserved processes between species.

The direction of the changes in conserved proteostasis processes in humans is consistent with a relation between loss of proteostasis and healthy aging (Figure 3). Although macroautophagy did not show a strong enrichment score in the Figure 2C (GO:0016236, enrichment score: 0.90), there is down-regulation of the conserved genes associated with macroautophagy (Figure 3A), translation (Figure 3B), and the proteasome complex (Figure 3C), which are important in the protein quality network. Similar results are observed in hippocampus, although not with a strong signal as in skeletal muscle (Figure S7). The changes during healthy aging in both tissues are rather subtle but significant (Figure 3, Table S7).

**Figure 3.**
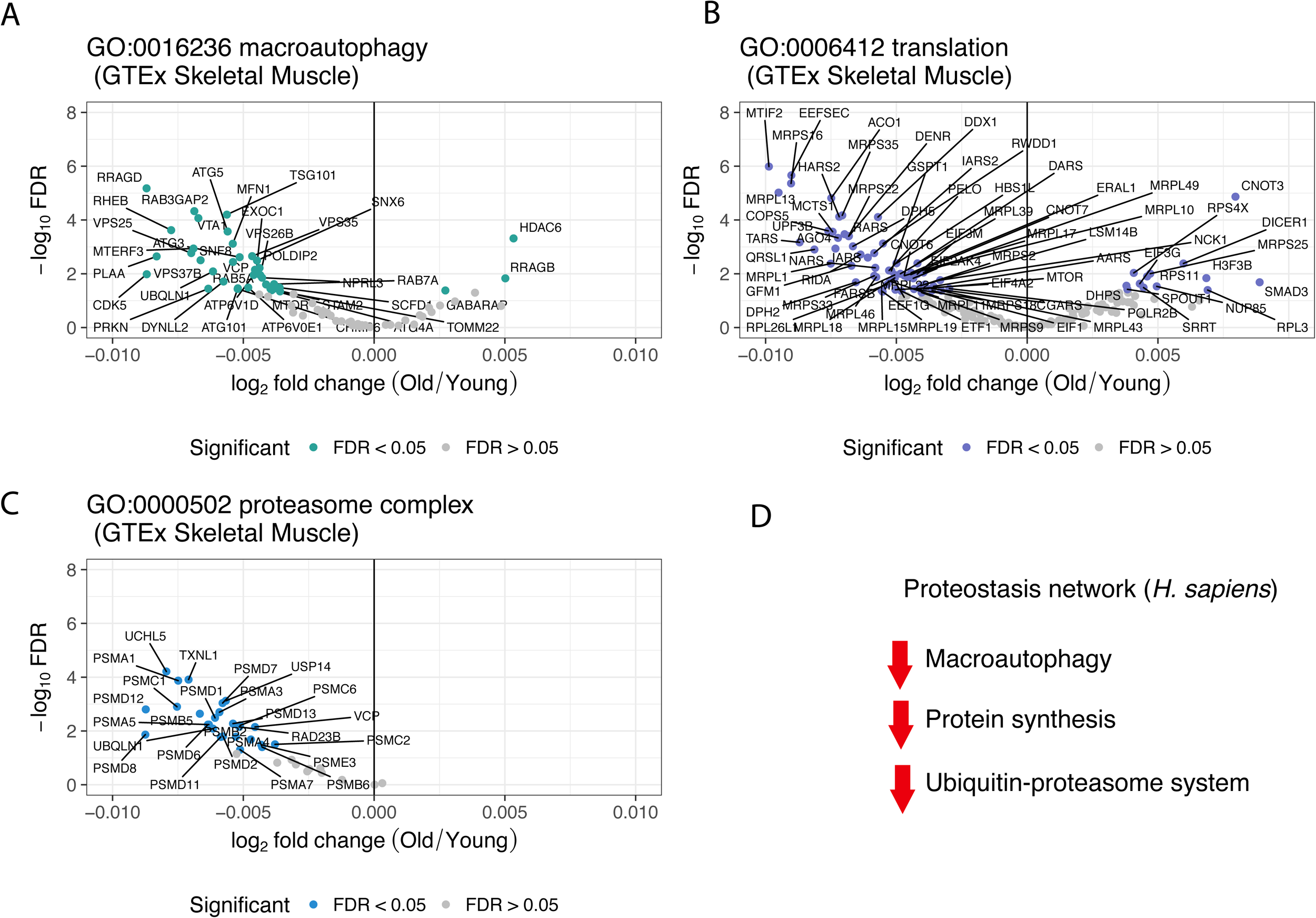
Gene expression changes in the main aspects of the proteostasis network in healthy aging human skeletal muscle. Conserved genes from macroautophagy (A), translation (B) and proteasome complex (C) in GTEx skeletal muscle data. Grey, conserved genes that are not significant (FDR > 0.05) in human GTEx skeletal muscle data. The x-axis of the volcano plots shows the log2 of age-regression coefficient (log2 slope, Formula 1) across the samples in GTEx data (see Methods; Formula 1), named log2 fold-change. (D) Schematic outline of the gene expression direction of the proteostasis-linked processes in aging human muscle.

### Fuctional characterization of cross-species age-related network identifies candidate genes related to healthy aging

To characterize age-related processes at a systems-level and to prioritize conserved marker genes associated with normal aging, we constructed probabilistic networks. These were based on prioritization of co-expression links between conserved age-related genes across four species. These genes became nodes in the multi-species network. Thus the connections between the conserved age-related genes are based on evolutionary conservation, and prioritized according to the their co-expression in each species.

Our integrative network analysis initially identified 20 and 14 modules for skeletal muscle and hippocampus, respectively. We randomized our networks 100 times based on the same number of conserved genes per experiment and obtained significantly higher numbers of gene-gene connections than in the original network (permutation test, p = 0.0198) (Figure S9). Thus aging networks appear to be lowly connected. We focused only on the modules larger than 10 genes; there were 12 such modules per tissue. These modules ranged in size from 16 (M7 hippocampus) to 191 genes (M12 hippocampus) (Figure 4A and 4B, Table S8). The networks were summarized to module level (module as a node), and we observed strong inter-modular associations. This analysis provided several levels of information. First, it provided a small number of coherent gene modules that represent distinct transcriptional responses to aging, confirming the existence of a conserved modular system. Second, it detected conserved marker genes affected during aging, discussed below.

**Figure 4.**
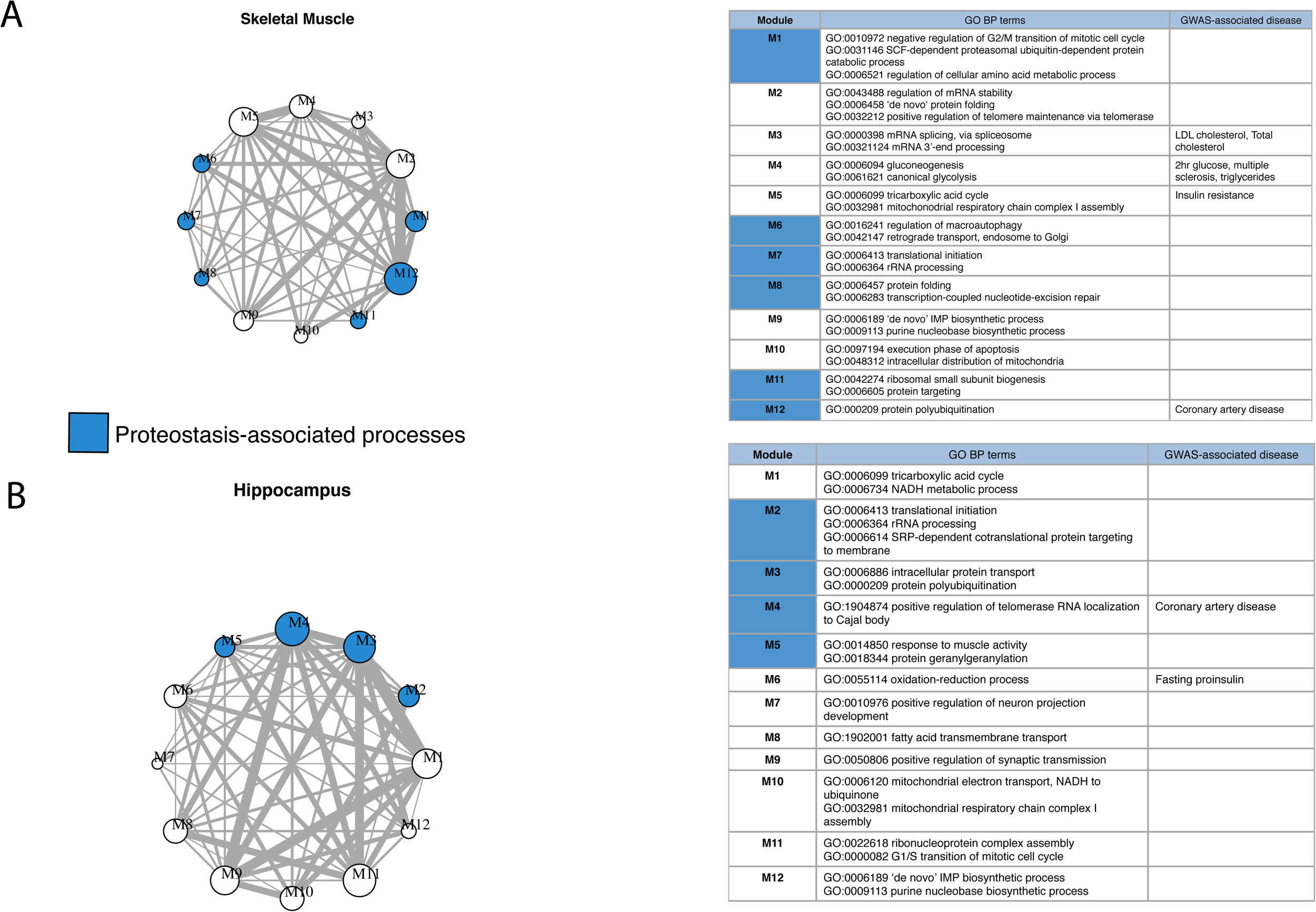
Cross-species aging-associated skeletal muscle and hippocampus functional modules and GO enrichments. Module networks of skeletal muscle (A) and hippocampus (B) with GO and GWAS enrichments for modules of size greater than 10. The tables on the right show top GO BP terms (FDR < 0.1) enriched in the skeletal (upper panel) and hippocampus (lower panel) modules. The GWAS-associated disease column in the same table contains associations to the module passing a threshold of FDR < 0.2.

To determine which of the conserved aging-associated modules are related to the main components of the proteostasis network, we carried out functional enrichment analysis on these modules, based on human gene annotations. The enrichments were highly significant for all modules (FDR < 0.01), and confirmed the inter-modular associations (Table S8). Not all of the modules were related to proteostasis. Interestingly, M1, M10 and M5 in the skeletal muscle network share strong associations with mitochondrion organization and distribution, regulation of cellular amino acid metabolic process and ubiquitin protein catabolic process, while M2 and M3 in hippocampus share associations with different types of protein transport. Other modules (M1, M6, M7, M8, M11, M12 in skeletal muscle; M2, M3, M4, M5, M12 in hippocampus) support the impact of healthy aging on genes related to the proteostasis-linked processes. This included processes related to protein polyubiquitination (GO:0000209), translational initiation (GO:0006413), protein transport (GO:0015031), regulation of macroautophagy (GO:0016241), and proteasome-mediated ubiquitin-dependent protein catabolic process (GO:0043161). In skeletal muscle tissue there were also a strong enrichment in splicing process (M3). Moreover, the connection between M2, M10 and M6 in hippocampus, and between M1, M5 and M12 in skeletal muscle indicates that there is a connection between mitochondrial and proteostasis-related processes, recently shown to occur also in amyloid-beta proteotoxic diseases (Sorrentino *et al*., 2017), and during mitochondrial stress(Labbadia and Morimoto, 2015; D’amico, Sorrentino and Auwerx, 2017; Sorrentino, Menzies and Auwerx, 2018).

To investigate the relevance of proteostasis-linked modules to age-related diseases, we performed enrichment analysis based on genes coming from 22 GWAS studies (See Methods, Table S9). M3, M4, M5 and M12 of skeletal muscle showed enrichment in coronary artery disease, triglycerides, 2hr glucose, multiple sclerosis and cholesterol-related diseases, while M4 and M6 of hippocampus showed enrichment in coronary artery disease and fasting proinsulin, respectively. Skeletal muscle module M12 is particularly interesting because its genes are not only enriched in GWAS studies but also have strong involvement in proteostasis (Figure S10A). Similarly, hippocampus module M4 is interesting due to enrichment in both GWAS and in one of the proteostasis processes (Figure 5B).

**Figure 5.**
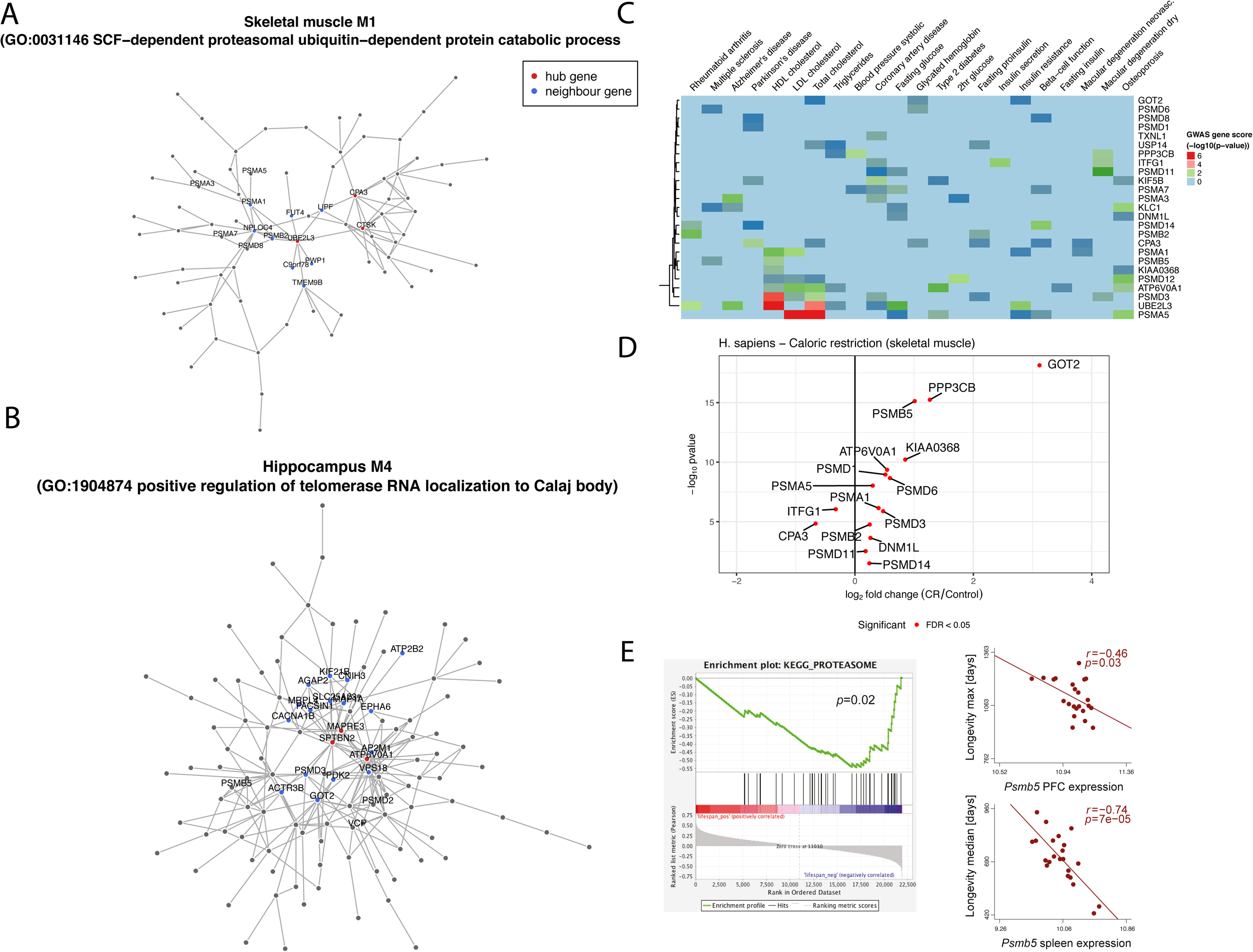
Module architectures and prioritization of candidate genes. (A-B) Architecture of modules related to protein polyubiquitination (M2; A) and positive regulation of telomerase RNA (M4; B) with hub genes (in red) and their neighboring genes (in black), in skeletal muscle (A) and hippocampus (B). (C) GWAS heatmap of the conserved proteostasis-related genes that were prioritized in modules. The heatmap shows the strength of association of each gene (hubs and neighbouring genes from the interested modules) with GWAS. (D) Volcano plot of the prioritized and conserved genes in human dietary restriction dataset. (E) Validation plots for *PSMB5* gene in independent mouse studies, taken at 72 and 78 days of age. The x-axis represents the expression values of the gene in 35 strains of LXS (upper scatterplot) and BXD (lower scatterplot), and y-axis maximum (upper scatterplot) and median (lower scatterplot) lifespan of that strain. The left panel shows GSEA enrichment relation between proteasome complex and lifespan.

To further characterize these modules, we studied how conserved modular genes associated with proteostasis and age-related GWAS diseases are changed in expression in humans, as a long-lived species. We looked deeper into the gene composition of two modules, M1 associated with SCF-dependent proteasomal ubiquitin-dependent protein catabolic process (79 genes) and M4 associated with positive regulation of telomerase RNA localization to Calaj body (155 genes) from the skeletal muscle and hippocampus networks, respectively. We defined network hubs, genes that exhibit a significantly high number of connections with other genes in the network, for each of these modules in muscle (Figure 5A, S10A) and hippocampus (Figure 5B, S10B). We focused on the hubs with the highest scores in each module and examined their neighborhood. The top ranked genes in M1 of the skeletal muscle were *CTSK, UBE2L3 and CPA3* (Figure 5A). They are associated with protein quality network, related to protein degradation. Interestingly, the neighboring genes *PSMB2* and *PSMA1* are associated with the proteasome complex (Figure 5A). The top ranked genes in M4 in skeletal muscle were related to the translational initiation process, with *MAPRE3, SPTBN2* and *ATP6V0A1* as hub genes. Their network neighbors were tightly connected to the cytoskeleton and protein transportation (Figure 5B).

Other modules also show links to metabolism and to proteostasis. For example muscle module M12 and hippocampus module M3 are associated with the protein polyubiquitination process (Figure S10). The top-ranked hub genes in muscle M12 were *DDX3X, KIF5B* and *USP7* (Figure S9A). Those genes are related to DNA damage, translation and transport regulation in the cell. In the hippocampus module M3 (Figure 9B), the three hub genes (*PPP3CB, DNM1L* and *ITFG1*) are involved in hydrolase activity, apoptosis and programmed necrosis and modulating T-cell function. Although the hub genes with the highest scores were strongly related to metabolism and to tissue-specific functions in each of these two modules, their network neighborhood is associated with the protein quality control network. More specifically, the *PSMB5* and *PSMD3* genes are related to the proteasome complex and are connected to hub genes.

We combined this hub gene analysis with GWAS association gene scores, and observed that *PSMB5, UBE2L3*, and *PSMD3* (Figure 5C, Table S9) are important in many age-related diseases or phenotypes, such as Alzheimer’s disease, HDL cholesterol, LDL cholesterol, triglycerides, and insulin resistance. Other genes related to translation and proteasome complex were also strongly associated to such diseases, such as *PSMB5* with multiple sclerosis (Pascal (Lamparter *et al*., 2016) gene score: *p-value* = 0.0348) and HDL cholesterol (Pascal gene score: *p-value* = 0.0155). Finally, we observed that the prioritized genes associated with age-related diseases from conserved functional modules change in opposite directions with healthy aging and with caloric restriction (Figure 5D). This differential expression is consistent with a causal role in these age related diseases, given the attenuating effect of caloric restriction on aging.

### Validation of marker genes using independent mouse studies

We analyzed the association of the expression levels of candidate genes with lifespan in different tissues of mouse recombinant inbred lines used for population genetics analyses, such as the BXD (Andreux *et al*., 2012) and LXS (Liao *et al*., 2010) strains. We observed an inverse correlation between transcript levels of *PSMB5* (Figure 5E) in the spleen of the BXD strains (average age at the time of transcript analysis 78 days; p = 7.14 × 10^−5^) and in the prefrontal cortex of LXS lines (average age of 72-days; p = 0.03), and lifespan longevity. This correlation was consistent even after correction for the population structures with mixed models (Kang *et al*., 2008). Thus lower expression of PSMB5 is linked to lifespan. Consistent with this, the GSEA showed down-regulation of the proteasome complex during the lifespan of the mice (Figure 5E, left panel).

## Discussion

The challenge of detecting underlying mechanisms of healthy aging that are evolutionarily conserved is thought to be a key impediment for understanding human aging biology (Fontana *et al*., 2010). In this work, we coupled evolutionary and functional information of healthy aging gene expression studies to identify conserved age-related systems-level changes. We identified conserved functional modules by integration of co-expression networks, and we prioritized genes highlighted by GWAS of age-related diseases and traits. The observations on several functional levels allowed us to highlight the role of proteostasis, which includes all processes related to protein quality control network, as a strong core process associated with normal aging.

Previous observations restricted to a small number of evolutionarily conserved genes with large effects in aging, or in age-related diseases, provided some evidence that aging mechanisms might be conserved among animals (de Magalhães, Curado and Church, 2009). However, transcriptome level correlations of expression changes in aging between species are very low in our gene-level results, in accordance with other studies (Zahn *et al*., 2006; Smith *et al*., 2008; Alexey A Fushan *et al*., 2015). Yet the process of aging appears overall conserved, with notably common effects of interventions, such as caloric restriction, showing similar effects across species ranging from nematodes, flies, to mammals (Gems and Partridge, 2013). The solution to this apparent paradox seems to be that pathways are evolutionarily conserved in aging (Smith *et al*., 2008), even when single genes are not. Indeed, we have found strong similarities in age-related gene sets between human and other species.

Beyond individual pathways, the modular nature of aging has been previously reported at several levels, such as by protein-protein interaction network analysis during human and fruit fly brain aging (Xue *et al*., 2007), human longevity network construction and identifying modules (Budovsky *et al*., 2006), mouse age-related gene co-expression modules identification (Southworth, Owen and Kim, 2009), or aging and age-related diseases cluster detection in human aging (Fernandes *et al*., 2016). Integrating co-expression networks across species, we identified 10 and 13 evolutionarily conserved functional modules for skeletal muscle and hippocampus, respectively. These conserved modules are not only enriched in processes known to be involved in healthy aging, such as immune-related pathways, they significantly overlap with results from age-related GWASs. The latter is of particular relevance, since finding causality for aging in GWAS is difficult, given its highly multifactorial nature (McDaid *et al*., 2017). Of note, these modules can be tissue-specific, for example related to energy and amino acids in muscle (Figure 5A). Thus, aging is an evolutionarily conserved modular process, and this modularity is tissue-specific.

An advantage of our approach is that it allows us to detect with good confidence processes whose changes in aging are quite subtle. This is important because healthy aging is not a dramatic process, akin to embryonic development or cancer, but a gradual change in tissues and cell types which keep their defining characteristics. In other words, old muscle and young muscle are very similar at the molecular level, as shown, e.g., by the log-fold change scale in Fig. 3: a log2 age-related regression coefficient (Formula 1) of −0.005 corresponds to a decrease of only 1.0035 fold. Yet we are able to detect processes associated to these changes with strong confidence, and these processes are mostly known in to be age-related. The largest changes, thus easiest to detect, include metabolism (Finkel, 2015), transcription (Roy *et al*., 2002), translation (Steffen and Dillin, 2016), and immune response. Changes in expression for proteostasis-related genes are weaker, yet integrating at a systems level between species provided us with a strong signal.

More broadly, our results strengthen the case for further investigation into the molecular program that links proteostasis to healthy aging. This is in line with “loss of proteostasis” as one of the nine proposed hallmarks of aging (López-Otín *et al*., 2013; Walther *et al*., 2015). Aging involves a deregulation of the protein quality control network, and this is conserved between distant species. Changes in protein synthesis and protein degradation processes have already been linked to several age-related diseases, most notably Alzheimer’s and Parkinson’s disease (Morimoto and Cuervo, 2014). They may be fundamental to the response to normal aging because the accumulation of somatic and germline mutations can alter fine modulation of the protein homeostasis network and produce pathological alterations (Woodruff and Thompson, 2003; Khodakarami *et al*., 2015). Thus proteostasis provides a link between somatic genome-level changes and the phenotypic impact of aging. Our results show that during healthy or normal aging, the alterations in proteostasis network are rather subtle and discrete, by contrast to the strong down-regulation of metabolic processes. This suggests that perhaps there is a cascade of triggered pathways as aging proceeds (Tomaru *et al*., 2012). Moreover, we detect evolutionarily conserved links inside modules between mitochondrial deregulation (hub genes) and protein homeostasis (neighboring genes) in normal aging, consistent with recent advances in the field (D’amico, Sorrentino and Auwerx, 2017; Labbadia *et al*., 2017; Sorrentino, Menzies and Auwerx, 2018).

The main evolutionarily conserved gene candidates from proteostasis, *PSMB5* and *PSMD3*, are related to the proteasome. These two genes were tightly connected to metabolic hub genes in skeletal muscle and to filament organization genes in the hippocampus. The proteasome complex is down-regulated during aging in our results, and in a transgenic mouse mutant proteasome dysfunction led to shorter lifespan (Schmidt and Finley, 2014). In the database of gene expression Bgee (Bastian *et al*., 2008), human PSMB5 and PSMD3 have top expression in gastrocnemius muscle, with weaker expression in old age. Moreover, both genes showed significant association in GWAS studies with metabolic and disease traits. The *PSMB5* gene was validated by comparing mice strains, and the *PSMD3* gene was related with coronary artery disease, HDL cholesterol and fasting proinsulin, all indicators of healthspan, and would also be worthwhile to explore further.

The association with caloric restriction studies strengthens the functional contribution to aging of the processes we identified. We observed that the gene-set signals were both evolutionarily conserved in caloric restriction, and shared between healthy aging and caloric restriction experiments. Genes related to proteostasis showed opposite directions in expression changes between human healthy aging and caloric restriction. This indicates that these functions are maintained during caloric restriction in humans but lost during aging, and reinforces the case for a causal link between proteostasis and healthy aging. Our observations are consistent with previous research in *C. elegans*, reporting improvement of proteostasis during caloric restriction treatments and extension of the lifespan (Depuydt *et al*., 2013; Chondrogianni *et al*., 2015). Notably, *PSMB5* and *PSMD3* follow this trend in caloric restriction relative to healthy aging, further suggesting that they are prime candidates to study genes underlying functional modules in healthy aging.

Integrating biological processes based on evolutionary conservation allows distinguishing relevant signals from noise, despite the weak patterns in aging transcriptomes. Moreover, the fact that a process is similarly involved in aging in very different species strengthens the case for causality. This provides a promising foundation to search for relevant biomarkers of healthy aging of specific tissues, e.g. further analysis of directions of change in homologous tissues, in different model organisms.

In summary, the large-scale, comprehensive gene expression characterization in our study provides insights in underlying evolutionarily conserved mechanisms in normal aging. While metabolic and certain tissue-specific pathways play a crucial role in aging, processes affecting the protein quality control network also show very consistent signal. Using both evolutionary and functional information, we detected conserved functional modules that allowed us to identify core proteostasis-related genes. These genes were implicated as important hits in age-related GWAS studies (Gomes, 2013). Together, the integrative systems-level approach facilitated the identification of conserved modularity of aging, and of candidate genes for future normal aging biomarkers.

## Supplemetal Data

### Supplement Figures are in Supplemental document

**Table S1. Expression datasets used in aging and caloric restriction analysis**. This table contains 2 sheets, corresponding to aging and dietary restriction experiments.

**Table S2. Differential expression statistics in skeletal muscle (human, mouse), hippocampus (human, mouse), whole body (fly, worm) for age-related experiments and skeletal muscle (human, mouse) and whole body (fly, worm) for dietary restriction**. This table contains 6 sheets, each sheet corresponds for tissue and species. In each sheet, rows correspond to genes with no cutoffs applied. The columns provide differential expression statistics for all the samples (GTEx) and two-group comparisons (model organisms).

**Table S3. Overlap between the 1-to-1 conserved age-related orthologs between human and model organisms**.

**Table S4. List of orthologous genes from integrative analysis**. This table contains 3 sheets, corresponding to muscle, hippocampus and dietary restriction experiments that were integrated based on orthologous groups. The columns represent name of orthogroups, combined p-values across species from Fisher’s combined probability test, original p-values from differential expression analysis per species and annotations of genes. The rows contain genes that are representative per orthologous group for each species.

**Table S5. Summarized clusters based on GO semantic similarity method**. This table contains 3 sheets, corresponding to muscle, hippocampus and dietary restriction GO analysis. The file shows the GO enrichments and categorization to higher (more general) GO terms.

**Table S6. Proteostasis-linked processes enriched in 2 tissues and dietary restriction experiments**. This table contains 3 sheets, corresponding to muscle, hippocampus and dietary restriction GO analysis for proteostasis-linked processes.

**Table S7. Significant conserved genes from human GTEx in proteostasis quality network for skeletal muscle and hippocampus**. This table contains 6 sheets for each part of the protein quality network (macroautophagy, translation and proteasome complex) per tissue.

**Table S8. Summary of the statistics from network analysis**. This table contains 5 sheets of the information about the sizes of the all modules and GO and GWAS enrichments in each tissue for proteostasis-linked modules.

**Table S9. Summary of mapping the GWAS traits for selected modules**. This table contains the gene-level p-values from the PASCAL tool for the heatmap of Figure 5C for selected 22 GWAS age-related studies.

## METHODS

### Data selection

To obtain a representative set of aging gene expression experiments, a set of raw RNA-seq and microarray datasets of four species (*H. sapiens, M. musculus, D. melanogaster, C. elegans*) were downloaded from the GEO database (Barrett *et al*., 2013) and SRA database (Leinonen *et al*., 2011) (Table S1). For observing aging gene expression signatures in human and mouse, we selected hippocampus and skeletal muscle tissues. The aging gene expression experiments for fly and worm were available as whole-body experiments. All the healthy or control samples came from two extreme age groups (young and old adults) that are counted from sexual maturity. This corresponds to 20-30 years old humans, 3-4 months old mice, 4-5 days old flies and 3-6 days old worms (see Figure 1B) in young adults. In old adult age group, this corresponds to 60-70 years old humans, 20-24 months old mice, 40-50 days old flies and 12-14 days old worms. The sample size per age group was 3-6 replicates. The GTEx V6p read counts were used as *H. sapiens* aging experiment (V6p dbGaP accession phs000424.v6.p1, release date: October, 2016). The information about the sample ages was obtained through dbGAP annotation files of the GTEx project (restricted access). Two RNA-seq datasets were matched for *M. musculus* and *C. elegans*; and the microarray platforms included were from Affymetrix: Mouse 430 A/2.0, GeneChip Drosophila Genome array and *C. elegans* Genome array.

### GTEx v6p analysis

From the downloaded GTEx V6p data, we extracted the gene read counts values for protein-coding genes by using Ensembl (release 91). For each tissue, the lowly expressed genes were excluded from data analysis according to the GTEx pipeline (Mele *et al*., 2015). Prior to the age-related differential expression analysis, we used the PEER algorithm (Stegle *et al*., 2012) in a two-step approach to account for known covariates as well as for hidden factors present in GTEx V6p data per tissue. From covariate files (Brain_Hippocampus_Analysis.covariates.txt and Muscle_Skeletal_Analysis.covariates.txt), we used information about the three genotype principal components. From phenotype file (phs000424.v6.pht002742.v6.p1.c1.GTEx_Subject_Phenotypes.GRU.txt), we used information about age, gender, ischemic time and BMI information. From attribute file (phs000424.v6.pht002743.v6.p1.c1.GTEx_Sample_Attributes.GRU.txt), we extracted information about the sample associations with interested tissues, hippocampus and skeletal muscle. In the first step, the PEER algorithm discovers patterns of common variation; it created 15 and 35 assumed global hidden factors for hippocampus and skeletal muscle, respectively. In addition to global hidden factors, we provided age, BMI, sex and ischemic time as known covariates in PEER model. In the second step those hidden factors (gene expression principal components) that showed significant Pearson’s correlation coefficient with age (p-value < 0.05) were excluded. The number of hidden factors that did not significantly correlate in hippocampus was 7/15 and in skeletal muscle were 22/35 that were selected for further linear model analysis. The sum of remaining hidden factors and known covariates were included in a linear regression model to obtain the genes differentially expressed during age in GTEx V6p data for each tissue (Formula 1).

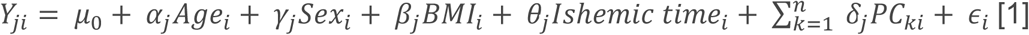

where, Y_ji_ is the expression of a gene *j* in a sample *i*, where *Age, Sex, BMI, Ischemic time* of sample *i*, with their regression coefficients *α, γ, β, θ. PC*_*ki*_ (1 < k < *n*) is the value of the *k*-hidden factors for the *i*-th sample with regression coefficient *δ*; *n* is a total number of factors that was not correlated with age, *ε*_*i*_ is the error term, and *µ*_*0*_ is the regression intercept. If *α* > 0, gene *j* was treated as up-regulated, otherwise gene *j* was treated as down-regulated. The linear model (Formula 1) was performed in *limma voom*, and the p-values were corrected for multiple testing by performing false discovery rate (FDR) correction using Benjamini-Hochberg method.

### Aging datasets microarray analysis

For microarray datasets (both aging and caloric restriction experiments) from skeletal muscle of *M. musculus* and whole-body of *D. melanogaster*, raw Affymetrix .CEL files were downloaded from the GEO database and preprocessed using RMA normalization algorithm (Irizarry *et al*., 2003) (Table S1). In case of multiple probes mapping to the genes on the array, the average of the probes was taken in further analysis. The annotation was used from Ensembl release 91. In order to identify the features that exhibit the most variation in the dataset, principal component analysis (PCA) was performed on the expression matrices to detect outlier samples, gender and other batches.

### Aging datasets RNA-seq analysis

For RNA-seq datasets from two model organisms, *M. musculus* and *C. elegans*, the .sra files were downloaded from the SRA database (Leinonen *et al*., 2011). Both datasets were sequenced on Illumina HiSeq 2000 with read length 50nt. The reads were mapped to species-specific reference genomes (*M. musculus*: GRCm38.p5, *C. elegans*: WBCel235) using kallisto v0.43.1 (for index building: kallisto index –i genome.idx genome.cdna.all.fa (k-mer = 31, default option); for mapping: kallisto quant -i genome.idx –o output.file –single –l 200 –s 20 single.end.fastq.file) (Bray *et al*., 2016). Both *M. musculus* and *C. elegans* had single-end RNA-seq libraries in the experiments (Table S1.). The transcript abundances were summarized at the gene-level (Soneson, Love and Robinson, 2015). For both species, we used GTF gene annotation files that were downloaded from Ensembl ftp site (release 91) (Aken *et al*., 2016). The transcript abundances were summarized at the gene-level to lengthscaledTPMs using tximport v1.6.0 (Soneson, Love and Robinson, 2015) and used as an input to *limma voom*. The gene-level read counts were further analyzed in R v3.4.3. The read counts were normalized by total number of all mappable reads (library size) for each gene. The *limma voom* results in a matrix of normalized gene expression values on log2 scale. The counts and normalized log2 *limma voom* expression values were used as a raw input for all the analysis. Outlier samples were checked by principal component analysis. For each species, genes that showed expression below 1 count per million (cpm < 1) in the group of replicates were excluded from downstream analysis.

### Identification of age-related differentially expressed genes

To be able to obtain differentially expressed genes from different experiments that were normalized, we had to account for the possible batches present. Since we are not aware of all the batches in the studies, we used Surrogate Variable Analysis (SVA) to correct for batches (Leek and Storey, 2007) in microarray data analysis. The SVA method borrows the information across gene expression levels to estimate the large-scale effects of all factors absent from the model directly from the data. After species-specific expression matrices were corrected, they served as input into linear model analysis implemented in *limma* (Affymetrix) or *limma voom* (RNA-seq) (Law *et al*., 2014), for finding age-related differentially expressed genes between two extreme aging groups, young and old. Briefly, *limma* uses moderate t-statistics that includes moderated standard errors across genes, therefore effectively borrowing strength from other genes to obtain the inference about each gene. The statistical significance of putatively age-dependent genes was determined with a false discovery rate (FDR) of 10%.

### Caloric restriction datasets microarray analysis

The GEO database was used to download caloric restriction datasets (Table S1). Only muscle tissue was available in *H. sapiens*, therefore we selected correspondingly muscle tissue in mouse, but whole body in fly and worm. The datasets were normalized using RMA normalization algorithm (Irizarry *et al*., 2003) (Table S1). In case of multiple probes mapping to the genes on the array, the average of the probes was taken in further analysis. The annotation was used from Ensembl release 91. To call differentially expressed genes, we used *limma* between caloric restriction and control samples. The statistical significance of putatively age-dependent genes was determined with a false discovery rate (FDR) of 5%.

### Age group alignments between species

For deriving one-to-one orthologs, human genes were mapped to the homologs in the respective species using biomaRt v2.34.2. After detection of significant age-associated differentially expressed genes, we overlapped one-to-one orthologous genes between the species in order to observe the consistency of age groups between species. We took the *limma voom* corrected expression matrix for GTEx V6p and the expression matrices of model organisms, and selected only genes that were differentially expressed with an FDR of 5%. We then accounted for the laboratory batch effect by applying Combat on expression matrices (Leek *et al*., 2012).

### Gene-level analysis

To examine the relationship between aging in human and model organisms on single-gene level, we mapped one-to-one orthologous genes from human to model organisms and between the organisms downloaded from Ensembl (Aken *et al*., 2016). We calculated Spearman correlations between sets of matched differentially expressed orthologous genes, between log2 fold-changes (Supplementary Figure S2). No cutoff for fold change was used.

### Constructing homologous quadruplets and enrichment analysis

We downloaded hierarchical orthologous groups (HOGs, in further text referring to orthologous groups (OG)) across four species from the OMA (orthologous matrix analysis) database (Altenhoff *et al*., 2015) at the Bilatera level (*Amphimedon queenslandica* (*Cnidaria*) was used as a metazoan outgroup), which resulted in 3232 orthologous groups. Briefly, hierarchical orthologous groups are gene families that contain orthologs (genes related by speciation) and in-paralogs (genes related by duplication) at the taxonomic level which orthologous groups were defined. The sizes of orthologous groups in this study range from 4 to 246 genes. We filtered age-related genes per orthologous group per species in order to obtain representative species-specific genes per group. The genes within orthologous group were selected according to the *P* values from differentially expression analysis (Rittschof *et al*., 2014). We applied Bonferroni correction on each orthologous group to the differential expression *P* values in order to correct for the size of the orthologous group. We then combined the corrected differential gene expression *P* values across species using Fisher’s combined probability test generating a new *P* value from χ^2^ distribution with 2k degrees of freedom (Formula 2).

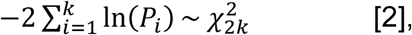

where *P*_*i*_ is species-specific gene *P* value from differential expression analysis within a OG.

We adjusted combined Fisher *P* values for multiple testing, and filtered orthologous groups with FDR of 10% for further analysis. This resulted in 2010 and 2075 common OGs for skeletal muscle and hippocampus, respectively. In caloric restriction experiments, we detected 1962 common OGs.

We performed general GO enrichment analysis using Fisher’s test (topGO R package) on significant orthologous group genes and based on human gene set annotation to find functional enrichment of OGs in GO ‘biological process’ terms. To summarize the significantly enriched top 100 GO terms into main ones, we used the Wang GO semantic similarity method (Wang *et al*., 2007) that takes into account the hierarchy of gene ontology, and performed hierarchical clustering (11 clusters for skeletal muscle and 13 clusters for hippocampus, 10 clusters for caloric restriction) on the semantic matrix for both aging and caloric restriction experiments (Table S5). The clusters were then named according to the common term of the cluster. We associated proteostasis-linked processes to GO terms associated with ‘translation’, ‘protein folding’, ‘proteasome assembly’, ‘macroautophagy’, ‘proteasome complex’, ‘endoplasmic reticulum’, ‘lysosome’ and others.

To perform the randomizations, we selected random genes from the differential expression matrices with the same number as the number of orthologous groups selected for skeletal muscle and hippocampus. The p-values associated with the random genes per species were then combined with the Fisher’s combined test. The GO enrichment analysis was performed as for the observed data with focus on the ‘biological process’ and based on the human annotation. The procedure was repeated 100 times (Figure S6).

### Prioritization of OG gene pairs in multi-species co-expression network

We aimed to detect gene sets that are perturbed in aging in different species. We selected the genes from previously formed significant age-related OGs per species and constructed the species-specific co-expression networks by calculating Pearson correlation coefficient between age-related OGs genes. In the resulting species-specific co-expression network, nodes represent genes and edges connect genes that are above a set significant threshold from Pearson correlation calculation (*P* value < 0.05). Only positively correlated genes were taken into account, while the negatively correlated genes and genes correlating under the threshold were set to zero. Negatively correlated genes might be interesting to detect complex regulatory patterns, but are beyond the scope of this study. The cross-species network was obtained as follows (Stuart *et al*., 2003). Each co-expression link was assigned a rank within the species according to the Pearson correlation value. We then divided the species-specific ranks by the total number of OGs per tissue to normalize the ranks across the species (Formula 3, example for human, but same for other species).

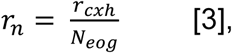

where *r*_*n*_ is normalized gene pair rank, *r*_*cxh*_ is the rank of coexpression link in human and *N*_*eog*_ is the number of common evolutionary orthologous groups selected for tissue.

The final gene-pair list was then obtained by integrating human, mouse, fly and worm ranked lists using robust aggregation, originally made for comparing two lists (Kolde *et al*., 2012). Briefly, using beta probability distribution on order statistics, we asked how probable is the co-expression link by taking into account the ranks of all four species. This method assigns a *P* value to each co-expression link in an aggregated list, indicating how much better it is ranked compared to the null model (random ordering). This yielded networks with 2887 and 3353 significant gene-pairs (edges) (*P* value < 0.001) for skeletal muscle and hippocampus, respectively.

To confirm that the integrated age-related multi-species networks are significant, we selected randomly collected genes from each species. The numbers of selected genes was the same as in the OGs. We then formed the quadruplets and performed the same integration analysis as before. We repeated the procedure 100 times, and obtained 100 randomly integrated multi-species networks (Figure S7). In both cases, random and original analysis, the annotation was based on human.

### Clustering the integrated cross-species network

In order to identify aging-associated functional modules, we created networks containing 1142 nodes (2887 edges) in skeletal muscle and 1098 nodes (3353 edges) in hippocampus, from our prioritized gene pair list based on orthology and all edges between them. The negative logarithm (base 10) of *P* values from aggregated list was assigned as edge weights in both integrated networks. We decomposed the skeletal muscle and hippocampus integrated networks into components and the further analysis was restricted to analysis of a giant component. The giant component contained 1050 genes (nodes) in skeletal muscle and 1067 genes (nodes) in hippocampus. As before, we used human annotation. The modules within the cross-species networks of each tissue were obtained by using a multilevel community algorithm that takes into account edge weights (Yang, Algesheimer and Tessone, 2016) from igraph (Csárdi & Nepusz 2006). Briefly, the multilevel algorithm (Blondel *et al*., 2008) takes into account each node as its own and assigns it to the community with which it achieves the highest contribution to modularity. To obtain Figure 4, we summarized groups of module nodes to single meta-nodes according to their multilevel-algorithm calculated module membership, and showed the inter-modular connectivity using a circular layout. We selected the modules with size greater than 10, which returned 12 modules per tissue-specific cross-species network. We checked the functional enrichment of genes within selected modules in every network using Gene Ontology through topGO R package (See Figure 4).

Moreover, we downloaded the pre-calculated file of gene-level summary statistics from 37 GWASs from the Pascal method (Lamparter *et al*., 2016). We selected 22 out of 37 GWAS studies (Marbach *et al*., 2016) (Table S9) that are associated with metabolic and neurological age-related diseases. To perform enrichment of the module genes within GWAS age-related diseases categories, we selected top-ranking genes (GWAS gene score < 0.1) within each disease and formed the categories for enrichment. We ran enrichment analysis on final network modules to find disease-related modules (adjusted p-value < 0.2). The human genome was used as a background gene set.

Finally, we used Kleinberg’s hub centrality score to determine the hub genes within interested modules and observed the hub-gene neighborhood. The final genes were then selected to show their *P* value association within GWAS studies (Figure 5C, Table S9).

### LXS and BXD mouse data

Male and female mice from those strains were fed with normal ad libitum diet, and median and maximum lifespan were calculated to represent longevity across strains. Microarray data as well as lifespan data were downloaded from GeneNetwork.org. Microarray data from prefrontal cortex of LXS mice was generated by Dr. Michael Miles using animals with the average age of 72 days (GN Accession: GN130). Microarray data from spleen of BXD mice was generated by Dr. Robert W. Williams using animals with the average age of 78 days (GN Accession: GN283). Microarray data from hippocampus of BXD mice was generated by Dr. Gerd Kempermann and Dr. Robert W. Williams using animals with the average age of 70 days (GN Accession: GN110). To correct for the population structure within the strains, a linear mixed model approach was applied. For enrichment analysis, genes were ranked based on their Pearson correlation coefficients with the lifespan data of the BXD strains, and Gene Set Enrichment Analysis (GSEA) was performed to find the enriched gene sets correlated with the lifespan (Subramanian et al., 2005).

## Author contributions

Conceptualization, A.K. and M.R.R.; Methodology, A.K.; Investigation, A.K.; Preprocessing datasets: A.K, Formal Analysis, A.K.; Validation analysis: H.L.; Writing – Original Draft, A.K.; Writing – Review & Editing, A.K., H.L., V.S., J.A., Z.K. and M.R.R.; Funding Acquisition, M.R.R.; Supervision, M.R.R.

The authors declare that they have no conflict of interest.

## Acknowledgements

We thank Natasha Glover for help with the OMA database. We thank AgingX collaborators, Matthias Morf and all members of the Robinson-Rechavi group for helpful comments. We thank Thomas Flatt for critical reading of the manuscript. We are grateful to the teams that made their data available through Gene expression omnibus (GEO) and the database of Genotypes and Phenotypes (dbGAP) repositories. The computations were performed at the Vital-IT (http://www.vital-it.ch) center for high-performance computing of the SIB Swiss Institute of Bioinformatics. This work was supported by the AgingX project from SystemsX.ch, the EPFL, the Velux Stiftung (J.A.), and the NIH (R01AG043930).

