## Supplementary Materials for "Cross-species functional modules link proteostasis to human normal aging"

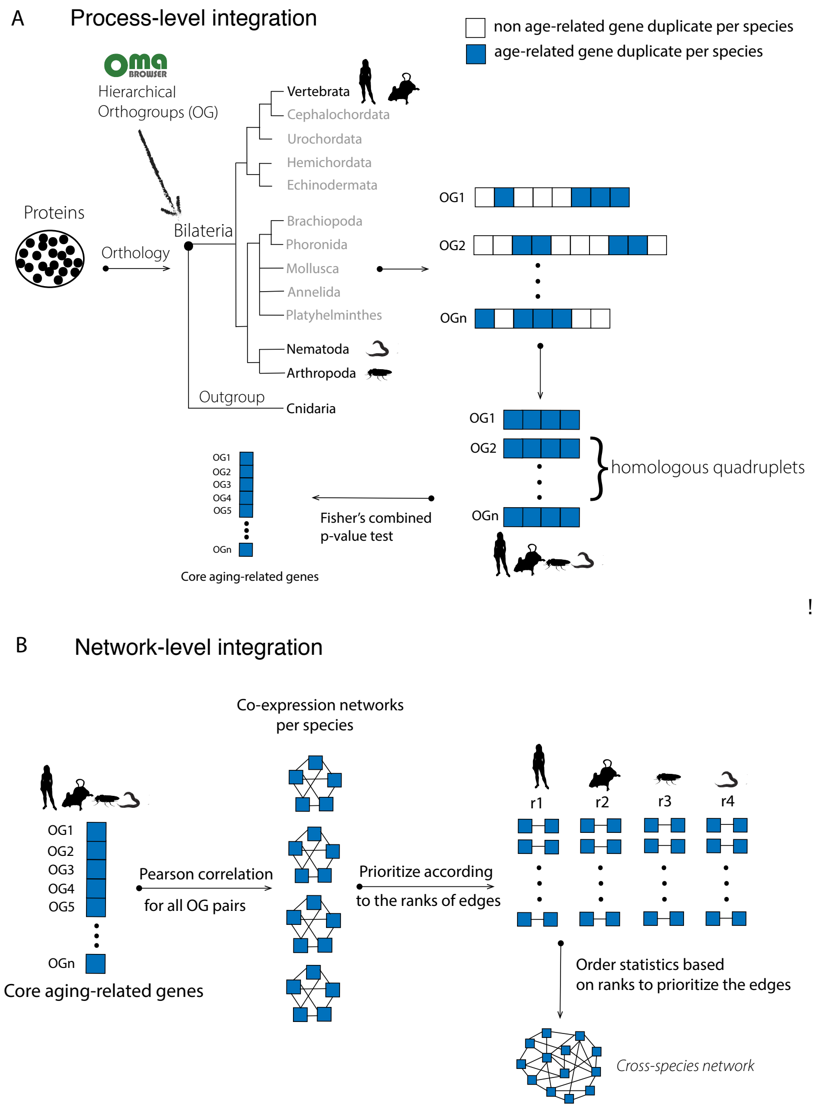


**Figure S1. Detailed overview on the statistical integrative cross-species approach.**
**(A)** Process-level integration. The integration is done based on the selection of the gene set families conserved across 4 species; minimum p-values from age-related differential expression analysis are used to define "age-related" genes. The p-values are combined using Fisher's combined test. **(B)** Network-level integration. The obtained age-related conserved genes were used for the integration of gene co-expression networks across species based on n-order statistics.

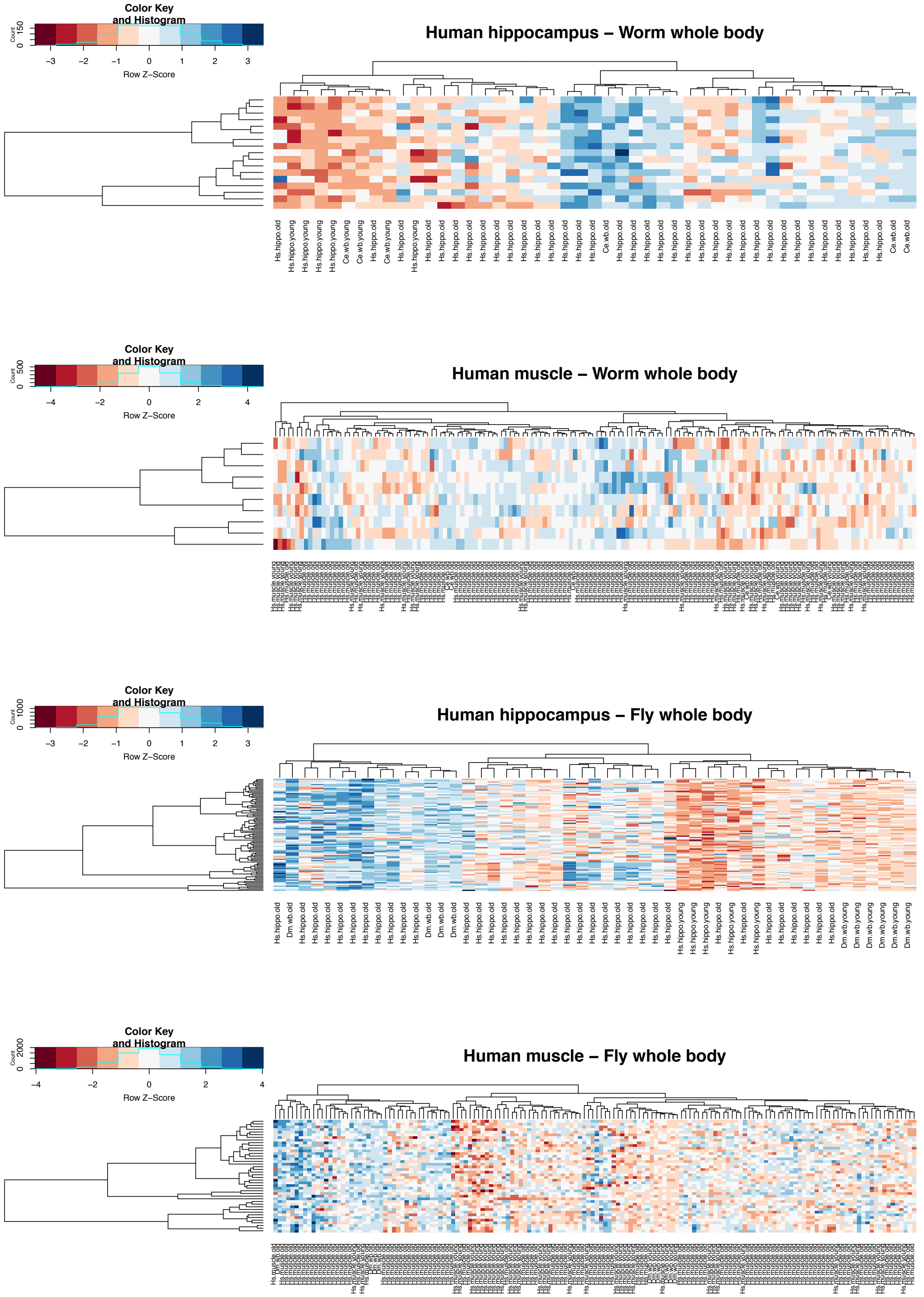


**Figure S2. Clustering of the age-related samples between human (20-30y; 61-70y) and other species.** The selected top differentially expressed genes that are orthologous between each species show alignments between the young and old samples.

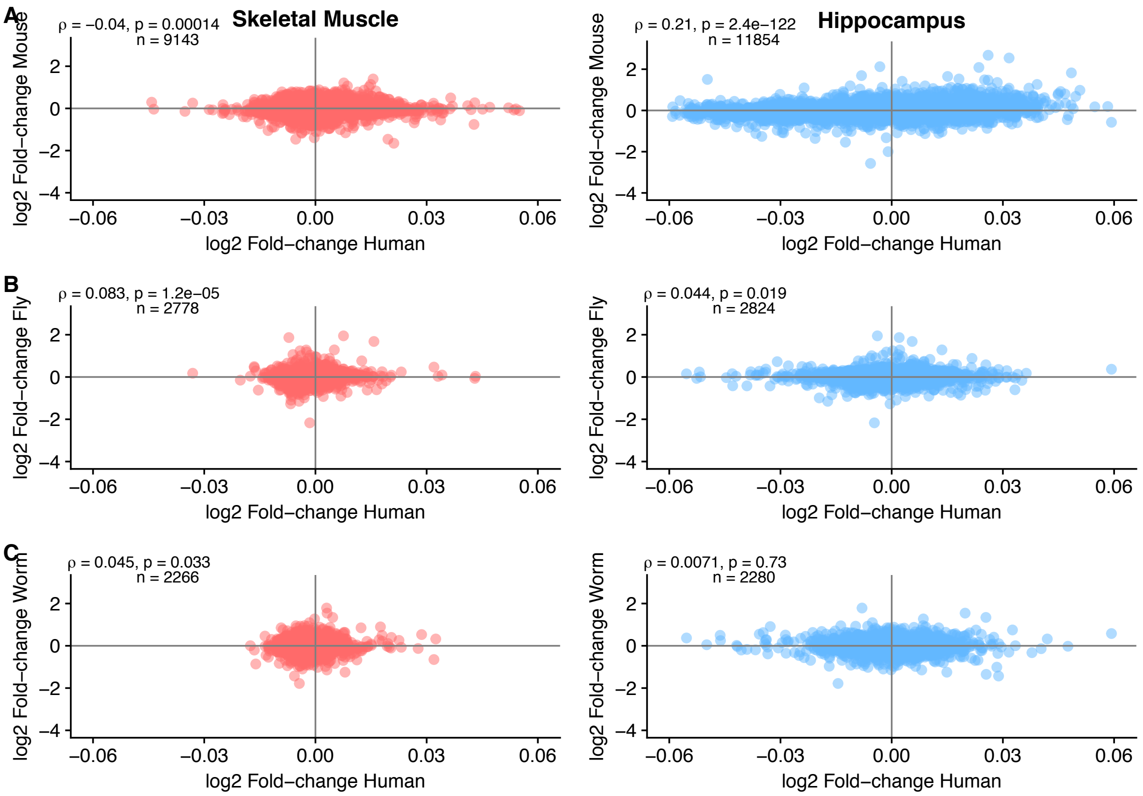


**Figure S3. The scatterplots of pairwise species comparison from single-gene level analysis. (A) Human-Mouse, (B) Human-Fly, (C) Human-Worm**. No cut-off was applied. There is a weak correlation between the 1-1 orthologous genes between human and other species. This indicates that the gene-level changes in aging are species-specific.

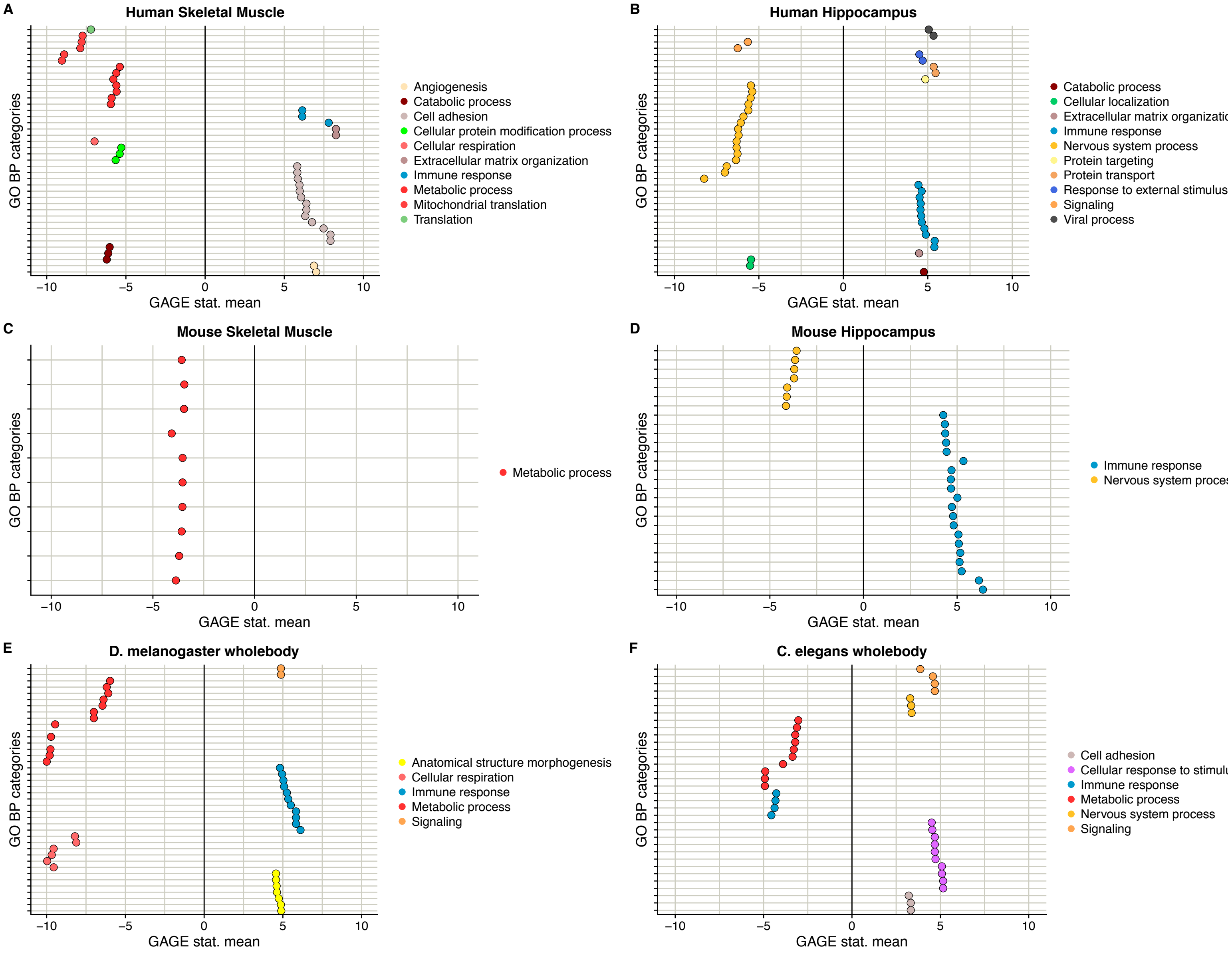


**Figure S4. GSEA per single-species. The panels (A-F) show the enrichments in GO BP categories (FDR < 0.20) in normal aging per species.** The GSEA plots show strong enrichment in tissue-specific processes that are perturbed during aging process.

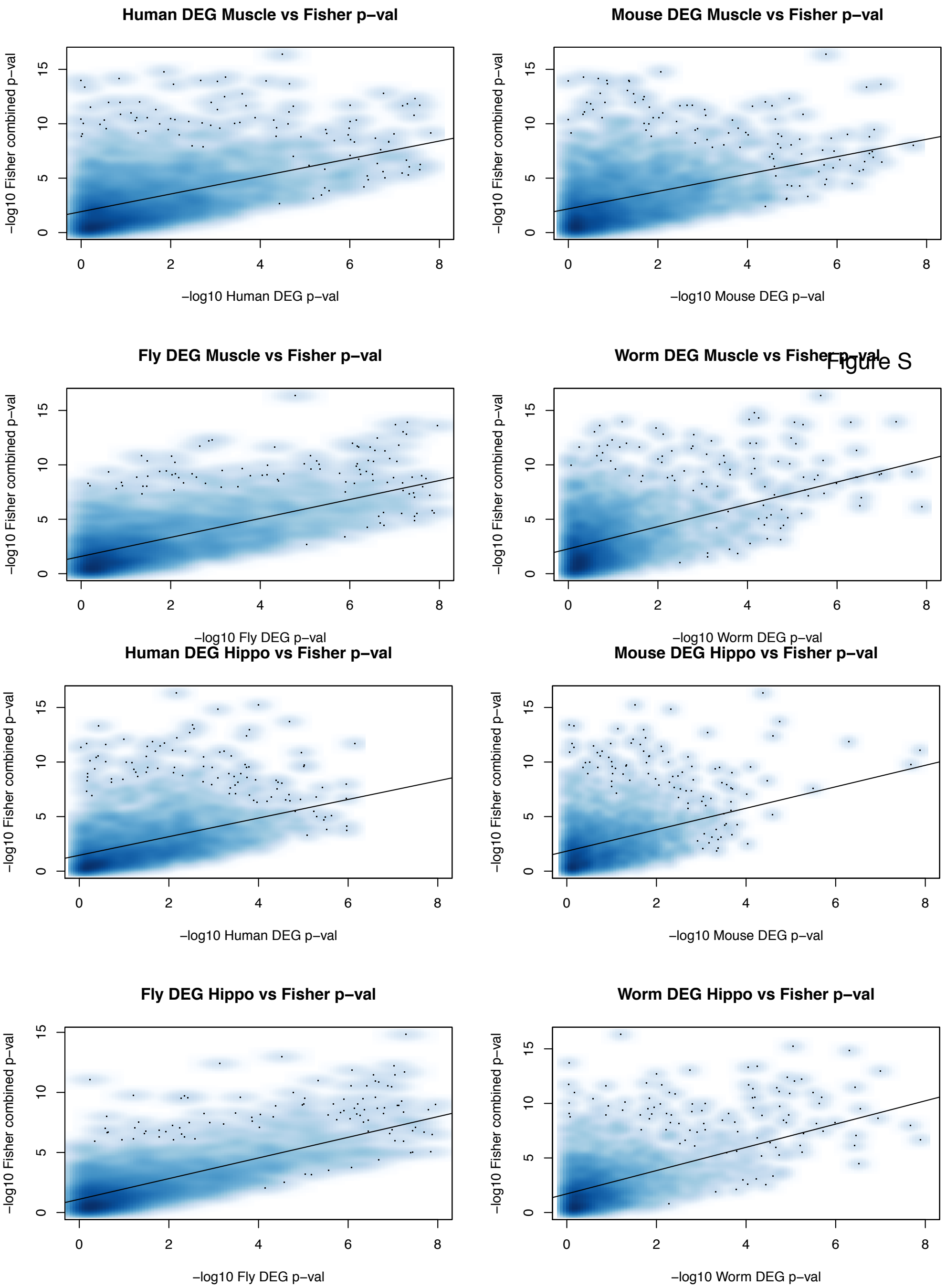


**Figure S5. The scatterplots of p-values from single-species differential expression analysis, against the p-values integrated using Fisher's combined test.** The Fisher's method gives more conservative than classical (per species), meaning that some genes found differentially expressed in species when combined are more significant.

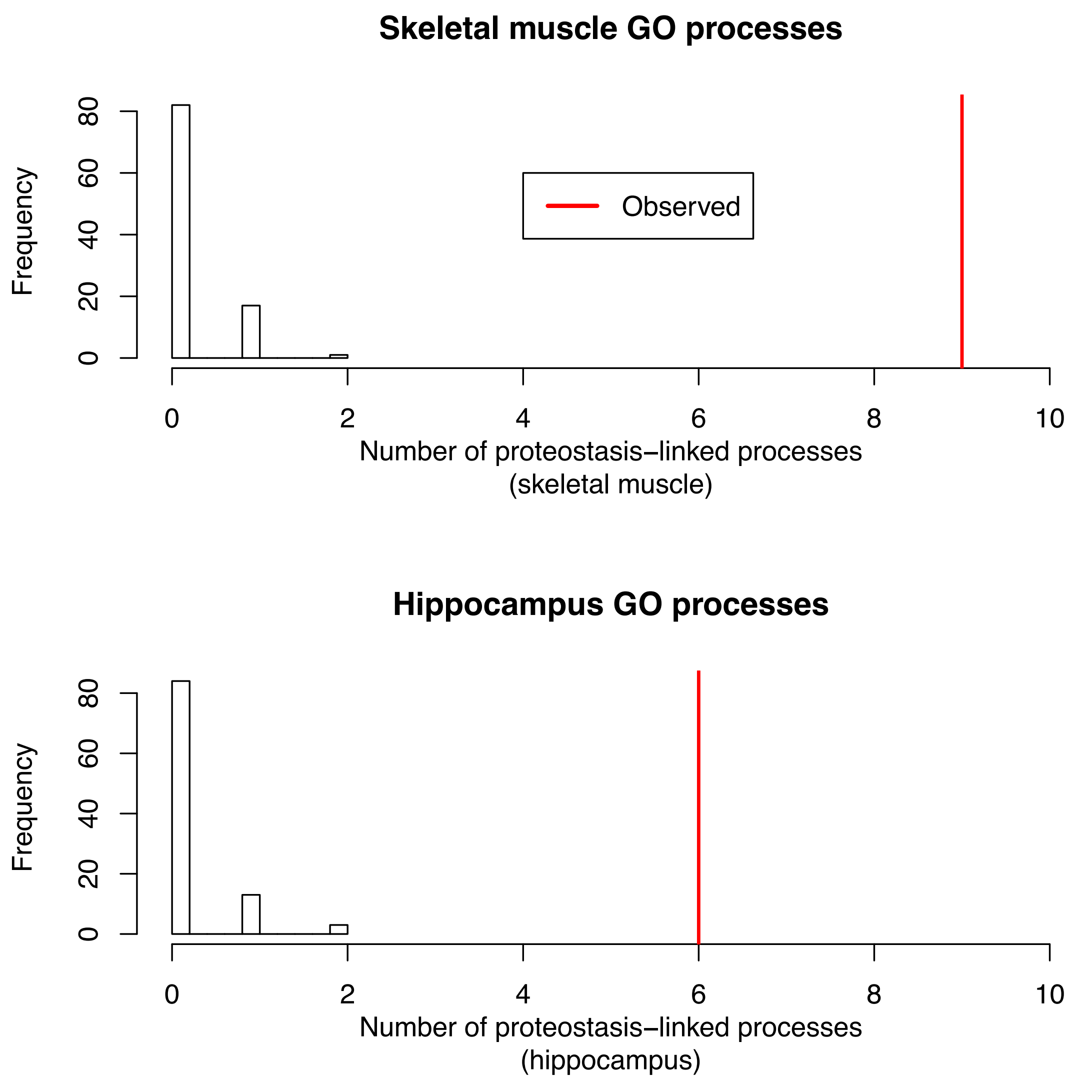


**Figure S6. Randomization on the processes-level results (100 permutations) in both skeletal muscle and hippocampus data.** We observe that the proteostasis-linked processes are appearing more than expected by chance.

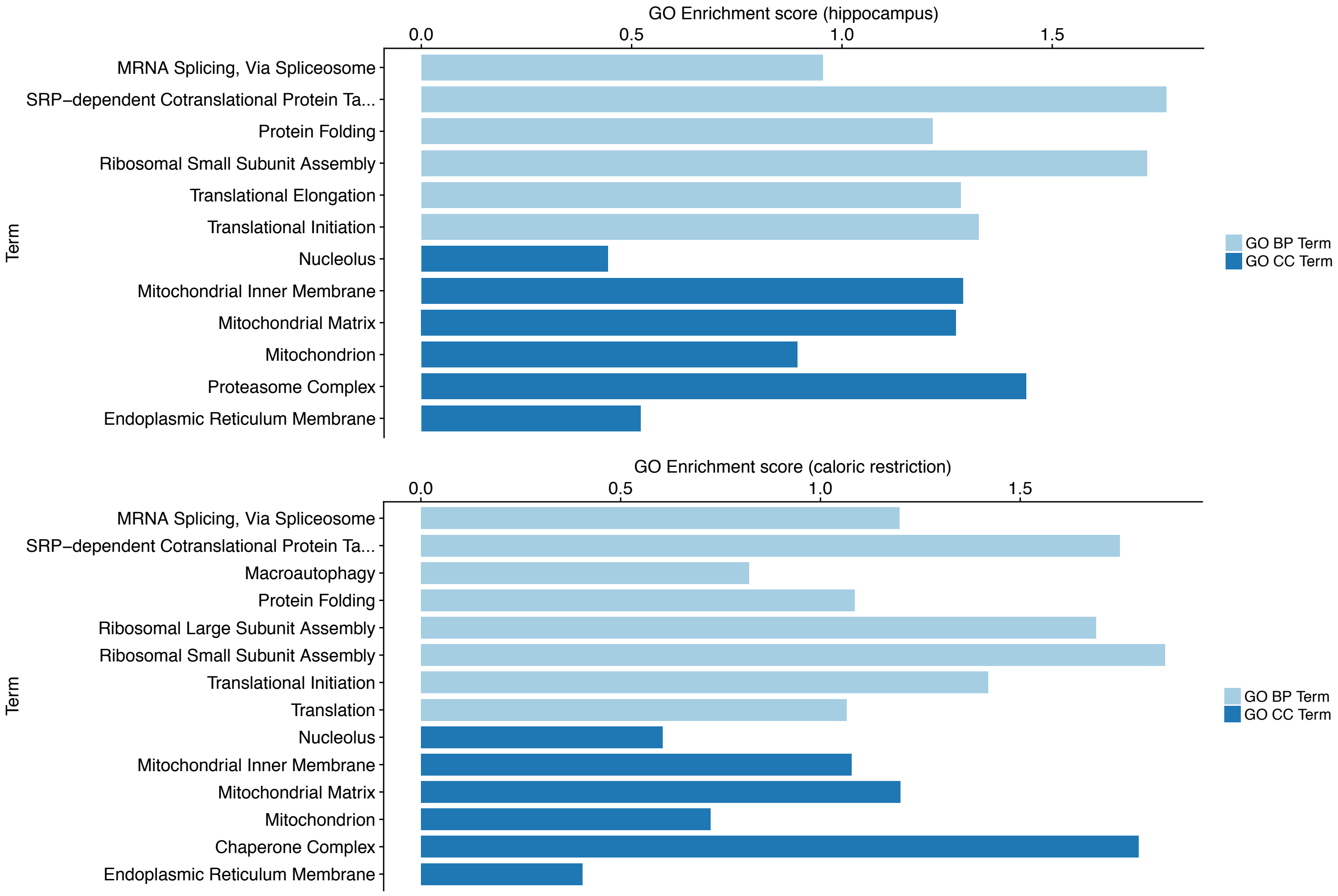


**Figure S7. Proteostasis-linked processes enriched in hippocampus and caloric restriction experiments in human.** The log2 GO enrichment scores are shown for both ‘biological process’ and ‘cellular component’ categories that are related to proteostasis processes.

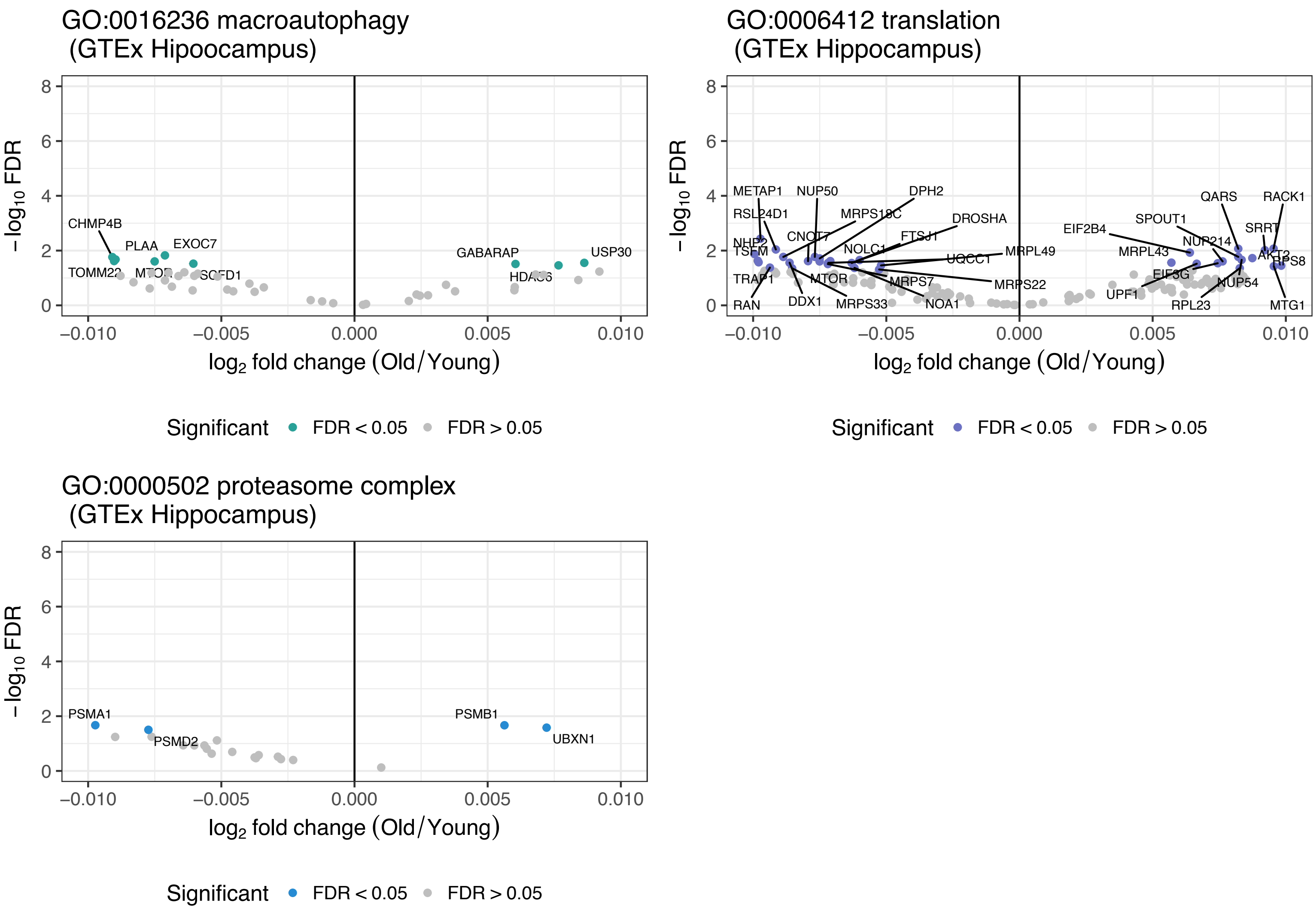


**Figure S8. Volcano plots of conserved gene expression in human hippocampus.** The genes are annotated to human genome. The signal of loss of proteostasis in hippocampus is not as strong as in skeletal muscle (Figure 3).

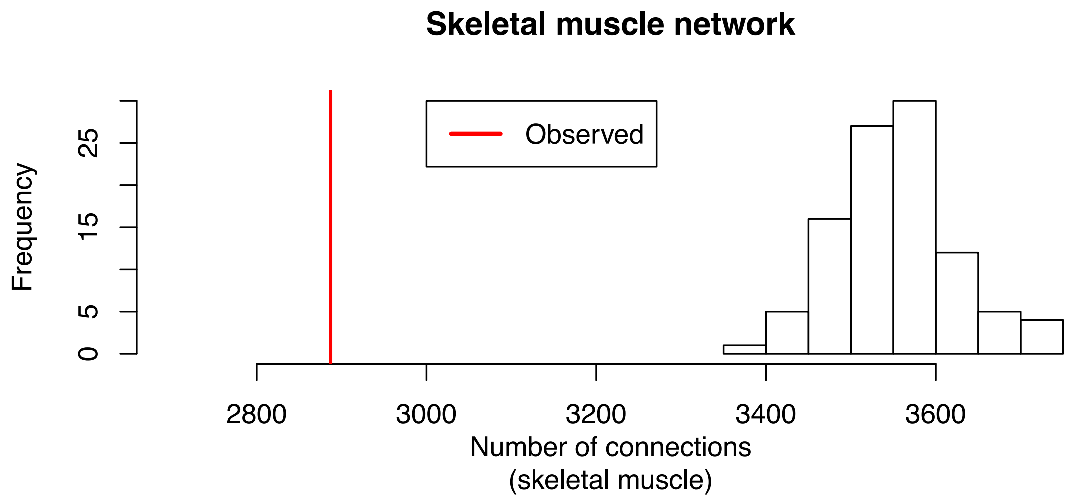


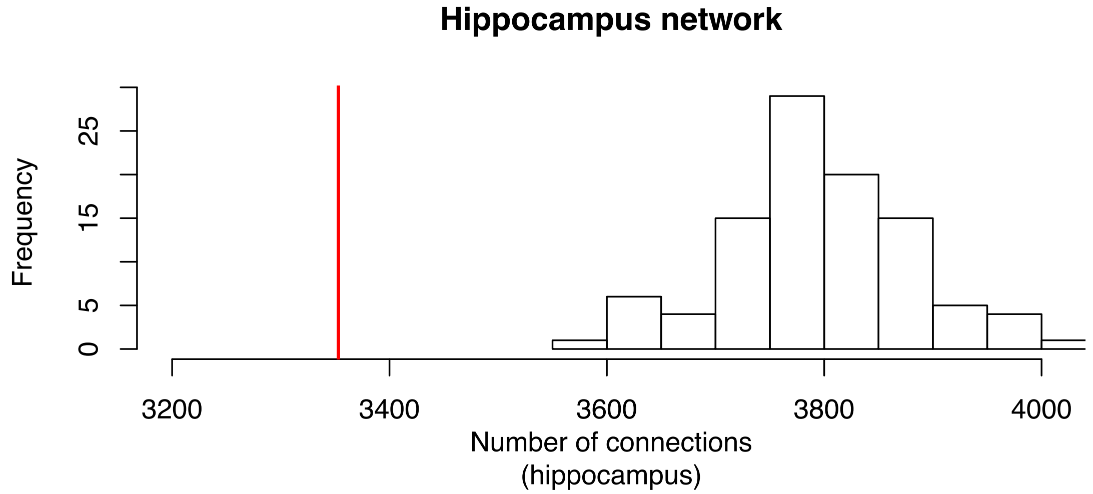


**Figure S9. The number of connections and number of modules from random networks results (100 permutations) in both skeletal muscle and hippocampus data.** The conserved aging co-expression networks show low number of connections than when the integration is performed on the random genes.

**Skeletal muscle M12**
**(GO:000209 protein polyubiquitination)**

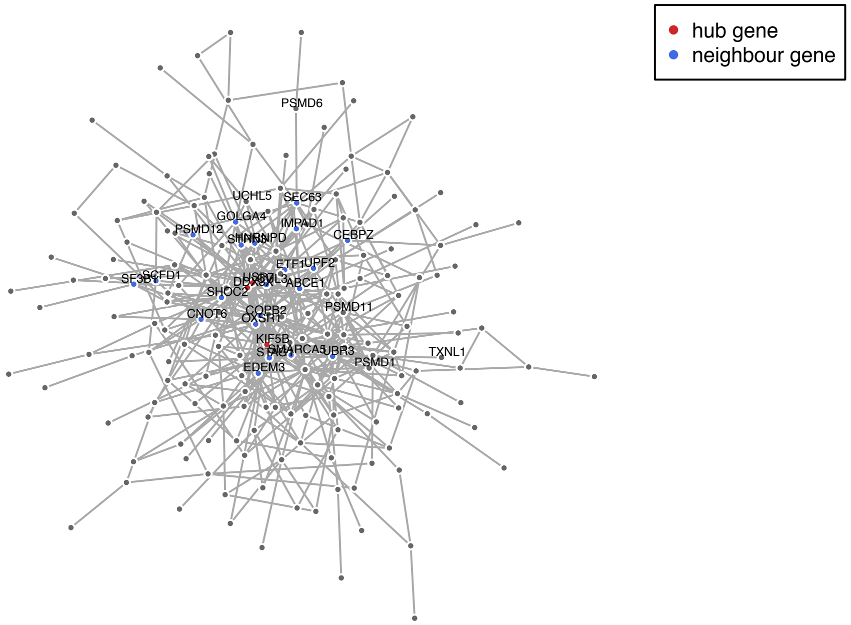


**Hippocampus M3**
**(GO:0000209 protein polyubiquitination)**

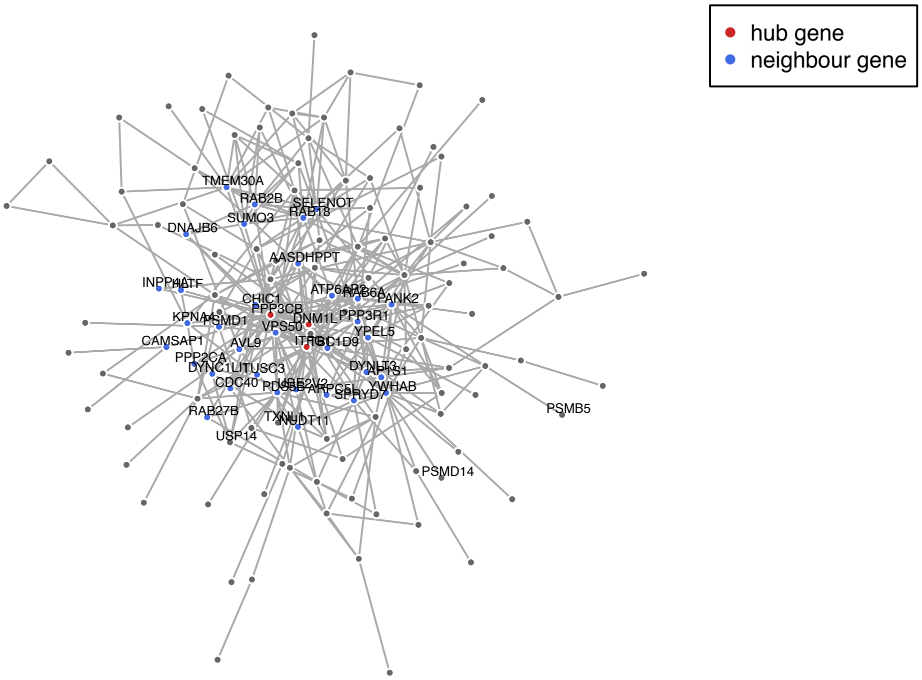


**Figure S10. Additional significantly enriched modules (skeletal muscle (A) on oxidation-reduction process; hippocampus (B) on translational initiation) associated with proteostasis-linked processes and age-related GWAS**. Their hub genes and genes part of the proteasome complex are shown in Figure 5C.
