## Supplementary Materials for "Cross-species functional modules link proteostasis to human normal aging"

### Main Figures + Supplement (Differential expression part)

*Andrea Komljenović*

*3/9/2018*

```
## --------------------------------
# Load the packages

packages <- c( "affy", "affyPLM", "biomaRt", "limma",
               "tidyr", "gplots", "ggplot2", "reshape2",
               "Biobase", "downloader", "sva", "dplyr", "RColorBrewer", "cowplot",
               "igraph", "WriteXLS", "org.Hs.eg.db", "org.Mm.eg.db", "org.Dm.eg.db", "org.Ce.eg.db", "gag
               "cowplot", "ggrepel", "plyr", "data.table", "dplyr", "ComplexHeatmap", "circlize", "RColo
lapply(packages, library, character.only = TRUE)
```

```
path <- "~/Project1/manuscript_GSEA/data_preprocessing/"
```

Loading differentially expressed matrices.

```
# --------------------------
# Human
deg.muscle.human.aging <-
  readRDS(paste0(path, "hsapiens/aging_data/diff_exp/topTable_Muscle_Skeletal_GTEx_V6p.rds"))
deg.hippo.human.aging <-
  readRDS(paste0(path, "hsapiens/aging_data/diff_exp/topTable_Brain_Hippocampus_GTEx_V6p.rds"))


# ---------------------------
# Mouse

# name it correctly + consistency - TO DO
deg.mouse.muscle <-
  readRDS(paste0(path, "mmusculus/aging_data/skeletal_muscle/diffexp_aging_mouse_skeletal_muscle_aging.
deg.muscle.mouse.aging <- deg.mouse.muscle$oldvsyoung

# change here
deg.hippo.mouse.aging <- readRDS(paste0(path, "mmusculus/aging_data/hippocampus/diff_expression_mmuscul

# ---------------------------
# Fly - wb
deg.fly <-
  readRDS(paste0(path, "dmelanogaster/aging_data/diff_expression_fly_aging.rds"))
deg.wholebody.fly.aging <- deg.fly$oldvsyoung

# ---------------------------
# Worm - wb
deg.wholebody.worm.aging <-
  readRDS(paste0(path, "celegans/aging_data/diff_expression_celegans_aging.rds"))
```

```
## Dietary restriction
deg.human.dr <-
```

```
  readRDS(paste0(path, "hsapiens/caloric_restriction_data/differential_expression_human_dietary_restric
## Dietary restriction
deg.mouse.dr <-
  readRDS(paste0(path, "mmusculus/caloric_restriction_data/differential_expression_mouse_dietary_restri
## Dietary restriction - rename this
deg.fly.dr <-
  readRDS(paste0(path, "/dmelanogaster/caloric_restriction_data/diff_exp_fly_dietary_restriction.rds"))
## Dietary restriction
deg.worm.dr <-
  readRDS(paste0(path, "/celegans/caloric_restriction_data/diff_exp_worm_dietary_restriction.rds"))
```

**Barplots of the differential expression analysis**

```
# -----
# FUNCTION
number.of.degs <- function(toptable, cutoff){
    up <- toptable[which(sign(toptable$logFC) == 1 & toptable$adj.P.Val < cutoff),]
    dn <- toptable[which(sign(toptable$logFC) == -1 & toptable$adj.P.Val < cutoff),]
    cat("The number of genes that are downregulated:", dim(dn)[1], "\n")
    cat("The number of genes that are upregulated:", dim(up)[1], "\n")
  #  cat("The number of genes that are downregulated:", dim(dn)[1], "\n")

    return(list(downregulated = dim(dn)[1], upregulated = dim(up)[1]))
}



## ----------------
## preparing the datasets for plotting
list.degs.species <- list(human.muscle = deg.muscle.human.aging,
                          human.hippo = deg.hippo.human.aging,
                          mouse.muscle = deg.muscle.mouse.aging,
                          mouse.hippo = deg.hippo.mouse.aging,
                          fly.wholebody = deg.wholebody.fly.aging,
                          worm.wholebody = deg.wholebody.worm.aging)

signif.expressed.genes <- lapply(list.degs.species, function(x) number.of.degs(x, 0.1))
```

```
## The number of genes that are downregulated: 2540
## The number of genes that are upregulated: 2513
## The number of genes that are downregulated: 2978
## The number of genes that are upregulated: 3105
## The number of genes that are downregulated: 1271
## The number of genes that are upregulated: 1184
## The number of genes that are downregulated: 718
## The number of genes that are upregulated: 921
## The number of genes that are downregulated: 2344
## The number of genes that are upregulated: 2413
## The number of genes that are downregulated: 1634
## The number of genes that are upregulated: 1904
```

```
vec <- unlist(signif.expressed.genes)
ind <- seq(1, length(vec), by = 2)
```

```r
vec[ind] <- vec[ind]*(-1) # to give negative sign for downregulated ones
names(vec) <- ""

# plotting histogram
rnaseq <- data.frame(
    Dataset = c(
    rep("H.sapiens - Skeletal Muscle", 2), rep("H.sapiens - Hippocampus", 2),
  rep("M.musculus - Skeletal Muscle", 2), rep("M.musculus - Hippocampus", 2),
  rep("D.melanogaster - Whole Body", 2),
  rep("C.elegans - Whole Body", 2)),
    Status = c(rep(c("Downregulated","Upregulated"), 6)),
  # differentially expressed genes
    deg = vec)

# make V1 an ordered factor
rnaseq$Dataset <- factor(rnaseq$Dataset, levels = unique(rnaseq$Dataset))

# title <- "Aging"

# pdf("~/Project1/manuscript_GSEA/results/Figure1C_barplot_differential_exp_update.pdf", 7, 3)
ggplot(rnaseq, aes(as.factor(Dataset), deg, fill = Status)) +
  geom_bar(position= "identity", colour="grey50", stat="identity", width=0.8) +
  coord_flip() + scale_x_discrete(limits = rev(levels(rnaseq$Dataset))) +
  labs(x = "",
       y = "Number of detected differentially expressed genes") +
  scale_fill_brewer(type="qual", palette="Pastel1") +
  theme_bw() +
  theme(
    panel.grid.major = element_blank(),
    panel.grid.minor = element_blank(),
    panel.border = element_blank(),
    panel.background = element_blank(),
    axis.ticks.y=element_blank(),
    legend.title=element_blank(),axis.text.x=element_blank(),
    axis.ticks.x=element_blank()) +
    geom_hline(yintercept=0) +
    geom_text(label = abs(rnaseq$deg),
                                   hjust = "center",
                                   vjust = "bottom")
```

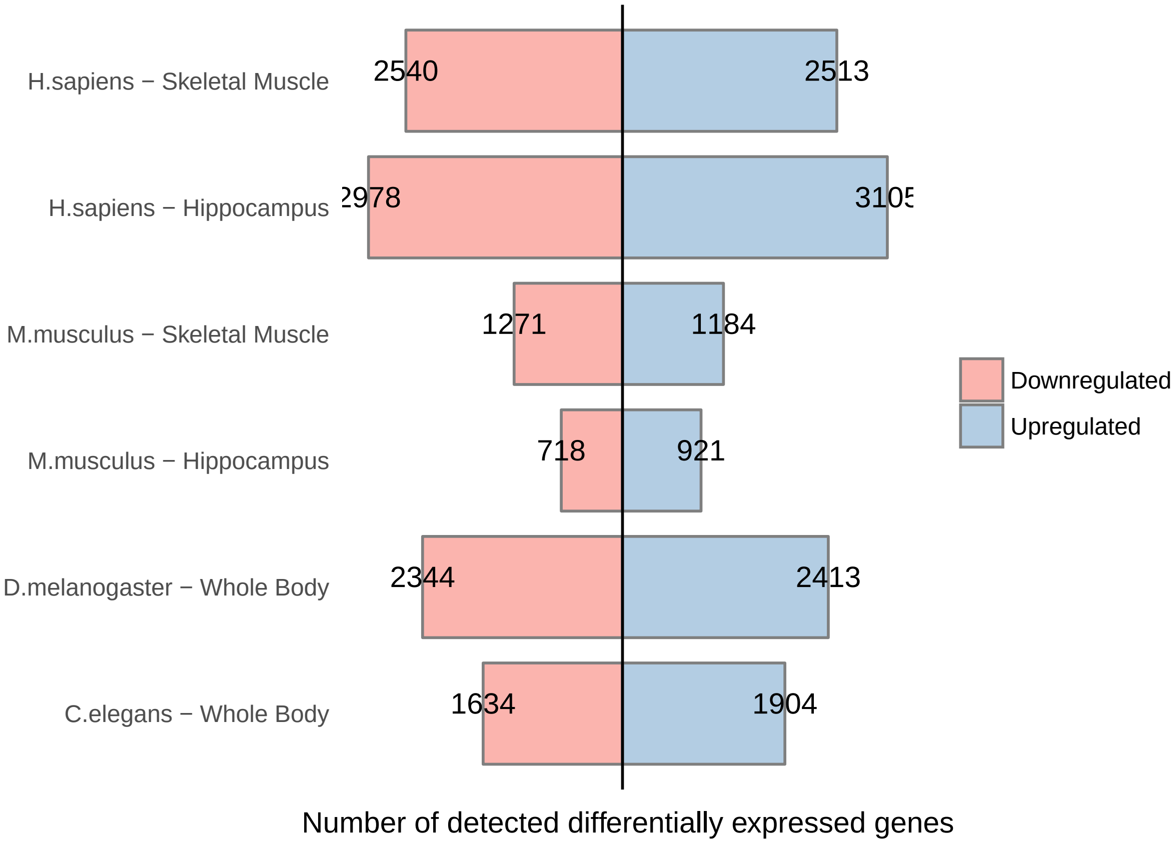


```
# dev.off()
```

##### Alignment of the stages between species

Generating Fig S2. heatmaps of clustering the young and old samples for human and other species, in order to do age alignments.

```
### defining the orthology
orthology.to.human.pc <- function(mart.human){
  # library(biomaRt)
        orth.mouse <-
          getBM(attributes = c("ensembl_gene_id", "external_gene_name", "mmusculus_homolog_ensembl_gene
        "mmusculus_homolog_orthology_type"), filters = "with_mmusculus_homolog", values= TRUE,
                    mart = mart.human, bmHeader = FALSE, uniqueRows = TRUE)

    orth.drosophila <-
      getBM(attributes = c("ensembl_gene_id", "external_gene_name", "dmelanogaster_homolog_ensembl_gene
        "dmelanogaster_homolog_orthology_type"), filters = "with_dmelanogaster_homolog", values= TRUE,
                    mart = mart.human, bmHeader = FALSE, uniqueRows = TRUE)

    orth.celegans <-
      getBM(attributes = c("ensembl_gene_id", "external_gene_name", "celegans_homolog_ensembl_gene",
        "celegans_homolog_orthology_type"), filters = "with_celegans_homolog", values= TRUE,
                    mart = mart.human, bmHeader = FALSE, uniqueRows = TRUE)

    protein.coding <-
```

```r
      getBM(attributes = c("ensembl_gene_id", "gene_biotype"),
            values = TRUE, mart = mart.human,  bmHeader = FALSE, uniqueRows = TRUE)

    orth.mouse <- merge(orth.mouse, protein.coding, by = "ensembl_gene_id")
    orth.drosophila <- merge(orth.drosophila, protein.coding, by = "ensembl_gene_id")
    orth.celegans <- merge(orth.celegans, protein.coding, by = "ensembl_gene_id")
        return(list(ToMouse = orth.mouse, ToDmel = orth.drosophila, ToCele = orth.celegans))
}


### for selecting the  ages
select.ages.gtex <- function(path.annotation.file, expression){

  require(data.table)

  subject.phenotypes <-
    as.data.frame(fread(paste0(path.annotation.file,
    "phs000424.v6.pht002742.v6.p1.c1.GTEx_Subject_Phenotypes.GRU.txt"), header = TRUE))
    colnames(subject.phenotypes) <- as.vector(as.matrix(subject.phenotypes[1,]))
    subject.phenotypes <- subject.phenotypes[-1,]

    ages <- as.character(subject.phenotypes$AGE);
    names(ages) <- subject.phenotypes$SUBJID
    # 20-30 years; 61-70 years

    # this removes exact ages
    young.gtex <- rep("young", length(ages[which(ages >= "20" & ages <= "30")]))
    names(young.gtex) <- names(ages[which(ages >= "20" & ages <= "30")])

    old.gtex <- rep("old", length(ages[which(ages >= "61" & ages <= "70")]))
    names(old.gtex) <- names(ages[which(ages >= "61" & ages <= "70")])

    selected.ages.gtex <- c( young.gtex, old.gtex )

    expression.human <-
      expression[, na.omit(match( names(selected.ages.gtex), colnames(expression) ))]

    selected.ages.gtex.tissue <-
      selected.ages.gtex[na.omit(match(colnames(expression.human),names(selected.ages.gtex)))]
    colnames(expression.human) <-
      selected.ages.gtex.tissue

    return(expression.human)
}



select.signif.genes <- function( diff.expression.human, diff.expression.sp2,
                                 common.orthologs.human.sp2,
                                 ensembl.sp2.gnames, cutoff = NULL ) {
    # here it is always 1-1 orthologs
    # human
    significant.human <-
```

```
      diff.expression.human[diff.expression.human$logFC > 0 &
                              diff.expression.human$adj.P.Val < cutoff,]
    # species2
    significant.sp2 <-
      diff.expression.sp2[diff.expression.sp2$logFC > 0 &
                            diff.expression.sp2$adj.P.Val < cutoff,]
    msig1 <- merge(common.orthologs.human.sp2, significant.sp2,
                   by.x = ensembl.sp2.gnames, by.y = "row.names" )
    msig2 <- merge(msig1, significant.human,
                   by.x = "ensembl_gene_id", by.y = "row.names" )

    return(msig2)
}


# for supplement data
select.signif.genes.sp <- function( diff.expression.human, diff.expression.sp2,
                                     common.orthologs.human.sp2,
                                    ensembl.sp2.gnames, cutoff = NULL, tissue.name ) {

    # here it is always 1-1 orthologs

  # human
    significant.human <- diff.expression.human[diff.expression.human$adj.P.Val < cutoff,]
  cat("Number of significant DEGs in human dataset:", nrow(significant.human), "\n")
    # species2
    significant.sp2 <- diff.expression.sp2[diff.expression.sp2$adj.P.Val < cutoff,]
  cat("Number of significant DEGs in mouse dataset:", nrow(significant.sp2), "\n")


  deg.number <- paste(nrow(significant.human), nrow(significant.sp2), sep = "/")

    msig1 <-
      merge(common.orthologs.human.sp2,
                   significant.sp2, by.x = ensembl.sp2.gnames, by.y = "row.names" )
    msig2 <-
      merge(msig1, significant.human,
            by.x = "ensembl_gene_id", by.y = "row.names" )
  cat("Number of common DEGs based on 1-1 orthologs:", nrow(msig2), "\n")
  # calculate overlap
  overlap.percentage <- (nrow(msig2)/nrow(significant.human))*100
  df <- data.frame(Tissue = tissue.name, DEGnumber = deg.number,
                   OneToOneOrthologsNumber = nrow(common.orthologs.human.sp2),
                   Overlap.percentage = overlap.percentage)
    return(df)
}

# the colnames of expression matrices should be named as they wanted to be shown in heatmap
clustering.samples <- function( merged.significant.genes, expression.human,
                                expression.sp2, ensembl.sp2.gnames,
    filename, heatmap.title, plot = FALSE ){

    library(sva)
```

```
    colnames.human <- colnames(expression.human)
    colnames.sp2 <- colnames(expression.sp2)

    batch <- c(rep("1", ncol(expression.human)), rep("2", ncol(expression.sp2)))
    modcombat <- model.matrix(~1, data=as.data.frame(batch))


    mm <-
      merge(merged.significant.genes, expression.human,
            by.x = "ensembl_gene_id", by.y = "row.names")
    mm2 <- merge(mm, expression.sp2,
            by.x = ensembl.sp2.gnames, by.y = "row.names")
    # remove the diff. exp results
    mm2 <- mm2[, -c(1:17)]
    colnames(mm2) <- c(colnames.human, colnames.sp2)

    combat.expression.data <- ComBat(dat = mm2,
                                     batch = batch,
                                     mod = modcombat,
                                     par.prior = TRUE, prior.plots = FALSE)
    # there is few outliers but looks ok

  hclust.comp <- function(x) hclust(x, method="complete")

  if(plot) {
    pdf(filename, 15, 5)
        hclust.comp <- function(x) hclust(x, method="complete")
        gplots::heatmap.2(as.matrix(combat.expression.data), scale = "row",
                                            labRow = FALSE,
                                            col=brewer.pal(11,"RdBu"), trace="none",
                                            margins =c(12,9), hclustfun=hclust.comp,
                                            main = heatmap.title)
        dev.off()

  } else {

    gplots::heatmap.2(as.matrix(combat.expression.data), scale = "row",
                      labRow = FALSE,
                      col=brewer.pal(11,"RdBu"), trace="none",
                      margins =c(12,9), hclustfun=hclust.comp,
                      main = heatmap.title)

  }

    return(combat.expression.data)

}
```

Load the expression matrices:

```
##### ------------------------------------------------------------------
## Load expression data

# ----------------------
```

```
# human
exp.human.hippo <-
  readRDS(paste0(path, "/hsapiens/aging_data/exp_mat/voom_Brain_Hippocampus_GTEx_V6p.rds"))
exp.human.muscle <-
  readRDS(paste0(path, "hsapiens/aging_data/exp_mat/voom_Muscle_Skeletal_GTEx_V6p.rds"))


# for gtex data, selected only the youngest and the oldest samples (extremes) to show the alignments
expression.human.hippo <-
  select.ages.gtex("~/Documents/Lausanne/GTEx_annotation/", exp.human.hippo)
colnames(expression.human.hippo) <-
  paste0("Hs.hippo.", colnames(expression.human.hippo)) # 41

expression.human.muscle <-
  select.ages.gtex("~/Documents/Lausanne/GTEx_annotation/", exp.human.muscle)
colnames(expression.human.muscle) <-
  paste0("Hs.muscle.", colnames(expression.human.muscle)) # 142


# -------------------------
# mouse
expression.mouse.muscle <-
  readRDS(paste0(path, "/mmusculus/aging_data/skeletal_muscle/expression_matrix_mouse_skeletal_muscle_a
expression.mouse.hippo <-
  readRDS(paste0(path, "mmusculus/aging_data/hippocampus/voom_mouse_expression_matrix_hippocampus_aging

colnames(expression.mouse.muscle) <- c(rep("young",4),rep("old", 5))
colnames(expression.mouse.muscle) <- paste0("Mm.muscle.", colnames(expression.mouse.muscle))
colnames(expression.mouse.hippo) <- paste0("Mm.hippo.", colnames(expression.mouse.hippo))


# ------------------------
# fly
expression.microarray.fly <- readRDS(paste0(path, "/dmelanogaster/aging_data/expression_matrix_fly_agin
expression.microarray.fly <- expression.microarray.fly[[2]]
colnames(expression.microarray.fly) <- paste0("Dm.wb.", colnames(expression.microarray.fly))



# -----------------------
# worm
expression.wb.worm <-
  readRDS(paste0(path, "/celegans/aging_data/voom_celegans_expression_matrix_aging.rds"))
expression.wb.worm <- expression.wb.worm[, !(colnames(expression.wb.worm) %in% "too_young")]
colnames(expression.wb.worm) <- paste0("Ce.wb.", colnames(expression.wb.worm))
```

Define 1-1 orthologs between species:

```
library(biomaRt)
human.ensembl <- useMart(biomart="ENSEMBL_MART_ENSEMBL",
                         host="www.ensembl.org",
                         path="/biomart/martservice",
                         dataset="hsapiens_gene_ensembl",
                         version = "Ensembl Genes 91")
```

```
# orthology to human on protein coding genes only
common.human.orthologs.pc <- orthology.to.human.pc(human.ensembl)
common.human.orthologs.final.pc <- lapply(common.human.orthologs.pc,
                                          function(x) x[which(x[,4]== "ortholog_one2one"),])
common.human.orthologs.one2one <- lapply(common.human.orthologs.final.pc,
                                         function(x) x[which(x[,5]== "protein_coding"),])
```

Plotting the heatmap for Human-Mouse for Figure 2A.

```
# 52 genes
signif.muscle <- select.signif.genes(deg.muscle.human.aging, deg.muscle.mouse.aging,
                                                                        common.human.orthologs.one2one$
                                                                        "mmusculus_homolog_ensembl_gene


# 94 genes
signif.hippo <- select.signif.genes(deg.hippo.human.aging, deg.hippo.mouse.aging,
                                                                            common.human.orthologs.one2
                                                                            "mmusculus_homolog_ensembl_



clusters.muscle.human.mouse <- clustering.samples( signif.muscle,
                                                   expression.human.muscle, expression.mouse.muscle,
                                                             "mmusculus_homolog_ensembl_gene",
                                                             "~/Project1/manuscript_GSEA/results/Figure
                                                             "Human - Mouse skeletal muscle", plot = FA
```

```
## Found2batches

## Adjusting for0covariate(s) or covariate level(s)

## Standardizing Data across genes

## Fitting L/S model and finding priors

## Finding parametric adjustments

## Adjusting the Data
```

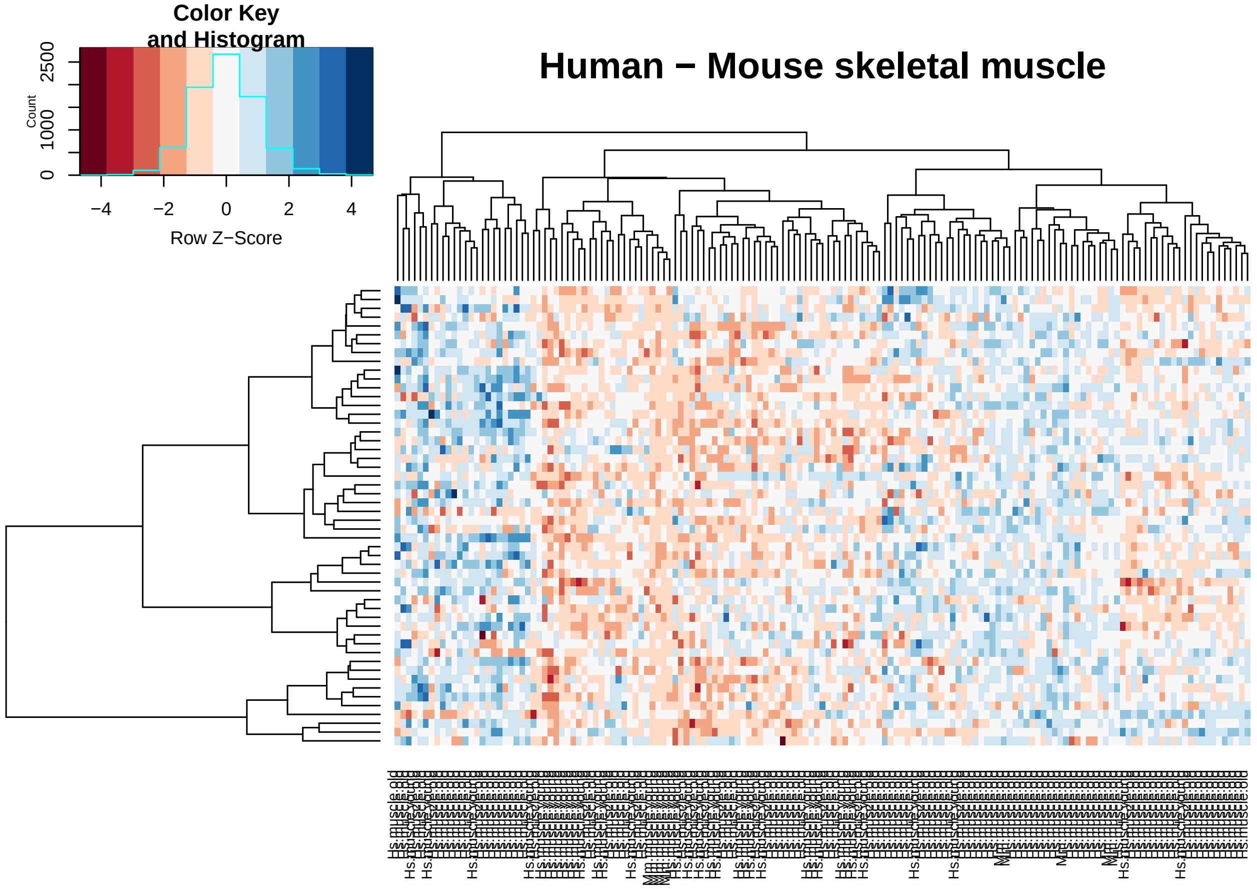


```
clusters.hippo.human.mouse <- clustering.samples( signif.hippo,
                                                  expression.human.hippo, expression.mouse.hippo,
                                                            "mmusculus_homolog_ensembl_gene",
                                                  "~/Project1/manuscript_GSEA/results/Figure2A_heatmap_I
                                                            "Human - Mouse hippocampus", plot = FALSE)
```

```
## Found2batches

## Adjusting for0covariate(s) or covariate level(s)

## Standardizing Data across genes

## Fitting L/S model and finding priors

## Finding parametric adjustments

## Adjusting the Data
```

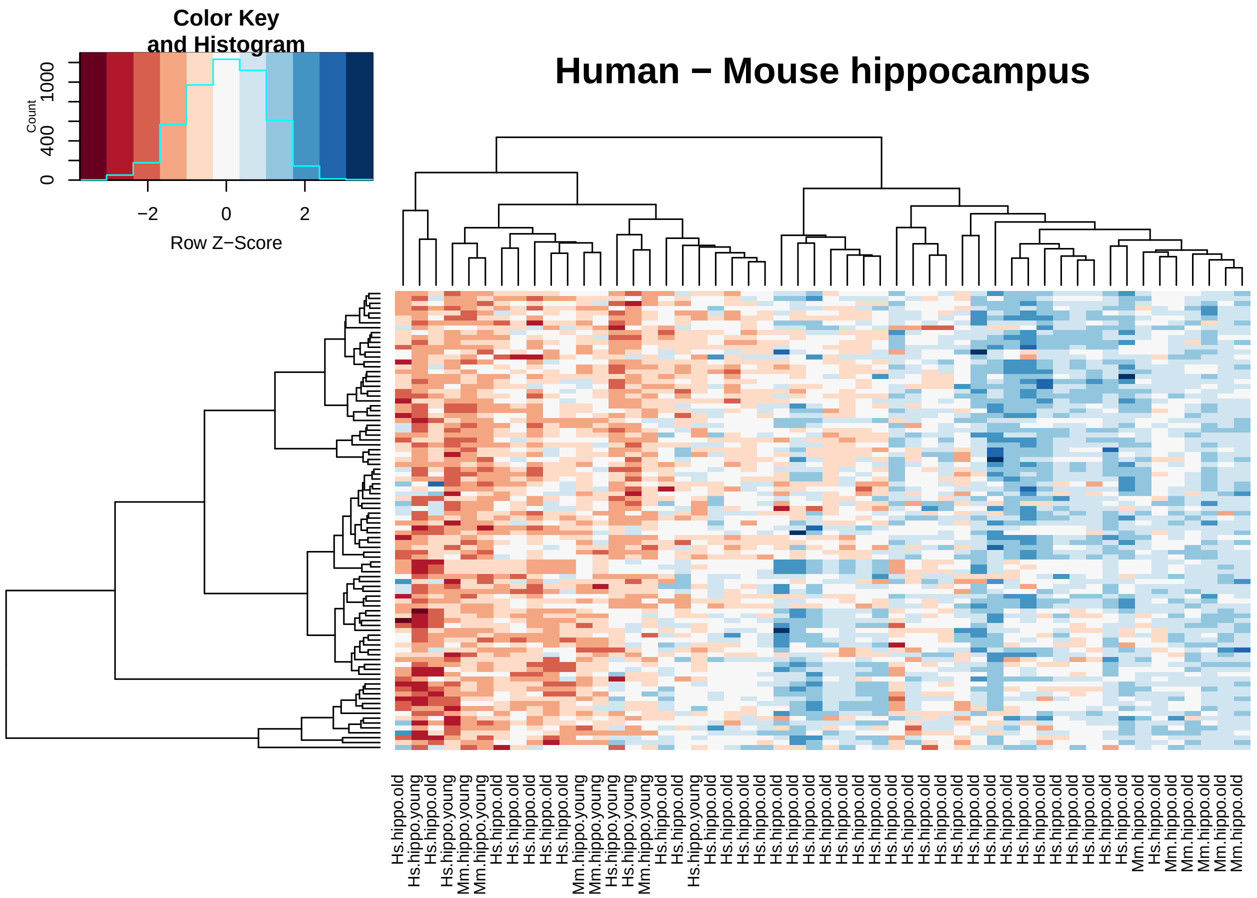


Plotting for Supplement Figures S2: Human-Fly, Human-Worm:

```
# 42
signif.muscle.human.fly <- select.signif.genes(deg.muscle.human.aging, deg.wholebody.fly.aging,
                                                common.human.orthologs.one2one$ToDmel,
                                                "dmelanogaster_homolog_ensembl_gene", cutoff = 0.05)

# 87
signif.hippo.human.fly <- select.signif.genes(deg.hippo.human.aging, deg.wholebody.fly.aging,
                                               common.human.orthologs.one2one$ToDmel,
                                               "dmelanogaster_homolog_ensembl_gene", cutoff = 0.05)

clusters.human.fly.muscle <- clustering.samples( signif.muscle.human.fly, expression.human.muscle, expre
                                                "dmelanogaster_homolog_ensembl_gene",
                                                "~/Project1/manuscript_GSEA/results/FigureS2_heatmap_hu
                                                "Human muscle - Fly whole body", plot = FALSE)
```

```
## Found2batches

## Adjusting for0covariate(s) or covariate level(s)

## Standardizing Data across genes

## Fitting L/S model and finding priors

## Finding parametric adjustments

## Adjusting the Data
```

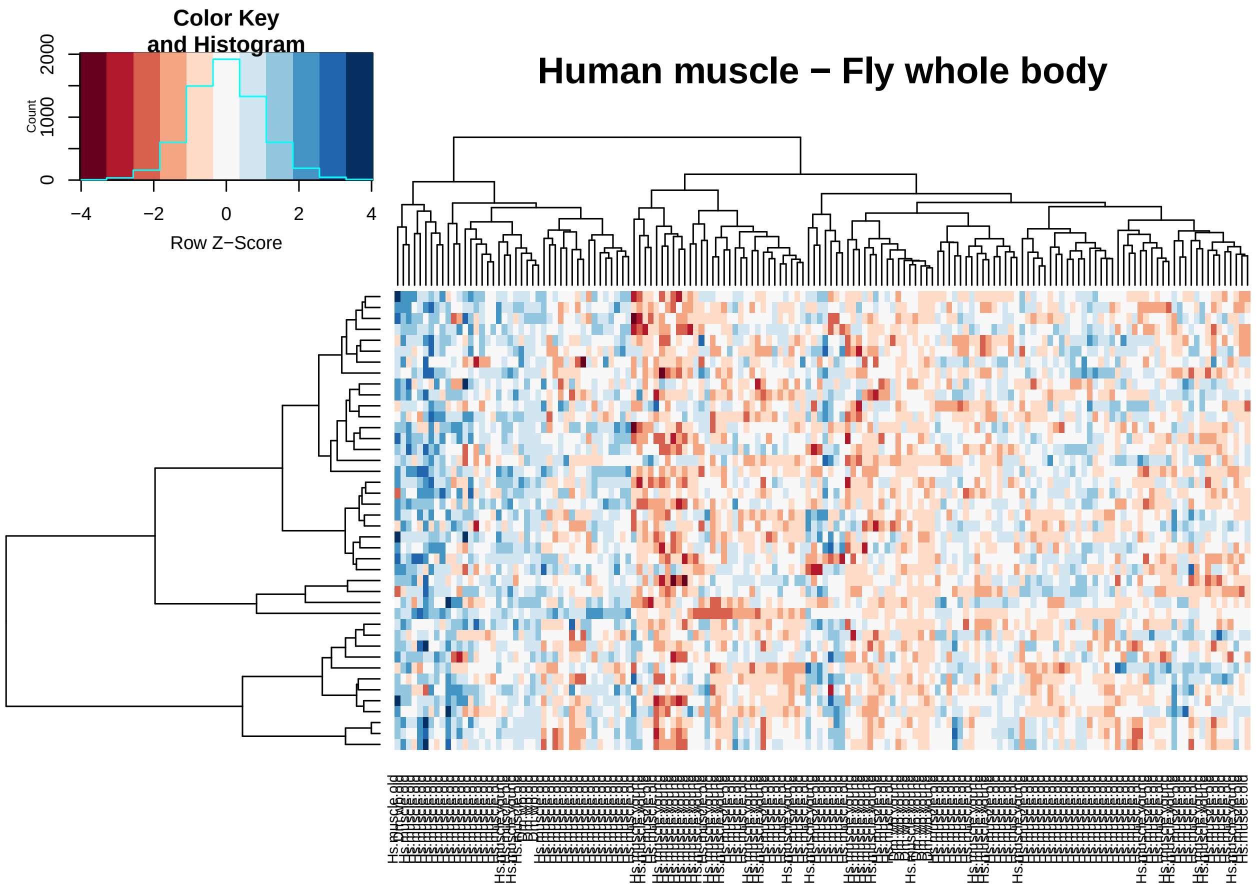


```
clusters.human.fly.hippo <- clustering.samples( signif.hippo.human.fly , expression.human.hippo, express
                                               "dmelanogaster_homolog_ensembl_gene",
                                               "~/Project1/manuscript_GSEA/results/FigureS2_heatmap_hu
                                               "Human hippocampus - Fly whole body", plot = FALSE)
```

```
## Found2batches

## Adjusting for0covariate(s) or covariate level(s)

## Standardizing Data across genes

## Fitting L/S model and finding priors

## Finding parametric adjustments

## Adjusting the Data
```

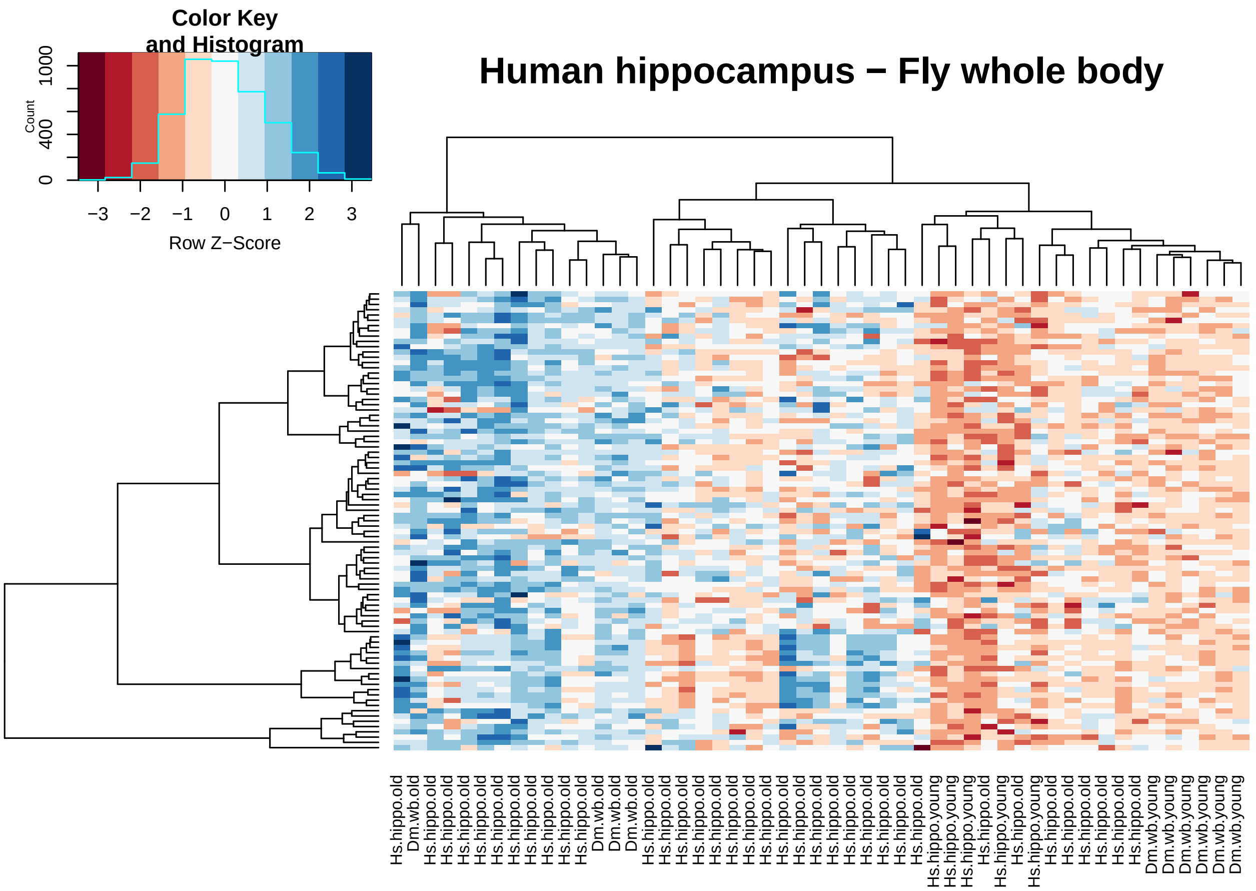


```
## here I had to change to the 0.05 because I get only 2 genes when cutoff 0.01

# 10
signif.muscle.human.worm <- select.signif.genes(deg.muscle.human.aging, deg.wholebody.worm.aging,
                                                              common.human.orthologs.one2one$ToCele,

# 17
signif.hippo.human.worm <- select.signif.genes(deg.hippo.human.aging, deg.wholebody.worm.aging,
                                                             common.human.orthologs.one2one$ToCele, "


clusters.human.worm.muscle <- clustering.samples( signif.muscle.human.worm, expression.human.muscle, ex
                                                "celegans_homolog_ensembl_gene",
                                                "~/Project1/manuscript_GSEA/results/FigureS2_heatmap_
                                                "Human muscle - Worm whole body", plot = FALSE)
```

```
## Found2batches

## Adjusting for0covariate(s) or covariate level(s)

## Standardizing Data across genes

## Fitting L/S model and finding priors

## Finding parametric adjustments

## Adjusting the Data
```

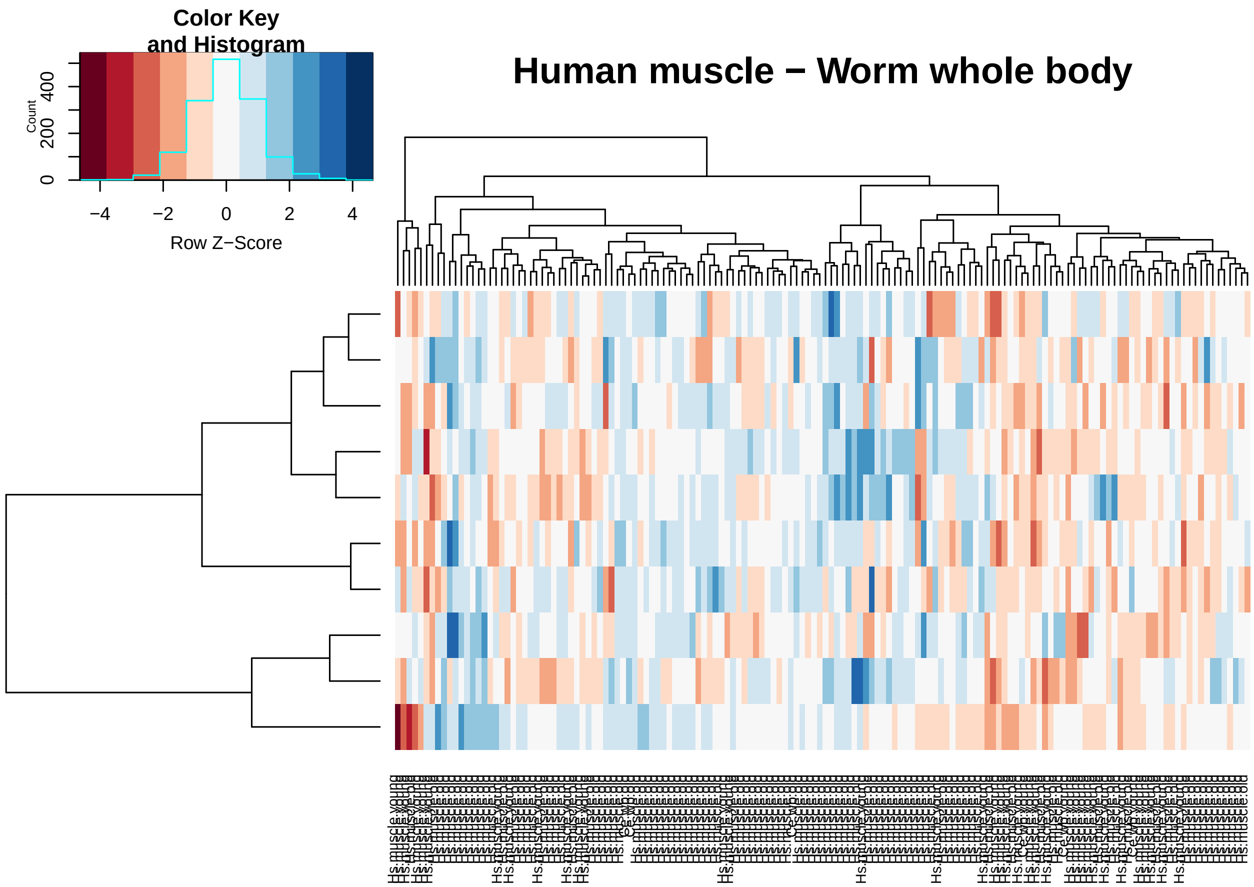


```
clusters.human.worm.hippo <- clustering.samples( signif.hippo.human.worm, expression.human.hippo,express
                                                 "celegans_homolog_ensembl_gene",
                                                 "~/Project1/manuscript_GSEA/results/FigureS2_heatmap_hu
                                                 "Human hippocampus - Worm whole body", plot = FALSE)
```

```
## Found2batches

## Adjusting for0covariate(s) or covariate level(s)

## Standardizing Data across genes

## Fitting L/S model and finding priors

## Finding parametric adjustments

## Adjusting the Data
```

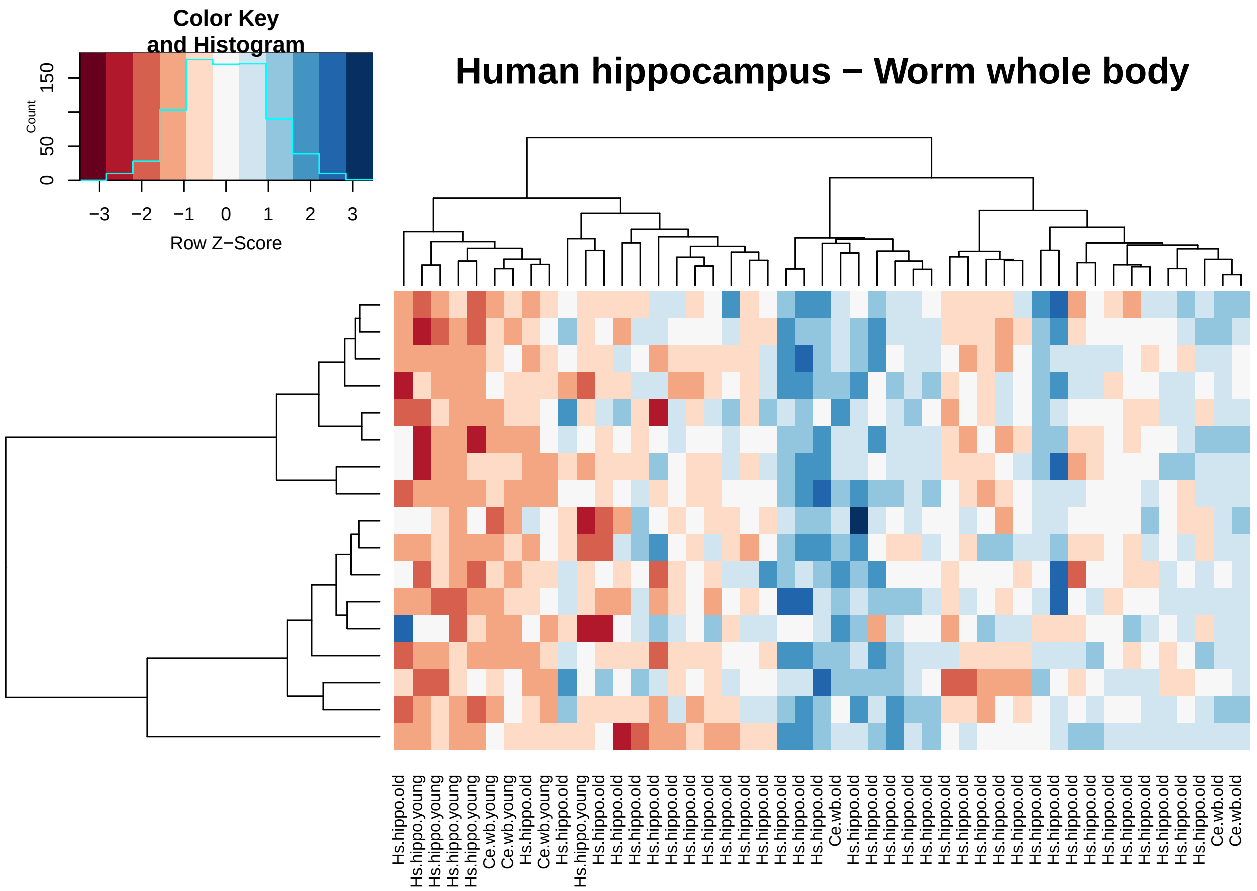


**Table S2. Differential expression summary stats per species**

```
annotation.deg.matrices <- function(genome, differential.exp.matrix){

  differential.exp.matrix$gene.names <- mapIds(genome,
                                               keys=rownames(differential.exp.matrix),
                                               column="SYMBOL",
                                               keytype="ENSEMBL",
                                               multiVals="first")
  # differential.exp.matrix <- #differential.exp.matrix[complete.cases(differential.exp.matrix$gene_nam
  return(differential.exp.matrix)

}



anno.deg.human.muscle <- annotation.deg.matrices(org.Hs.eg.db, deg.muscle.human.aging)
```

```
## 'select()' returned 1:many mapping between keys and columns
```

```
anno.deg.human.hippo <- annotation.deg.matrices(org.Hs.eg.db, deg.hippo.human.aging)
```

```
## 'select()' returned 1:many mapping between keys and columns
```

```
# mouse
anno.deg.mouse.muscle <- annotation.deg.matrices(org.Mm.eg.db, deg.muscle.mouse.aging)
```

```
## 'select()' returned 1:many mapping between keys and columns
```

```
anno.deg.mouse.hippo <- annotation.deg.matrices(org.Mm.eg.db, deg.hippo.mouse.aging)
```

```
## 'select()' returned 1:many mapping between keys and columns
```

```
# fly
anno.deg.fly.wholebody <- annotation.deg.matrices(org.Dm.eg.db, deg.wholebody.fly.aging)
```

```
## 'select()' returned 1:many mapping between keys and columns
```

```
# worm
anno.deg.worm.wholebody <- annotation.deg.matrices(org.Ce.eg.db, deg.wholebody.worm.aging)
```

```
## 'select()' returned 1:many mapping between keys and columns
```

```
differential.expression.gene.list <-
  list(DEG_muscle_human_aging = anno.deg.human.muscle,
       DEG_hippo_human_aging = anno.deg.human.hippo,
       DEG_muscle_mouse_aging = anno.deg.mouse.muscle,
       DEG_hippo_mouse_aging = anno.deg.mouse.hippo,
       DEG_wholebody_fly_aging = anno.deg.fly.wholebody,
       DEG_wholebody_worm_aging = anno.deg.worm.wholebody)

differential.expression.gene.list<-
  lapply(differential.expression.gene.list,
         function(x) {  x$ENSEMBL <- rownames(x) ; x })

#### ---------------------------------------------
# Supplement Table S2

# WriteXLS(differential.expression.gene.list,
#          ExcelFileName = "~/Project1/manuscript_GSEA/supplementary_data/Table_S2.xlsx", SheetNames = N
```

**Table S3. Percentage of overlap pairwise species human vs others.**

```
### muscle
human.mouse.muscle <- select.signif.genes.sp(deg.muscle.human.aging, deg.muscle.mouse.aging,
                                common.human.orthologs.one2one$ToMouse, "mmusculus_homolog_ensembl_gene
                cutoff = 0.1, "Skeletal Muscle (Human)/Skeletal Muscle (Mouse)") #778
```

```
## Number of significant DEGs in human dataset: 5053
## Number of significant DEGs in mouse dataset: 2455
## Number of common DEGs based on 1-1 orthologs: 778
```

```
human.fly.muscle <- select.signif.genes.sp(deg.muscle.human.aging, deg.wholebody.fly.aging,
                                common.human.orthologs.one2one$ToDmel, "dmelanogaster_homolog_ensembl_ge
                cutoff = 0.1, "Skeletal Muscle (Human)/Whole Body (Fly)") #395
```

```
## Number of significant DEGs in human dataset: 5053
## Number of significant DEGs in mouse dataset: 4757
## Number of common DEGs based on 1-1 orthologs: 395
```

```
human.worm.muscle <- select.signif.genes.sp(deg.muscle.human.aging,deg.wholebody.worm.aging,
                                common.human.orthologs.one2one$ToCele, "celegans_homolog_ensembl_gene",
                cutoff = 0.1, "Skeletal Muscle (Human)/Whole Body (Worm)") #103
```

```
## Number of significant DEGs in human dataset: 5053
```

```
## Number of significant DEGs in mouse dataset: 3538
## Number of common DEGs based on 1-1 orthologs: 103
```

```r
#### hippocampus
human.mouse.hippo <- select.signif.genes.sp(deg.hippo.human.aging, deg.hippo.mouse.aging,
                                               common.human.orthologs.one2one$ToMouse, "mmusculus_homol
                                cutoff = 0.1, "Hippocampus (Human)/Hippocampus (Mouse)") #662
```

```
## Number of significant DEGs in human dataset: 6083
## Number of significant DEGs in mouse dataset: 1639
## Number of common DEGs based on 1-1 orthologs: 662
```

```r
human.fly.hippo <- select.signif.genes.sp(deg.hippo.human.aging, deg.wholebody.fly.aging,
                                               common.human.orthologs.one2one$ToDmel, "dmelanogaster_ho
                                cutoff = 0.1, "Hippocampus (Human)/Whole Body (Fly)") #379
```

```
## Number of significant DEGs in human dataset: 6083
## Number of significant DEGs in mouse dataset: 4757
## Number of common DEGs based on 1-1 orthologs: 379
```

```r
human.worm.hippo <- select.signif.genes.sp(deg.hippo.human.aging, deg.wholebody.worm.aging,
                                               common.human.orthologs.one2one$ToCele, "celegans_homolog
                                cutoff = 0.1, "Hippocampus (Human)/Whole Body (Worm)") # 112
```

```
## Number of significant DEGs in human dataset: 6083
## Number of significant DEGs in mouse dataset: 3538
## Number of common DEGs based on 1-1 orthologs: 112
```

```r
## supplement table S3
overlap <- do.call("rbind", list(human.mouse.muscle, human.fly.muscle,
                                 human.worm.muscle, human.mouse.hippo,
                                 human.fly.hippo, human.worm.hippo))

# WriteXLS(overlap,
#         ExcelFileName = "~/Project1/manuscript_GSEA/supplementary_data/Table_S3.xlsx",
#         SheetNames = NULL, row.names = FALSE)
```

Fig S3. Correlations on orthologous gene-levels.

Single gene-level analysis.

```r
## -------------------
# FUNCTIONS
## -------------------

defining.cutoff <- function(single.gene.df, cutoff = NULL){
      return(single.gene.df$sp1.sp2[which(single.gene.df$sp1.sp2$adj.P.Val.x < cutoff &
                                            single.gene.df$sp1.sp2$adj.P.Val.y < cutoff),])
}


labeling.plot.cutoff <- function(correlation, merged.mat, plot.object){
  # require(cowplot)
  label.sp <- substitute(paste(rho, " = ", estimate, ", p = ", pvalue),
                         list(estimate = signif(correlation$estimate, 2),
                              pvalue = signif(correlation$p.value, 2)))
  label.n <- substitute(paste("n = ", n), list(n = dim(merged.mat)[1]))
```

```r
  pp <- ggdraw(plot.object) + draw_label(label.sp, .25, .9, size = 10)
  ppp <- ggdraw(pp) + draw_label(label.n, .3, .85, size = 10)

  return(ppp)
}

labeling.plot.gs <- function(correlation, plot.object){
  # require(cowplot)
  label.sp <- substitute(paste(rho, " = ", estimate, ", p = ", pvalue),
                         list(estimate = signif(correlation$corr$estimate, 2),
                              pvalue = signif(correlation$corr$p.value, 2)))
  label.n <- substitute(paste("n = ", n), list(n = dim(correlation$sp1.sp2)[1]))

  pp <- ggdraw(plot.object) + draw_label(label.sp, .3, .9, size = 10)
  ppp <- ggdraw(pp)  + draw_label(label.n, .3, .85, size = 10)

  return(ppp)
}



correlation.spearman <- function(matrix.species){
  print(cor.test(matrix.species$logFC.x, matrix.species$logFC.y, method = "spearman"))
  correlation <- cor.test(matrix.species$logFC.x, matrix.species$logFC.y, method = "spearman")
  return(correlation)
}

plotting.singlegene <- function(data, xname, yname, colour.dots, colour.lm, gtitle){
  ggplot(data, aes(x = logFC.x, y = logFC.y)) +
    geom_point(size = 2.5, alpha = 0.5, colour = colour.dots) +
    labs(list(title = gtitle,  x = xname, y = yname)) + xlim(c(-0.06, 0.06) ) + ylim(c(-4,3)) +
    theme(text = element_text(size=12)) +
    geom_hline(yintercept=0, colour = "grey50") +
    geom_vline(xintercept = 0, colour = "grey50")
}

correlations.singlegene.level.aging <- function(species1, species2, orthologs.relationships,
                                                ensembl.id.names.species, xtext, ytext){

  merged.ortho.species1 <- merge(orthologs.relationships, species1, by.x = "ensembl_gene_id", by.y= "ro
  merged.species1.species2 <- merge(merged.ortho.species1, species2, by.x = ensembl.id.names.species, b
  cat("Number of common genes between the species after filtering:", dim(merged.species1.species2)[1],

  cat("Summary of", xtext, ytext, "\n")
  print(cor.test(merged.species1.species2$logFC.x, merged.species1.species2$logFC.y, method = "spearman
  correlation <- cor.test(merged.species1.species2$logFC.x, merged.species1.species2$logFC.y, method =

  return(list(corr = correlation, sp1.sp2 = merged.species1.species2))
}

map.the.entrez <- function(toptable, genome.db){
  toptable$entrez = AnnotationDbi::mapIds(genome.db,
                                          keys=rownames(toptable),
```

```r
                                                column="ENTREZID",
                                                keytype="ENSEMBL",
                                                multiVals="first")
  return(toptable)
}
```

```
## Muscle

# Human -> Mouse
corr.HumanMouse.SingleGene.Muscle <- correlations.singlegene.level.aging( deg.muscle.human.aging,
        deg.muscle.mouse.aging, common.human.orthologs.one2one$ToMouse, "mmusculus_homolog_ensembl_gene
```

```
## Number of common genes between the species after filtering: 9287
## Summary of Human Mouse
##
##  Spearman's rank correlation rho
##
## data:  merged.species1.species2$logFC.x and merged.species1.species2$logFC.y
## S = 1.3879e+11, p-value = 0.0001329
## alternative hypothesis: true rho is not equal to 0
## sample estimates:
##         rho
## -0.03963933
```

```r
# Human -> Fly
corr.HumanFly.SingleGene.Muscle <- correlations.singlegene.level.aging( deg.muscle.human.aging,
    deg.wholebody.fly.aging, common.human.orthologs.one2one$ToDmel, "dmelanogaster_homolog_ensembl_gene
```

```
## Number of common genes between the species after filtering: 2808
## Summary of Human Mouse
##
##  Spearman's rank correlation rho
##
## data:  merged.species1.species2$logFC.x and merged.species1.species2$logFC.y
## S = 3385500000, p-value = 1.189e-05
## alternative hypothesis: true rho is not equal to 0
## sample estimates:
##        rho
## 0.08254312
```

```r
# Human -> Worm
corr.HumanWorm.SingleGene.Muscle <- correlations.singlegene.level.aging( deg.muscle.human.aging,
        deg.wholebody.worm.aging, common.human.orthologs.one2one$ToCele, "celegans_homolog_ensembl_gene
```

```
## Number of common genes between the species after filtering: 2302
## Summary of Human Mouse
##
##  Spearman's rank correlation rho
##
## data:  merged.species1.species2$logFC.x and merged.species1.species2$logFC.y
## S = 1943300000, p-value = 0.03402
## alternative hypothesis: true rho is not equal to 0
## sample estimates:
##        rho
## 0.04418579
```

```
## Hippocampus

# Human -> Mouse
corr.HumanMouse.SingleGene.Hippo <- correlations.singlegene.level.aging( deg.hippo.human.aging,
        deg.hippo.mouse.aging, common.human.orthologs.one2one$ToMouse, "mmusculus_homolog_ensembl_gene"
```

```
## Number of common genes between the species after filtering: 12067
## Summary of Human Mouse
##
##  Spearman's rank correlation rho
##
## data:  merged.species1.species2$logFC.x and merged.species1.species2$logFC.y
## S = 2.3095e+11, p-value < 2.2e-16
## alternative hypothesis: true rho is not equal to 0
## sample estimates:
##       rho
## 0.2113838
```

```
# Human -> Fly
corr.HumanFly.SingleGene.Hippo <- correlations.singlegene.level.aging( deg.hippo.human.aging,
    deg.wholebody.fly.aging, common.human.orthologs.one2one$ToDmel, "dmelanogaster_homolog_ensembl_gene
```

```
## Number of common genes between the species after filtering: 2860
## Summary of Human Fly
##
##  Spearman's rank correlation rho
##
## data:  merged.species1.species2$logFC.x and merged.species1.species2$logFC.y
## S = 3727300000, p-value = 0.01852
## alternative hypothesis: true rho is not equal to 0
## sample estimates:
##        rho
## 0.04403345
```

```
# Human -> Worm
corr.HumanWorm.SingleGene.Hippo <- correlations.singlegene.level.aging( deg.hippo.human.aging,
             deg.wholebody.worm.aging, common.human.orthologs.one2one$ToCele, "celegans_homolog_ensembl_
```

```
## Number of common genes between the species after filtering: 2325
## Summary of Human Worm
##
##  Spearman's rank correlation rho
##
## data:  merged.species1.species2$logFC.x and merged.species1.species2$logFC.y
## S = 2080500000, p-value = 0.7439
## alternative hypothesis: true rho is not equal to 0
## sample estimates:
##         rho
## 0.006780104
```

```
## ------------------------------
# putting it under the cutoff of 0.05 to remove zeros

## cutoff-free

# skeletal muscle
```

```r
human.mouse.singlegenes.muscle <- defining.cutoff(corr.HumanMouse.SingleGene.Muscle, cutoff = 0.99)
human.fly.singlegenes.muscle <- defining.cutoff(corr.HumanFly.SingleGene.Muscle, cutoff = 0.99)
human.worm.singlegenes.muscle <- defining.cutoff(corr.HumanWorm.SingleGene.Muscle, cutoff = 0.99)


# hippocampus
human.mouse.singlegenes.hippo <- defining.cutoff(corr.HumanMouse.SingleGene.Hippo, cutoff = 0.99)
human.fly.singlegenes.hippo <- defining.cutoff(corr.HumanFly.SingleGene.Hippo, cutoff = 0.99)
human.worm.singlegenes.hippo <- defining.cutoff(corr.HumanWorm.SingleGene.Hippo, cutoff = 0.99)
```

```r
## ------------
# check the correlations after cutoff

## muscle
cutoff.corrs.human.mouse.muscle <- correlation.spearman(human.mouse.singlegenes.muscle)
```

```
##
##  Spearman's rank correlation rho
##
## data:  matrix.species$logFC.x and matrix.species$logFC.y
## S = 1.3246e+11, p-value = 0.0001374
## alternative hypothesis: true rho is not equal to 0
## sample estimates:
##         rho
## -0.03986407
```

```r
cutoff.corrs.human.fly.muscle <- correlation.spearman(human.fly.singlegenes.muscle)
```

```
##
##  Spearman's rank correlation rho
##
## data:  matrix.species$logFC.x and matrix.species$logFC.y
## S = 3276300000, p-value = 1.169e-05
## alternative hypothesis: true rho is not equal to 0
## sample estimates:
##        rho
## 0.08305571
```

```r
cutoff.corrs.human.worm.muscle <- correlation.spearman(human.worm.singlegenes.muscle)
```

```
##
##  Spearman's rank correlation rho
##
## data:  matrix.species$logFC.x and matrix.species$logFC.y
## S = 1852200000, p-value = 0.03269
## alternative hypothesis: true rho is not equal to 0
## sample estimates:
##        rho
## 0.04487102
```

```r
#### hippocampus
cutoff.corrs.human.mouse.hippo <- correlation.spearman(human.mouse.singlegenes.hippo)
```

```
##
##  Spearman's rank correlation rho
##
```

```
## data:  matrix.species$logFC.x and matrix.species$logFC.y
## S = 2.1832e+11, p-value < 2.2e-16
## alternative hypothesis: true rho is not equal to 0
## sample estimates:
##       rho
## 0.2135748
```

```
cutoff.corrs.human.fly.hippo <- correlation.spearman(human.fly.singlegenes.hippo)
```

```
##
##  Spearman's rank correlation rho
##
## data:  matrix.species$logFC.x and matrix.species$logFC.y
## S = 3588100000, p-value = 0.01918
## alternative hypothesis: true rho is not equal to 0
## sample estimates:
##        rho
## 0.04407119
```

```
cutoff.corrs.human.worm.hippo <- correlation.spearman(human.worm.singlegenes.hippo)
```

```
##
##  Spearman's rank correlation rho
##
## data:  matrix.species$logFC.x and matrix.species$logFC.y
## S = 1961400000, p-value = 0.7346
## alternative hypothesis: true rho is not equal to 0
## sample estimates:
##        rho
## 0.00710321
```

Saving the plots for the Supplementary Figure

```
# Human - Mouse
p1.muscle.cutoff <- plotting.singlegene(human.mouse.singlegenes.muscle,
             "log2 Fold-change Human", "log2 Fold-change Mouse", "indianred1", "red", "Skeletal Muscle")
pp1.muscle.cutoff <- labeling.plot.cutoff(cutoff.corrs.human.mouse.muscle, human.mouse.singlegenes.muscl
```

```
## Warning: Removed 3 rows containing missing values (geom_point).
```

```
p1.hippo.cutoff <- plotting.singlegene(human.mouse.singlegenes.hippo, "log2 Fold-change Human", "log2 Fo
pp1.hippo.cutoff <- labeling.plot.cutoff(cutoff.corrs.human.mouse.hippo, human.mouse.singlegenes.hippo,
```

```
## Warning: Removed 13 rows containing missing values (geom_point).
```

```
# Human - Fly - here it is ok
p2.muscle.cutoff <- plotting.singlegene(human.fly.singlegenes.muscle, "log2 Fold-change Human", "log2 Fo
pp2.muscle.cutoff <- labeling.plot.cutoff(cutoff.corrs.human.fly.muscle, human.fly.singlegenes.muscle, p
```

```
## Warning: Removed 1 rows containing missing values (geom_point).
```

```
p2.hippo.cutoff <- plotting.singlegene(human.fly.singlegenes.hippo,  "log2 Fold-change Human", "log2 Fol
pp2.hippo.cutoff <- labeling.plot.cutoff(cutoff.corrs.human.fly.hippo, human.fly.singlegenes.hippo, p2.h


# Human - Worm
p3.muscle.cutoff <- plotting.singlegene(human.worm.singlegenes.muscle,  "log2 Fold-change Human", "log2
pp3.muscle.cutoff <- labeling.plot.cutoff(cutoff.corrs.human.worm.muscle, human.worm.singlegenes.muscle
```

```r
p3.hippo.cutoff <- plotting.singlegene(human.worm.singlegenes.hippo,  "log2 Fold-change Human", "log2 Fo
pp3.hippo.cutoff <- labeling.plot.cutoff(cutoff.corrs.human.worm.hippo, human.worm.singlegenes.hippo, p3
```

```r
# pdf("~/Project1/manuscript_GSEA/results/FigureS3_Single_Gene_Only.pdf", 10, 7)
plot_grid(pp1.muscle.cutoff, pp1.hippo.cutoff,
                    pp2.muscle.cutoff, pp2.hippo.cutoff,
          pp3.muscle.cutoff, pp3.hippo.cutoff,
                    labels = c("A", "", "B", "", "C", ""), nrow = 3, align = "h")
```

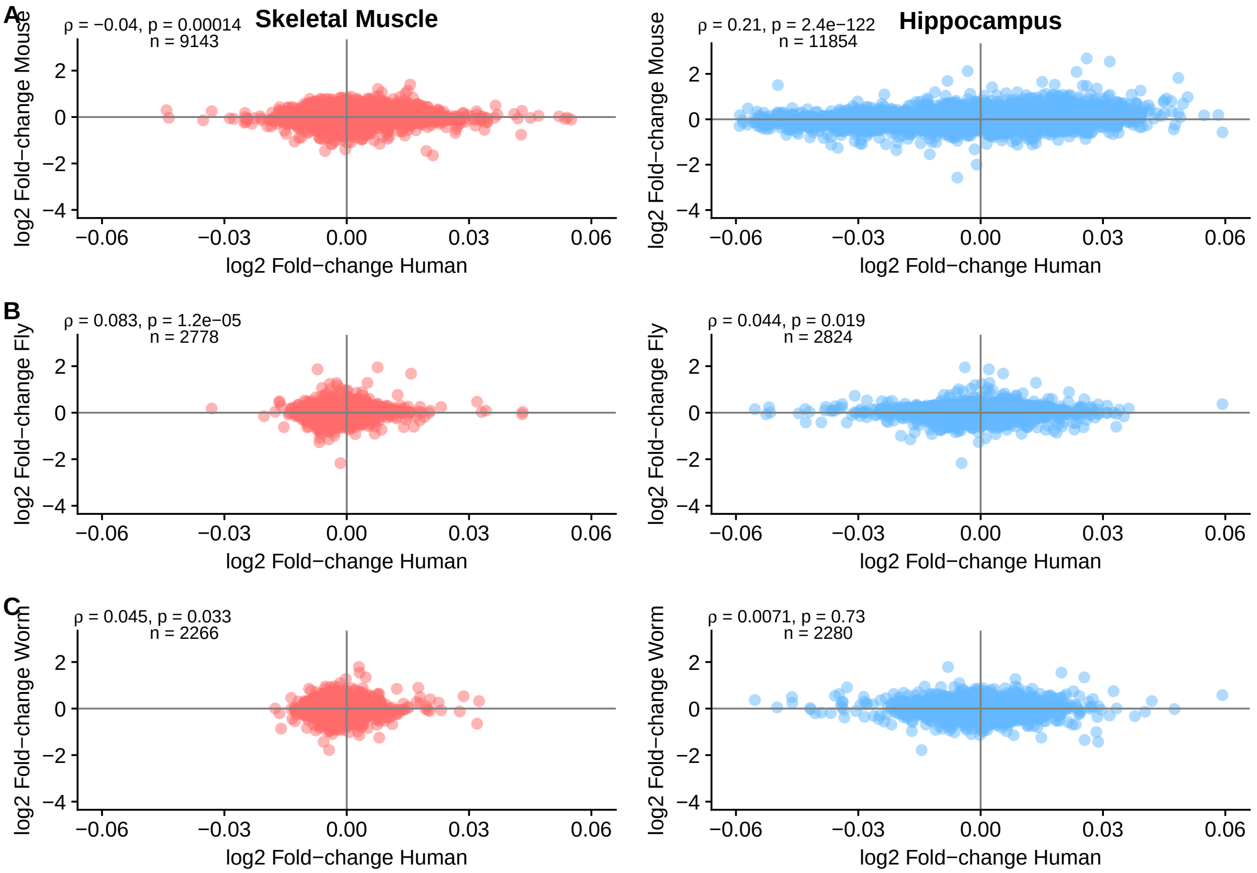


```r
# dev.off()
```

##### Single-species GSEA

Generating Figure S4.

```r
# functions for the analysis
go.gsea.analysis <- function(species, toptable){


  if(species == "Human"){
    library(org.Hs.eg.db)
    go.human <- go.gsets(species)
    go.human.sets <- go.human$go.sets
    go.human.subs <- go.human$go.subs

    cat("Getting GO categories.\n")
```

```r
    gobpsets.human <- go.human.sets[go.human.subs$BP]
    goccsets.human <- go.human.sets[go.human.subs$CC]
    gomfsets.human <- go.human.sets[go.human.subs$MF]

    toptable <- map.the.entrez(toptable, org.Hs.eg.db)
    tt.pc.order <- toptable[order(toptable$adj.P.Val),]
    gene.stats <- foldchange.tvalues(tt.pc.order)

    cat("Showing logFoldChanges and t-values.\n")
    print(lapply(gene.stats, head))

    fc <- fun.analysis.go(foldchangesstats = gene.stats[[1]],
                          tstats = NULL, gobpsets.human, gomfsets.human, goccsets.human)
    t <- fun.analysis.go(tstats = gene.stats[[2]],
                         foldchangesstats = NULL, gobpsets.human, gomfsets.human, goccsets.human)

  } else if(species == "Mouse") {

    library(org.Mm.eg.db)
    go.mouse <- go.gsets(species)
    go.mouse.sets <- go.mouse$go.sets
    go.mouse.subs <- go.mouse$go.subs

    cat("Getting GO categories.\n")
    gobpsets.mouse <- go.mouse.sets[go.mouse.subs$BP]
    goccsets.mouse <- go.mouse.sets[go.mouse.subs$CC]
    gomfsets.mouse <- go.mouse.sets[go.mouse.subs$MF]

    toptable <- map.the.entrez(toptable, org.Mm.eg.db)
    tt.pc.order <- toptable[order(toptable$adj.P.Val),]
    gene.stats <- foldchange.tvalues(tt.pc.order)

    cat("Showing logFoldChanges and t-values.\n")
    print(lapply(gene.stats, head))

    fc <- fun.analysis.go(foldchangesstats = gene.stats[[1]],
                          tstats = NULL, gobpsets.mouse, gomfsets.mouse, goccsets.mouse)
    t <- fun.analysis.go(tstats = gene.stats[[2]],
                         foldchangesstats = NULL, gobpsets.mouse, gomfsets.mouse, goccsets.mouse)


  } else if(species == "Worm") {
    library(org.Ce.eg.db)
    go.worm <- go.gsets(species)
    go.worm.sets <- go.worm$go.sets
    go.worm.subs <- go.worm$go.subs

    cat("Getting GO categories.\n")
    gobpsets.worm <- go.worm.sets[go.worm.subs$BP]
    goccsets.worm <- go.worm.sets[go.worm.subs$CC]
    gomfsets.worm <- go.worm.sets[go.worm.subs$MF]

    toptable <- map.the.entrez(toptable, org.Ce.eg.db)
```

```r
    tt.pc.order <- toptable[order(toptable$adj.P.Val),]
    gene.stats <- foldchange.tvalues(tt.pc.order)

    cat("Showing logFoldChanges and t-values.\n")
    print(lapply(gene.stats, head))

    fc <- fun.analysis.go(foldchangesstats = gene.stats[[1]],
                          tstats = NULL, gobpsets.worm, gomfsets.worm, goccsets.worm)
    t <- fun.analysis.go(tstats = gene.stats[[2]],
                         foldchangesstats = NULL, gobpsets.worm, gomfsets.worm, goccsets.worm)


  } else if(species == "Fly") {
    library(org.Dm.eg.db)
    go.fly <- go.gsets(species)
    go.fly.sets <- go.fly$go.sets
    go.fly.subs <- go.fly$go.subs

    cat("Getting GO categories.\n")
    gobpsets.fly <- go.fly.sets[go.fly.subs$BP]
    goccsets.fly <- go.fly.sets[go.fly.subs$CC]
    gomfsets.fly <- go.fly.sets[go.fly.subs$MF]

    toptable <- map.the.entrez(toptable, org.Dm.eg.db)
    tt.pc.order <- toptable[order(toptable$adj.P.Val),]
    gene.stats <- foldchange.tvalues(tt.pc.order)

    cat("Showing logFoldChanges and t-values.\n")
    print(lapply(gene.stats, head))

    fc <- fun.analysis.go(foldchangesstats = gene.stats[[1]],
                          tstats = NULL, gobpsets.fly, gomfsets.fly, goccsets.fly)
    t <- fun.analysis.go(tstats = gene.stats[[2]],
                         foldchangesstats = NULL, gobpsets.fly, gomfsets.fly, goccsets.fly)
  }

  return(list(foldchange.analysis = fc, tstat.analysis = t))
}


fun.analysis.go <- function(tstats = NULL, foldchangesstats = NULL,
                            gobpsets.species, gomfsets.species, goccsets.species){
  cat("Calculating the functional analysis.\n")

  if(length(tstats) > 1){
    cat("Calculating the functional analysis on t-values.\n")
    gobpres.healthy <- gage(tstats,  gsets=gobpsets.species, same.dir=TRUE, use.fold = FALSE, ref = NUL
    gomfres.healthy<- gage(tstats,   gsets=gomfsets.species, same.dir=TRUE, use.fold = FALSE, ref = NUL
    goccres.healthy <- gage(tstats,  gsets=goccsets.species, same.dir=TRUE, use.fold = FALSE, ref = NUL

  } else {

    cat("Calculating the functional analysis on logFC values.\n")
```

```
    gobpres.healthy <- gage(foldchangesstats,  gsets=gobpsets.species, same.dir=TRUE, ref = NULL, samp =
    gomfres.healthy <- gage(foldchangesstats,  gsets=gomfsets.species, same.dir=TRUE, ref = NULL, samp =
    goccres.healthy <- gage(foldchangesstats, gsets=goccsets.species, same.dir=TRUE, ref = NULL, samp =

  }

  return(list(gobpres.healthy, gomfres.healthy, goccres.healthy))

}

significant.go <- function(results.go.species) {
  results.go <- results.go.species[[2]]
  greater.go <- list()
  less.go <- list()
  for(i in 1:3){
    # this cutoff is because of the mouse data
    greater.go[[i]] <- results.go[[i]]$greater[results.go[[i]]$greater[,4] < 0.20, ]
    less.go[[i]] <- results.go[[i]]$less[results.go[[i]]$less[,4] < 0.20, ]
  }

  return(list(up.go = greater.go, down.go = less.go))

}


choose.go <- function(significant.gos, ont = "BP", number.of.sets = 20){

  if(ont == "BP"){
    ups <- significant.gos$up.go[[1]][1:number.of.sets, ]
    downs <- significant.gos$down.go[[1]][1:number.of.sets, ]
  } else if (ont == "MF"){
    ups <- significant.gos$up.go[[2]][1:number.of.sets, ]
    downs <- significant.gos$down.go[[2]][1:number.of.sets, ]
  } else {
    ups <- significant.gos$up.go[[3]][1:number.of.sets, ]
    downs <- significant.gos$down.go[[3]][1:number.of.sets, ]
  }

  df <- as.data.frame(rbind(ups, downs))
  df$Name <- rownames(df)
  return(df)

}


foldchange.tvalues <- function(toptable){
  foldchanges <- toptable$logFC
  names(foldchanges) <- toptable$entrez
  tvalues <- toptable$t
  names(tvalues) <- toptable$entrez
  return(list(foldchanges, tvalues))
}
```

```r
#### Species-specific analysis

# -----------------------------------------------------
# GSEA on GO ontology
# -----------------------------------------------------
library(gage)

# GSEA now
results.go.human.muscle <- go.gsea.analysis("Human", deg.muscle.human.aging)
```

```
## Gene ID type for 'human' is: 'EG'

##

## Getting GO categories.

## 'select()' returned 1:many mapping between keys and columns

## Showing logFoldChanges and t-values.
## [[1]]
##      221061       10655        8701       54102       55885      442117
## -0.02602730 -0.03311675  0.06730171 -0.04387005  0.04556766  0.06637557
##
## [[2]]
##    221061     10655      8701     54102     55885    442117
## -9.854277 -9.365321  8.469465 -8.434936  7.888115  7.697907
##
## Calculating the functional analysis.
## Calculating the functional analysis on logFC values.
## Calculating the functional analysis.
## Calculating the functional analysis on t-values.
```

```r
results.go.mouse.muscle <- go.gsea.analysis("Mouse", deg.muscle.mouse.aging)
```

```
## Gene ID type for 'mouse' is: 'EG'

## Getting GO categories.

## 'select()' returned 1:many mapping between keys and columns

## Showing logFoldChanges and t-values.
## [[1]]
##     118449      21912     110880      56495      66469      70510
## -1.1810740 -1.0756396 -1.4619566 -1.1449355 -0.7841583 -0.7509859
##
## [[2]]
##    118449     21912    110880     56495     66469     70510
## -16.08017 -15.36940 -14.24539 -14.00831 -13.64479 -13.40626
##
## Calculating the functional analysis.
## Calculating the functional analysis on logFC values.
## Calculating the functional analysis.
## Calculating the functional analysis on t-values.
```

```r
results.go.human.hippo <- go.gsea.analysis("Human", deg.hippo.human.aging)
```

```
## Gene ID type for 'human' is: 'EG'

## Getting GO categories.
```

```
## 'select()' returned 1:many mapping between keys and columns
## Showing logFoldChanges and t-values.
## [[1]]
##       27287      266722      115992       22843      285220         347
##  0.08460573 -0.06411874  0.01738765 -0.06430807 -0.05484216  0.03129872
##
## [[2]]
##     27287    266722    115992     22843    285220       347
##  7.086007 -6.782670  6.080744 -5.978037 -5.915660  5.910514
##
## Calculating the functional analysis.
## Calculating the functional analysis on logFC values.
## Calculating the functional analysis.
## Calculating the functional analysis on t-values.
```

```r
results.go.mouse.hippo <- go.gsea.analysis("Mouse", deg.hippo.mouse.aging)
```

```
## Gene ID type for 'mouse' is: 'EG'
## Getting GO categories.
## 'select()' returned 1:many mapping between keys and columns
## Showing logFoldChanges and t-values.
## [[1]]
##     12268      <NA>     78781     55990     93961     93880
## 2.2492003 1.6899556 1.0632543 2.6770648 0.5585991 1.0268640
##
## [[2]]
##    12268     <NA>    78781    55990    93961    93880
## 16.81314 15.17006 13.27444 12.97582 12.51056 12.39425
##
## Calculating the functional analysis.
## Calculating the functional analysis on logFC values.
## Calculating the functional analysis.
## Calculating the functional analysis on t-values.
```

```r
results.go.dmelano.wholebody <- go.gsea.analysis("Fly", deg.wholebody.fly.aging)
```

```
## Gene ID type for 'fly' is: 'EG'
## Getting GO categories.
## 'select()' returned 1:many mapping between keys and columns
## Showing logFoldChanges and t-values.
## [[1]]
##     59235     35409     36410     44921     39225     34800
## -2.731647 -2.248416 -4.169935  3.940811  2.146957 -1.578291
##
## [[2]]
##     59235     35409     36410     44921     39225     34800
## -41.34455 -35.32211 -30.91917  28.68678  26.37980 -26.30645
##
## Calculating the functional analysis.
## Calculating the functional analysis on logFC values.
## Calculating the functional analysis.
## Calculating the functional analysis on t-values.
```

```r
results.go.celegans.wholebody <- go.gsea.analysis("Worm",deg.wholebody.worm.aging)
```

```
## Gene ID type for 'worm' is: 'EG'
## Getting GO categories.
## 'select()' returned 1:many mapping between keys and columns
## Showing logFoldChanges and t-values.
## [[1]]
##    174652    178245    177447    178971    179804    184206
## -3.560177 -3.297889 -3.198386 -3.109974 -3.434989 -3.364338
##
## [[2]]
##    174652    178245    177447    178971    179804    184206
## -36.67322 -34.32467 -34.26308 -30.11533 -29.80401 -29.61442
##
## Calculating the functional analysis.
## Calculating the functional analysis on logFC values.
## Calculating the functional analysis.
## Calculating the functional analysis on t-values.
```

```r
# get significant ones - FDR: 0.10 - not possible
# up.go; down.go
signif.human.muscle <- significant.go(results.go.human.muscle)
signif.human.hippo <- significant.go(results.go.human.hippo)

signif.mouse.muscle <- significant.go(results.go.mouse.muscle)
signif.mouse.hippo <- significant.go(results.go.mouse.hippo)

signif.dmelano <- significant.go(results.go.dmelano.wholebody)
signif.celegans <- significant.go(results.go.celegans.wholebody)


### filter the gos - now it is without go semantics - top 20 categories
human.muscle.bp <- choose.go(signif.human.muscle)
human.hippo.bp <- choose.go(signif.human.hippo)

mouse.muscle.bp <- choose.go(signif.mouse.muscle)
#head(mouse.muscle.bp)
mouse.hippo.bp <- choose.go(signif.mouse.hippo)

dmelano.bp <- choose.go(signif.dmelano)
cele.bp <- choose.go(signif.celegans)
```

```r
# ------------------------------
# human
human.muscle.bp$GO.ID <- do.call(c, lapply(strsplit(human.muscle.bp$Name, " "), '[', 1))
human.muscle.bp$Annot <- c(rep("Extracellular matrix organization",2), rep("Cell adhesion", 2),
                          "Immune response", "Cell adhesion", rep("Angiogenesis",2), rep("Cell adhesion
                          rep("Immune response", 2),
                          rep("Cell adhesion",6), rep("Mitochondrial translation", 5), "Translation",
                          "Cellular respiration", rep("Catabolic process", 3),
                          rep("Metabolic process", 2), "Cellular protein modification process",
                          rep("Metabolic process", 4), "Cellular protein modification process",
                          "Metabolic process", "Cellular protein modification process")
```

```r
human.muscle.bp.new <- human.muscle.bp[order(human.muscle.bp$Annot), ]
human.muscle.bp.new$Annot <- factor(human.muscle.bp.new$Annot, levels=unique(human.muscle.bp.new$Annot)
human.muscle.bp.new$Name <- factor(human.muscle.bp.new$Name, levels=unique(human.muscle.bp.new$Name))


## human hippocampus
human.hippo.bp$GO.ID <- do.call(c, lapply(strsplit(human.hippo.bp$Name, " "), '[', 1))
human.hippo.bp$Annot <- c(rep("Immune response", 2),  "Viral process", rep("Protein transport", 2), "Vi
                          rep("Immune response", 2), "Protein targeting",
                          "Response to external stimulus", "Catabolic process",
                          rep("Immune response", 6), "Response to external stimulus",
                          "Extracellular matrix organization", "Immune response",
                          rep("Nervous system process", 3) , "Signaling",
                          rep("Nervous system process", 10), "Cellular localization",
                          "Nervous system process", "Cellular localization",
                          "Signaling", rep("Nervous system process",2))
human.hippo.bp.new <- human.hippo.bp[order(human.hippo.bp$Annot), ]
human.hippo.bp.new$Annot <- factor(human.hippo.bp.new$Annot, levels=unique(human.hippo.bp.new$Annot))
human.hippo.bp.new$Name <- factor(human.hippo.bp.new$Name, levels=unique(human.hippo.bp.new$Name))


# -------------------------------
# mouse muscle
mouse.muscle.bp$GO.ID <- do.call(c, lapply(strsplit(mouse.muscle.bp$Name, " "), '[', 1))
# check can you remove NA's
mouse.muscle.bp$Annot <- c(rep(NA, 10), rep(NA, 10), rep("Metabolic process", 10), rep(NA, 10))
mouse.muscle.bp.new <- mouse.muscle.bp[order(mouse.muscle.bp$Annot), ]
mouse.muscle.bp.new <- mouse.muscle.bp.new[complete.cases(mouse.muscle.bp.new$Annot), ]
mouse.muscle.colors <- c("Metabolic process" = "firebrick1")
mouse.muscle.bp.new$Annot <- factor(mouse.muscle.bp.new$Annot, levels=unique(mouse.muscle.bp.new$Annot)
mouse.muscle.bp.new$Name <- factor(mouse.muscle.bp.new$Name, levels=unique(mouse.muscle.bp.new$Name))


### hippocampus
mouse.hippo.bp$GO.ID <- do.call(c, lapply(strsplit(mouse.hippo.bp$Name, " "), '[', 1))
mouse.hippo.bp <- mouse.hippo.bp[complete.cases(mouse.hippo.bp),]
mouse.hippo.bp$Annot <- c(rep("Immune response", 20),
                          rep("Nervous system process", 7))
mouse.hippo.bp.new <- mouse.hippo.bp[order(mouse.hippo.bp$Annot), ]
mouse.hippo.bp.new$Annot <- factor(mouse.hippo.bp.new$Annot, levels=unique(mouse.hippo.bp.new$Annot))
mouse.hippo.bp.new$Name <- factor(mouse.hippo.bp.new$Name, levels=unique(mouse.hippo.bp.new$Name))


# -------------------------------
# dmelanogaster
dmelano.bp$GO.ID <- do.call(c, lapply(strsplit(dmelano.bp$Name, " "), '[', 1))
dmelano.bp$Annot <- c(rep("Immune response", 10), rep("Anatomical structure morphogenesis", 2),
                      rep("Signaling", 2), "Immune response",
                        rep("Anatomical structure morphogenesis", 5),
                      rep("Metabolic process", 7), rep("Cellular respiration", 6),
                      rep("Metabolic process", 7))
dmele.bp.new <- dmelano.bp[order(dmelano.bp$Annot), ]
```

```
dmele.bp.new$Annot <- factor(dmele.bp.new$Annot, levels=unique(dmele.bp.new$Annot))
dmele.bp.new$Name <- factor(dmele.bp.new$Name, levels=unique(dmele.bp.new$Name))


# -----------------------------
# celegans
# remove NAs
cele.bp <- cele.bp[complete.cases(cele.bp),]
cele.bp$GO.ID <- do.call(c, lapply(strsplit(cele.bp$Name, " "), '[', 1))
cele.bp$Annot <- c( rep("Cellular response to stimulus", 5), "Signaling",
                    rep("Cellular response to stimulus", 3), "Signaling",
                      rep("Cellular response to stimulus", 2),
                    rep("Signaling", 2), rep("Nervous system process", 2),
                    rep("Cell adhesion", 2),
                      "Nervous system process", "Cell adhesion",
                    rep("Metabolic process", 3), rep("Immune response", 4),
                    rep("Metabolic process", 7) )

cele.bp.new <- cele.bp[order(cele.bp$Annot), ]

cele.bp.new$Annot <- factor(cele.bp.new$Annot, levels=unique(cele.bp.new$Annot))
cele.bp.new$Name <- factor(cele.bp.new$Name, levels=unique(cele.bp.new$Name))

### check the colors
united.colors.of.genesets <- c("Cellular localization" = "springgreen3",
                "Cellular response to stimulus" = "mediumorchid1",
                "Signaling" = "tan1",
                "Translation" = "palegreen3",
                "Anatomical structure morphogenesis" = "yellow",
                #"Nucleoside metabolic process" = "tomato1",
                "Cellular protein modification process" = "green",
                "Cellular respiration" = "indianred1",
                "Immune response" = "deepskyblue3",
                "Nervous system process" = "goldenrod1",
                "Metabolic process" ="firebrick1",
                "Defense response" =  "dodgerblue",
                "Catabolic process" =  "darkred",
                "Cellular respiration" = "indianred1",
                "Immune response" = "deepskyblue3",
                "Mitochondrial translation" = "brown1",
                "Cofactor biosynthetic process" = "coral1",
                "Extracellular matrix organization" = "rosybrown",
                "Angiogenesis" = "moccasin",
                "Cell adhesion" = "mistyrose3",
                "Protein transport" = "sandybrown",
                "Protein targeting" = "khaki1",
                "Cellular localization" = "olivedrab",
                "Response to external stimulus" = "royalblue",
                "Viral process" = "grey30")
```

Plotting the GSEA results.

```
p1.human.muscle <- ggplot(human.muscle.bp.new, aes(stat.mean, Name)) +
                    geom_point(shape = 19, size =3, colour = "black") + geom_point(shape = 19, size = 2
```

```
                    aes(colour = Annot)) +
                    scale_colour_manual(values = united.colors.of.genesets)  +
                    ylab("GO BP categories") + xlab("GAGE stat. mean") +
                    theme(axis.text.y=element_blank(), legend.position = "right",
                          panel.grid.minor=element_line(color="ivory3",size=0.5),
                          panel.grid.major=element_line(color="ivory3",size=0.5),
                          legend.title=element_blank()) +
                    geom_vline(xintercept = 0) +
                    xlim(c(-10, 10))  +
                    labs(title = "Human Skeletal Muscle")


p1.human.hippo <- ggplot(human.hippo.bp.new, aes(stat.mean, Name)) +
                  geom_point(shape = 19, size =3, colour = "black") + geom_point(shape = 19, size = 2.5
                  aes(colour = Annot)) +
                  scale_colour_manual(values = united.colors.of.genesets)  +
                  ylab("GO BP categories") + xlab("GAGE stat. mean") +
                  theme(axis.text.y=element_blank(), legend.position = "right",
                        panel.grid.minor=element_line(color="ivory3",size=0.5),
                        panel.grid.major=element_line(color="ivory3",size=0.5),
                        legend.title=element_blank()) +
                  geom_vline(xintercept = 0) +
                  xlim(c(-10, 10))  +
                  labs(title = "Human Hippocampus")


p1.mouse.muscle <- ggplot(mouse.muscle.bp.new, aes(stat.mean, Name)) +
                   geom_point(shape = 19, size =3, colour = "black") + geom_point(shape = 19, size = 2.
                   aes(colour = Annot)) +
                   scale_colour_manual(values = united.colors.of.genesets)  +
                   ylab("GO BP categories") + xlab("GAGE stat. mean") +
                   theme(axis.text.y=element_blank(), legend.position = "right",
                         panel.grid.minor=element_line(color="ivory3",size=0.5),
                         panel.grid.major=element_line(color="ivory3",size=0.5),
                         legend.title=element_blank()) +
                   geom_vline(xintercept = 0) +
                   xlim(c(-10, 10))  +
                   labs(title = "Mouse Skeletal Muscle")


p1.mouse.hippo <- ggplot(mouse.hippo.bp.new, aes(stat.mean, Name)) +
                  geom_point(shape = 19, size =3, colour = "black") + geom_point(shape = 19, size = 2.5
                  aes(colour = Annot)) +
                  scale_colour_manual(values = united.colors.of.genesets)  +
                  ylab("GO BP categories") + xlab("GAGE stat. mean") +
                  theme(axis.text.y=element_blank(), legend.position = "right",
                        panel.grid.minor=element_line(color="ivory3",size=0.5),
                        panel.grid.major=element_line(color="ivory3",size=0.5),
                        legend.title=element_blank()) +
                  geom_vline(xintercept = 0) +
                  xlim(c(-10, 10))  +
                  labs(title = "Mouse Hippocampus")
```

```
p1.dmele <- ggplot(dmele.bp.new, aes(stat.mean, Name)) +
            geom_point(shape = 19, size =3, colour = "black") + geom_point(shape = 19, size = 2.5,
            aes(colour = Annot)) +
            scale_colour_manual(values = united.colors.of.genesets)  +
            ylab("GO BP categories") + xlab("GAGE stat. mean") +
            theme(axis.text.y=element_blank(), legend.position = "right",
                  panel.grid.minor=element_line(color="ivory3",size=0.5),
                  panel.grid.major=element_line(color="ivory3",size=0.5),
                  legend.title=element_blank()) +
            geom_vline(xintercept = 0) +
            xlim(c(-10, 10))  +
            labs(title = "D. melanogaster wholebody")


p1.cele <- ggplot(cele.bp.new, aes(stat.mean, Name)) +
           geom_point(shape = 19, size =3, colour = "black") + geom_point(shape = 19, size = 2.5,
           aes(colour = Annot)) +
           scale_colour_manual(values = united.colors.of.genesets)  +
           ylab("GO BP categories") + xlab("GAGE stat. mean") +
           theme(axis.text.y=element_blank(), legend.position = "right",
                 panel.grid.minor=element_line(color="ivory3",size=0.5),
                 panel.grid.major=element_line(color="ivory3",size=0.5),
                 legend.title=element_blank()) +
           geom_vline(xintercept = 0) +
           xlim(c(-10, 10))  +
           labs(title = "C. elegans wholebody")


### figure S4. for GSEA

# pdf("~/Project1/manuscript_GSEA/results/Figure_S4.pdf", 17, 13, useDingbats = FALSE)
plot_grid(p1.human.muscle, p1.human.hippo,
          p1.mouse.muscle, p1.mouse.hippo,
          p1.dmele, p1.cele,
          labels = c("A", "B", "C", "D", "E", "F"), nrow = 3, align = "h")
```

```
## Warning: Removed 2 rows containing missing values (geom_point).

## Warning: Removed 2 rows containing missing values (geom_point).
```

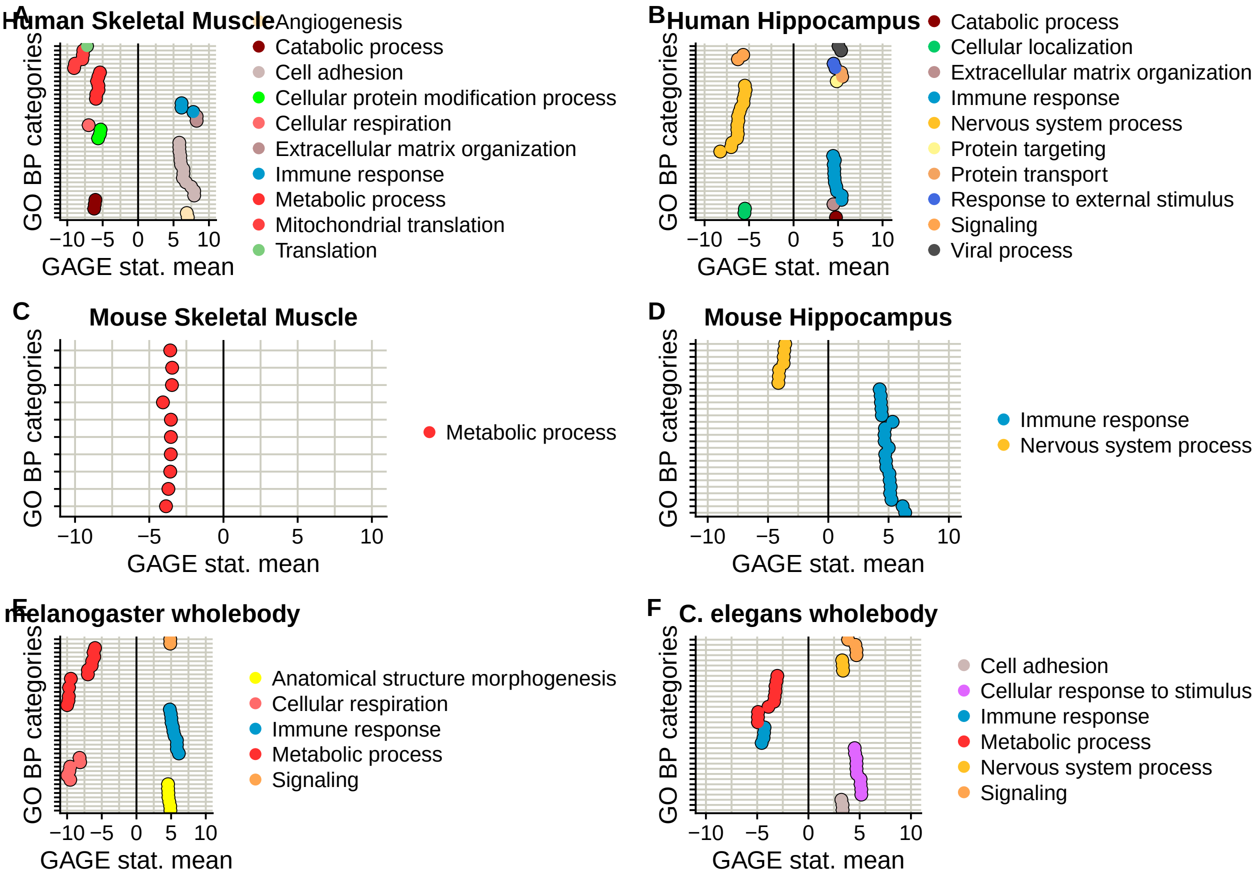
A
Human Skeletal Muscle
GO BP categories
−10 −5 0 5 10
GAGE stat. mean
Angiogenesis
Catabolic process
Cell adhesion
Cellular protein modification process
Cellular respiration
Extracellular matrix organization
Immune response
Metabolic process
Mitochondrial translation
Translation
B
Human Hippocampus
GO BP categories
−10 −5 0 5 10
GAGE stat. mean
Catabolic process
Cellular localization
Extracellular matrix organization
Immune response
Nervous system process
Protein targeting
Protein transport
Response to external stimulus
Signaling
Viral process
C
Mouse Skeletal Muscle
GO BP categories
−10 −5 0 5 10
GAGE stat. mean
Metabolic process
D
Mouse Hippocampus
GO BP categories
−10 −5 0 5 10
GAGE stat. mean
Immune response
Nervous system process
E
melanogaster wholebody
GO BP categories
−10 −5 0 5 10
GAGE stat. mean
Anatomical structure morphogenesis
Cellular respiration
Immune response
Metabolic process
Signaling
F
C. elegans wholebody
GO BP categories
−10 −5 0 5 10
GAGE stat. mean
Cell adhesion
Cellular response to stimulus
Immune response
Metabolic process
Nervous system process
Signaling

```
# dev.off()
```

### Main Figures + Supplement (Differential expression part)

*Andrea Komljenović*

*3/9/2018*

```
## --------------------------------
# Load the packages

packages <- c( "affy", "affyPLM", "biomaRt", "limma",
               "tidyr", "gplots", "ggplot2", "reshape2",
               "Biobase", "downloader", "sva", "dplyr", "RColorBrewer", "cowplot",
               "igraph", "WriteXLS", "org.Hs.eg.db", "org.Mm.eg.db", "org.Dm.eg.db", "org.Ce.eg.db", "gag
               "cowplot", "ggrepel", "plyr", "data.table", "dplyr", "ComplexHeatmap", "circlize", "RColo
lapply(packages, library, character.only = TRUE)
```

```
path <- "~/Project1/manuscript_GSEA/data_preprocessing/"
```

Loading differentially expressed matrices.

```
# --------------------------
# Human
deg.muscle.human.aging <-
  readRDS(paste0(path, "hsapiens/aging_data/diff_exp/topTable_Muscle_Skeletal_GTEx_V6p.rds"))
deg.hippo.human.aging <-
  readRDS(paste0(path, "hsapiens/aging_data/diff_exp/topTable_Brain_Hippocampus_GTEx_V6p.rds"))


# ---------------------------
# Mouse

# name it correctly + consistency - TO DO
deg.mouse.muscle <-
  readRDS(paste0(path, "mmusculus/aging_data/skeletal_muscle/diffexp_aging_mouse_skeletal_muscle_aging.r
deg.muscle.mouse.aging <- deg.mouse.muscle$oldvsyoung

# change here
deg.hippo.mouse.aging <- readRDS(paste0(path, "mmusculus/aging_data/hippocampus/diff_expression_mmusculu

# ---------------------------
# Fly - wb
deg.fly <-
  readRDS(paste0(path, "dmelanogaster/aging_data/diff_expression_fly_aging.rds"))
deg.wholebody.fly.aging <- deg.fly$oldvsyoung

# ---------------------------
# Worm - wb
deg.wholebody.worm.aging <-
  readRDS(paste0(path, "celegans/aging_data/diff_expression_celegans_aging.rds"))
```

```
## Dietary restriction
deg.human.dr <-
```

```
  readRDS(paste0(path, "hsapiens/caloric_restriction_data/differential_expression_human_dietary_restric
## Dietary restriction
deg.mouse.dr <-
  readRDS(paste0(path, "mmusculus/caloric_restriction_data/differential_expression_mouse_dietary_restri
## Dietary restriction - rename this
deg.fly.dr <-
  readRDS(paste0(path, "/dmelanogaster/caloric_restriction_data/diff_exp_fly_dietary_restriction.rds"))
## Dietary restriction
deg.worm.dr <-
  readRDS(paste0(path, "/celegans/caloric_restriction_data/diff_exp_worm_dietary_restriction.rds"))
```

##### Barplots of the differential expression analysis

```
# -----
# FUNCTION
number.of.degs <- function(toptable, cutoff){
    up <- toptable[which(sign(toptable$logFC) == 1 & toptable$adj.P.Val < cutoff),]
    dn <- toptable[which(sign(toptable$logFC) == -1 & toptable$adj.P.Val < cutoff),]
    cat("The number of genes that are downregulated:", dim(dn)[1], "\n")
    cat("The number of genes that are upregulated:", dim(up)[1], "\n")
  #  cat("The number of genes that are downregulated:", dim(dn)[1], "\n")

    return(list(downregulated = dim(dn)[1], upregulated = dim(up)[1]))
}




## ----------------
## preparing the datasets for plotting
list.degs.species <- list(human.muscle = deg.muscle.human.aging,
                          human.hippo = deg.hippo.human.aging,
                          mouse.muscle = deg.muscle.mouse.aging,
                          mouse.hippo = deg.hippo.mouse.aging,
                          fly.wholebody = deg.wholebody.fly.aging,
                          worm.wholebody = deg.wholebody.worm.aging)

signif.expressed.genes <- lapply(list.degs.species, function(x) number.of.degs(x, 0.1))
```

```
## The number of genes that are downregulated: 2540
## The number of genes that are upregulated: 2513
## The number of genes that are downregulated: 2978
## The number of genes that are upregulated: 3105
## The number of genes that are downregulated: 1271
## The number of genes that are upregulated: 1184
## The number of genes that are downregulated: 718
## The number of genes that are upregulated: 921
## The number of genes that are downregulated: 2344
## The number of genes that are upregulated: 2413
## The number of genes that are downregulated: 1634
## The number of genes that are upregulated: 1904
```

```
vec <- unlist(signif.expressed.genes)
ind <- seq(1, length(vec), by = 2)
```

```r
vec[ind] <- vec[ind]*(-1) # to give negative sign for downregulated ones
names(vec) <- ""

# plotting histogram
rnaseq <- data.frame(
    Dataset = c(
    rep("H.sapiens - Skeletal Muscle", 2), rep("H.sapiens - Hippocampus", 2),
  rep("M.musculus - Skeletal Muscle", 2), rep("M.musculus - Hippocampus", 2),
  rep("D.melanogaster - Whole Body", 2),
  rep("C.elegans - Whole Body", 2)),
    Status = c(rep(c("Downregulated","Upregulated"), 6)),
  # differentially expressed genes
    deg = vec)

# make V1 an ordered factor
rnaseq$Dataset <- factor(rnaseq$Dataset, levels = unique(rnaseq$Dataset))

# title <- "Aging"

# pdf("~/Project1/manuscript_GSEA/results/Figure1C_barplot_differential_exp_update.pdf", 7, 3)
ggplot(rnaseq, aes(as.factor(Dataset), deg, fill = Status)) +
  geom_bar(position= "identity", colour="grey50", stat="identity", width=0.8) +
  coord_flip() + scale_x_discrete(limits = rev(levels(rnaseq$Dataset))) +
  labs(x = "",
      y = "Number of detected differentially expressed genes") +
  scale_fill_brewer(type="qual", palette="Pastel1") +
  theme_bw() +
  theme(
    panel.grid.major = element_blank(),
    panel.grid.minor = element_blank(),
    panel.border = element_blank(),
    panel.background = element_blank(),
    axis.ticks.y=element_blank(),
    legend.title=element_blank(),axis.text.x=element_blank(),
    axis.ticks.x=element_blank()) +
    geom_hline(yintercept=0) +
    geom_text(label = abs(rnaseq$deg),
                                     hjust = "center",
                                     vjust = "bottom")
```

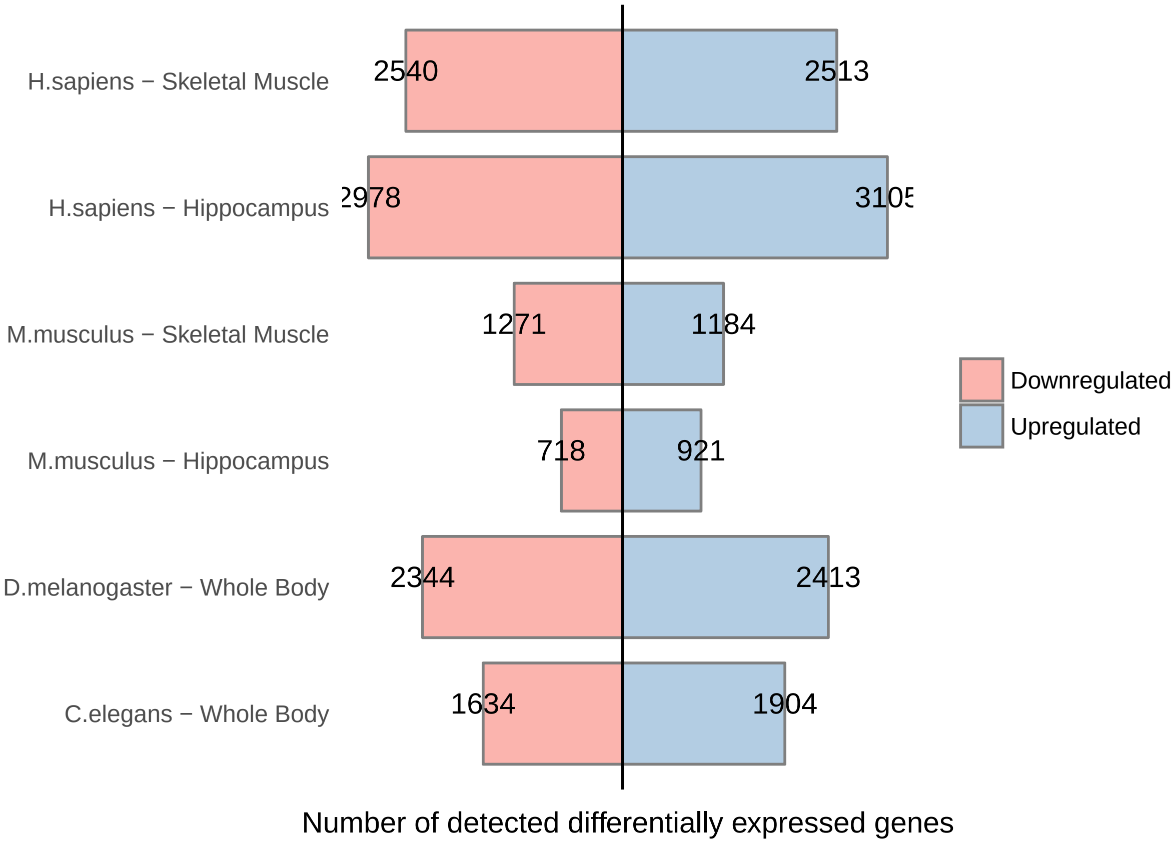


```
# dev.off()
```

##### Alignment of the stages between species

Generating Fig S2. heatmaps of clustering the young and old samples for human and other species, in order to do age alignments.

```
### defining the orthology
orthology.to.human.pc <- function(mart.human){
  # library(biomaRt)
        orth.mouse <-
          getBM(attributes = c("ensembl_gene_id", "external_gene_name", "mmusculus_homolog_ensembl_gene
        "mmusculus_homolog_orthology_type"), filters = "with_mmusculus_homolog", values= TRUE,
                    mart = mart.human, bmHeader = FALSE, uniqueRows = TRUE)

    orth.drosophila <-
      getBM(attributes = c("ensembl_gene_id", "external_gene_name", "dmelanogaster_homolog_ensembl_gene
        "dmelanogaster_homolog_orthology_type"), filters = "with_dmelanogaster_homolog", values= TRUE,
                    mart = mart.human, bmHeader = FALSE, uniqueRows = TRUE)

    orth.celegans <-
      getBM(attributes = c("ensembl_gene_id", "external_gene_name", "celegans_homolog_ensembl_gene",
        "celegans_homolog_orthology_type"), filters = "with_celegans_homolog", values= TRUE,
                    mart = mart.human, bmHeader = FALSE, uniqueRows = TRUE)

    protein.coding <-
```

```
      getBM(attributes = c("ensembl_gene_id", "gene_biotype"),
            values = TRUE, mart = mart.human,  bmHeader = FALSE, uniqueRows = TRUE)

    orth.mouse <- merge(orth.mouse, protein.coding, by = "ensembl_gene_id")
    orth.drosophila <- merge(orth.drosophila, protein.coding, by = "ensembl_gene_id")
    orth.celegans <- merge(orth.celegans, protein.coding, by = "ensembl_gene_id")
        return(list(ToMouse = orth.mouse, ToDmel = orth.drosophila, ToCele = orth.celegans))
}


### for selecting the  ages
select.ages.gtex <- function(path.annotation.file, expression){

  require(data.table)

  subject.phenotypes <-
    as.data.frame(fread(paste0(path.annotation.file,
    "phs000424.v6.pht002742.v6.p1.c1.GTEx_Subject_Phenotypes.GRU.txt"), header = TRUE))
    colnames(subject.phenotypes) <- as.vector(as.matrix(subject.phenotypes[1,]))
    subject.phenotypes <- subject.phenotypes[-1,]

    ages <- as.character(subject.phenotypes$AGE);
    names(ages) <- subject.phenotypes$SUBJID
    # 20-30 years; 61-70 years

    # this removes exact ages
    young.gtex <- rep("young", length(ages[which(ages >= "20" & ages <= "30")]))
    names(young.gtex) <- names(ages[which(ages >= "20" & ages <= "30")])

    old.gtex <- rep("old", length(ages[which(ages >= "61" & ages <= "70")]))
    names(old.gtex) <- names(ages[which(ages >= "61" & ages <= "70")])

    selected.ages.gtex <- c( young.gtex, old.gtex )

    expression.human <-
      expression[, na.omit(match( names(selected.ages.gtex), colnames(expression) ))]

    selected.ages.gtex.tissue <-
      selected.ages.gtex[na.omit(match(colnames(expression.human),names(selected.ages.gtex)))]
    colnames(expression.human) <-
      selected.ages.gtex.tissue

    return(expression.human)
}



select.signif.genes <- function( diff.expression.human, diff.expression.sp2,
                                 common.orthologs.human.sp2,
                                 ensembl.sp2.gnames, cutoff = NULL ) {
    # here it is always 1-1 orthologs
    # human
    significant.human <-
```

```r
    diff.expression.human[diff.expression.human$logFC > 0 &
                            diff.expression.human$adj.P.Val < cutoff,]
  # species2
  significant.sp2 <-
    diff.expression.sp2[diff.expression.sp2$logFC > 0 &
                          diff.expression.sp2$adj.P.Val < cutoff,]
  msig1 <- merge(common.orthologs.human.sp2, significant.sp2,
                 by.x = ensembl.sp2.gnames, by.y = "row.names" )
  msig2 <- merge(msig1, significant.human,
                 by.x = "ensembl_gene_id", by.y = "row.names" )

  return(msig2)
}


# for supplement data
select.signif.genes.sp <- function( diff.expression.human, diff.expression.sp2,
                                     common.orthologs.human.sp2,
                                    ensembl.sp2.gnames, cutoff = NULL, tissue.name ) {

  # here it is always 1-1 orthologs

 # human
  significant.human <- diff.expression.human[diff.expression.human$adj.P.Val < cutoff,]
 cat("Number of significant DEGs in human dataset:", nrow(significant.human), "\n")
  # species2
  significant.sp2 <- diff.expression.sp2[diff.expression.sp2$adj.P.Val < cutoff,]
 cat("Number of significant DEGs in mouse dataset:", nrow(significant.sp2), "\n")


 deg.number <- paste(nrow(significant.human), nrow(significant.sp2), sep = "/")

  msig1 <-
    merge(common.orthologs.human.sp2,
                 significant.sp2, by.x = ensembl.sp2.gnames, by.y = "row.names" )
  msig2 <-
    merge(msig1, significant.human,
          by.x = "ensembl_gene_id", by.y = "row.names" )
 cat("Number of common DEGs based on 1-1 orthologs:", nrow(msig2), "\n")
 # calculate overlap
 overlap.percentage <- (nrow(msig2)/nrow(significant.human))*100
 df <- data.frame(Tissue = tissue.name, DEGnumber = deg.number,
                 OneToOneOrthologsNumber = nrow(common.orthologs.human.sp2),
                 Overlap.percentage = overlap.percentage)
  return(df)
}

# the colnames of expression matrices should be named as they wanted to be shown in heatmap
clustering.samples <- function( merged.significant.genes, expression.human,
                                expression.sp2, ensembl.sp2.gnames,
  filename, heatmap.title, plot = FALSE ){

  library(sva)
```

```
    colnames.human <- colnames(expression.human)
    colnames.sp2 <- colnames(expression.sp2)

    batch <- c(rep("1", ncol(expression.human)), rep("2", ncol(expression.sp2)))
    modcombat <- model.matrix(~1, data=as.data.frame(batch))


    mm <-
      merge(merged.significant.genes, expression.human,
            by.x = "ensembl_gene_id", by.y = "row.names")
    mm2 <- merge(mm, expression.sp2,
            by.x = ensembl.sp2.gnames, by.y = "row.names")
    # remove the diff. exp results
    mm2 <- mm2[, -c(1:17)]
    colnames(mm2) <- c(colnames.human, colnames.sp2)

    combat.expression.data <- ComBat(dat = mm2,
                                     batch = batch,
                                     mod = modcombat,
                                     par.prior = TRUE, prior.plots = FALSE)
    # there is few outliers but looks ok

  hclust.comp <- function(x) hclust(x, method="complete")

  if(plot) {
    pdf(filename, 15, 5)
        hclust.comp <- function(x) hclust(x, method="complete")
        gplots::heatmap.2(as.matrix(combat.expression.data), scale = "row",
                                            labRow = FALSE,
                                            col=brewer.pal(11,"RdBu"), trace="none",
                                            margins =c(12,9), hclustfun=hclust.comp,
                                            main = heatmap.title)
        dev.off()

  } else {

    gplots::heatmap.2(as.matrix(combat.expression.data), scale = "row",
                      labRow = FALSE,
                      col=brewer.pal(11,"RdBu"), trace="none",
                      margins =c(12,9), hclustfun=hclust.comp,
                      main = heatmap.title)

  }

    return(combat.expression.data)

}
```

Load the expression matrices:

```
##### ------------------------------------------------------------------
## Load expression data

# ----------------------
```

```r
# human
exp.human.hippo <-
  readRDS(paste0(path, "/hsapiens/aging_data/exp_mat/voom_Brain_Hippocampus_GTEx_V6p.rds"))
exp.human.muscle <-
  readRDS(paste0(path, "hsapiens/aging_data/exp_mat/voom_Muscle_Skeletal_GTEx_V6p.rds"))


# for gtex data, selected only the youngest and the oldest samples (extremes) to show the alignments
expression.human.hippo <-
  select.ages.gtex("~/Documents/Lausanne/GTEx_annotation/", exp.human.hippo)
colnames(expression.human.hippo) <-
  paste0("Hs.hippo.", colnames(expression.human.hippo)) # 41

expression.human.muscle <-
  select.ages.gtex("~/Documents/Lausanne/GTEx_annotation/", exp.human.muscle)
colnames(expression.human.muscle) <-
  paste0("Hs.muscle.", colnames(expression.human.muscle)) # 142


# -------------------------
# mouse
expression.mouse.muscle <-
  readRDS(paste0(path, "/mmusculus/aging_data/skeletal_muscle/expression_matrix_mouse_skeletal_muscle_a
expression.mouse.hippo <-
  readRDS(paste0(path, "mmusculus/aging_data/hippocampus/voom_mouse_expression_matrix_hippocampus_aging

colnames(expression.mouse.muscle) <- c(rep("young",4),rep("old", 5))
colnames(expression.mouse.muscle) <- paste0("Mm.muscle.", colnames(expression.mouse.muscle))
colnames(expression.mouse.hippo) <- paste0("Mm.hippo.", colnames(expression.mouse.hippo))


# ------------------------
# fly
expression.microarray.fly <- readRDS(paste0(path, "/dmelanogaster/aging_data/expression_matrix_fly_agin
expression.microarray.fly <- expression.microarray.fly[[2]]
colnames(expression.microarray.fly) <- paste0("Dm.wb.", colnames(expression.microarray.fly))



# -----------------------
# worm
expression.wb.worm <-
  readRDS(paste0(path, "/celegans/aging_data/voom_celegans_expression_matrix_aging.rds"))
expression.wb.worm <- expression.wb.worm[, !(colnames(expression.wb.worm) %in% "too_young")]
colnames(expression.wb.worm) <- paste0("Ce.wb.", colnames(expression.wb.worm))
```

Define 1-1 orthologs between species:

```r
library(biomaRt)
human.ensembl <- useMart(biomart="ENSEMBL_MART_ENSEMBL",
                         host="www.ensembl.org",
                         path="/biomart/martservice",
                         dataset="hsapiens_gene_ensembl",
                         version = "Ensembl Genes 91")
```

```
# orthology to human on protein coding genes only
common.human.orthologs.pc <- orthology.to.human.pc(human.ensembl)
common.human.orthologs.final.pc <- lapply(common.human.orthologs.pc,
                                    function(x) x[which(x[,4]== "ortholog_one2one"),])
common.human.orthologs.one2one <- lapply(common.human.orthologs.final.pc,
                                   function(x) x[which(x[,5]== "protein_coding"),])
```

Plotting the heatmap for Human-Mouse for Figure 2A.

```
# 52 genes
signif.muscle <- select.signif.genes(deg.muscle.human.aging, deg.muscle.mouse.aging,
                                     common.human.orthologs.one2one$[illegible]
                                     "mmusculus_homolog_ensembl_gene[illegible]



# 94 genes
signif.hippo <- select.signif.genes(deg.hippo.human.aging, deg.hippo.mouse.aging,
                                     common.human.orthologs.one2[illegible]
                                     "mmusculus_homolog_ensembl_[illegible]




clusters.muscle.human.mouse <- clustering.samples( signif.muscle,
                               expression.human.muscle, expression.mouse.muscle,
                                    "mmusculus_homolog_ensembl_gene",
                                    "~/Project1/manuscript_GSEA/results/Figure[illegible]
                                    "Human - Mouse skeletal muscle", plot = FA[illegible]
```

```
## Found2batches

## Adjusting for0covariate(s) or covariate level(s)

## Standardizing Data across genes

## Fitting L/S model and finding priors

## Finding parametric adjustments

## Adjusting the Data
```

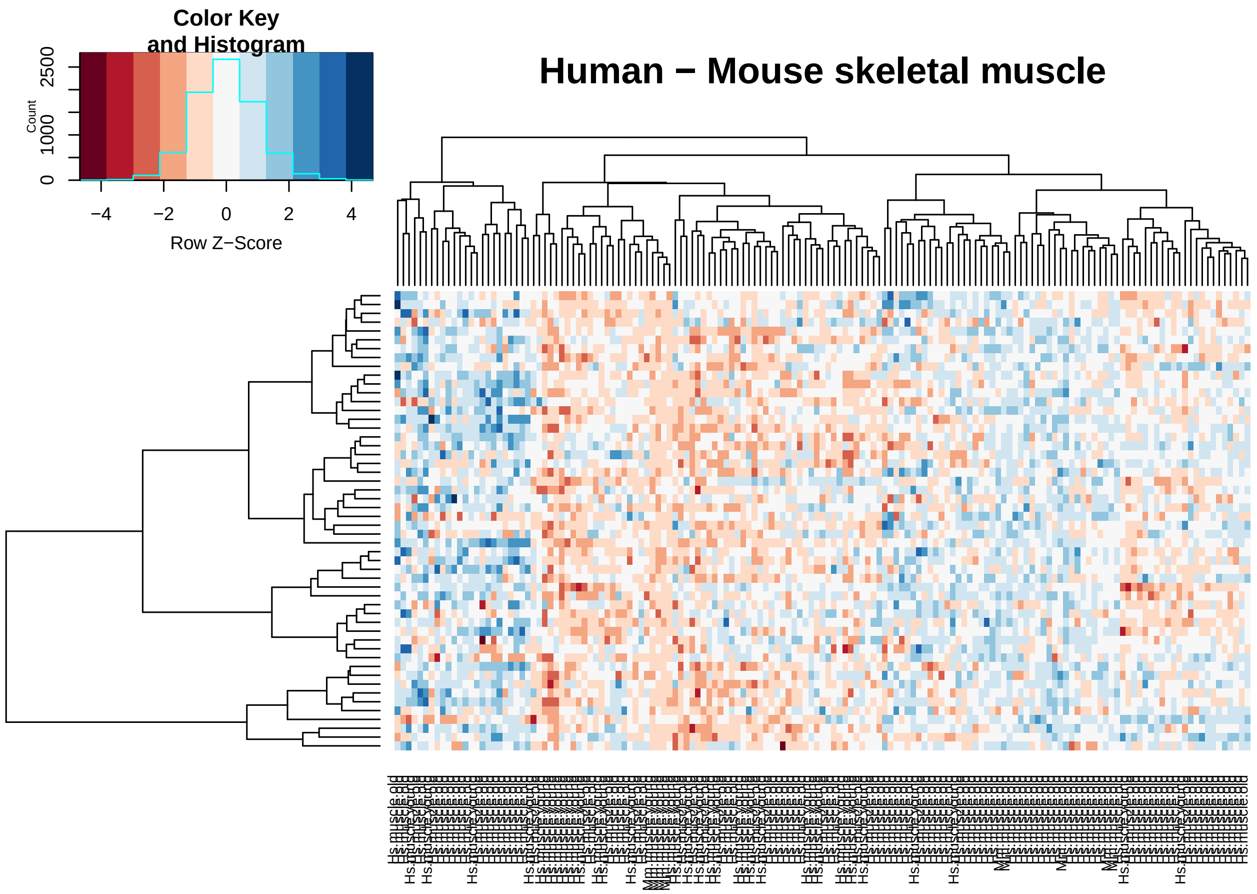


```
clusters.hippo.human.mouse <- clustering.samples( signif.hippo,
                                                   expression.human.hippo, expression.mouse.hippo,
                                                             "mmusculus_homolog_ensembl_gene",
                                                   "~/Project1/manuscript_GSEA/results/Figure2A_heatmap_I
                                                             "Human - Mouse hippocampus", plot = FALSE)
```

```
## Found2batches
## Adjusting for0covariate(s) or covariate level(s)
## Standardizing Data across genes
## Fitting L/S model and finding priors
## Finding parametric adjustments
## Adjusting the Data
```

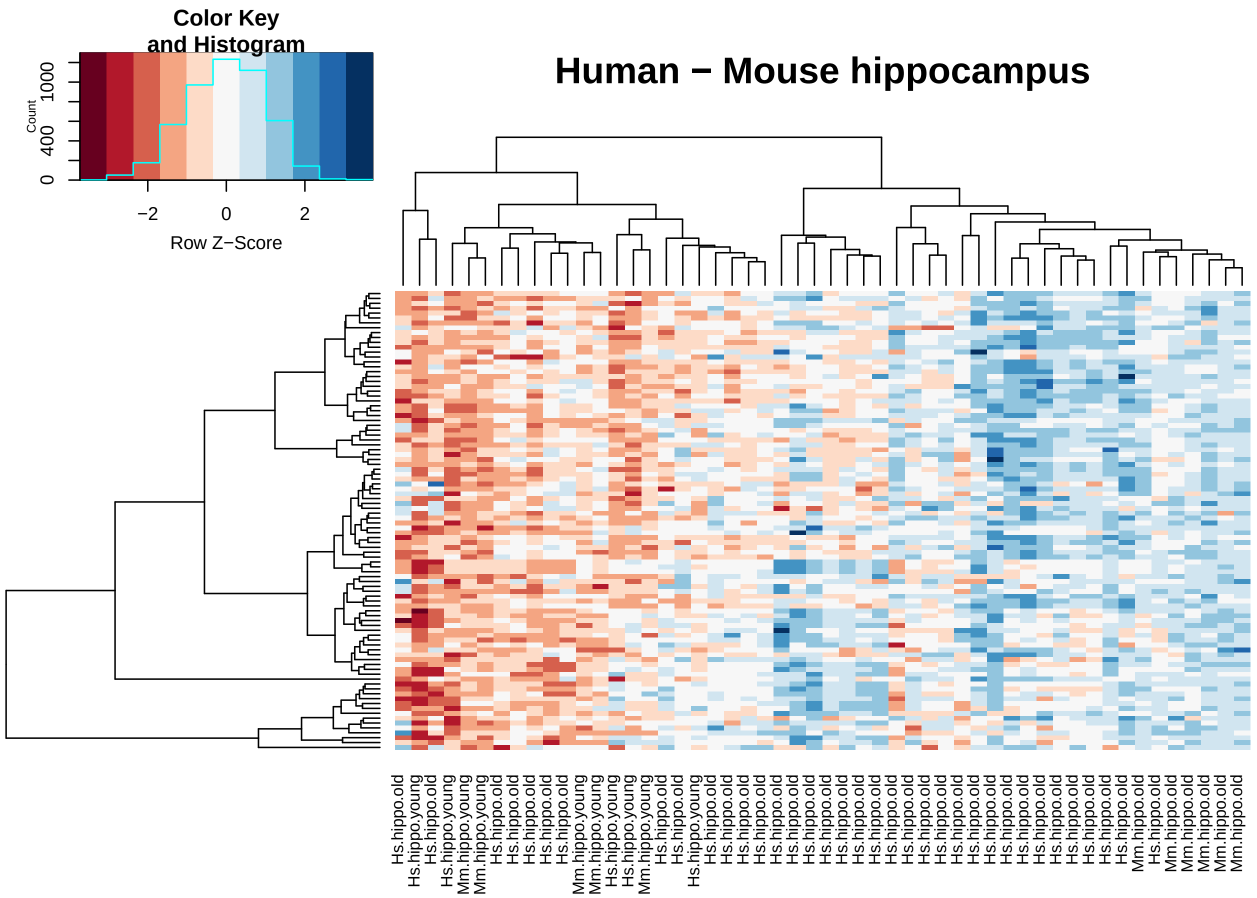


Plotting for Supplement Figures S2: Human-Fly, Human-Worm:

```
# 42
signif.muscle.human.fly <- select.signif.genes(deg.muscle.human.aging, deg.wholebody.fly.aging,
                                                common.human.orthologs.one2one$ToDmel,
                                                "dmelanogaster_homolog_ensembl_gene", cutoff = 0.05)

# 87
signif.hippo.human.fly <- select.signif.genes(deg.hippo.human.aging, deg.wholebody.fly.aging,
                                               common.human.orthologs.one2one$ToDmel,
                                               "dmelanogaster_homolog_ensembl_gene", cutoff = 0.05)

clusters.human.fly.muscle <- clustering.samples( signif.muscle.human.fly, expression.human.muscle, expre
                                                 "dmelanogaster_homolog_ensembl_gene",
                                                 "~/Project1/manuscript_GSEA/results/FigureS2_heatmap_hu
                                                 "Human muscle - Fly whole body", plot = FALSE)
```

```
## Found2batches

## Adjusting for0covariate(s) or covariate level(s)

## Standardizing Data across genes

## Fitting L/S model and finding priors

## Finding parametric adjustments

## Adjusting the Data
```

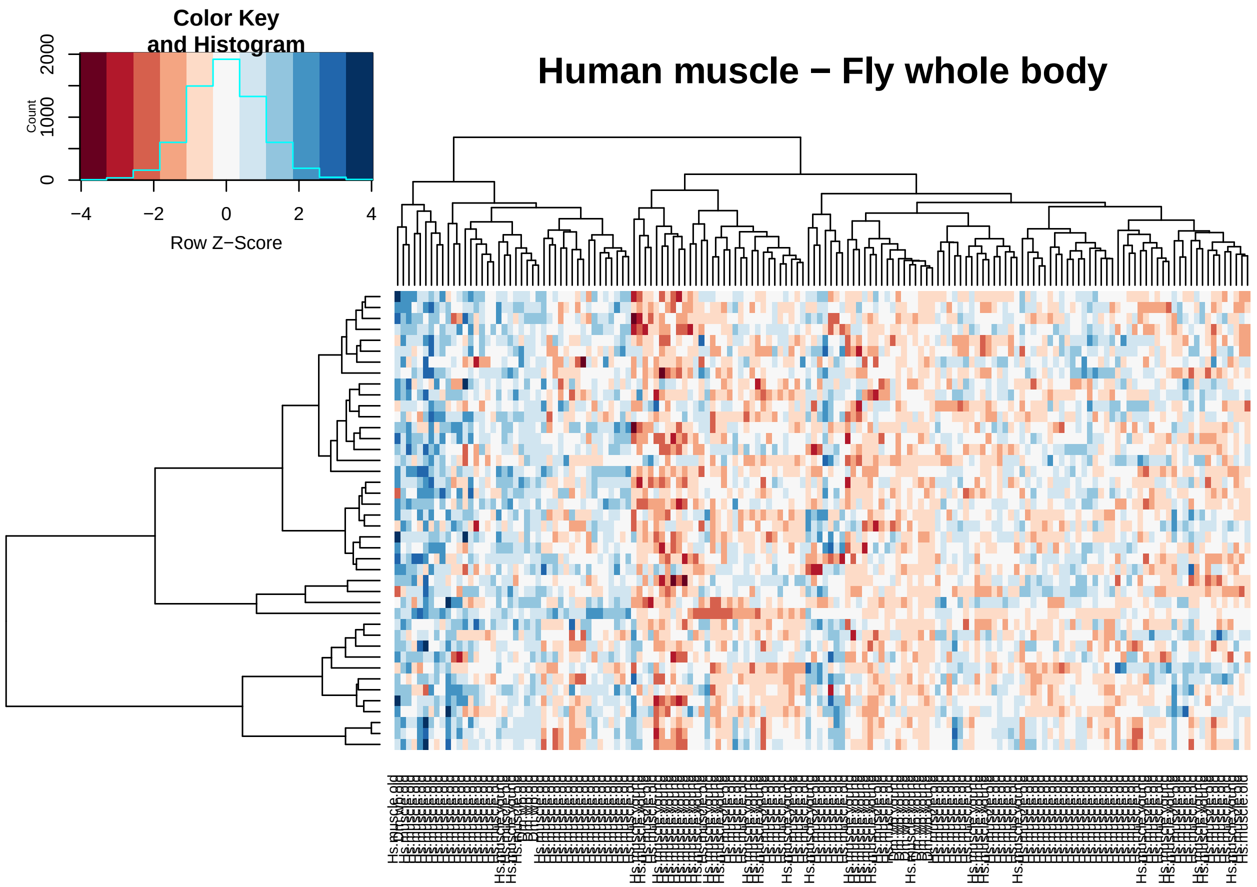


```
clusters.human.fly.hippo <- clustering.samples( signif.hippo.human.fly , expression.human.hippo, express
                                              "dmelanogaster_homolog_ensembl_gene",
                                              "~/Project1/manuscript_GSEA/results/FigureS2_heatmap_hu
                                              "Human hippocampus - Fly whole body", plot = FALSE)
```

```
## Found2batches
## Adjusting for0covariate(s) or covariate level(s)
## Standardizing Data across genes
## Fitting L/S model and finding priors
## Finding parametric adjustments
## Adjusting the Data
```

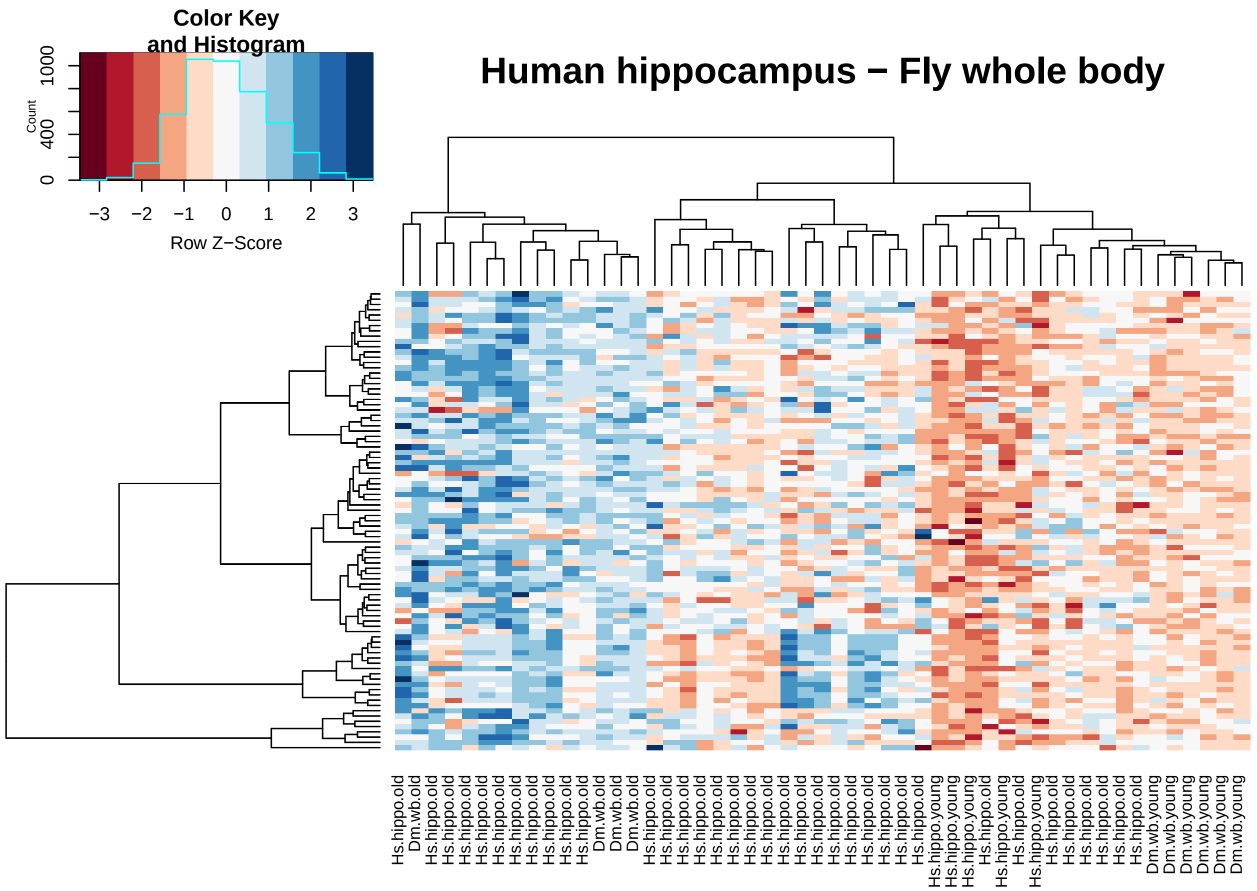


```
## here I had to change to the 0.05 because I get only 2 genes when cutoff 0.01

# 10
signif.muscle.human.worm <- select.signif.genes(deg.muscle.human.aging, deg.wholebody.worm.aging,
                                                             common.human.orthologs.one2one$ToCele,

# 17
signif.hippo.human.worm <- select.signif.genes(deg.hippo.human.aging, deg.wholebody.worm.aging,
                                                             common.human.orthologs.one2one$ToCele, "


clusters.human.worm.muscle <- clustering.samples( signif.muscle.human.worm, expression.human.muscle, ex
                                                  "celegans_homolog_ensembl_gene",
                                                  "~/Project1/manuscript_GSEA/results/FigureS2_heatmap_
                                                  "Human muscle - Worm whole body", plot = FALSE)
```

```
## Found2batches

## Adjusting for0covariate(s) or covariate level(s)

## Standardizing Data across genes

## Fitting L/S model and finding priors

## Finding parametric adjustments

## Adjusting the Data
```

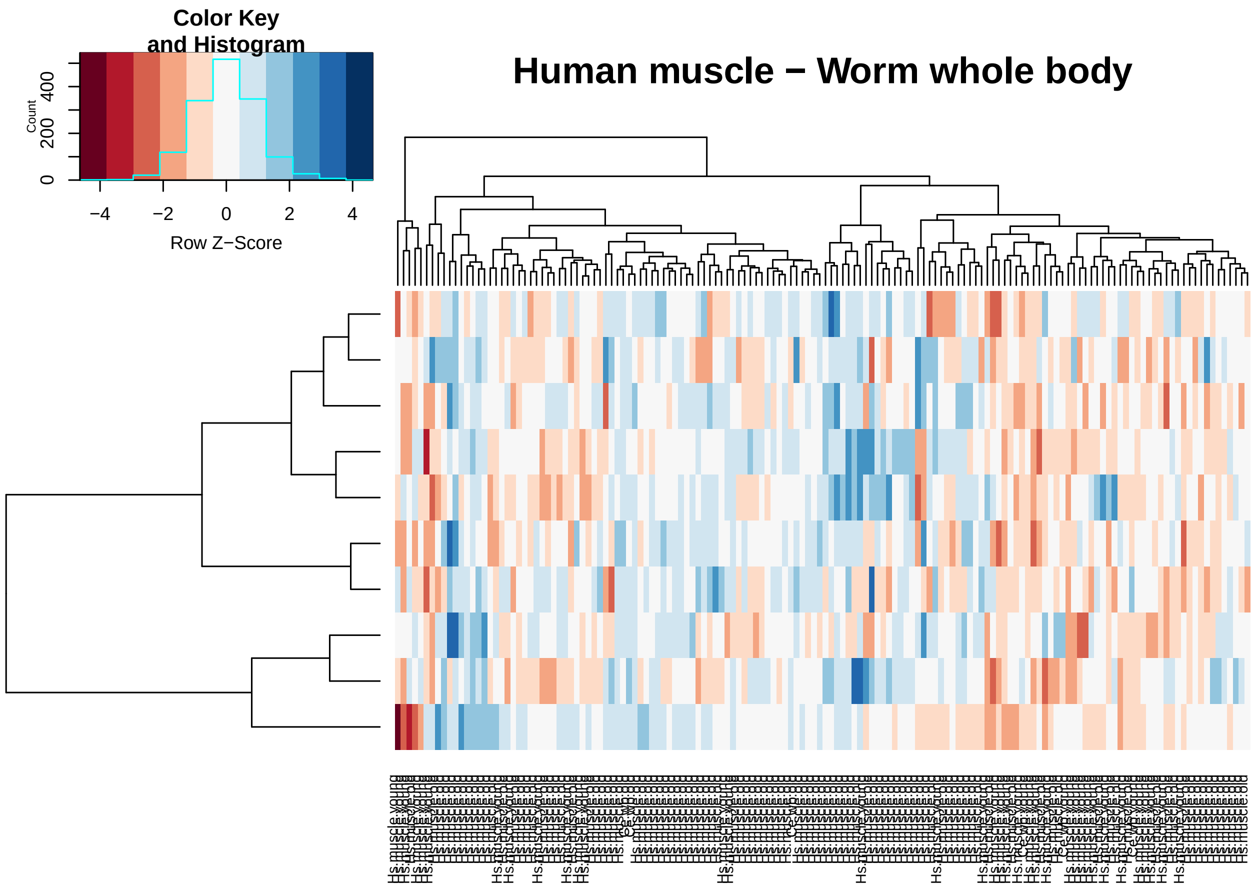


```
clusters.human.worm.hippo <- clustering.samples( signif.hippo.human.worm, expression.human.hippo,express
                                                 "celegans_homolog_ensembl_gene",
                                                 "~/Project1/manuscript_GSEA/results/FigureS2_heatmap_hu
                                                 "Human hippocampus - Worm whole body", plot = FALSE)
```

```
## Found2batches
## Adjusting for0covariate(s) or covariate level(s)
## Standardizing Data across genes
## Fitting L/S model and finding priors
## Finding parametric adjustments
## Adjusting the Data
```

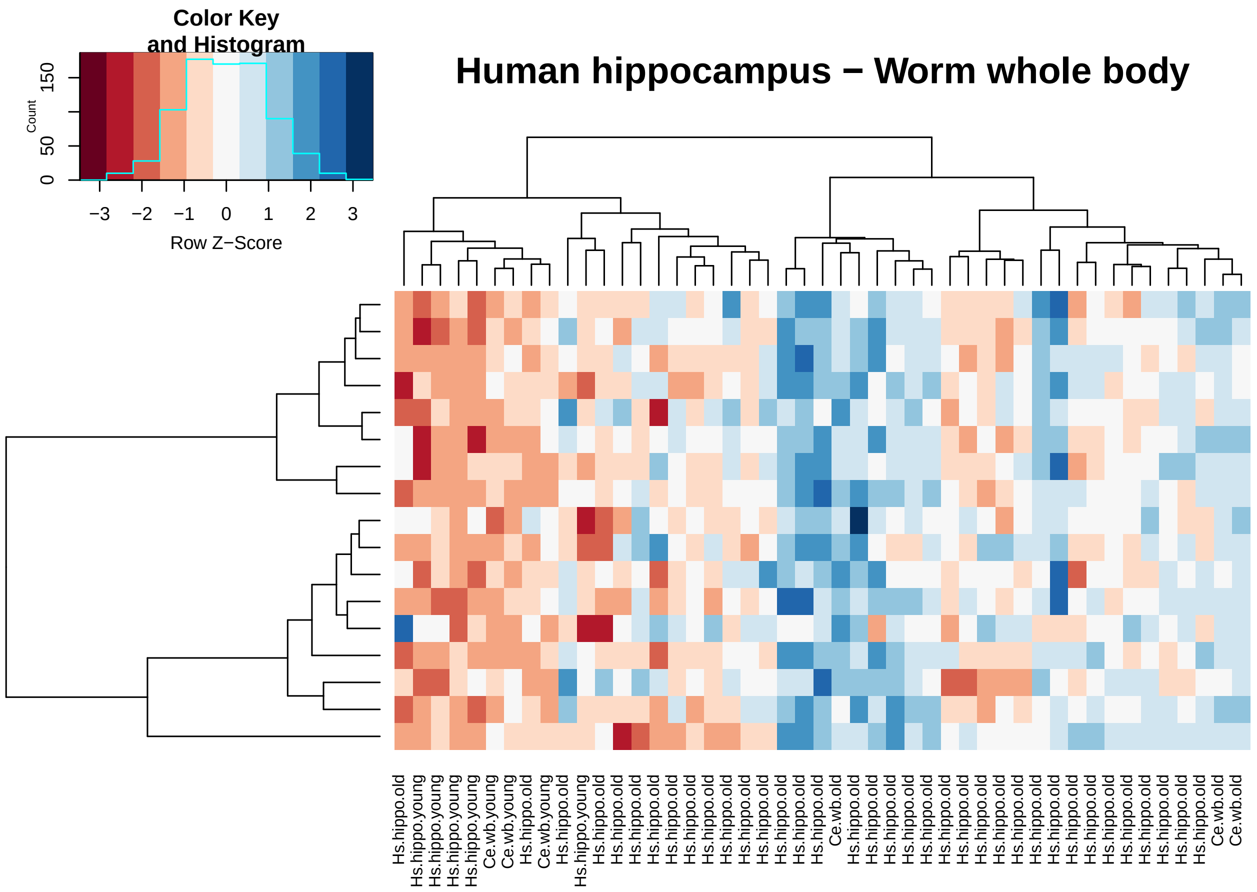


**Table S2. Differential expression summary stats per species**

```
annotation.deg.matrices <- function(genome, differential.exp.matrix){

  differential.exp.matrix$gene.names <- mapIds(genome,
                                               keys=rownames(differential.exp.matrix),
                                               column="SYMBOL",
                                               keytype="ENSEMBL",
                                               multiVals="first")
  # differential.exp.matrix <- #differential.exp.matrix[complete.cases(differential.exp.matrix$gene_nam
  return(differential.exp.matrix)

}



anno.deg.human.muscle <- annotation.deg.matrices(org.Hs.eg.db, deg.muscle.human.aging)
```

```
## 'select()' returned 1:many mapping between keys and columns
```

```
anno.deg.human.hippo <- annotation.deg.matrices(org.Hs.eg.db, deg.hippo.human.aging)
```

```
## 'select()' returned 1:many mapping between keys and columns
```

```
# mouse
anno.deg.mouse.muscle <- annotation.deg.matrices(org.Mm.eg.db, deg.muscle.mouse.aging)
```

```
## 'select()' returned 1:many mapping between keys and columns
```

```
anno.deg.mouse.hippo <- annotation.deg.matrices(org.Mm.eg.db, deg.hippo.mouse.aging)
```

```
## 'select()' returned 1:many mapping between keys and columns
```

```
# fly
anno.deg.fly.wholebody <- annotation.deg.matrices(org.Dm.eg.db, deg.wholebody.fly.aging)
```

```
## 'select()' returned 1:many mapping between keys and columns
```

```
# worm
anno.deg.worm.wholebody <- annotation.deg.matrices(org.Ce.eg.db, deg.wholebody.worm.aging)
```

```
## 'select()' returned 1:many mapping between keys and columns
```

```
differential.expression.gene.list <-
  list(DEG_muscle_human_aging = anno.deg.human.muscle,
       DEG_hippo_human_aging = anno.deg.human.hippo,
       DEG_muscle_mouse_aging = anno.deg.mouse.muscle,
       DEG_hippo_mouse_aging = anno.deg.mouse.hippo,
       DEG_wholebody_fly_aging = anno.deg.fly.wholebody,
       DEG_wholebody_worm_aging = anno.deg.worm.wholebody)

differential.expression.gene.list<-
  lapply(differential.expression.gene.list,
         function(x) {  x$ENSEMBL <- rownames(x) ; x })

#### ----------------------------------------------
# Supplement Table S2

# WriteXLS(differential.expression.gene.list,
#          ExcelFileName = "~/Project1/manuscript_GSEA/supplementary_data/Table_S2.xlsx", SheetNames = N
```

**Table S3. Percentage of overlap pairwise species human vs others.**

```
### muscle
human.mouse.muscle <- select.signif.genes.sp(deg.muscle.human.aging, deg.muscle.mouse.aging,
                                common.human.orthologs.one2one$ToMouse, "mmusculus_homolog_ensembl_gene
                cutoff = 0.1, "Skeletal Muscle (Human)/Skeletal Muscle (Mouse)") #778
```

```
## Number of significant DEGs in human dataset: 5053
## Number of significant DEGs in mouse dataset: 2455
## Number of common DEGs based on 1-1 orthologs: 778
```

```
human.fly.muscle <- select.signif.genes.sp(deg.muscle.human.aging, deg.wholebody.fly.aging,
                                common.human.orthologs.one2one$ToDmel, "dmelanogaster_homolog_ensembl_g
                cutoff = 0.1, "Skeletal Muscle (Human)/Whole Body (Fly)") #395
```

```
## Number of significant DEGs in human dataset: 5053
## Number of significant DEGs in mouse dataset: 4757
## Number of common DEGs based on 1-1 orthologs: 395
```

```
human.worm.muscle <- select.signif.genes.sp(deg.muscle.human.aging,deg.wholebody.worm.aging,
                                common.human.orthologs.one2one$ToCele, "celegans_homolog_ensembl_gene",
                cutoff = 0.1, "Skeletal Muscle (Human)/Whole Body (Worm)") #103
```

```
## Number of significant DEGs in human dataset: 5053
```

```
## Number of significant DEGs in mouse dataset: 3538
## Number of common DEGs based on 1-1 orthologs: 103
```

```
#### hippocampus
human.mouse.hippo <- select.signif.genes.sp(deg.hippo.human.aging, deg.hippo.mouse.aging,
                                                common.human.orthologs.one2one$ToMouse, "mmusculus_homol
                                cutoff = 0.1, "Hippocampus (Human)/Hippocampus (Mouse)") #662
```

```
## Number of significant DEGs in human dataset: 6083
## Number of significant DEGs in mouse dataset: 1639
## Number of common DEGs based on 1-1 orthologs: 662
```

```
human.fly.hippo <- select.signif.genes.sp(deg.hippo.human.aging, deg.wholebody.fly.aging,
                                                common.human.orthologs.one2one$ToDmel, "dmelanogaster_ho
                                cutoff = 0.1, "Hippocampus (Human)/Whole Body (Fly)") #379
```

```
## Number of significant DEGs in human dataset: 6083
## Number of significant DEGs in mouse dataset: 4757
## Number of common DEGs based on 1-1 orthologs: 379
```

```
human.worm.hippo <- select.signif.genes.sp(deg.hippo.human.aging, deg.wholebody.worm.aging,
                                                common.human.orthologs.one2one$ToCele, "celegans_homolog
                                cutoff = 0.1, "Hippocampus (Human)/Whole Body (Worm)") # 112
```

```
## Number of significant DEGs in human dataset: 6083
## Number of significant DEGs in mouse dataset: 3538
## Number of common DEGs based on 1-1 orthologs: 112
```

```
## supplement table S3
overlap <- do.call("rbind", list(human.mouse.muscle, human.fly.muscle,
                                 human.worm.muscle, human.mouse.hippo,
                                 human.fly.hippo, human.worm.hippo))

# WriteXLS(overlap,
#         ExcelFileName = "~/Project1/manuscript_GSEA/supplementary_data/Table_S3.xlsx",
#         SheetNames = NULL, row.names = FALSE)
```

Fig S3. Correlations on orthologous gene-levels.

Single gene-level analysis.

```
## --------------------
# FUNCTIONS
## --------------------

defining.cutoff <- function(single.gene.df, cutoff = NULL){
      return(single.gene.df$sp1.sp2[which(single.gene.df$sp1.sp2$adj.P.Val.x < cutoff &
                                            single.gene.df$sp1.sp2$adj.P.Val.y < cutoff),])
}


labeling.plot.cutoff <- function(correlation, merged.mat, plot.object){
  # require(cowplot)
  label.sp <- substitute(paste(rho, " = ", estimate, ", p = ", pvalue),
                         list(estimate = signif(correlation$estimate, 2),
                              pvalue = signif(correlation$p.value, 2)))
  label.n <- substitute(paste("n = ", n), list(n = dim(merged.mat)[1]))
```

```r
  pp <- ggdraw(plot.object) + draw_label(label.sp, .25, .9, size = 10)
  ppp <- ggdraw(pp) + draw_label(label.n, .3, .85, size = 10)

  return(ppp)
}

labeling.plot.gs <- function(correlation, plot.object){
  # require(cowplot)
  label.sp <- substitute(paste(rho, " = ", estimate, ", p = ", pvalue),
                         list(estimate = signif(correlation$corr$estimate, 2),
                              pvalue = signif(correlation$corr$p.value, 2)))
  label.n <- substitute(paste("n = ", n), list(n = dim(correlation$sp1.sp2)[1]))

  pp <- ggdraw(plot.object) + draw_label(label.sp, .3, .9, size = 10)
  ppp <- ggdraw(pp)  + draw_label(label.n, .3, .85, size = 10)

  return(ppp)
}



correlation.spearman <- function(matrix.species){
  print(cor.test(matrix.species$logFC.x, matrix.species$logFC.y, method = "spearman"))
  correlation <- cor.test(matrix.species$logFC.x, matrix.species$logFC.y, method = "spearman")
  return(correlation)
}

plotting.singlegene <- function(data, xname, yname, colour.dots, colour.lm, gtitle){
  ggplot(data, aes(x = logFC.x, y = logFC.y)) +
    geom_point(size = 2.5, alpha = 0.5, colour = colour.dots) +
    labs(list(title = gtitle,  x = xname, y = yname)) + xlim(c(-0.06, 0.06) ) + ylim(c(-4,3)) +
    theme(text = element_text(size=12)) +
    geom_hline(yintercept=0, colour = "grey50") +
    geom_vline(xintercept = 0, colour = "grey50")
}

correlations.singlegene.level.aging <- function(species1, species2, orthologs.relationships,
                                               ensembl.id.names.species, xtext, ytext){

  merged.ortho.species1 <- merge(orthologs.relationships, species1, by.x = "ensembl_gene_id", by.y= "ro
  merged.species1.species2 <- merge(merged.ortho.species1, species2, by.x = ensembl.id.names.species, b
  cat("Number of common genes between the species after filtering:", dim(merged.species1.species2)[1],

  cat("Summary of", xtext, ytext, "\n")
  print(cor.test(merged.species1.species2$logFC.x, merged.species1.species2$logFC.y, method = "spearman
  correlation <- cor.test(merged.species1.species2$logFC.x, merged.species1.species2$logFC.y, method =

  return(list(corr = correlation, sp1.sp2 = merged.species1.species2))
}

map.the.entrez <- function(toptable, genome.db){
  toptable$entrez = AnnotationDbi::mapIds(genome.db,
                                         keys=rownames(toptable),
```

```
                                                     column="ENTREZID",
                                                     keytype="ENSEMBL",
                                                     multiVals="first")
  return(toptable)
}
```

```
## Muscle

# Human -> Mouse
corr.HumanMouse.SingleGene.Muscle <- correlations.singlegene.level.aging( deg.muscle.human.aging,
        deg.muscle.mouse.aging, common.human.orthologs.one2one$ToMouse, "mmusculus_homolog_ensembl_gene
```

```
## Number of common genes between the species after filtering: 9287
## Summary of Human Mouse
##
##  Spearman's rank correlation rho
##
## data:  merged.species1.species2$logFC.x and merged.species1.species2$logFC.y
## S = 1.3879e+11, p-value = 0.0001329
## alternative hypothesis: true rho is not equal to 0
## sample estimates:
##         rho
## -0.03963933
```

```
# Human -> Fly
corr.HumanFly.SingleGene.Muscle <- correlations.singlegene.level.aging( deg.muscle.human.aging,
    deg.wholebody.fly.aging, common.human.orthologs.one2one$ToDmel, "dmelanogaster_homolog_ensembl_gene
```

```
## Number of common genes between the species after filtering: 2808
## Summary of Human Mouse
##
##  Spearman's rank correlation rho
##
## data:  merged.species1.species2$logFC.x and merged.species1.species2$logFC.y
## S = 3385500000, p-value = 1.189e-05
## alternative hypothesis: true rho is not equal to 0
## sample estimates:
##        rho
## 0.08254312
```

```
# Human -> Worm
corr.HumanWorm.SingleGene.Muscle <- correlations.singlegene.level.aging( deg.muscle.human.aging,
        deg.wholebody.worm.aging, common.human.orthologs.one2one$ToCele, "celegans_homolog_ensembl_gene
```

```
## Number of common genes between the species after filtering: 2302
## Summary of Human Mouse
##
##  Spearman's rank correlation rho
##
## data:  merged.species1.species2$logFC.x and merged.species1.species2$logFC.y
## S = 1943300000, p-value = 0.03402
## alternative hypothesis: true rho is not equal to 0
## sample estimates:
##        rho
## 0.04418579
```

```
## Hippocampus

# Human -> Mouse
corr.HumanMouse.SingleGene.Hippo <- correlations.singlegene.level.aging( deg.hippo.human.aging,
        deg.hippo.mouse.aging, common.human.orthologs.one2one$ToMouse, "mmusculus_homolog_ensembl_gene"
```

```
## Number of common genes between the species after filtering: 12067
## Summary of Human Mouse
##
##  Spearman's rank correlation rho
##
## data:  merged.species1.species2$logFC.x and merged.species1.species2$logFC.y
## S = 2.3095e+11, p-value < 2.2e-16
## alternative hypothesis: true rho is not equal to 0
## sample estimates:
##       rho
## 0.2113838
```

```
# Human -> Fly
corr.HumanFly.SingleGene.Hippo <- correlations.singlegene.level.aging( deg.hippo.human.aging,
    deg.wholebody.fly.aging, common.human.orthologs.one2one$ToDmel, "dmelanogaster_homolog_ensembl_gene
```

```
## Number of common genes between the species after filtering: 2860
## Summary of Human Fly
##
##  Spearman's rank correlation rho
##
## data:  merged.species1.species2$logFC.x and merged.species1.species2$logFC.y
## S = 3727300000, p-value = 0.01852
## alternative hypothesis: true rho is not equal to 0
## sample estimates:
##        rho
## 0.04403345
```

```
# Human -> Worm
corr.HumanWorm.SingleGene.Hippo <- correlations.singlegene.level.aging( deg.hippo.human.aging,
             deg.wholebody.worm.aging, common.human.orthologs.one2one$ToCele, "celegans_homolog_ensembl_
```

```
## Number of common genes between the species after filtering: 2325
## Summary of Human Worm
##
##  Spearman's rank correlation rho
##
## data:  merged.species1.species2$logFC.x and merged.species1.species2$logFC.y
## S = 2080500000, p-value = 0.7439
## alternative hypothesis: true rho is not equal to 0
## sample estimates:
##         rho
## 0.006780104
```

```
## ------------------------------
# putting it under the cutoff of 0.05 to remove zeros

## cutoff-free

# skeletal muscle
```

```r
human.mouse.singlegenes.muscle <- defining.cutoff(corr.HumanMouse.SingleGene.Muscle, cutoff = 0.99)
human.fly.singlegenes.muscle <- defining.cutoff(corr.HumanFly.SingleGene.Muscle, cutoff = 0.99)
human.worm.singlegenes.muscle <- defining.cutoff(corr.HumanWorm.SingleGene.Muscle, cutoff = 0.99)


# hippocampus
human.mouse.singlegenes.hippo <- defining.cutoff(corr.HumanMouse.SingleGene.Hippo, cutoff = 0.99)
human.fly.singlegenes.hippo <- defining.cutoff(corr.HumanFly.SingleGene.Hippo, cutoff = 0.99)
human.worm.singlegenes.hippo <- defining.cutoff(corr.HumanWorm.SingleGene.Hippo, cutoff = 0.99)
```

```r
## ------------
# check the correlations after cutoff

## muscle
cutoff.corrs.human.mouse.muscle <- correlation.spearman(human.mouse.singlegenes.muscle)
```

```
##
##  Spearman's rank correlation rho
##
## data:  matrix.species$logFC.x and matrix.species$logFC.y
## S = 1.3246e+11, p-value = 0.0001374
## alternative hypothesis: true rho is not equal to 0
## sample estimates:
##         rho
## -0.03986407
```

```r
cutoff.corrs.human.fly.muscle <- correlation.spearman(human.fly.singlegenes.muscle)
```

```
##
##  Spearman's rank correlation rho
##
## data:  matrix.species$logFC.x and matrix.species$logFC.y
## S = 3276300000, p-value = 1.169e-05
## alternative hypothesis: true rho is not equal to 0
## sample estimates:
##        rho
## 0.08305571
```

```r
cutoff.corrs.human.worm.muscle <- correlation.spearman(human.worm.singlegenes.muscle)
```

```
##
##  Spearman's rank correlation rho
##
## data:  matrix.species$logFC.x and matrix.species$logFC.y
## S = 1852200000, p-value = 0.03269
## alternative hypothesis: true rho is not equal to 0
## sample estimates:
##        rho
## 0.04487102
```

```r
#### hippocampus
cutoff.corrs.human.mouse.hippo <- correlation.spearman(human.mouse.singlegenes.hippo)
```

```
##
##  Spearman's rank correlation rho
##
```

```
## data:  matrix.species$logFC.x and matrix.species$logFC.y
## S = 2.1832e+11, p-value < 2.2e-16
## alternative hypothesis: true rho is not equal to 0
## sample estimates:
##       rho
## 0.2135748
```

```
cutoff.corrs.human.fly.hippo <- correlation.spearman(human.fly.singlegenes.hippo)
```

```
##
##  Spearman's rank correlation rho
##
## data:  matrix.species$logFC.x and matrix.species$logFC.y
## S = 3588100000, p-value = 0.01918
## alternative hypothesis: true rho is not equal to 0
## sample estimates:
##        rho
## 0.04407119
```

```
cutoff.corrs.human.worm.hippo <- correlation.spearman(human.worm.singlegenes.hippo)
```

```
##
##  Spearman's rank correlation rho
##
## data:  matrix.species$logFC.x and matrix.species$logFC.y
## S = 1961400000, p-value = 0.7346
## alternative hypothesis: true rho is not equal to 0
## sample estimates:
##        rho
## 0.00710321
```

Saving the plots for the Supplementary Figure

```
# Human - Mouse
p1.muscle.cutoff <- plotting.singlegene(human.mouse.singlegenes.muscle,
             "log2 Fold-change Human", "log2 Fold-change Mouse", "indianred1", "red", "Skeletal Muscle"
pp1.muscle.cutoff <- labeling.plot.cutoff(cutoff.corrs.human.mouse.muscle, human.mouse.singlegenes.muscl
```

```
## Warning: Removed 3 rows containing missing values (geom_point).
```

```
p1.hippo.cutoff <- plotting.singlegene(human.mouse.singlegenes.hippo, "log2 Fold-change Human", "log2 F
pp1.hippo.cutoff <- labeling.plot.cutoff(cutoff.corrs.human.mouse.hippo, human.mouse.singlegenes.hippo,
```

```
## Warning: Removed 13 rows containing missing values (geom_point).
```

```
# Human - Fly - here it is ok
p2.muscle.cutoff <- plotting.singlegene(human.fly.singlegenes.muscle, "log2 Fold-change Human", "log2 F
pp2.muscle.cutoff <- labeling.plot.cutoff(cutoff.corrs.human.fly.muscle, human.fly.singlegenes.muscle, 
```

```
## Warning: Removed 1 rows containing missing values (geom_point).
```

```
p2.hippo.cutoff <- plotting.singlegene(human.fly.singlegenes.hippo,  "log2 Fold-change Human", "log2 Fo
pp2.hippo.cutoff <- labeling.plot.cutoff(cutoff.corrs.human.fly.hippo, human.fly.singlegenes.hippo, p2.


# Human - Worm
p3.muscle.cutoff <- plotting.singlegene(human.worm.singlegenes.muscle,  "log2 Fold-change Human", "log2
pp3.muscle.cutoff <- labeling.plot.cutoff(cutoff.corrs.human.worm.muscle, human.worm.singlegenes.muscle
```

```
p3.hippo.cutoff <- plotting.singlegene(human.worm.singlegenes.hippo,  "log2 Fold-change Human", "log2 Fo
pp3.hippo.cutoff <- labeling.plot.cutoff(cutoff.corrs.human.worm.hippo, human.worm.singlegenes.hippo, p3
```

```
# pdf("~/Project1/manuscript_GSEA/results/FigureS3_Single_Gene_Only.pdf", 10, 7)
plot_grid(pp1.muscle.cutoff, pp1.hippo.cutoff,
                    pp2.muscle.cutoff, pp2.hippo.cutoff,
          pp3.muscle.cutoff, pp3.hippo.cutoff,
                    labels = c("A", "", "B", "", "C", ""), nrow = 3, align = "h")
```

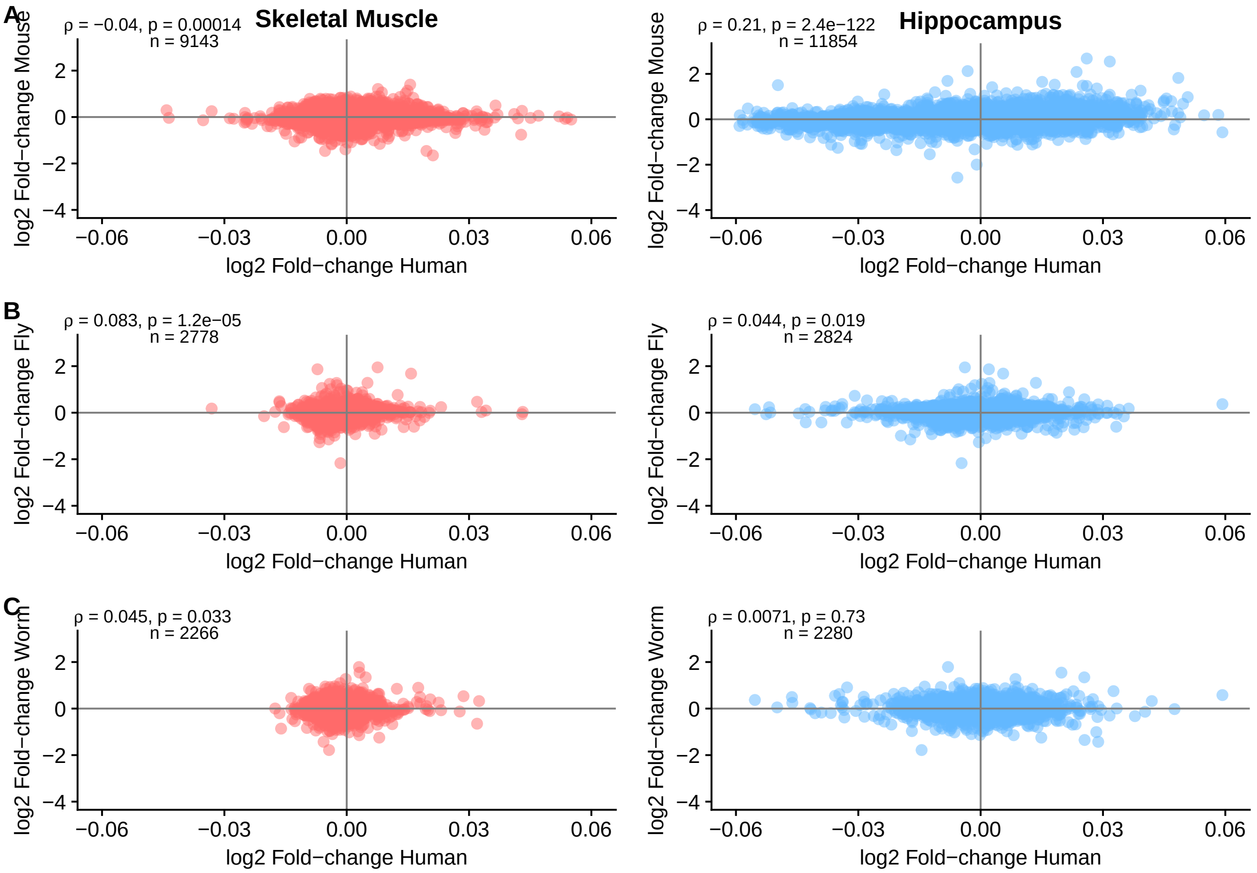


```
# dev.off()
```

##### Single-species GSEA

Generating Figure S4.

```
# functions for the analysis
go.gsea.analysis <- function(species, toptable){


  if(species == "Human"){
    library(org.Hs.eg.db)
    go.human <- go.gsets(species)
    go.human.sets <- go.human$go.sets
    go.human.subs <- go.human$go.subs

    cat("Getting GO categories.\n")
```

```
  gobpsets.human <- go.human.sets[go.human.subs$BP]
  goccsets.human <- go.human.sets[go.human.subs$CC]
  gomfsets.human <- go.human.sets[go.human.subs$MF]

  toptable <- map.the.entrez(toptable, org.Hs.eg.db)
  tt.pc.order <- toptable[order(toptable$adj.P.Val),]
  gene.stats <- foldchange.tvalues(tt.pc.order)

  cat("Showing logFoldChanges and t-values.\n")
  print(lapply(gene.stats, head))

  fc <- fun.analysis.go(foldchangesstats = gene.stats[[1]],
                      tstats = NULL, gobpsets.human, gomfsets.human, goccsets.human)
  t <- fun.analysis.go(tstats = gene.stats[[2]],
                     foldchangesstats = NULL, gobpsets.human, gomfsets.human, goccsets.human)

} else if(species == "Mouse") {

  library(org.Mm.eg.db)
  go.mouse <- go.gsets(species)
  go.mouse.sets <- go.mouse$go.sets
  go.mouse.subs <- go.mouse$go.subs

  cat("Getting GO categories.\n")
  gobpsets.mouse <- go.mouse.sets[go.mouse.subs$BP]
  goccsets.mouse <- go.mouse.sets[go.mouse.subs$CC]
  gomfsets.mouse <- go.mouse.sets[go.mouse.subs$MF]

  toptable <- map.the.entrez(toptable, org.Mm.eg.db)
  tt.pc.order <- toptable[order(toptable$adj.P.Val),]
  gene.stats <- foldchange.tvalues(tt.pc.order)

  cat("Showing logFoldChanges and t-values.\n")
  print(lapply(gene.stats, head))

  fc <- fun.analysis.go(foldchangesstats = gene.stats[[1]],
                      tstats = NULL, gobpsets.mouse, gomfsets.mouse, goccsets.mouse)
  t <- fun.analysis.go(tstats = gene.stats[[2]],
                     foldchangesstats = NULL, gobpsets.mouse, gomfsets.mouse, goccsets.mouse)


} else if(species == "Worm") {
  library(org.Ce.eg.db)
  go.worm <- go.gsets(species)
  go.worm.sets <- go.worm$go.sets
  go.worm.subs <- go.worm$go.subs

  cat("Getting GO categories.\n")
  gobpsets.worm <- go.worm.sets[go.worm.subs$BP]
  goccsets.worm <- go.worm.sets[go.worm.subs$CC]
  gomfsets.worm <- go.worm.sets[go.worm.subs$MF]

  toptable <- map.the.entrez(toptable, org.Ce.eg.db)
```

```r
    tt.pc.order <- toptable[order(toptable$adj.P.Val),]
    gene.stats <- foldchange.tvalues(tt.pc.order)

    cat("Showing logFoldChanges and t-values.\n")
    print(lapply(gene.stats, head))

    fc <- fun.analysis.go(foldchangesstats = gene.stats[[1]],
                          tstats = NULL, gobpsets.worm, gomfsets.worm, goccsets.worm)
    t <- fun.analysis.go(tstats = gene.stats[[2]],
                         foldchangesstats = NULL, gobpsets.worm, gomfsets.worm, goccsets.worm)


  } else if(species == "Fly") {
    library(org.Dm.eg.db)
    go.fly <- go.gsets(species)
    go.fly.sets <- go.fly$go.sets
    go.fly.subs <- go.fly$go.subs

    cat("Getting GO categories.\n")
    gobpsets.fly <- go.fly.sets[go.fly.subs$BP]
    goccsets.fly <- go.fly.sets[go.fly.subs$CC]
    gomfsets.fly <- go.fly.sets[go.fly.subs$MF]

    toptable <- map.the.entrez(toptable, org.Dm.eg.db)
    tt.pc.order <- toptable[order(toptable$adj.P.Val),]
    gene.stats <- foldchange.tvalues(tt.pc.order)

    cat("Showing logFoldChanges and t-values.\n")
    print(lapply(gene.stats, head))

    fc <- fun.analysis.go(foldchangesstats = gene.stats[[1]],
                          tstats = NULL, gobpsets.fly, gomfsets.fly, goccsets.fly)
    t <- fun.analysis.go(tstats = gene.stats[[2]],
                         foldchangesstats = NULL, gobpsets.fly, gomfsets.fly, goccsets.fly)
  }

  return(list(foldchange.analysis = fc, tstat.analysis = t))
}


fun.analysis.go <- function(tstats = NULL, foldchangesstats = NULL,
                            gobpsets.species, gomfsets.species, goccsets.species){
  cat("Calculating the functional analysis.\n")

  if(length(tstats) > 1){
    cat("Calculating the functional analysis on t-values.\n")
    gobpres.healthy <- gage(tstats,  gsets=gobpsets.species, same.dir=TRUE, use.fold = FALSE, ref = NULL
    gomfres.healthy<- gage(tstats,   gsets=gomfsets.species, same.dir=TRUE, use.fold = FALSE, ref = NULL
    goccres.healthy <- gage(tstats,  gsets=goccsets.species, same.dir=TRUE, use.fold = FALSE, ref = NULL

  } else {

    cat("Calculating the functional analysis on logFC values.\n")
```

```
    gobpres.healthy <- gage(foldchangesstats,  gsets=gobpsets.species, same.dir=TRUE, ref = NULL, samp =
    gomfres.healthy <- gage(foldchangesstats,  gsets=gomfsets.species, same.dir=TRUE, ref = NULL, samp =
    goccres.healthy <- gage(foldchangesstats, gsets=goccsets.species, same.dir=TRUE, ref = NULL, samp =

  }

  return(list(gobpres.healthy, gomfres.healthy, goccres.healthy))

}

significant.go <- function(results.go.species) {
  results.go <- results.go.species[[2]]
  greater.go <- list()
  less.go <- list()
  for(i in 1:3){
    # this cutoff is because of the mouse data
    greater.go[[i]] <- results.go[[i]]$greater[results.go[[i]]$greater[,4] < 0.20, ]
    less.go[[i]] <- results.go[[i]]$less[results.go[[i]]$less[,4] < 0.20, ]
  }

  return(list(up.go = greater.go, down.go = less.go))

}


choose.go <- function(significant.gos, ont = "BP", number.of.sets = 20){

  if(ont == "BP"){
    ups <- significant.gos$up.go[[1]][1:number.of.sets, ]
    downs <- significant.gos$down.go[[1]][1:number.of.sets, ]
  } else if (ont == "MF"){
    ups <- significant.gos$up.go[[2]][1:number.of.sets, ]
    downs <- significant.gos$down.go[[2]][1:number.of.sets, ]
  } else {
    ups <- significant.gos$up.go[[3]][1:number.of.sets, ]
    downs <- significant.gos$down.go[[3]][1:number.of.sets, ]
  }

  df <- as.data.frame(rbind(ups, downs))
  df$Name <- rownames(df)
  return(df)

}


foldchange.tvalues <- function(toptable){
  foldchanges <- toptable$logFC
  names(foldchanges) <- toptable$entrez
  tvalues <- toptable$t
  names(tvalues) <- toptable$entrez
  return(list(foldchanges, tvalues))
}
```

```r
#### Species-specific analysis

# ----------------------------------------------------
# GSEA on GO ontology
# ----------------------------------------------------
library(gage)

# GSEA now
results.go.human.muscle <- go.gsea.analysis("Human", deg.muscle.human.aging)
```

```
## Gene ID type for 'human' is: 'EG'

##

## Getting GO categories.

## 'select()' returned 1:many mapping between keys and columns

## Showing logFoldChanges and t-values.
## [[1]]
##      221061      10655        8701       54102       55885      442117
## -0.02602730 -0.03311675  0.06730171 -0.04387005  0.04556766  0.06637557
##
## [[2]]
##    221061     10655      8701     54102     55885    442117
## -9.854277 -9.365321  8.469465 -8.434936  7.888115  7.697907
##
## Calculating the functional analysis.
## Calculating the functional analysis on logFC values.
## Calculating the functional analysis.
## Calculating the functional analysis on t-values.
```

```r
results.go.mouse.muscle <- go.gsea.analysis("Mouse", deg.muscle.mouse.aging)
```

```
## Gene ID type for 'mouse' is: 'EG'

## Getting GO categories.

## 'select()' returned 1:many mapping between keys and columns

## Showing logFoldChanges and t-values.
## [[1]]
##     118449      21912     110880      56495      66469      70510
## -1.1810740 -1.0756396 -1.4619566 -1.1449355 -0.7841583 -0.7509859
##
## [[2]]
##    118449     21912    110880     56495     66469     70510
## -16.08017 -15.36940 -14.24539 -14.00831 -13.64479 -13.40626
##
## Calculating the functional analysis.
## Calculating the functional analysis on logFC values.
## Calculating the functional analysis.
## Calculating the functional analysis on t-values.
```

```r
results.go.human.hippo <- go.gsea.analysis("Human", deg.hippo.human.aging)
```

```
## Gene ID type for 'human' is: 'EG'

## Getting GO categories.
```

```
## 'select()' returned 1:many mapping between keys and columns

## Showing logFoldChanges and t-values.
## [[1]]
##       27287      266722      115992       22843      285220         347
##  0.08460573 -0.06411874  0.01738765 -0.06430807 -0.05484216  0.03129872
##
## [[2]]
##     27287    266722    115992     22843    285220       347
##  7.086007 -6.782670  6.080744 -5.978037 -5.915660  5.910514
##
## Calculating the functional analysis.
## Calculating the functional analysis on logFC values.
## Calculating the functional analysis.
## Calculating the functional analysis on t-values.
```

```
results.go.mouse.hippo <- go.gsea.analysis("Mouse", deg.hippo.mouse.aging)
```

```
## Gene ID type for 'mouse' is: 'EG'

## Getting GO categories.

## 'select()' returned 1:many mapping between keys and columns

## Showing logFoldChanges and t-values.
## [[1]]
##     12268      <NA>     78781     55990     93961     93880
## 2.2492003 1.6899556 1.0632543 2.6770648 0.5585991 1.0268640
##
## [[2]]
##    12268     <NA>    78781    55990    93961    93880
## 16.81314 15.17006 13.27444 12.97582 12.51056 12.39425
##
## Calculating the functional analysis.
## Calculating the functional analysis on logFC values.
## Calculating the functional analysis.
## Calculating the functional analysis on t-values.
```

```
results.go.dmelano.wholebody <- go.gsea.analysis("Fly", deg.wholebody.fly.aging)
```

```
## Gene ID type for 'fly' is: 'EG'

## Getting GO categories.

## 'select()' returned 1:many mapping between keys and columns

## Showing logFoldChanges and t-values.
## [[1]]
##     59235     35409     36410     44921     39225     34800
## -2.731647 -2.248416 -4.169935  3.940811  2.146957 -1.578291
##
## [[2]]
##     59235     35409     36410     44921     39225     34800
## -41.34455 -35.32211 -30.91917  28.68678  26.37980 -26.30645
##
## Calculating the functional analysis.
## Calculating the functional analysis on logFC values.
## Calculating the functional analysis.
## Calculating the functional analysis on t-values.
```

```r
results.go.celegans.wholebody <- go.gsea.analysis("Worm",deg.wholebody.worm.aging)
```

```
## Gene ID type for 'worm' is: 'EG'

## Getting GO categories.

## 'select()' returned 1:many mapping between keys and columns

## Showing logFoldChanges and t-values.
## [[1]]
##    174652    178245    177447    178971    179804    184206
## -3.560177 -3.297889 -3.198386 -3.109974 -3.434989 -3.364338
##
## [[2]]
##    174652    178245    177447    178971    179804    184206
## -36.67322 -34.32467 -34.26308 -30.11533 -29.80401 -29.61442
##
## Calculating the functional analysis.
## Calculating the functional analysis on logFC values.
## Calculating the functional analysis.
## Calculating the functional analysis on t-values.
```

```r
# get significant ones - FDR: 0.10 - not possible
# up.go; down.go
signif.human.muscle <- significant.go(results.go.human.muscle)
signif.human.hippo <- significant.go(results.go.human.hippo)

signif.mouse.muscle <- significant.go(results.go.mouse.muscle)
signif.mouse.hippo <- significant.go(results.go.mouse.hippo)

signif.dmelano <- significant.go(results.go.dmelano.wholebody)
signif.celegans <- significant.go(results.go.celegans.wholebody)


### filter the gos - now it is without go semantics - top 20 categories
human.muscle.bp <- choose.go(signif.human.muscle)
human.hippo.bp <- choose.go(signif.human.hippo)

mouse.muscle.bp <- choose.go(signif.mouse.muscle)
#head(mouse.muscle.bp)
mouse.hippo.bp <- choose.go(signif.mouse.hippo)

dmelano.bp <- choose.go(signif.dmelano)
cele.bp <- choose.go(signif.celegans)
```

```r
# ------------------------------
# human
human.muscle.bp$GO.ID <- do.call(c, lapply(strsplit(human.muscle.bp$Name, " "), '[', 1))
human.muscle.bp$Annot <- c(rep("Extracellular matrix organization",2), rep("Cell adhesion", 2),
                           "Immune response", "Cell adhesion", rep("Angiogenesis",2), rep("Cell adhesion
                           rep("Immune response", 2),
                           rep("Cell adhesion",6), rep("Mitochondrial translation", 5), "Translation",
                           "Cellular respiration", rep("Catabolic process", 3),
                           rep("Metabolic process", 2), "Cellular protein modification process",
                           rep("Metabolic process", 4), "Cellular protein modification process",
                           "Metabolic process", "Cellular protein modification process")
```

```r
human.muscle.bp.new <- human.muscle.bp[order(human.muscle.bp$Annot), ]
human.muscle.bp.new$Annot <- factor(human.muscle.bp.new$Annot, levels=unique(human.muscle.bp.new$Annot))
human.muscle.bp.new$Name <- factor(human.muscle.bp.new$Name, levels=unique(human.muscle.bp.new$Name))


## human hippocampus
human.hippo.bp$GO.ID <- do.call(c, lapply(strsplit(human.hippo.bp$Name, " "), '[', 1))
human.hippo.bp$Annot <- c(rep("Immune response", 2),  "Viral process", rep("Protein transport", 2), "Vi
                          rep("Immune response", 2), "Protein targeting",
                          "Response to external stimulus", "Catabolic process",
                          rep("Immune response", 6), "Response to external stimulus",
                          "Extracellular matrix organization", "Immune response",
                          rep("Nervous system process", 3) , "Signaling",
                          rep("Nervous system process", 10), "Cellular localization",
                          "Nervous system process", "Cellular localization",
                          "Signaling", rep("Nervous system process",2))
human.hippo.bp.new <- human.hippo.bp[order(human.hippo.bp$Annot), ]
human.hippo.bp.new$Annot <- factor(human.hippo.bp.new$Annot, levels=unique(human.hippo.bp.new$Annot))
human.hippo.bp.new$Name <- factor(human.hippo.bp.new$Name, levels=unique(human.hippo.bp.new$Name))


# -------------------------------
# mouse muscle
mouse.muscle.bp$GO.ID <- do.call(c, lapply(strsplit(mouse.muscle.bp$Name, " "), '[', 1))
# check can you remove NA's
mouse.muscle.bp$Annot <- c(rep(NA, 10), rep(NA, 10), rep("Metabolic process", 10), rep(NA, 10))
mouse.muscle.bp.new <- mouse.muscle.bp[order(mouse.muscle.bp$Annot), ]
mouse.muscle.bp.new <- mouse.muscle.bp.new[complete.cases(mouse.muscle.bp.new$Annot), ]
mouse.muscle.colors <- c("Metabolic process" = "firebrick1")
mouse.muscle.bp.new$Annot <- factor(mouse.muscle.bp.new$Annot, levels=unique(mouse.muscle.bp.new$Annot))
mouse.muscle.bp.new$Name <- factor(mouse.muscle.bp.new$Name, levels=unique(mouse.muscle.bp.new$Name))


### hippocampus
mouse.hippo.bp$GO.ID <- do.call(c, lapply(strsplit(mouse.hippo.bp$Name, " "), '[', 1))
mouse.hippo.bp <- mouse.hippo.bp[complete.cases(mouse.hippo.bp),]
mouse.hippo.bp$Annot <- c(rep("Immune response", 20),
                          rep("Nervous system process", 7))
mouse.hippo.bp.new <- mouse.hippo.bp[order(mouse.hippo.bp$Annot), ]
mouse.hippo.bp.new$Annot <- factor(mouse.hippo.bp.new$Annot, levels=unique(mouse.hippo.bp.new$Annot))
mouse.hippo.bp.new$Name <- factor(mouse.hippo.bp.new$Name, levels=unique(mouse.hippo.bp.new$Name))


# -------------------------------
# dmelanogaster
dmelano.bp$GO.ID <- do.call(c, lapply(strsplit(dmelano.bp$Name, " "), '[', 1))
dmelano.bp$Annot <- c(rep("Immune response", 10), rep("Anatomical structure morphogenesis", 2),
                      rep("Signaling", 2), "Immune response",
                        rep("Anatomical structure morphogenesis", 5),
                      rep("Metabolic process", 7), rep("Cellular respiration", 6),
                      rep("Metabolic process", 7))
dmele.bp.new <- dmelano.bp[order(dmelano.bp$Annot), ]
```

```
dmele.bp.new$Annot <- factor(dmele.bp.new$Annot, levels=unique(dmele.bp.new$Annot))
dmele.bp.new$Name <- factor(dmele.bp.new$Name, levels=unique(dmele.bp.new$Name))


# ------------------------------
# celegans
# remove NAs
cele.bp <- cele.bp[complete.cases(cele.bp),]
cele.bp$GO.ID <- do.call(c, lapply(strsplit(cele.bp$Name, " "), '[', 1))
cele.bp$Annot <- c( rep("Cellular response to stimulus", 5), "Signaling",
                    rep("Cellular response to stimulus", 3), "Signaling",
                      rep("Cellular response to stimulus", 2),
                    rep("Signaling", 2), rep("Nervous system process", 2),
                    rep("Cell adhesion", 2),
                      "Nervous system process", "Cell adhesion",
                    rep("Metabolic process", 3), rep("Immune response", 4),
                    rep("Metabolic process", 7) )

cele.bp.new <- cele.bp[order(cele.bp$Annot), ]

cele.bp.new$Annot <- factor(cele.bp.new$Annot, levels=unique(cele.bp.new$Annot))
cele.bp.new$Name <- factor(cele.bp.new$Name, levels=unique(cele.bp.new$Name))

### check the colors
united.colors.of.genesets <- c("Cellular localization" = "springgreen3",
                "Cellular response to stimulus" = "mediumorchid1",
                "Signaling" = "tan1",
                "Translation" = "palegreen3",
                "Anatomical structure morphogenesis" = "yellow",
                #"Nucleoside metabolic process" = "tomato1",
                "Cellular protein modification process" = "green",
                "Cellular respiration" = "indianred1",
                "Immune response" = "deepskyblue3",
                "Nervous system process" = "goldenrod1",
                "Metabolic process" ="firebrick1",
                "Defense response" =  "dodgerblue",
                "Catabolic process" =  "darkred",
                "Cellular respiration" = "indianred1",
                "Immune response" = "deepskyblue3",
                "Mitochondrial translation" = "brown1",
                "Cofactor biosynthetic process" = "coral1",
                "Extracellular matrix organization" = "rosybrown",
                "Angiogenesis" = "moccasin",
                "Cell adhesion" = "mistyrose3",
                "Protein transport" = "sandybrown",
                "Protein targeting" = "khaki1",
                "Cellular localization" = "olivedrab",
                "Response to external stimulus" = "royalblue",
                "Viral process" = "grey30")
```

Plotting the GSEA results.

```
p1.human.muscle <- ggplot(human.muscle.bp.new, aes(stat.mean, Name)) +
                    geom_point(shape = 19, size =3, colour = "black") + geom_point(shape = 19, size = 2
```

```
                    aes(colour = Annot)) +
                    scale_colour_manual(values = united.colors.of.genesets)  +
                    ylab("GO BP categories") + xlab("GAGE stat. mean") +
                    theme(axis.text.y=element_blank(), legend.position = "right",
                          panel.grid.minor=element_line(color="ivory3",size=0.5),
                          panel.grid.major=element_line(color="ivory3",size=0.5),
                          legend.title=element_blank()) +
                    geom_vline(xintercept = 0) +
                    xlim(c(-10, 10))  +
                    labs(title = "Human Skeletal Muscle")


p1.human.hippo <- ggplot(human.hippo.bp.new, aes(stat.mean, Name)) +
                  geom_point(shape = 19, size =3, colour = "black") + geom_point(shape = 19, size = 2.5
                  aes(colour = Annot)) +
                  scale_colour_manual(values = united.colors.of.genesets)  +
                  ylab("GO BP categories") + xlab("GAGE stat. mean") +
                  theme(axis.text.y=element_blank(), legend.position = "right",
                        panel.grid.minor=element_line(color="ivory3",size=0.5),
                        panel.grid.major=element_line(color="ivory3",size=0.5),
                        legend.title=element_blank()) +
                  geom_vline(xintercept = 0) +
                  xlim(c(-10, 10))  +
                  labs(title = "Human Hippocampus")


p1.mouse.muscle <- ggplot(mouse.muscle.bp.new, aes(stat.mean, Name)) +
                   geom_point(shape = 19, size =3, colour = "black") + geom_point(shape = 19, size = 2.
                   aes(colour = Annot)) +
                   scale_colour_manual(values = united.colors.of.genesets)  +
                   ylab("GO BP categories") + xlab("GAGE stat. mean") +
                   theme(axis.text.y=element_blank(), legend.position = "right",
                         panel.grid.minor=element_line(color="ivory3",size=0.5),
                         panel.grid.major=element_line(color="ivory3",size=0.5),
                         legend.title=element_blank()) +
                   geom_vline(xintercept = 0) +
                   xlim(c(-10, 10))  +
                   labs(title = "Mouse Skeletal Muscle")


p1.mouse.hippo <- ggplot(mouse.hippo.bp.new, aes(stat.mean, Name)) +
                  geom_point(shape = 19, size =3, colour = "black") + geom_point(shape = 19, size = 2.5
                  aes(colour = Annot)) +
                  scale_colour_manual(values = united.colors.of.genesets)  +
                  ylab("GO BP categories") + xlab("GAGE stat. mean") +
                  theme(axis.text.y=element_blank(), legend.position = "right",
                        panel.grid.minor=element_line(color="ivory3",size=0.5),
                        panel.grid.major=element_line(color="ivory3",size=0.5),
                        legend.title=element_blank()) +
                  geom_vline(xintercept = 0) +
                  xlim(c(-10, 10))  +
                  labs(title = "Mouse Hippocampus")
```

```r
p1.dmele <- ggplot(dmele.bp.new, aes(stat.mean, Name)) +
            geom_point(shape = 19, size =3, colour = "black") + geom_point(shape = 19, size = 2.5,
            aes(colour = Annot)) +
            scale_colour_manual(values = united.colors.of.genesets)  +
            ylab("GO BP categories") + xlab("GAGE stat. mean") +
            theme(axis.text.y=element_blank(), legend.position = "right",
                  panel.grid.minor=element_line(color="ivory3",size=0.5),
                  panel.grid.major=element_line(color="ivory3",size=0.5),
                  legend.title=element_blank()) +
            geom_vline(xintercept = 0) +
            xlim(c(-10, 10))  +
            labs(title = "D. melanogaster wholebody")


p1.cele <- ggplot(cele.bp.new, aes(stat.mean, Name)) +
           geom_point(shape = 19, size =3, colour = "black") + geom_point(shape = 19, size = 2.5,
           aes(colour = Annot)) +
           scale_colour_manual(values = united.colors.of.genesets)  +
           ylab("GO BP categories") + xlab("GAGE stat. mean") +
           theme(axis.text.y=element_blank(), legend.position = "right",
                 panel.grid.minor=element_line(color="ivory3",size=0.5),
                 panel.grid.major=element_line(color="ivory3",size=0.5),
                 legend.title=element_blank()) +
           geom_vline(xintercept = 0) +
           xlim(c(-10, 10))  +
           labs(title = "C. elegans wholebody")


### figure S4. for GSEA

# pdf("~/Project1/manuscript_GSEA/results/Figure_S4.pdf", 17, 13, useDingbats = FALSE)
plot_grid(p1.human.muscle, p1.human.hippo,
          p1.mouse.muscle, p1.mouse.hippo,
          p1.dmele, p1.cele,
          labels = c("A", "B", "C", "D", "E", "F"), nrow = 3, align = "h")
```

```
## Warning: Removed 2 rows containing missing values (geom_point).

## Warning: Removed 2 rows containing missing values (geom_point).
```

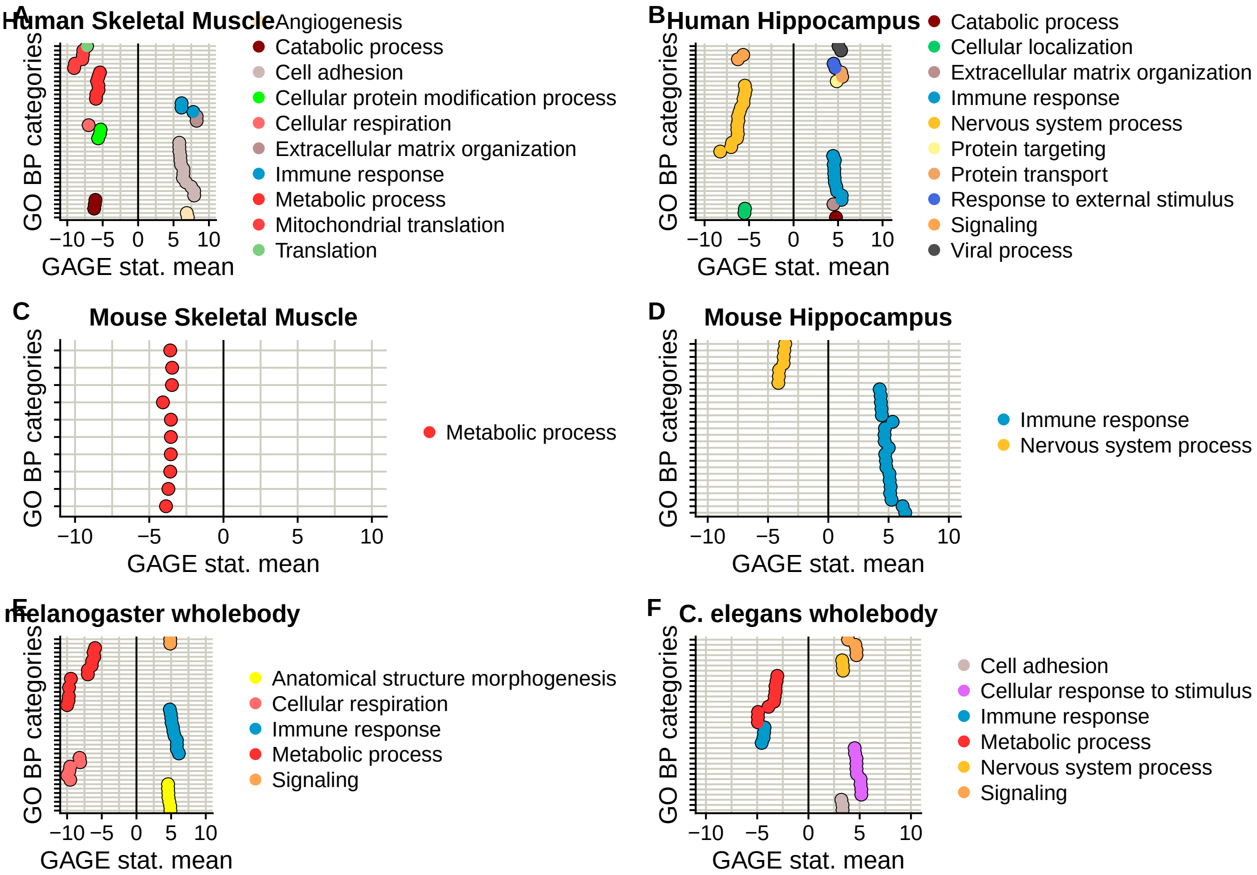
A
Human Skeletal Muscle
GO BP categories
GAGE stat. mean
−10 −5 0 5 10
Angiogenesis
Catabolic process
Cell adhesion
Cellular protein modification process
Cellular respiration
Extracellular matrix organization
Immune response
Metabolic process
Mitochondrial translation
Translation
B
Human Hippocampus
Catabolic process
Cellular localization
Extracellular matrix organization
Immune response
Nervous system process
Protein targeting
Protein transport
Response to external stimulus
Signaling
Viral process
C
Mouse Skeletal Muscle
Metabolic process
D
Mouse Hippocampus
Immune response
Nervous system process
E
melanogaster wholebody
Anatomical structure morphogenesis
Cellular respiration
Immune response
Metabolic process
Signaling
F
C. elegans wholebody
Cell adhesion
Cellular response to stimulus
Immune response
Metabolic process
Nervous system process
Signaling

```
# dev.off()
```

### Main Figures + Supplement (Process part)

*Andrea Komljenović*

*3/9/2018*

Part II

```
## ---------------------------------
# Load the packages

packages <- c( "affy", "affyPLM", "biomaRt", "limma",
               "tidyr", "gplots", "ggplot2", "reshape2",
               "Biobase", "downloader", "sva", "dplyr", "RColorBrewer", "cowplot",
               "igraph", "WriteXLS", "org.Hs.eg.db", "org.Mm.eg.db",
               "org.Dm.eg.db", "org.Ce.eg.db", "gage",
               "cowplot", "ggrepel", "plyr", "data.table",
               "dplyr", "ComplexHeatmap", "circlize", "RColorBrewer")
lapply(packages, library, character.only = TRUE)
```

```
# -----------------------------------------------------------------
## Figure 2
#### integration for skeletal muscle, hippocampus and dietary restriction experiments


# ------------
# function to choose the genes in the orthologous groups with the lowest p-value from DEA
choose.the.lowest.pval <- function(list){
    return(lapply(list, function(x) x[which.min(x$P.Value),] ))
}


# -------------
# function for fisher method of the p-value combination
fisher.method <- function(p.values){
  df <- 2*length(p.values)
  return(pchisq( -2*sum(log(p.values)), df, lower.tail=FALSE))
}


# -------------
## annotation orthogroup
# add.species.column
annotation.orthogroups <- function(combined.dataframe.fisher,
                                   add.species.column, selected.genes.species){

  ## according to species
  fisher.combn.species <- combined.dataframe.fisher
  ## adding the rownames of each species ensembl gene ids
  rownames(fisher.combn.species) <- unlist(lapply(selected.genes.species, function(x) rownames(x)))
  ### fisher order per species - because of the enrichments
  fisher.combn.species.ordered <- fisher.combn.species[order(fisher.combn.species),, drop = FALSE]
  return(fisher.combn.species.ordered)
```

```
}


### -------------------------------------
# selecting the gene duplicates per species
# species.annotation:
# differential.expression.species: the table from differential expression analysis
# species.name: "Human", "Mouse", "Fly", "Worm"
selection.gene.duplicates.per.species <- function(species.annotation,
                                                  differential.expression.species, species.name){

  species <- lapply(species.annotation, function(x)
    differential.expression.species[na.omit(match(x, rownames(differential.expression.species))),])
  # correct the size of the gene set family
  species.gene.set.size.corrected <- lapply(species, function(x) { x$OG.adj.P.Val <- p.adjust(x$P.Value
                                                                                               method =
  ## Choose the lowest p-value in the gene-set family related to differential gene expression analysis
  species.lowest.pvalue.deg <- choose.the.lowest.pval(species.gene.set.size.corrected)
  # remove the empty dataframes from the list
  species.lowest.pvalue.df <- species.lowest.pvalue.deg[sapply(species.lowest.pvalue.deg,
                                                               function(x) dim(x)[1]) > 0]

  # quick check for the numbers
  cat(paste0("Number of selected OGs in ", species.name, ":" ),
      length(species.lowest.pvalue.df), "\n")

  return(species.lowest.pvalue.df)
}


# -------------------------------------
# proteostasis-linked processes
cc.proteostasis <- c("GO:0005783", # endoplasmic reticulum
                     "GO:0005789", # endoplasmic reticulum membrane
                      "GO:0101031", # chaperone complex
                      "GO:0005776", # autophagosome
                      "GO:0005764", # lysosome
                      "GO:0000502", # proteasome complex
                      "GO:0005739", # mitochondrion
                      "GO:0005759", # mitochondrial matrix
                      "GO:0005743", # mitochondrial inner membrane
                      "GO:0005730", # nucleolus
                      "GO:0005840") # ribosome

bp.proteostasis <- c(   "GO:0006412",     # translation
                        "GO:0006413",  # translational initiation
                        "GO:0006414",  # translational elongation
                        "GO:0006415",  # translational termination - didn't exist
                        "GO:0000028",   # ribosomal small subunit assembly
                        "GO:0000027",   # ribosomal large subunit assembly

                        "GO:0061684",  # chaperone-mediated autophagy
                        "GO:0006457",  # protein folding
```

```
                        "GO:0016236",  # Macroautophagy
                        "GO:0006614",  # SRP-dependent cotranslational protein targeting to membrane
                        "GO:0000398",  # mRNA splicing, via splicesome
                        "GO:0006508",  # proteolysis
                        "GO:0043248",  # proteasome assembly
                        "GO:0051290",  # protein heterotetramerization
                        "GO:0051289",  # protein homotetramerization
                        "GO:0006497",  # protein lipidation
                        "GO:0000209"   # protein polyubiquitination
                   )


### --------------------------------------
# intersect the orthologous groups between each species
integration.pvalues.across.species <- function(annotation.human, differential.human,
                                               annotation.mouse, differential.mouse,
                                               annotation.fly, differential.fly,
                                               annotation.worm, differential.worm){

  ## the intersection happens per tissue
  human <- selection.gene.duplicates.per.species(annotation.human, differential.human, "Human")
  mouse <- selection.gene.duplicates.per.species(annotation.mouse, differential.mouse, "Mouse")
  dmelano <- selection.gene.duplicates.per.species(annotation.fly, differential.fly, "Fly")
  cele <- selection.gene.duplicates.per.species(annotation.worm, differential.worm, "Worm")

  # how many common orthologous groups are
  intersected.tissue <- Reduce(intersect, list(names(human), names(mouse), names(dmelano), names(cele))
  cat("Number of selected OGs common to species:", length(intersected.tissue), "\n")


  dataframe.pvalues.species <-
    data.frame(human.pval = do.call(rbind, human[intersected.tissue])$OG.adj.P.Val,
                                    mouse.pval = do.call(rbind, mouse[intersected.tissue])$OG.adj.P.Val
                                    fly.pval = do.call(rbind, dmelano[intersected.tissue])$OG.adj.P.Val
                                    worm.pval = do.call(rbind, cele[intersected.tissue])$OG.adj.P.Val,
                                    human.logfc = do.call(rbind, human[intersected.tissue])$logFC,
                                    mouse.logfc = do.call(rbind, mouse[intersected.tissue])$logFC,
                                    fly.logfc = do.call(rbind, dmelano[intersected.tissue])$logFC,
                                    worm.logfc = do.call(rbind, cele[intersected.tissue])$logFC,
                                    human.gene.name = unlist(lapply(human[intersected.tissue], function
                                    mouse.gene.name = unlist(lapply(mouse[intersected.tissue], function
                                    fly.gene.name = unlist(lapply(dmelano[intersected.tissue], function
                                    worm.gene.name = unlist(lapply(cele[intersected.tissue], function(x
  rownames(dataframe.pvalues.species) <- intersected.tissue


  ### INTEGRATION: fisher p value combinations
  # focus only on first 4 columns with pvals
  dataframe.fishers.combined.pvalues <-
    as.data.frame(apply(dataframe.pvalues.species[,1:4], 1, fisher.method))
  colnames(dataframe.fishers.combined.pvalues) <- "combn.P.Val"
```

```r
  # giving them the annotations
  human.ordered.fishers.pvals <- annotation.orthogroups(dataframe.fishers.combined.pvalues,
                                                         "human.gene.name", human[intersected.tissue])
  mouse.ordered.fishers.pvals <- annotation.orthogroups(dataframe.fishers.combined.pvalues,
                                                        "mouse.gene.name", mouse[intersected.tissue])
  fly.ordered.fishers.pvals <- annotation.orthogroups(dataframe.fishers.combined.pvalues,
                                                      "fly.gene.name", dmelano[intersected.tissue])
  worm.ordered.fishers.pvals <- annotation.orthogroups(dataframe.fishers.combined.pvalues,
                                                       "worm.gene.name", cele[intersected.tissue])

  # first list (fishers.pvals) is the normal order, second is ordered according to the pvals
  return(list(raw.pvals.species = dataframe.pvalues.species,
              fishers.pvals = dataframe.fishers.combined.pvalues,
              fisher.pvals.human = human.ordered.fishers.pvals,
              fisher.pvals.mouse = mouse.ordered.fishers.pvals,
              fisher.pvals.dmelano = fly.ordered.fishers.pvals,
              fisher.pvals.cele = worm.ordered.fishers.pvals))
}


## annotation - changed to latest release
annotation.eogs <- function(columns, genome){
    require(biomaRt)
    # ensembl <- useEnsembl(biomart="ensembl", dataset=genome, version = 85)
    ensembl <- useMart(biomart="ENSEMBL_MART_ENSEMBL", host="www.ensembl.org",
                       path="/biomart/martservice",dataset=genome,
                                 version = "Ensembl Genes 91")
    annotation <- getBM(attributes = c("ensembl_gene_id", "gene_biotype", "description"),
                        values= columns, filter = "ensembl_gene_id",
                        mart = ensembl)
    annotation <- annotation[annotation$gene_biotype == "protein_coding", ]
    return(annotation)
}


final.annotation <- function(dataframe.pvals) {

  ## annotation
  anno.human <- annotation.eogs(dataframe.pvals$human.gene.name, "hsapiens_gene_ensembl")
  anno.mouse <- annotation.eogs(dataframe.pvals$mouse.gene.name, "mmusculus_gene_ensembl")
  anno.fly <- annotation.eogs(dataframe.pvals$fly.gene.name, "dmelanogaster_gene_ensembl")
  anno.worm <- annotation.eogs(dataframe.pvals$worm.gene.name, "celegans_gene_ensembl")

  ## associate with  dataframe
  mk <- match(dataframe.pvals$human.gene.name, anno.human$ensembl_gene_id)
  dataframe.pvals$human.gene.annotation <- anno.human[mk, "description"]

  mkm <- match(dataframe.pvals$mouse.gene.name, anno.mouse$ensembl_gene_id)
  dataframe.pvals$mouse.gene.annotation <- anno.mouse[mkm, "description"]

  mkf <- match(dataframe.pvals$fly.gene.name, anno.fly$ensembl_gene_id)
  dataframe.pvals$fly.gene.annotation <- anno.fly[mkf, "description"]
```

```r
  mkw <- match(dataframe.pvals$worm.gene.name, anno.worm$ensembl_gene_id)
  dataframe.pvals$worm.gene.annotation <- anno.worm[mkw, "description"]

  return(dataframe.pvals)
}






#### ------------------------------------
## GO enrichments for specific tissues
# input files:
# tissue.ogs: oma groups (evolutionary conserved gene sets across 4 species) per tissue
# toptable.tissue: dataframe from tissue specific differential expression analysis
# genome: genome from specific species
go.tissue.species.enrichment <- function(tissue.ogs, toptable.tissue, genome){

   tissue.ogs$FDR <- p.adjust(tissue.ogs$combn.P.Val, method = "fdr")
   tissue.ogs.sig <- tissue.ogs[which(tissue.ogs$FDR < 0.10), ]
   go.tissue.enrichment <- go.enrichment(rownames(toptable.tissue), rownames(tissue.ogs.sig), genome)
   return(go.tissue.enrichment)

 }



go.enrichment <- function(total.gene.names, significant.genes, genome){
  library(topGO)
  all.genes <- total.gene.names
  diff.genes <- significant.genes
  # giving the priority here to genes of interest
  relevant.genes <- factor(as.integer(all.genes %in% diff.genes))
  names(relevant.genes) <- all.genes
  head(relevant.genes)

  ont <- c("BP", "MF", "CC")

  ont.res <- list()
  for(i in 1:length(ont)){
    # forming the dataset
    GOdata <- new("topGOdata", ontology = ont[i], allGenes = relevant.genes,
                  geneSel = function(p) p < 1e-2, description = "Test",
                  annot = annFUN.org, mapping=genome, ID="Ensembl")

    resultFisher <- runTest(GOdata, algorithm = "elim", statistic = "fisher")

    res <- GenTable(GOdata, elimFisher = resultFisher, topNodes = 100) # make it bigger
    # this is correction for the processes
    corrected <- p.adjust(as.numeric(res$elimFisher), method="fdr")
    res$FDR <- corrected
    ont.res[[i]] <- res
```

```r
  }

  names(ont.res) <- ont
  return(ont.res)

}



#### for doing the heatmaps of GO clusters to aggregate towards bigger terms
# semantic matrix - the matrix from Wang method
# k - number of the clusters
# title - title of the heatmap
# ont - which ontology to use (BP, MF, CC)
heatmap.plotting <- function(semantic.matrix, k, title, ont, pdf.name){
  library(gplots)
  library(dendextend)
  # library(dendextendRcpp)
  library(RColorBrewer)
  library(GO.db)
  library(goseq)

  hclustfunc <- function(x) hclust(x, method="complete")
  distfunc <- function(x) dist(x,method="euclidean")


  # clustering according to the rows and columns and semantic matrix obtained from Wang method
  Rowv  <- semantic.matrix %>% distfunc %>% hclustfunc %>% as.dendrogram
  # set("branches_k_color", k = cl) %>% set("branches_lwd", 4) %>%
  # ladderize
  Colv  <- semantic.matrix %>% t %>% distfunc %>% hclustfunc %>% as.dendrogram
  # set("branches_k_color", k = cl) %>% set("branches_lwd", 4) %>%
  # ladderize


  #pdf(paste0(title, ".pdf"), 10, 10)
  pdf(pdf.name, 10, 10)
  heatmap.2(semantic.matrix,
           scale="none",
           Rowv =  Rowv,
           Colv =  Colv,
           col = colorRampPalette(brewer.pal(9,"Blues"))(100),
           key=TRUE,
           key.title = "",
           key.xlab = "GO semantic similarity score",
           key.par=list(mgp=c(3, 1, 0),
                        mar=c(5, 1.5, 5, 1.5)),
           symkey=FALSE,
           density.info="none",
           trace="none",
           cexRow=0.5,
           hclust=hclustfunc,
           distfun=distfunc,
```

```r
          lmat = rbind(c(0,3),c(2,1),c(0,4)),
          lwid = c(1.5,4),
          lhei = c(1.5,4,1),
          main = title)
  dev.off()

  # extract cluters
  find.clusters <- hclustfunc(distfunc(semantic.matrix))
  clusters <- cutree(find.clusters, k)


  if(ont == "BP"){
    parentsGO <- as.list(GOBPPARENTS)
    # remove the ones that do not have parents
    parentsGO <- parentsGO[!is.na(parentsGO)]
  }else if(ont == "MF"){
    parentsGO <- as.list(GOMFPARENTS)
    # remove the ones that do not have parents
    parentsGO <- parentsGO[!is.na(parentsGO)]
  }else{
    parentsGO <- as.list(GOCCPARENTS)
    # remove the ones that do not have parents
    parentsGO <- parentsGO[!is.na(parentsGO)]
  }

  term <- list()
  for(i in 1:k){
    cl <- parentsGO[names(clusters)[clusters == i]]
    clr <- goseq:::reversemapping(cl)
    term[[i]] <- cbind(Term(GOTERM[names(clr)]), sapply(clr, length))
    names(term)[i] <- paste("Cluster", i, sep = "_")

  }

  term <- lapply(term, function(x) {  colnames(x) <- c("Term", "Number.of.sets"); x  })
  return(term)

}


go.semantics <- function(topgo.process, species, ont){
  require(GOSemSim)
  d <- topgo.process$GO.ID
  dd <- godata(species, ont=ont, computeIC=FALSE)
  AGOsim <- matrix(0, nrow = length(d), ncol = length(d))
  # this is too slow - parallel computing helps
  for(i in 1:length(d)){
    for(j in 1:length(d)){
      # you can change the organism
      AGOsim[i,j] <- goSim(d[i],d[j], semData=dd, measure = "Wang")
    }
  }
```

```r
  rownames(AGOsim) <- colnames(AGOsim) <- d
  return(AGOsim)

}


# observed frequency
check.processes <- function(go.enrichment.data, bp.process, cc.process){

  df.cc.process <- go.enrichment.data$CC[na.omit(match(cc.process, go.enrichment.data$CC$GO.ID)), ]
  df.bp.process <-  go.enrichment.data$BP[na.omit(match(bp.process, go.enrichment.data$BP$GO.ID)), ]
  return(list(cc = df.cc.process, bp = df.bp.process))

}

# raw p-vals are after bonferroni correction
plotting.fisher.vs.raw <- function(eogs, main){
  par(mfrow = c(2,2))
  ph <- -log10(eogs$raw.pvals.species[,1])
  p2 <- -log10(eogs$fishers.pvals$combn.P.Val)

  fit1 <- lm(p2~ph)
  smoothScatter(ph,p2, main = main[1],
                xlab = "-log10 Human DEG p-val",
                ylab = "-log10 Fisher combined p-val", xlim = c(0,8))
  abline(fit1)

  pm <- -log10(eogs$raw.pvals.species[,2])
  fit2 <- lm(p2~pm)
  smoothScatter(pm, p2, main =  main[2],
                xlab = "-log10 Mouse DEG p-val",
                ylab = "-log10 Fisher combined p-val", xlim = c(0,8))
  abline(fit2)

  pf <- -log10(eogs$raw.pvals.species[,3])
  fit3 <- lm(p2~pf)
  smoothScatter(pf, p2, main =  main[3],
                xlab = "-log10 Fly DEG p-val",
                ylab = "-log10 Fisher combined p-val", xlim = c(0,8))
  abline(fit3)

  pw <- -log10(eogs$raw.pvals.species[,4])
  fit4 <- lm(p2~pw)
  smoothScatter(pw, p2, main =  main[4],
                xlab = "-log10 Worm DEG p-val",
                ylab = "-log10 Fisher combined p-val", xlim = c(0,8))
  abline(fit4)
}
```

##### Load the data

```r
path <- "~/Project1/manuscript_GSEA/data_preprocessing/"
```

**Analysis using OMA groups (hierarchical orthologous groups)**

```
# ---------------------------------------
# Load evolutionary orthologous groups - HOGs
oma.groups <-  readRDS(paste0(path, "/oma/oma_hogs.rds"))
length(oma.groups) # http://omabrowser.org/oma/home/
head(oma.groups)
# 3232 orthogroups

## sizes of the groups
range(lapply(oma.groups, function(x) length(x))) # from 4 to 246
# hist(do.call(c, lapply(oma.groups, function(x) length(x))))

# Human
human.oma.genes <- lapply(oma.groups, function(x) x[grepl("^ENSG", x, perl = TRUE)])
# M. musculus
mouse.oma.genes <- lapply(oma.groups, function(x) x[grepl("^ENSMUSG", x, perl = TRUE)])

# D. melanogaster
fly.oma.genes <- lapply(oma.groups, function(x) x[grepl("^FB", x, perl = TRUE)])
# C. elegans
worm.oma.genes <- lapply(oma.groups, function(x) x[grepl("^WB", x, perl = TRUE)])
```

Loading differentially expressed matrices.

```
# -------------------------
# Human
deg.muscle.human.aging <-
  readRDS(paste0(path, "hsapiens/aging_data/diff_exp/topTable_Muscle_Skeletal_GTEx_V6p.rds"))
deg.hippo.human.aging <-
  readRDS(paste0(path, "hsapiens/aging_data/diff_exp/topTable_Brain_Hippocampus_GTEx_V6p.rds"))


# --------------------------
# Mouse


deg.mouse.muscle <-
  readRDS(paste0(path,
                 "mmusculus/aging_data/skeletal_muscle/diffexp_aging_mouse_skeletal_muscle_aging.rds"))
deg.muscle.mouse.aging <- deg.mouse.muscle$oldvsyoung

# change here
deg.hippo.mouse.aging <-
  readRDS(paste0(path,
                 "mmusculus/aging_data/hippocampus/diff_expression_mmusculus_hippocampus_aging.rds"))

# --------------------------
# Fly - wb
deg.fly <-
  readRDS(paste0(path, "dmelanogaster/aging_data/diff_expression_fly_aging.rds"))
deg.wholebody.fly.aging <- deg.fly$oldvsyoung

# --------------------------
# Worm - wb
```

```
deg.wholebody.worm.aging <-
  readRDS(paste0(path, "celegans/aging_data/diff_expression_celegans_aging.rds"))
```

```
## Dietary restriction
deg.human.dr <-
  readRDS(paste0(path,
                 "hsapiens/caloric_restriction_data/differential_expression_human_dietary_restriction.r
## Dietary restriction
deg.mouse.dr <-
  readRDS(paste0(path,
                 "mmusculus/caloric_restriction_data/differential_expression_mouse_dietary_restriction.
## Dietary restriction - rename this
deg.fly.dr <-
  readRDS(paste0(path,
                 "/dmelanogaster/caloric_restriction_data/diff_exp_fly_dietary_restriction.rds"))
## Dietary restriction
deg.worm.dr <-
  readRDS(paste0(path,
                 "/celegans/caloric_restriction_data/diff_exp_worm_dietary_restriction.rds"))
```

##### Annotation of orthogroups

```
# Analysis

# muscle
muscle.common.eogs.oma <-
  integration.pvalues.across.species(human.oma.genes, deg.muscle.human.aging,
                                     mouse.oma.genes, deg.muscle.mouse.aging,
                                     fly.oma.genes, deg.wholebody.fly.aging,
                                     worm.oma.genes, deg.wholebody.worm.aging)
```

```
## Number of selected OGs in Human: 3069
## Number of selected OGs in Mouse: 2836
## Number of selected OGs in Fly: 3036
## Number of selected OGs in Worm: 3160
## Number of selected OGs common to species: 2511
```

```
nrow(muscle.common.eogs.oma[[1]]) # 2511 now
```

```
## [1] 2511
```

```
# hippocampus
hippocampus.common.eogs.oma <-
  integration.pvalues.across.species(human.oma.genes, deg.hippo.human.aging,
                                     mouse.oma.genes, deg.hippo.mouse.aging,
                                     fly.oma.genes, deg.wholebody.fly.aging,
                                     worm.oma.genes, deg.wholebody.worm.aging)
```

```
## Number of selected OGs in Human: 3096
## Number of selected OGs in Mouse: 3136
## Number of selected OGs in Fly: 3036
## Number of selected OGs in Worm: 3160
## Number of selected OGs common to species: 2800
```

```
nrow(hippocampus.common.eogs.oma[[1]]) # 2800 now
```

```
## [1] 2800
```

```r
# I have only on the muscle
muscle.common.eogs.oma.dr <-
  integration.pvalues.across.species(human.oma.genes, deg.human.dr,
                                     mouse.oma.genes, deg.mouse.dr,
                                     fly.oma.genes, deg.fly.dr,
                                     worm.oma.genes, deg.worm.dr)
```

```
## Number of selected OGs in Human: 2359
## Number of selected OGs in Mouse: 2836
## Number of selected OGs in Fly: 3113
## Number of selected OGs in Worm: 3100
## Number of selected OGs common to species: 1971
```

```r
nrow(muscle.common.eogs.oma.dr[[1]]) # 1971 now
```

```
## [1] 1971
```

The plots with the Fisher and raw p-values. Supplement S5. Fisher plots.

```r
# pdf("~/Project1/manuscript_GSEA/results/FigureS5A_Fisher_plots.pdf", 10, 7)
# par(frow = c(1, 2))
plotting.fisher.vs.raw(muscle.common.eogs.oma,
                       c("Human DEG Muscle vs Fisher p-val", "Mouse DEG Muscle vs Fisher p-val",
                         "Fly DEG Muscle vs Fisher p-val", "Worm DEG Muscle vs Fisher p-val"))
```

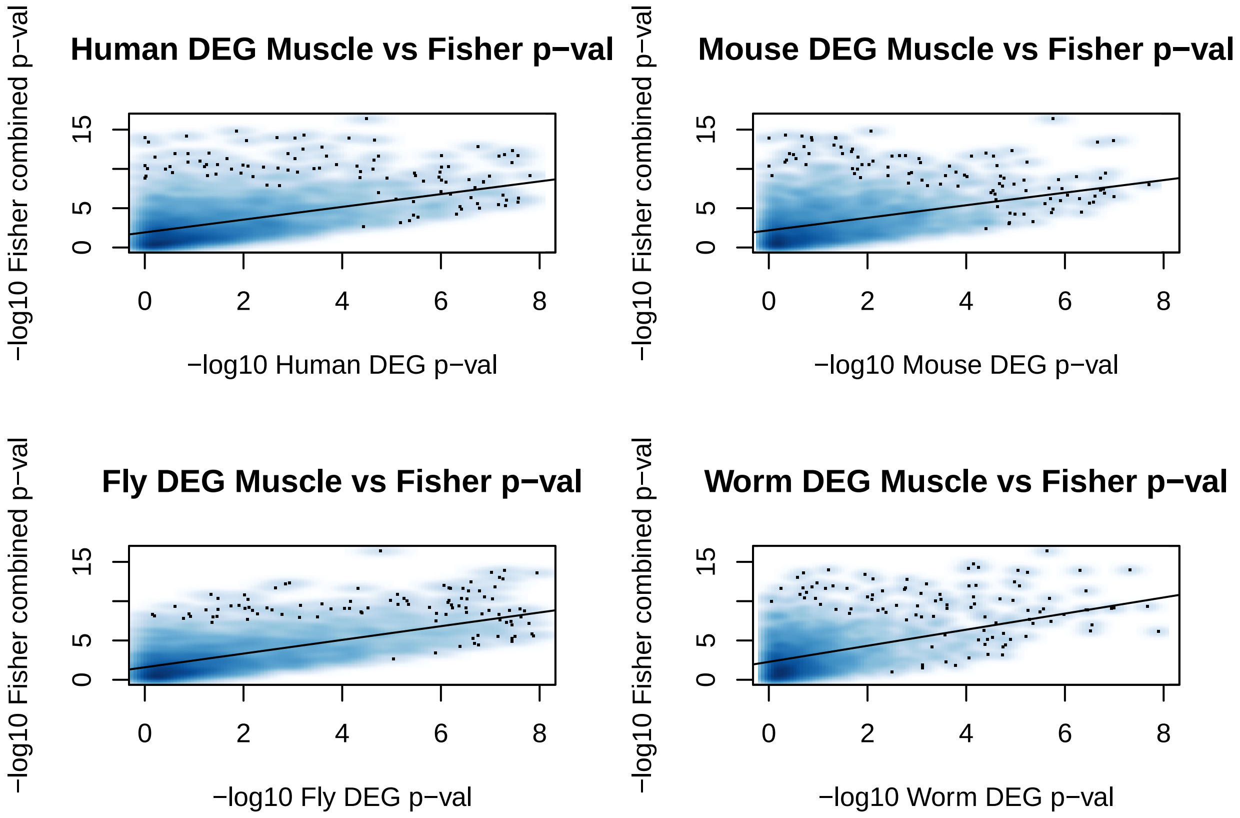


```r
# dev.off()
```

```r
# pdf("~/Project1/manuscript_GSEA/results/FigureS5B_Fisher_plots.pdf", 10, 7)
plotting.fisher.vs.raw(hippocampus.common.eogs.oma,
```

```
                          c("Human DEG Hippo vs Fisher p-val", "Mouse DEG Hippo vs Fisher p-val",
                            "Fly DEG Hippo vs Fisher p-val", "Worm DEG Hippo vs Fisher p-val"))
```

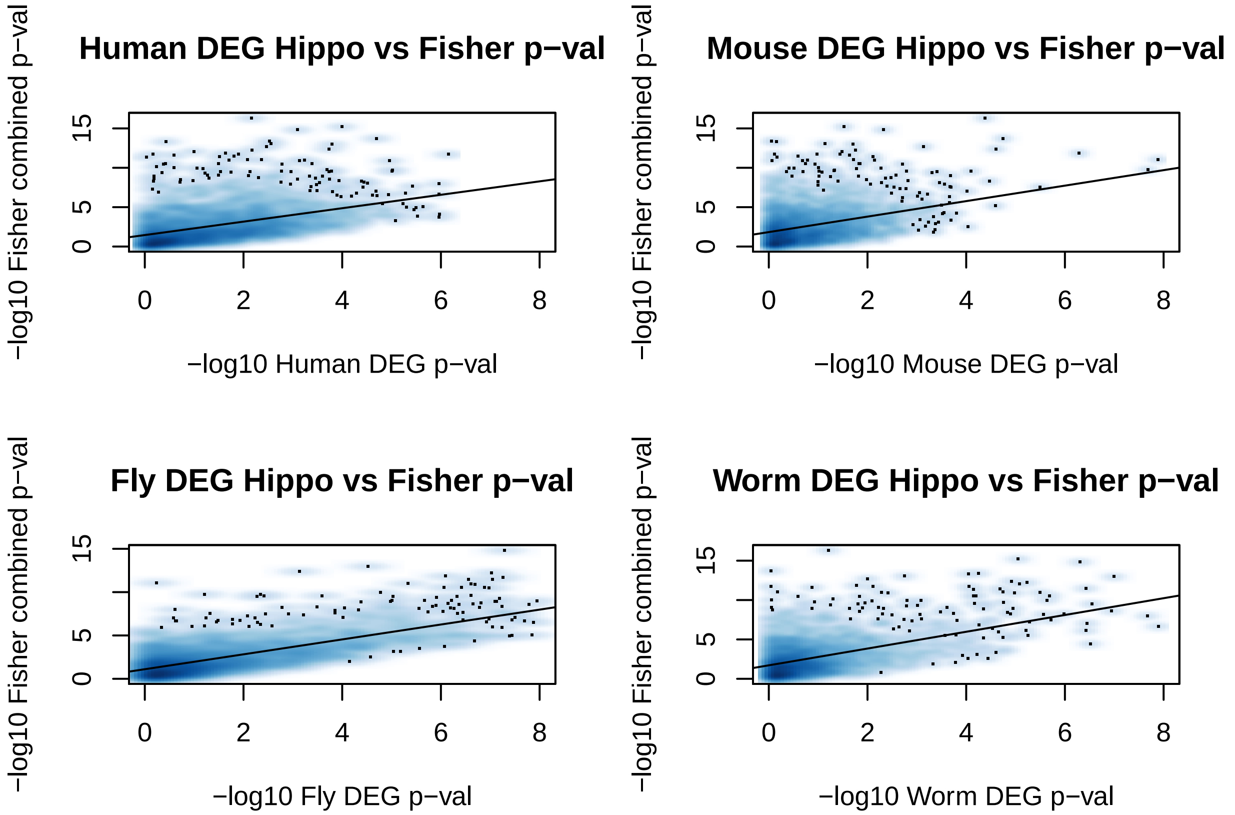


```
# dev.off()
```

###### Borrowing the information from species to human

General conservation of the processes across species using orthogroups. Here the mappings are on the human annotation, because we borrowed information from other species.

```
##  skeletal muscle
fisher.muscle.human.oma <- muscle.common.eogs.oma$fisher.pvals.human
human.go.muscle.oma <-  go.tissue.species.enrichment(fisher.muscle.human.oma, deg.muscle.human.aging, "
# GOSemSim v2.4.1
human.muscle.summary.go.oma <-  go.semantics(human.go.muscle.oma$BP, 'org.Hs.eg.db', "BP")
clusters.human.muscle.hog.oma <- heatmap.plotting(human.muscle.summary.go.oma, 11,
                                                  "Human HOG GO BP Muscle", "BP",
                                                  "~/Project1/manuscript_GSEA/results/check_Human_Muscl

names(clusters.human.muscle.hog.oma) <- c("Cellular respiration",
                                          "Transport",
                                          "Translation",
                                          "Transcription and post-transcriptional modifications",
                                          "Cell division",
                                          "Protein folding",
                                          "Protein ubiquitination",
                                          "Cofactor biosynthetic process",
```

```r
                                          "Immune response",
                                          "Cell differentiation",
                                          "Protein-containing complex assembly")


## hippocampus
fisher.hippo.human.oma <- hippocampus.common.eogs.oma$fisher.pvals.human
human.go.hippo.oma <- go.tissue.species.enrichment(fisher.hippo.human.oma, deg.hippo.human.aging, "org.
human.hippo.summary.go.oma <-  go.semantics(human.go.hippo.oma$BP, 'org.Hs.eg.db', "BP")
clusters.human.hippo.hog.oma <- heatmap.plotting(human.hippo.summary.go.oma, 13,
                                                 "Human HOG GO BP Hippo", "BP",
                                                 "~/Project1/manuscript_GSEA/results/check_Human_Hippo_

names(clusters.human.hippo.hog.oma) <- c( "Transport",
                                          "Translation",
                                          "Transcription and post-transcriptional modifications",
                                          "Transcription and post-transcriptional modifications", #
                                          "Cellular amino acid metabolic process",
                                          "Protein folding",
                                          "Cellular respiration",
                                          "Cofactor biosynthetic process",
                                          "Immune response",
                                         "Cofactor biosynthetic process",
                                         "Cell division",
                                         "Protein-containing complex assembly",
                                         "Protein ubiquitination" )




## dietary restriction
fisher.muscle.human.oma.dr <- muscle.common.eogs.oma.dr$fisher.pvals.human
human.go.muscle.oma.dr <- go.tissue.species.enrichment(fisher.muscle.human.oma.dr, deg.human.dr, "org.H
human.muscle.summary.go.oma.dr <-  go.semantics(human.go.muscle.oma.dr$BP, 'org.Hs.eg.db', "BP")


# dietary restriction - human - done
clusters.human.muscle.hog.oma.dr <- heatmap.plotting(human.muscle.summary.go.oma.dr, 10,
                                                     "Human HOG GO BP Muscle DR", "BP",
                                                     "~/Project1/manuscript_GSEA/results/check_Human_DR

names(clusters.human.muscle.hog.oma.dr) <- c( "Transport",
                                              "Translation",
                                              "Transcription and post-transcriptional modifications",
                                              "Cellular amino acid metabolic process",
                                              "Cell division",
                                              "Protein ubiquitination",
                                              "Protein-containing complex assembly",
                                              "Cell differentiation",
                                              "Ribosome biogenesis",
                                              "Immune response"
                                              )

## plotting the conserved processes
```

```r
# change this to caloric restriction

df.muscle <- data.frame(Name = names(clusters.human.muscle.hog.oma),
                        Size = sapply(clusters.human.muscle.hog.oma, nrow),
                        Tissue = rep("Skeletal muscle (aging)",length(clusters.human.muscle.hog.oma)))

df.hippo <- data.frame(Name = names(clusters.human.hippo.hog.oma),
                       Size = sapply(clusters.human.hippo.hog.oma, nrow),
                       Tissue = rep("Hippocampus (aging)",length(clusters.human.hippo.hog.oma)))

df.dietary <- data.frame(Name = names(clusters.human.muscle.hog.oma.dr),
                       Size = sapply(clusters.human.muscle.hog.oma.dr, nrow),
                       Tissue = rep("Skeletal muscle (caloric restriction)", length(clusters.human.muscl

integrated.pathways <- rbind(df.muscle, df.hippo, df.dietary)


# pdf("~/Project1/manuscript_GSEA/results/Figure2B_GOcategories_Aging_Caloric_Sep_re.pdf", 10, 7, useDi
p <- ggplot(integrated.pathways,aes(Tissue, Name, colour = factor(Tissue))) +
            scale_colour_brewer(palette="Paired") +
            geom_point(aes(size = Size)) + ylab("GO categories") + xlab("") +
            theme(axis.text.x=element_text(size="10", color="black",  angle = 45, hjust = 1),
                  panel.grid.minor=element_line(color="ivory3",size=0.5),
                  panel.grid.major=element_line(color="ivory3",size=0.5)) +
            labs(size = "No. of significant\ngene sets conserved\nacross 4 species")

p + guides(colour=FALSE)
```

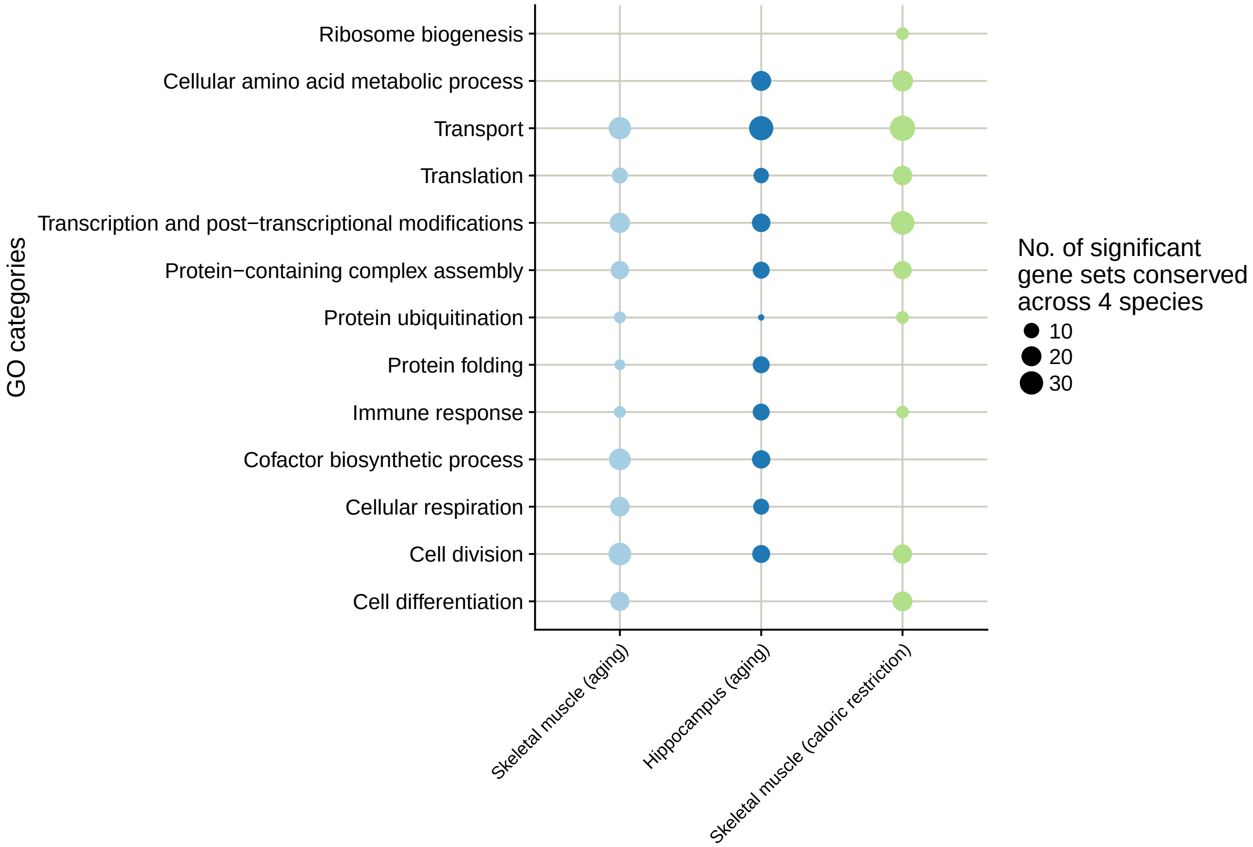


```
# dev.off()
```

Generating Figure 2C. with GO enrichments.

```
## ---------------------------

# here should go the significant data
preparing.proteostasis <- function(go.enrichment.data, bp.process, cc.process){

  df.human.cc.process <- go.enrichment.data$CC[na.omit(match(cc.process, go.enrichment.data$CC$GO.ID)),
  # calculating the enrichment score
  df.human.cc.process$LogEnrich <- log2(df.human.cc.process$Significant / df.human.cc.process$Expected)
  df.human.cc.process$Category <- "GO CC Term"


  df.human.bp.process <-  go.enrichment.data$BP[na.omit(match(bp.process, go.enrichment.data$BP$GO.ID))
  # calculate the enrichment score
  df.human.bp.process$LogEnrich <- log2(df.human.bp.process$Significant / df.human.bp.process$Expected)
  df.human.bp.process$Category <- "GO BP Term"

  ## to change to the capital letters
  simpleCap <- function(x) {
    s <- strsplit(x, " ")[[1]]
    paste(toupper(substring(s, 1,1)), substring(s, 2),
        sep="", collapse=" ")
  }

  # looking only in human
```

```r
  df.human.process <- rbind(df.human.cc.process, df.human.bp.process)
  df.human.process$Term <- sapply(df.human.process$Term, simpleCap)
  df.human.process$Term <- factor(df.human.process$Term, levels = df.human.process$Term)

  return(df.human.process)

}


### checking the proteostasis-linked processes in enriched integrated list
human.muscle.proteostasis <- preparing.proteostasis(human.go.muscle.oma,
                                                    bp.proteostasis, cc.proteostasis)
human.hippo.proteostasis <- preparing.proteostasis(human.go.hippo.oma,
                                                   bp.proteostasis, cc.proteostasis)
human.dr.proteostasis <- preparing.proteostasis(human.go.muscle.oma.dr,
                                                   bp.proteostasis, cc.proteostasis)


### plotting the enrichments scores for significant GO proteostasis-linked processes
## for muscle enrichments

# remove the background
ggplot(human.muscle.proteostasis, aes(Term, LogEnrich, fill = factor(Category))) + theme_classic() +
                      scale_fill_brewer("", palette="Paired") +
                      geom_bar(position= "identity", stat="identity", width=0.8) +
                      coord_flip() +
                      scale_y_continuous(position = "right") +
                      labs( y = "GO log2Enrichment score (skeletal muscle)") +
                      theme(axis.text.x = element_text(size=14),
                            axis.text.y = element_text(size=14),
                            text = element_text(size=14))
```

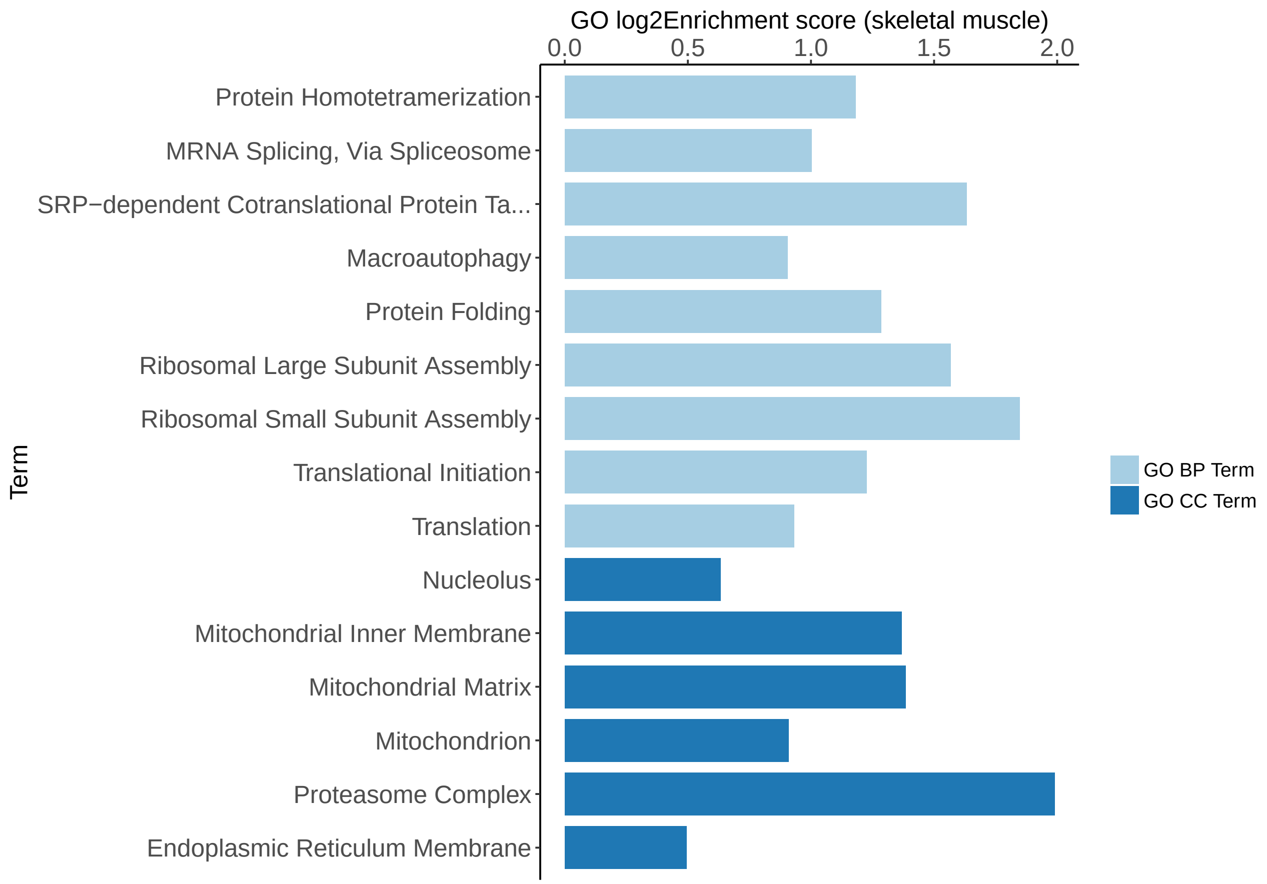


```
    # ggsave("~/Project1/manuscript_GSEA/results/Figure2C_Proteostasis_Enrichment_Human_Muscle.pdf",
```

Generating Figure 3. with GO terms but mapped to GTEx data. The genes are conserved across the species.

```
get.go.annotation <- function(deg.species.aging, fisher.pvals.species,
                              processes, dataframe.pvals.logfc,
                              species.name, species.name.logfc, genome, ont){

  fisher.pvals.species$FDR <- p.adjust(fisher.pvals.species$combn.P.Val, method = "fdr")
  species.significant.fisher.pvals <- fisher.pvals.species[which(fisher.pvals.species$FDR < 0.10), ]


  total.genes <- rownames(deg.species.aging)
  diff.genes <- rownames(species.significant.fisher.pvals)
  relevant.genes <- factor(as.integer(total.genes %in% diff.genes))
  names(relevant.genes) <- total.genes


  GOdata <- new("topGOdata", ontology = ont, allGenes = relevant.genes,
                geneSel = function(p) p < 1e-2, description = "Test",
                annot = annFUN.org, mapping=genome, ID="Ensembl")

  sel.terms.proteo <- processes
  # num.ann.genes.proteo <- countGenesInTerm(GOdata, sel.terms.proteo)
  ann.genes.proteo <- genesInTerm(GOdata, sel.terms.proteo)
# list of the terms based on human
  annotation.proteo <- lapply(ann.genes.proteo, function(x) x[x %in% diff.genes] )
```

```
  return(annotation.proteo)

}



## preparation for volcano plots
# to extract the annotation
volcano.plots.preparation <- function(differential.expression,
                                      common.eogs, processes, ont, annotation.fisher){

  annotation.ont <- get.go.annotation( differential.expression, common.eogs$fisher.pvals.human,
                                       processes, common.eogs$raw.pvals.species,
                                       "human.gene.name", "human.logfc", "org.Hs.eg.db", ont)
  annotation.stats <- lapply(annotation.ont, function(x)
                            annotation.fisher[na.omit(match(x, annotation.fisher$human.gene.name)), ])

  human.deg <- lapply(annotation.stats, function(x) {
                                df.deg <- differential.expression[match(x$human.gene.name,
                                                                        rownames(differential.expressio
                                df.deg$Gene <- mapIds(org.Hs.eg.db,
                                                        keys=rownames(df.deg),
                                                        column="SYMBOL",
                                                        keytype="ENSEMBL",
                                                        multiVals="first")
                                df.deg$Significant <- ifelse(df.deg$adj.P.Val < 0.05,
                                                             "FDR < 0.05", "FDR > 0.05")
                                return(df.deg)} )
  return(human.deg)


}


# ------------------------------------------
# annotation
annotation.muscle.oma <- final.annotation(muscle.common.eogs.oma$raw.pvals.species)
annotation.muscle.oma.fisher <- cbind(muscle.common.eogs.oma$fishers.pvals, annotation.muscle.oma)

# saveRDS(annotation.muscle.oma, file = "annotation_muscle_oma.rds")



annotation.hippo.oma <- final.annotation(hippocampus.common.eogs.oma$raw.pvals.species)
annotation.hippo.oma.fisher <- cbind(hippocampus.common.eogs.oma$fishers.pvals, annotation.hippo.oma)

# saveRDS(annotation.hippo.oma, file = "annotation_hippocampus_oma.rds")
```

```
### ------------------------------------------
# human skeletal muscle

human.muscle.bp.deg <- volcano.plots.preparation(deg.muscle.human.aging, muscle.common.eogs.oma,
                                                 bp.proteostasis, "BP", annotation.muscle.oma.fisher)
```

```r
human.muscle.cc.deg <- volcano.plots.preparation(deg.muscle.human.aging, muscle.common.eogs.oma,
                                                  cc.proteostasis, "CC", annotation.muscle.oma.fisher)


# ------------------------------------------
# hippocampus
human.hippo.bp.deg <- volcano.plots.preparation(deg.hippo.human.aging, hippocampus.common.eogs.oma,
                                                bp.proteostasis, "BP", annotation.hippo.oma.fisher)

human.hippo.cc.deg <- volcano.plots.preparation(deg.hippo.human.aging, hippocampus.common.eogs.oma,
                                                cc.proteostasis, "CC", annotation.hippo.oma.fisher)





renaming.pvals <- function(list.species){

  for(i in 1:length(list.species)){
      list.species[[i]]$Gene[list.species[[i]]$Significant == "FDR > 0.05"] <- NA
      list.species[[i]][is.na(list.species[[i]])] <- ""
  }

  return(list.species)
}


human.muscle.cc.deg <- renaming.pvals(human.muscle.cc.deg)
human.muscle.bp.deg <- renaming.pvals(human.muscle.bp.deg)

human.hippo.bp.deg <- renaming.pvals(human.hippo.bp.deg)
human.hippo.cc.deg <- renaming.pvals(human.hippo.cc.deg)
```

```r
# ----------------------------------------------
# important parts of the protein quality control network
# proteasome complex
# translation
# macroautophagy
protein.quality.control.network <- c( "GO:0000502", "GO:0006412", "GO:0016236" )

library(ggrepel)
library(cowplot)
# geom_text_repel(aes(wt, mpg, label = rownames(mtcars)))

p.proteasome.muscle <- ggplot(human.muscle.cc.deg[[protein.quality.control.network[1]]],
                              aes(x = logFC, y = -log10(adj.P.Val))) +
  geom_point(aes(color = Significant)) +
  geom_text_repel(data = subset(human.muscle.cc.deg[[protein.quality.control.network[1]]],
                                adj.P.Val < 0.05),
                                 aes(label = Gene), size = 2.5) +
  scale_color_manual(values = c("#268bd2", "grey")) +
  theme_bw(base_size = 12) + theme(legend.position = "bottom") +
  ggtitle("GO:0000502 proteasome complex\n (GTEx Skeletal Muscle)") +
  xlab( expression(log[2]~fold~change~(Old/Young))) + ylab(expression(-log[10]~FDR)) +
```

```r
  ylim(0, 8) + xlim(-0.01, 0.01) +
  geom_vline(xintercept = 0)


p.translation.muscle <- ggplot(human.muscle.bp.deg[[protein.quality.control.network[2]]],
                               aes(x = logFC, y = -log10(adj.P.Val))) +
        geom_point(aes(color = Significant)) +
        geom_text_repel(data = subset(human.muscle.bp.deg[[protein.quality.control.network[2]]],
                                      adj.P.Val < 0.05),
                                      aes(logFC, -log10(adj.P.Val),
                                      label = Gene), size = 2.5) +
        scale_color_manual(values = c("#6c71c4", "grey")) +
        theme_bw(base_size = 12) + theme(legend.position = "bottom") +
        ggtitle("GO:0006412 translation\n (GTEx Skeletal Muscle)") +
        xlab( expression(log[2]~fold~change~(Old/Young))) + ylab(expression(-log[10]~FDR)) +
        ylim(0, 8) + xlim(-0.01, 0.01) +
        geom_vline(xintercept = 0)



p.macroautophagy.muscle <- ggplot(human.muscle.bp.deg[[protein.quality.control.network[3]]],
                                  aes(x = logFC, y = -log10(adj.P.Val))) +
    geom_text_repel(
        data = subset(human.muscle.bp.deg[[protein.quality.control.network[3]]], adj.P.Val < 0.05),
        aes(label = Gene), size = 2.5) +
        geom_point(aes(color = Significant)) +
        scale_color_manual(values = c("#2aa198", "grey")) +
        theme_bw(base_size = 12) + theme(legend.position = "bottom") +
        ggtitle("GO:0016236 macroautophagy\n (GTEx Skeletal Muscle)") +
        xlab( expression(log[2]~fold~change~(Old/Young))) + ylab(expression(-log[10]~FDR)) +
        ylim(0, 8) + xlim(-0.01, 0.01) +
        geom_vline(xintercept = 0)

# pdf("~/Project1/manuscript_GSEA/results/Figure3ABC_volcano_plots_proteostasis_network_re.pdf",
#10, 7, useDingbats = FALSE)
plot_grid(p.macroautophagy.muscle,
          p.translation.muscle,
          p.proteasome.muscle, labels = c("", "", ""), align = "h")
```

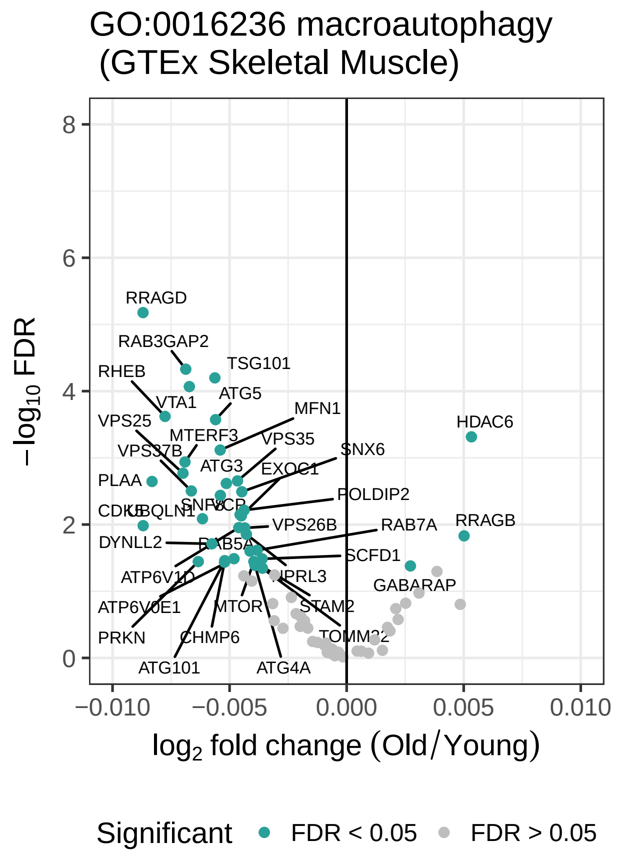
GO:0016236 macroautophagy
(GTEx Skeletal Muscle)
−log10 FDR
log2 fold change (Old/Young)
−0.010
−0.005
0.000
0.005
0.010
0
2
4
6
8
RRAGD
RAB3GAP2
RHEB
TSG101
VTA1
ATG5
VPS25
MTERF3
MFN1
VPS37B
VPS35
HDAC6
SNX6
ATG3
EXOC1
PLAA
POLDIP2
CDK5
UBQLN1
SNF8
VCP
VPS26B
RAB7A
RRAGB
DYNLL2
RAB5A
SCFD1
ATP6V1D
WDR41
MTOR
PRKN
CHMP6
ATG101
ATG4A
ATP6V0E1
IPRL3
STAM2
GABARAP
TOMM22
Significant
FDR < 0.05
FDR > 0.05

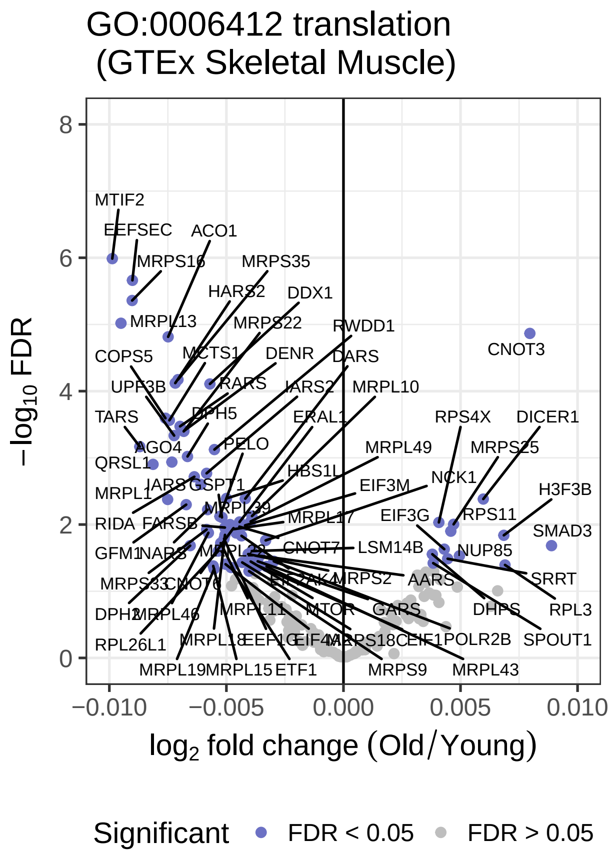
GO:0006412 translation
(GTEx Skeletal Muscle)
−log10 FDR
log2 fold change (Old/Young)
−0.010
−0.005
0.000
0.005
0.010
0
2
4
6
8
MTIF2
EEFSEC
ACO1
MRPS16
MRPS35
HARS2
DDX1
MRPL13
MRPS22
RWDD1
CNOT3
COPS5
MCTS1
DENR
DARS
UPF3B
RARS
IARS2
MRPL10
TARS
DPH5
ERAL1
RPS4X
DICER1
AGO4
PELO
MRPL49
MRPS25
QRSL1
HBS1L
NCK1
H3F3B
MRPL1
IARS
GSPT1
EIF3M
RIDA
FARSB
MRPL39
MRPL17
EIF3G
RPS11
SMAD3
GFM1
NARS
MRPL22
CNOT7
LSM14B
NUP85
MRPS33
CNOT6
EIF2AK4
MRPS2
AARS
SRRT
DPH2
MRPL46
MRPL11
MTOR
GARS
DHPS
RPL3
RPL26L1
MRPL18
EEF1G
EIF4A3
MRPS18C
EIF1
POLR2B
SPOUT1
MRPL19
MRPL15
ETF1
MRPS9
MRPL43
Significant
FDR < 0.05
FDR > 0.05

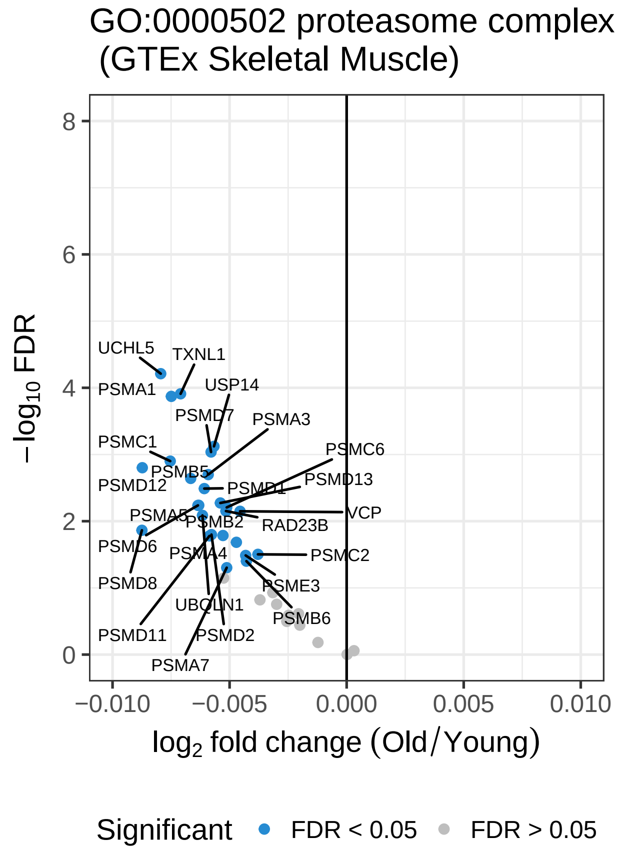
GO:0000502 proteasome complex
(GTEx Skeletal Muscle)
−log10 FDR
log2 fold change (Old/Young)
−0.010
−0.005
0.000
0.005
0.010
0
2
4
6
8
UCHL5
TXNL1
PSMA1
USP14
PSMD7
PSMA3
PSMC1
PSMC6
PSMB5
PSMD13
PSMD12
PSMD1
VCP
PSMA5
PSMB2
RAD23B
PSMD6
PSMA4
PSMC2
PSMD8
PSME3
UBQLN1
PSMB6
PSMD11
PSMD2
PSMA7
Significant
FDR < 0.05
FDR > 0.05

```
# dev.off()
```

**Supplementary Data.**

Generating the Table S4.

```
## tissue annotation
tissue.annotation.eogs <- function(experiment.common.eogs.oma){

  annotation.exp.oma <- final.annotation(experiment.common.eogs.oma$raw.pvals.species)
  annotation.exp.oma.fisher <- cbind(experiment.common.eogs.oma$fishers.pvals, annotation.exp.oma)

  # doesn't matter is it human or others
  fisher.tissue.human.oma <- experiment.common.eogs.oma$fisher.pvals.human
  tissue.human.eogs <- fisher.tissue.human.oma
  # correct integrated p vals
  tissue.human.eogs$FDR <- p.adjust(tissue.human.eogs$combn.P.Val,"fdr")
  # threshold seemed reasonable
  tissue.human.eogs.sig <- tissue.human.eogs[which(tissue.human.eogs$FDR < 0.10), ]

  print(nrow(tissue.human.eogs.sig))

  return(list(annotation.tissue = annotation.exp.oma.fisher,
              number.significant.genes = tissue.human.eogs.sig))
}



muscle.integration <- tissue.annotation.eogs(muscle.common.eogs.oma)
```

```
## [1] 2010
```

```
hippo.integration <- tissue.annotation.eogs(hippocampus.common.eogs.oma)
```

```
## [1] 2075
```

```
dr.integration <- tissue.annotation.eogs(muscle.common.eogs.oma.dr)
```

```
## [1] 1962
```

```
# fly is na
# check do you have rownames - ENSEMBL and gene names
integration.list <- list(Muscle_Healthy_Aging =  muscle.integration$annotation.tissue,
                         Hippocampus_Healthy_Aging = hippo.integration$annotation.tissue,
                         Dietary_restriction = dr.integration$annotation.tissue)



# Supplement Table S4 - TO DO
# WriteXLS(integration.list,
#          ExcelFileName = "~/Project1/manuscript_GSEA/supplementary_data/Table_S4.xlsx", SheetNames = N
```

Generating the Table S5.

```
library(plyr)
df.clusters.muscle.aging <-
```

```
  ldply(clusters.human.muscle.hog.oma, data.frame, .id = "Cluster.Name")
df.clusters.hippo.aging <-
  ldply(clusters.human.hippo.hog.oma, data.frame, .id = "Cluster.Name")
df.clusters.muscle.aging.dr <-
  ldply(clusters.human.muscle.hog.oma.dr, data.frame, .id = "Cluster.Name")

df.clusters.figure2 <-
  list(ConservedCl.MuscleAging = df.clusters.muscle.aging,
             ConservedCl.HippoAging = df.clusters.hippo.aging,
             ConservedCl.MuscleDR = df.clusters.muscle.aging.dr)


# WriteXLS(df.clusters.figure2,
#          ExcelFileName = "~/Project1/manuscript_GSEA/supplementary_data/Table_S5.xlsx", SheetNames = 
```

Generating Table S6.

```
df.proteostasis.processes <- list(Muscle_Human_Proteostasis = human.muscle.proteostasis,
                                                                  Hippo_Human_Proteostasis = human.hipp
                                                                  DR_Human_Proteostasis = human.dr.prot


# WriteXLS(df.proteostasis.processes,
#          ExcelFileName = "~/Project1/manuscript_GSEA/supplementary_data/Table_S6.xlsx", SheetNames = 
```

Generating Supplement Figure S7

```
## hippocampus enrichments
ggplot(human.hippo.proteostasis, aes(Term, LogEnrich, fill = factor(Category))) +
                      scale_fill_brewer("", palette="Paired") +
                      geom_bar(position= "identity", stat="identity", width=0.8) +
                      coord_flip() +
                      scale_y_continuous(position = "right") +
                      labs( y = "GO log2Enrichment score (hippocampus)") +
                      theme(axis.text.x = element_text(size=14),
                            axis.text.y = element_text(size=14),
                            text = element_text(size=14))
```

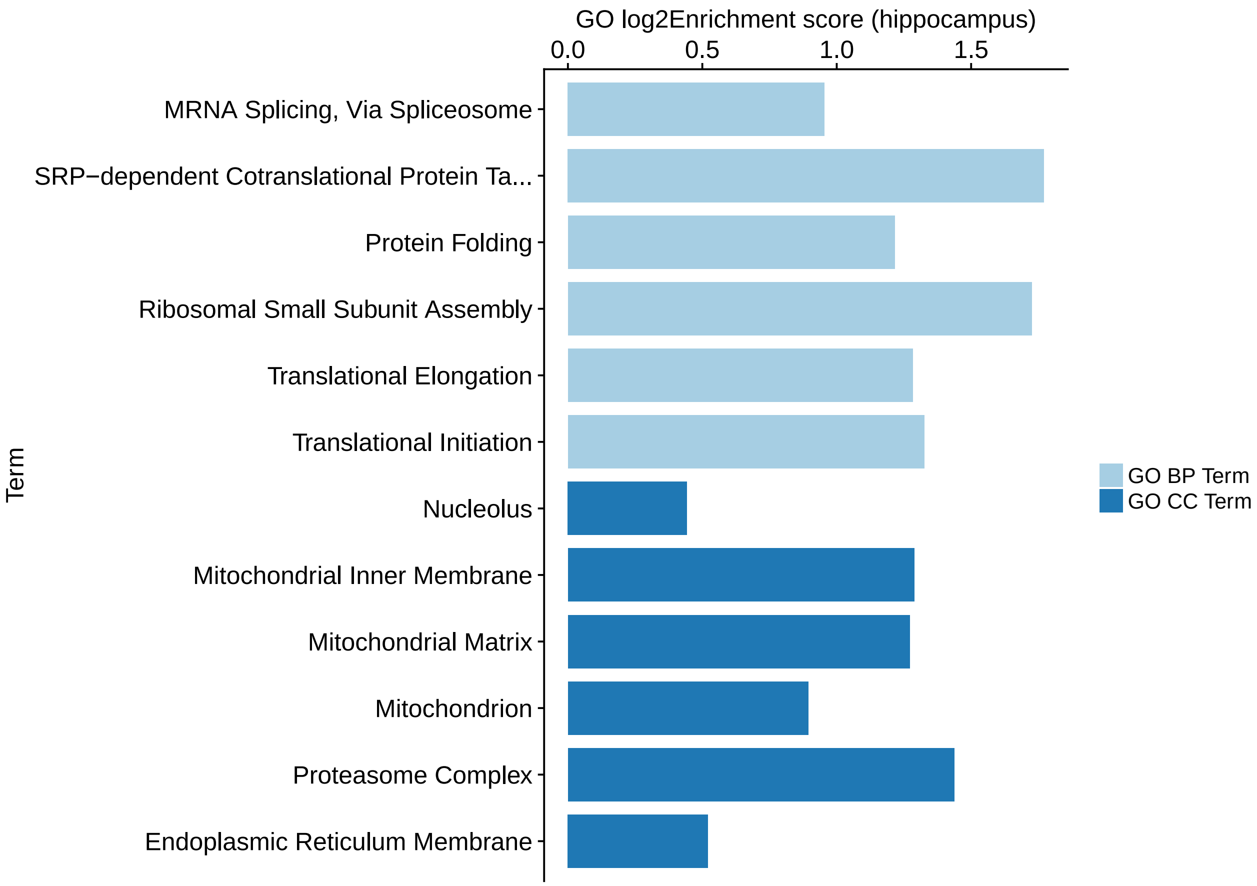


```
# ggsave("~/Project1/manuscript_GSEA/results/FigureS7A_Proteostasis_Enrichment_Human_Hippo.pdf", width


## dietary restriction
ggplot(human.dr.proteostasis, aes(Term, LogEnrich, fill = factor(Category))) +
                      scale_fill_brewer("", palette="Paired") +
                      geom_bar(position= "identity", stat="identity", width=0.8) +
                      coord_flip() +
                      scale_y_continuous(position = "right") +
                      labs( y = "GO log2Enrichment score (caloric restriction)") +
                      theme(axis.text.x = element_text(size=14),
                            axis.text.y = element_text(size=14),
                            text = element_text(size=14))
```

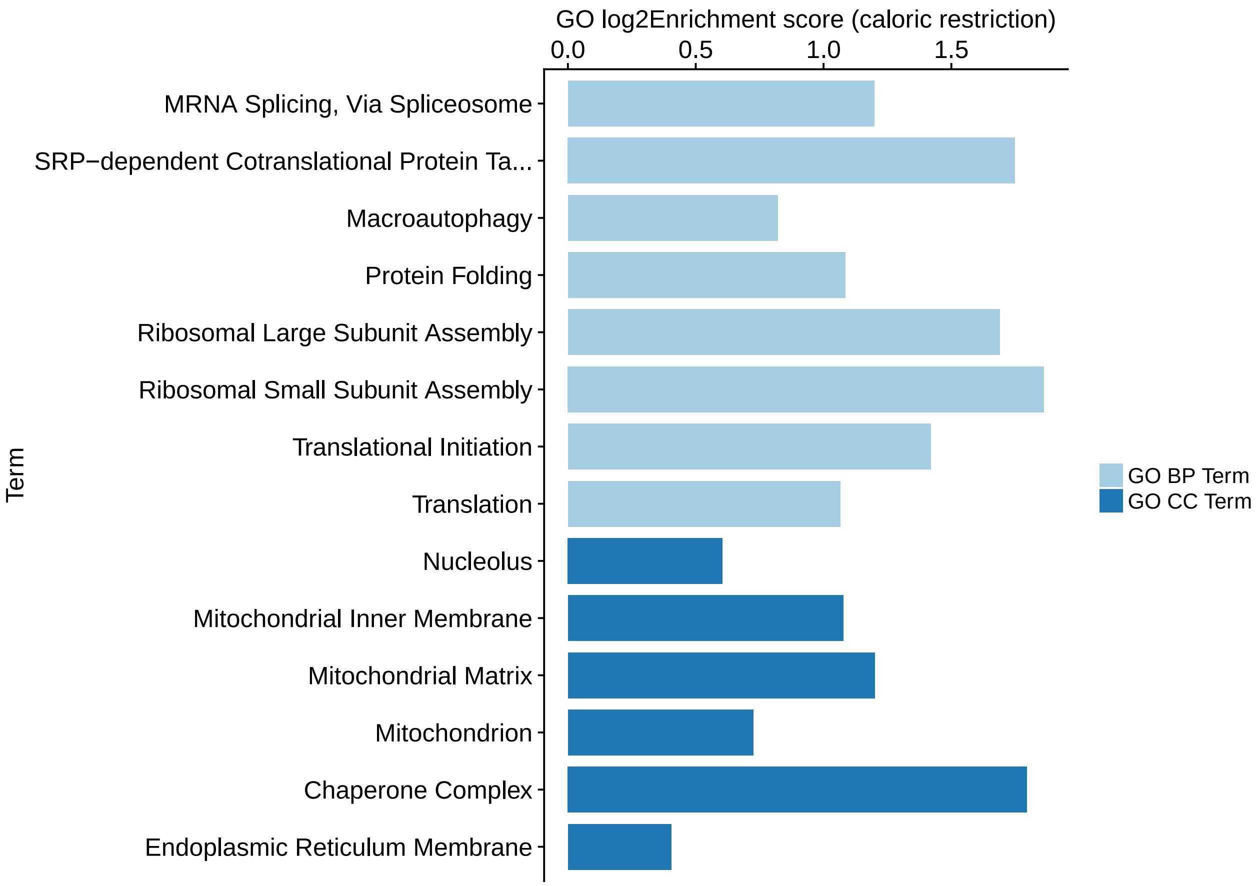


```
# ggsave("~/Project1/manuscript_GSEA/results/FigureS7B_Proteostasis_Enrichment_Human_DR.pdf", width =
```

Generating Supplementary Table S7.

```
muscle.proteasome <- human.muscle.cc.deg[[protein.quality.control.network[1]]]
muscle.translation <- human.muscle.bp.deg[[protein.quality.control.network[2]]]
muscle.macroautophagy <- human.muscle.bp.deg[[protein.quality.control.network[3]]]


## muscle
muscle.significant.proteasome <-
muscle.macroautophagy[muscle.macroautophagy$Significant == "FDR < 0.05", ]
muscle.significant.translation <- muscle.translation[muscle.translation$Significant == "FDR < 0.05", ]
muscle.significant.macroautophagy <- muscle.proteasome[muscle.proteasome$Significant == "FDR < 0.05", ]


# hippocampuss
hippo.proteasome <- human.hippo.cc.deg[[protein.quality.control.network[1]]]
hippo.translation <- human.hippo.bp.deg[[protein.quality.control.network[2]]]
hippo.macroautophagy <- human.hippo.bp.deg[[protein.quality.control.network[3]]]


hippo.significant.proteasome <- hippo.macroautophagy[hippo.macroautophagy$Significant == "FDR < 0.05",
hippo.significant.translation <- hippo.translation[hippo.translation$Significant == "FDR < 0.05", ]
hippo.significant.macroautophagy <- hippo.proteasome[hippo.proteasome$Significant == "FDR < 0.05", ]
```

```
# saving the evolutionary conserved genes
df.protein.quality.network <- list(Muscle_Human_Proteasome = muscle.significant.proteasome,
                                                            Muscle_Human_Translation = muscle.signific
                                                            Muscle_Human_Macroautophagy = muscle.signi

                                                            Hippo_Human_Proteasome = hippo.significant
                                                            Hippo_Human_Translation = hippo.significan
                                                            Hippo_Human_Macroautophagy = hippo.signifi


# WriteXLS(df.protein.quality.network,
#    ExcelFileName = "~/Project1/manuscript_GSEA/supplementary_data/Table_S7.xlsx", SheetNames = NULL)
```

Randomizations that are related to to observe proteostasis-linked process enrichment are performed on the cluster and saved in the script `randomized_processes_vitalit.R`.

```
setwd("/Users/akomljen/Project1/manuscript_GSEA/results/")
random.processes.muscle <- readRDS("list_of_random_processes_muscle.rds")
random.processes.hippo <- readRDS("list_of_random_processes_hippo.rds")

bp.proteostasis <- c(   "GO:0006412",     # translation
                        "GO:0006413",  # translational initiation
                        "GO:0006414",  # translational elongation
                        "GO:0006415",  # translational termination - didn't exist
                        "GO:0000028",   # ribosomal small subunit assembly
                        "GO:0000027",   # ribosomal large subunit assembly

                        "GO:0061684",  # chaperone-mediated autophagy
                        "GO:0006457",  # protein folding
                        "GO:0016236",  # Macroautophagy
                        "GO:0006614",  # SRP-dependent cotranslational protein targeting to membrane
                        "GO:0000398",  # mRNA splicing, via splicesome
                        "GO:0006508",  # proteolysis
                        "GO:0043248",  # proteasome assembly
                        "GO:0051290",  # protein heterotetramerization
                        "GO:0051289",  # protein homotetramerization
                        "GO:0006497",  # protein lipidation
                        "GO:0000209"   # protein polyubiquitination
                        )




list.random.proteostasis.muscle <-
  lapply(random.processes.muscle, function(x) x[na.omit(match(bp.proteostasis, x$GO.ID)),])

list.random.proteostasis.hippo <-
  lapply(random.processes.hippo, function(x) x[na.omit(match(bp.proteostasis, x$GO.ID)),])

summary.muscle.processes <- as.matrix(sapply(list.random.proteostasis.muscle, nrow))
summary.hippo.processes <- as.matrix(sapply(list.random.proteostasis.hippo, nrow))
```

```r
# go enrichment per species
muscle.common.eogs.oma <-
  integration.pvalues.across.species(human.oma.genes, deg.muscle.human.aging,
                                     mouse.oma.genes, deg.muscle.mouse.aging,
                                     fly.oma.genes, deg.wholebody.fly.aging,
                                     worm.oma.genes, deg.wholebody.worm.aging)
```

```
## Number of selected OGs in Human: 3069
## Number of selected OGs in Mouse: 2836
## Number of selected OGs in Fly: 3036
## Number of selected OGs in Worm: 3160
## Number of selected OGs common to species: 2511
```

```r
hippocampus.common.eogs.oma <-
  integration.pvalues.across.species(human.oma.genes, deg.hippo.human.aging,
                                      mouse.oma.genes, deg.hippo.mouse.aging,
                                      fly.oma.genes, deg.wholebody.fly.aging,
                                      worm.oma.genes, deg.wholebody.worm.aging)
```

```
## Number of selected OGs in Human: 3096
## Number of selected OGs in Mouse: 3136
## Number of selected OGs in Fly: 3036
## Number of selected OGs in Worm: 3160
## Number of selected OGs common to species: 2800
```

```r
fisher.muscle.human.oma <- muscle.common.eogs.oma$fisher.pvals.human
human.go.muscle.oma <-  go.tissue.species.enrichment(fisher.muscle.human.oma,
                                                     deg.muscle.human.aging, "org.Hs.eg.db")
```

```
##
## Building most specific GOs .....

##  ( 10856 GO terms found. )

##
## Build GO DAG topology .........

##  ( 14910 GO terms and 34803 relations. )

##
## Annotating nodes ...............

##  ( 12452 genes annotated to the GO terms. )

##
##          -- Elim Algorithm --
##
##      the algorithm is scoring 8802 nontrivial nodes
##      parameters:
##          test statistic: fisher
##          cutOff: 0.01

##
##   Level 20:  1 nodes to be scored    (0 eliminated genes)

##
##   Level 19:  2 nodes to be scored    (0 eliminated genes)

##
##   Level 18:  6 nodes to be scored    (0 eliminated genes)
```

```
##
##   Level 17:  14 nodes to be scored   (0 eliminated genes)
##
##   Level 16:  40 nodes to be scored   (0 eliminated genes)
##
##   Level 15:  100 nodes to be scored  (21 eliminated genes)
##
##   Level 14:  198 nodes to be scored  (156 eliminated genes)
##
##   Level 13:  362 nodes to be scored  (234 eliminated genes)
##
##   Level 12:  597 nodes to be scored  (538 eliminated genes)
##
##   Level 11:  858 nodes to be scored  (1140 eliminated genes)
##
##   Level 10:  1088 nodes to be scored (2136 eliminated genes)
##
##   Level 9:   1209 nodes to be scored (3104 eliminated genes)
##
##   Level 8:   1217 nodes to be scored (4021 eliminated genes)
##
##   Level 7:   1181 nodes to be scored (4628 eliminated genes)
##
##   Level 6:   934 nodes to be scored  (5488 eliminated genes)
##
##   Level 5:   560 nodes to be scored  (5737 eliminated genes)
##
##   Level 4:   295 nodes to be scored  (6623 eliminated genes)
##
##   Level 3:   116 nodes to be scored  (7162 eliminated genes)
##
##   Level 2:   23 nodes to be scored   (7587 eliminated genes)
##
##   Level 1:   1 nodes to be scored    (7587 eliminated genes)
##
## Building most specific GOs .....
##  ( 3774 GO terms found. )
##
## Build GO DAG topology .........
##  ( 4286 GO terms and 5520 relations. )
##
## Annotating nodes ..............
##  ( 12505 genes annotated to the GO terms. )
```

```
##
##           -- Elim Algorithm --
##
##       the algorithm is scoring 2380 nontrivial nodes
##       parameters:
##           test statistic: fisher
##           cutOff: 0.01
##
##   Level 15:  3 nodes to be scored    (0 eliminated genes)
##
##   Level 14:  6 nodes to be scored    (0 eliminated genes)
##
##   Level 13:  11 nodes to be scored   (15 eliminated genes)
##
##   Level 12:  37 nodes to be scored   (15 eliminated genes)
##
##   Level 11:  47 nodes to be scored   (15 eliminated genes)
##
##   Level 10:  78 nodes to be scored   (177 eliminated genes)
##
##   Level 9:   147 nodes to be scored  (196 eliminated genes)
##
##   Level 8:   252 nodes to be scored  (1636 eliminated genes)
##
##   Level 7:   411 nodes to be scored  (1686 eliminated genes)
##
##   Level 6:   625 nodes to be scored  (1813 eliminated genes)
##
##   Level 5:   452 nodes to be scored  (2195 eliminated genes)
##
##   Level 4:   228 nodes to be scored  (3597 eliminated genes)
##
##   Level 3:   69 nodes to be scored   (4792 eliminated genes)
##
##   Level 2:   13 nodes to be scored   (4956 eliminated genes)
##
##   Level 1:   1 nodes to be scored    (6946 eliminated genes)
##
## Building most specific GOs .....
##  ( 1553 GO terms found. )
##
## Build GO DAG topology .........
##  ( 1850 GO terms and 3486 relations. )
```

```
##
## Annotating nodes ..............
##  ( 13030 genes annotated to the GO terms. )
##
##           -- Elim Algorithm --
##
##       the algorithm is scoring 1210 nontrivial nodes
##       parameters:
##           test statistic: fisher
##           cutOff: 0.01
##
##   Level 17:  1 nodes to be scored    (0 eliminated genes)
##
##   Level 16:  9 nodes to be scored    (0 eliminated genes)
##
##   Level 15:  31 nodes to be scored   (9 eliminated genes)
##
##   Level 14:  60 nodes to be scored   (44 eliminated genes)
##
##   Level 13:  77 nodes to be scored   (487 eliminated genes)
##
##   Level 12:  129 nodes to be scored  (730 eliminated genes)
##
##   Level 11:  142 nodes to be scored  (1713 eliminated genes)
##
##   Level 10:  150 nodes to be scored  (5537 eliminated genes)
##
##   Level 9:   101 nodes to be scored  (5856 eliminated genes)
##
##   Level 8:   139 nodes to be scored  (5919 eliminated genes)
##
##   Level 7:   77 nodes to be scored   (8263 eliminated genes)
##
##   Level 6:   99 nodes to be scored   (9176 eliminated genes)
##
##   Level 5:   60 nodes to be scored   (10721 eliminated genes)
##
##   Level 4:   86 nodes to be scored   (10721 eliminated genes)
##
##   Level 3:   37 nodes to be scored   (10731 eliminated genes)
##
##   Level 2:   11 nodes to be scored   (10859 eliminated genes)
##
##   Level 1:   1 nodes to be scored    (10859 eliminated genes)
```

```
## Warning in p.adjust(as.numeric(res$elimFisher), method = "fdr"): NAs
## introduced by coercion
```

```
fisher.hippo.human.oma <- hippocampus.common.eogs.oma$fisher.pvals.human
human.go.hippo.oma <-  go.tissue.species.enrichment(fisher.hippo.human.oma,
                                                    deg.hippo.human.aging, "org.Hs.eg.db")
```

```
##
## Building most specific GOs .....

##  ( 11050 GO terms found. )

##
## Build GO DAG topology ..........

##  ( 15076 GO terms and 35165 relations. )

##
## Annotating nodes ...............

##  ( 13193 genes annotated to the GO terms. )

##
##          -- Elim Algorithm --
##
##      the algorithm is scoring 8638 nontrivial nodes
##      parameters:
##          test statistic: fisher
##          cutOff: 0.01

##
##   Level 19:  1 nodes to be scored    (0 eliminated genes)

##
##   Level 18:  3 nodes to be scored    (0 eliminated genes)

##
##   Level 17:  9 nodes to be scored    (0 eliminated genes)

##
##   Level 16:  32 nodes to be scored   (0 eliminated genes)

##
##   Level 15:  93 nodes to be scored   (22 eliminated genes)

##
##   Level 14:  194 nodes to be scored  (152 eliminated genes)

##
##   Level 13:  353 nodes to be scored  (250 eliminated genes)

##
##   Level 12:  571 nodes to be scored  (608 eliminated genes)

##
##   Level 11:  845 nodes to be scored  (1064 eliminated genes)

##
##   Level 10:  1071 nodes to be scored (1559 eliminated genes)

##
##   Level 9:   1194 nodes to be scored (2621 eliminated genes)
```

```
##
##   Level 8:   1207 nodes to be scored (3745 eliminated genes)
##
##   Level 7:   1168 nodes to be scored (4837 eliminated genes)
##
##   Level 6:   914 nodes to be scored  (5470 eliminated genes)
##
##   Level 5:   559 nodes to be scored  (5786 eliminated genes)
##
##   Level 4:   283 nodes to be scored  (7168 eliminated genes)
##
##   Level 3:   116 nodes to be scored  (7513 eliminated genes)
##
##   Level 2:   24 nodes to be scored   (7746 eliminated genes)
##
##   Level 1:   1 nodes to be scored    (7746 eliminated genes)
##
## Building most specific GOs .....
##  ( 3838 GO terms found. )
##
## Build GO DAG topology ..........
##  ( 4345 GO terms and 5602 relations. )
##
## Annotating nodes ...............
##  ( 13267 genes annotated to the GO terms. )
##
##           -- Elim Algorithm --
##
##       the algorithm is scoring 2425 nontrivial nodes
##       parameters:
##           test statistic: fisher
##           cutOff: 0.01
##
##   Level 16:  1 nodes to be scored    (0 eliminated genes)
##
##   Level 15:  3 nodes to be scored    (0 eliminated genes)
##
##   Level 14:  6 nodes to be scored    (0 eliminated genes)
##
##   Level 13:  13 nodes to be scored   (0 eliminated genes)
##
##   Level 12:  37 nodes to be scored   (57 eliminated genes)
##
##   Level 11:  49 nodes to be scored   (57 eliminated genes)
```

```
##
##   Level 10:  79 nodes to be scored   (147 eliminated genes)
##
##   Level 9:   161 nodes to be scored  (147 eliminated genes)
##
##   Level 8:   264 nodes to be scored  (1803 eliminated genes)
##
##   Level 7:   428 nodes to be scored  (1841 eliminated genes)
##
##   Level 6:   631 nodes to be scored  (2200 eliminated genes)
##
##   Level 5:   437 nodes to be scored  (2682 eliminated genes)
##
##   Level 4:   233 nodes to be scored  (4245 eliminated genes)
##
##   Level 3:   69 nodes to be scored   (5969 eliminated genes)
##
##   Level 2:   13 nodes to be scored   (7191 eliminated genes)
##
##   Level 1:   1 nodes to be scored    (8015 eliminated genes)
##
## Building most specific GOs .....
##  ( 1569 GO terms found. )
##
## Build GO DAG topology .........
##  ( 1858 GO terms and 3508 relations. )
##
## Annotating nodes ...............
##  ( 13876 genes annotated to the GO terms. )
##
##           -- Elim Algorithm --
##
##       the algorithm is scoring 1239 nontrivial nodes
##       parameters:
##           test statistic: fisher
##           cutOff: 0.01
##
##   Level 17:  1 nodes to be scored    (0 eliminated genes)
##
##   Level 16:  11 nodes to be scored   (0 eliminated genes)
##
##   Level 15:  30 nodes to be scored   (9 eliminated genes)
##
##   Level 14:  62 nodes to be scored   (40 eliminated genes)
```

```
##
##   Level 13:  81 nodes to be scored   (119 eliminated genes)

##
##   Level 12:  136 nodes to be scored  (390 eliminated genes)

##
##   Level 11:  154 nodes to be scored  (1436 eliminated genes)

##
##   Level 10:  154 nodes to be scored  (5811 eliminated genes)

##
##   Level 9:   107 nodes to be scored  (6119 eliminated genes)

##
##   Level 8:   137 nodes to be scored  (6125 eliminated genes)

##
##   Level 7:   78 nodes to be scored   (8517 eliminated genes)

##
##   Level 6:   101 nodes to be scored  (9445 eliminated genes)

##
##   Level 5:   55 nodes to be scored   (11174 eliminated genes)

##
##   Level 4:   85 nodes to be scored   (11176 eliminated genes)

##
##   Level 3:   35 nodes to be scored   (11188 eliminated genes)

##
##   Level 2:   11 nodes to be scored   (11325 eliminated genes)

##
##   Level 1:   1 nodes to be scored    (11325 eliminated genes)
## Warning in p.adjust(as.numeric(res$elimFisher), method = "fdr"): NAs
## introduced by coercion
```

```
human.go.muscle <- human.go.muscle.oma[[1]][na.omit(match(bp.proteostasis,
                                                          human.go.muscle.oma[[1]]$GO.ID)),]
human.go.hippo <- human.go.hippo.oma[[1]][na.omit(match(bp.proteostasis,
                                                        human.go.hippo.oma[[1]]$GO.ID)),]

obs.no.proteoprocess.muscle <- nrow(human.go.muscle)
obs.no.proteoprocess.hippo <- nrow(human.go.hippo)


# pdf("~/Project1/manuscript_GSEA/results/Supplementary_Fig_S6.pdf", 7,7)

par(mfrow = c(2,1))
# for number of connections
hist(as.numeric(summary.muscle.processes[,1]), xlim = c(0, 10),
xlab = "Number of proteostasis-linked processes\n(skeletal muscle)", main = "Skeletal muscle GO process

abline(v = obs.no.proteoprocess.muscle, col = "red", lwd = 2)
legend(4, 60, "Observed", lwd=2.5, col= "red")
```

```r
hist(as.numeric(summary.hippo.processes[,1]), xlim = c(0, 10),
xlab = "Number of proteostasis-linked processes\n(hippocampus)", main = "Hippocampus GO processes")

abline(v = obs.no.proteoprocess.hippo, col = "red", lwd = 2)
```

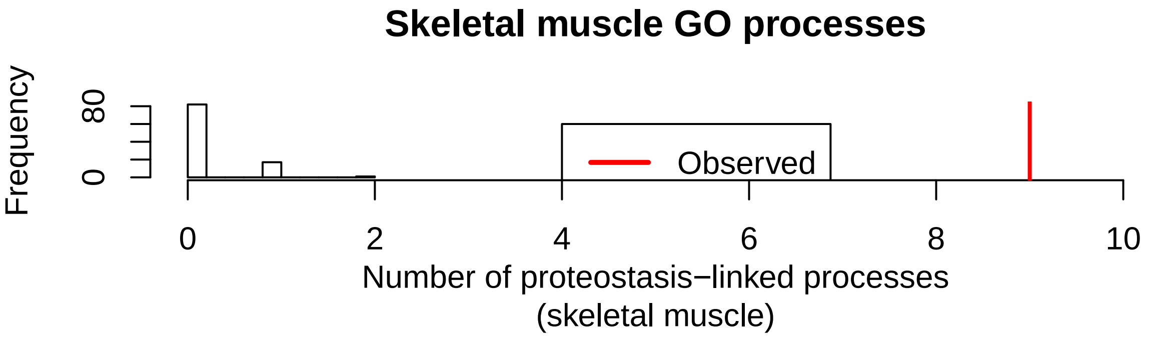


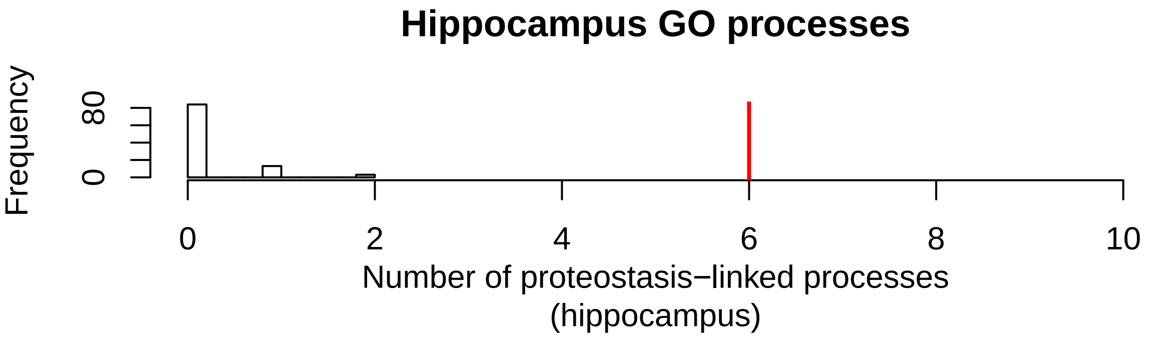


```r
# dev.off()
```

Saving expression matrices for Networks part.

```r
# --------------------------------------------
## saving the expression matrices for part III (network)

extract.eogs.per.species <- function(tissue.ogs, expression.matrix){

   tissue.ogs$FDR <- p.adjust(tissue.ogs$combn.P.Val, method = "fdr")
   tissue.ogs.sig <- tissue.ogs[which(tissue.ogs$FDR < 0.10), ]
   cat("Number of significant genes in orthogroup:", print(nrow(tissue.ogs.sig)), "\n")
   expression.matrix.oma <- expression.matrix[na.omit(match(rownames(tissue.ogs.sig),
                                                       rownames(expression.matrix))),]
   cat("Number of the rows in the expression matrix after filtering for OMA groups:",
       print(nrow(expression.matrix.oma)), "\n")
   return(expression.matrix.oma)
   # go.tissue.enrichment <- go.enrichment(rownames(toptable.tissue), rownames(tissue.ogs.sig), genome)
   # return(go.tissue.enrichment)

 }


# ------------------------
# Preparing data for Network part.
```

```
# Load expression matrices

# path <- "~/Project1/manuscript_GSEA/data_preprocessing/"

# human
# expression.human.muscle <-
# readRDS(paste0(path, "hsapiens/aging_data/exp_mat/voom_Muscle_Skeletal_GTEx_V6p.rds"))
# expression.human.hippo <-
# readRDS(paste0(path, "hsapiens/aging_data/exp_mat/voom_Brain_Hippocampus_GTEx_V6p.rds"))

# mouse
# expression.mouse.muscle <-
# readRDS(paste0(path, "mmusculus/aging_data/skeletal_muscle/expression_matrix_mouse_skeletal_muscle_ag
# expression.mouse.hippo <-
# readRDS(paste0(path, "mmusculus/aging_data/hippocampus/voom_mouse_expression_matrix_hippocampus_aging

# fly
# expression.fly.wb <-
# readRDS(paste0(path, "dmelanogaster/aging_data/expression_matrix_fly_aging.rds"))
# expression.fly.wholebody <- expression.fly.wb[[2]]

# worm
# expression.worm.wholebody <-
# readRDS(paste0(path, "celegans/aging_data/voom_celegans_expression_matrix_aging.rds"))


# --------------------
# filtering

# exp.matrix.human.muscle.oma <-
# extract.eogs.per.species(muscle.common.eogs.oma$fisher.pvals.human,
#                                 expression.human.muscle)
# exp.matrix.mouse.muscle.oma <-
# extract.eogs.per.species(muscle.common.eogs.oma$fisher.pvals.mouse,
#                           expression.mouse.muscle)
# exp.matrix.fly.muscle.oma <-
# extract.eogs.per.species(muscle.common.eogs.oma$fisher.pvals.dmelano,
#                           expression.fly.wholebody )
# exp.matrix.worm.muscle.oma <-
# extract.eogs.per.species(muscle.common.eogs.oma$fisher.pvals.cele,
#                           expression.worm.wb)


# exp.matrix.human.hippo.oma <-
# extract.eogs.per.species(hippocampus.common.eogs.oma$fisher.pvals.human,
#                          expression.human.hippo)
# exp.matrix.mouse.hippo.oma <-
# extract.eogs.per.species(hippocampus.common.eogs.oma$fisher.pvals.mouse,
#                           expression.mouse.hippo)
# exp.matrix.fly.hippo.oma <-
# extract.eogs.per.species(hippocampus.common.eogs.oma$fisher.pvals.dmelano,
#                           expression.fly.wholebody )
# exp.matrix.worm.hippo.oma <-
```

```
# extract.eogs.per.species(hippocampus.common.eogs.oma$fisher.pvals.cele,
#                           expression.worm.wb)

# it could have been saved in a list, but here is separate now
# saveRDS(exp.matrix.human.muscle.oma,
# file = paste0(path, "/expressionmat_for_networks/expmat_hsapiens_muscle_oma_sig.rds"))
# saveRDS(exp.matrix.mouse.muscle.oma,
# file = paste0(path, "/expressionmat_for_networks/expmat_mmusculus_muscle_oma_sig.rds"))
# saveRDS(exp.matrix.fly.muscle.oma,
# file = paste0(path, "/expressionmat_for_networks/expmat_dmelanogaster_muscle_oma_sig.rds"))
# saveRDS(exp.matrix.worm.muscle.oma,
# file = paste0(path, "/expressionmat_for_networks/expmat_celegans_muscle_oma_sig.rds"))


# saveRDS(exp.matrix.human.hippo.oma,
# file = paste0(path, "/expressionmat_for_networks/expmat_hsapiens_hippo_oma_sig.rds"))
# saveRDS(exp.matrix.mouse.hippo.oma,
# file = paste0(path, "/expressionmat_for_networks/expmat_mmusculus_hippo_oma_sig.rds"))
# saveRDS(exp.matrix.fly.hippo.oma,
# file = paste0(path, "/expressionmat_for_networks/expmat_dmelanogaster_hippo_oma_sig.rds"))
# saveRDS(exp.matrix.worm.hippo.oma,
# file = paste0(path, "/expressionmat_for_networks/expmat_celegans_hippo_oma_sig.rds"))
```

### Main Figures + Supplement (Network part)

*Andrea Komljenović*

*3/8/2018*

**Step III**

Loading the packages.

Functions for the networks and module search.

```
## ----------------
# FUNCTIONS for constructing networks
## ----------------


### CORRELATION / Co-expression MATRIX
significant.edges <- function(expression.matrix){
    require(Hmisc)
  siglevel <- 0.05
    corrmatr <- rcorr(t(expression.matrix),type="pearson")
    corrmat <- corrmatr$r
    diag(corrmat) <- 0
    corrmat[corrmatr$P > siglevel] <- 0
    corrmat[corrmat < 0 ] <- 0
    # flatten correlation matrix for plotting
    corrmat.flat <- flattenCorrMatrix(corrmat)
  return( corrmat.flat)

}

flattenCorrMatrix <- function(cormat) {
  ut <- upper.tri(cormat, diag = FALSE)
  return(data.frame(
    row = rownames(cormat)[row(cormat)[ut]],
    column = rownames(cormat)[col(cormat)[ut]],
    cor  =(cormat)[ut]
    ))
}



preprocess.edges <- function(expression.matrix, annotation.species, annotation){

    mm <- match(rownames(expression.matrix), annotation.species)
    annot <- annotation[mm,]

    rownames(expression.matrix) <- rownames(annot)
    corrmat <- significant.edges(expression.matrix)
    return(corrmat)
}
```

```r
# data integration of co-expression networks per species / per tissue
data.integration <- function(coexp.sp1, coexp.sp2, coexp.sp3, coexp.sp4, n.genes.tissue){
  # require(RobustRankAggreg)
  correlations <- list(human = coexp.sp1, mouse = coexp.sp2, fly = coexp.sp3, worm = coexp.sp4)
  cat("Number of gene pairs:", sapply(correlations, nrow), "\n")

  coexpr.mat <- lapply(correlations, function(corrs) {
                              rownames(corrs) <- paste(corrs$row, corrs$column , sep = ":")
                              return(corrs)
                      })
  # order per species
  sorted.coexpr.mat <- lapply(coexpr.mat, function(x) {
                            x <- x[order(x$cor, na.last = TRUE, decreasing = TRUE), ]
                            return(x) })
  sorted.correlations.rownames <- lapply(sorted.coexpr.mat, function(s) rownames(s))
  rank.matrix <- rankMatrix(sorted.correlations.rownames, N = n.genes.tissue, full = TRUE)
  aggregated.ranks <- aggregateRanks(rmat = rank.matrix, N = n.genes.tissue, method = "RRA")
  return(list(aggregation = aggregated.ranks, ranks.per.species = rank.matrix))
}


clean.modules <- function(subgraph.module, annotation.oma, process.human.symbol){
  tissue.community.module <- subgraph.module
  V(tissue.community.module)$size <- 3
  V(tissue.community.module)$frame.color <- "white"

  ### remap OGs to gene names
  remap.tissue.module <-
                  annotation.oma$human.gene.name[match(V(tissue.community.module)$name,
                  rownames(annotation.oma))]
  names(remap.tissue.module) <- mapIds(org.Hs.eg.db,
                                        keys=as.vector(remap.tissue.module),
                                        column="SYMBOL",
                                        keytype="ENSEMBL",
                                        multiVals="first")


  genes.process <- process.human.symbol[na.omit(match(names(remap.tissue.module),
                                                  process.human.symbol))]

  V(tissue.community.module)$label <- names(remap.tissue.module)
  hub.score.module.tissue <- hub.score(tissue.community.module)$vector
  ordered.hubs.module.tissue <- hub.score.module.tissue[order(hub.score.module.tissue,
                                                        decreasing = TRUE)][1:3]
  matched.hubs.tissue <- annotation.oma$human.gene.name[match(names(ordered.hubs.module.tissue),
                                                        rownames(annotation.oma))]

  names(matched.hubs.tissue) <- mapIds(org.Hs.eg.db,
                                      keys = as.vector(matched.hubs.tissue),
                                      column = "SYMBOL",
                                      keytype = "ENSEMBL",
                                      multiVals = "first")
```

```r
  return(list(module = tissue.community.module, hubs = matched.hubs.tissue, process = genes.process))

}



## plotting the modules
plot.modules <- function(genes.of.interest, community.module.of.interest,
                         neighbourhood.nodes, hubs.genes, plot.title){

  idx <- na.omit(match(genes.of.interest, V(community.module.of.interest)$label))
  V(community.module.of.interest)$label[idx] <- genes.of.interest
  V(community.module.of.interest)$label[-idx] <- ""
  # neighbouring genes
  V(community.module.of.interest)$color <- "grey40"
  V(community.module.of.interest)$color[match(neighbourhood.nodes,
                                              V(community.module.of.interest)$label)] <- "royalblue"
  # hub genes
  V(community.module.of.interest)$color[match(names(hubs.genes),
                                              V(community.module.of.interest)$label)] <- "firebrick3"

  tissue.layout <- layout.kamada.kawai(community.module.of.interest)
  plot(community.module.of.interest,
         layout = tissue.layout,
         main = plot.title,
         vertex.label.color = "black",
         vertex.label.cex = 0.6,
         vertex.size = 2.5,
         vertex.label.dist = 0.4,

         vertex.label.family = "Helvetica")

         legend("topright", c("hub gene", "neighbour gene"), pch = 16,
       col = c("firebrick3", "royalblue") )


}


checking.hubs <- function(module, ensembl.genes.protein.coding, annotation.oma){

  tissue.community.module <- module

  remap.tissue.module <-
              annotation.oma$human.gene.name[match(V(tissue.community.module)$name,
              rownames(annotation.oma))]

  gene.name.module <-
    ensembl.genes.protein.coding[match(remap.tissue.module,
            ensembl.genes.protein.coding$ensembl_gene_id),]
  V(tissue.community.module)$label <- gene.name.module$external_gene_name
  V(tissue.community.module)$name <- gene.name.module$external_gene_name
```

```
  hub.score.module <- hub.score(tissue.community.module)$vector
  ordered.hubs.module <- hub.score.module[order(hub.score.module, decreasing = TRUE)][1:20]
  ensembl.module.hubs <-
    ensembl.genes.protein.coding[match(names(ordered.hubs.module [1:20]),
    ensembl.genes.protein.coding$external_gene_name),]
  return(ensembl.module.hubs)

}
```

Loading the expression matrices for networks and annotation of the orthogroups.

```
path.oma.exp <- "~/Project1/manuscript_GSEA/data_preprocessing/expressionmat_for_networks/"

# ------------------------
# skeletal muscle
# human
expMatHuman.OG.muscle.oma.sig <- readRDS(paste0(path.oma.exp, "expmat_hsapiens_muscle_oma_sig.rds"))

# mouse
expMatMouse.OG.muscle.oma.sig <- readRDS(paste0(path.oma.exp,"expmat_mmusculus_muscle_oma_sig.rds"))
## add sample names
colnames(expMatMouse.OG.muscle.oma.sig) <- c(rep("young", 4), rep("old",5))

# fly
expMatFly.OG.muscle.oma.sig <- readRDS(paste0(path.oma.exp, "expmat_dmelanogaster_muscle_oma_sig.rds"))

# worm
expMatWorm.OG.muscle.oma.sig <- readRDS(paste0(path.oma.exp, "expmat_celegans_muscle_oma_sig.rds"))
# remove too young samples
expMatWorm.OG.muscle.oma.sig <- expMatWorm.OG.muscle.oma.sig[, -c(1:3)]


# -----------------------
# hippocampus significant
# human
expMatHuman.OG.hippo.oma.sig <- readRDS(paste0(path.oma.exp, "expmat_hsapiens_hippo_oma_sig.rds"))

# mouse
expMatMouse.OG.hippo.oma.sig <- readRDS(paste0(path.oma.exp,"expmat_mmusculus_hippo_oma_sig.rds"))

# fly
expMatFly.OG.hippo.oma.sig <- readRDS(paste0(path.oma.exp, "expmat_dmelanogaster_hippo_oma_sig.rds"))

# worm
expMatWorm.OG.hippo.oma.sig <- readRDS(paste0(path.oma.exp, "expmat_celegans_hippo_oma_sig.rds"))
# remove too young samples
expMatWorm.OG.hippo.oma.sig <- expMatWorm.OG.hippo.oma.sig[, -c(1:3)]



# ---------------------
# annotation
annotation.muscle.oma <- readRDS("~/Project1/manuscript_GSEA/results/annotation_muscle_oma.rds")
annotation.hippo.oma <- readRDS("~/Project1/manuscript_GSEA/results/annotation_hippocampus_oma.rds")
```

```r
# skeletal muscle
human.corr.muscle.oma.sig <- preprocess.edges(expMatHuman.OG.muscle.oma.sig,
                                              annotation.muscle.oma$human.gene.name,
                                              annotation.muscle.oma)
mouse.corr.muscle.oma.sig <- preprocess.edges(expMatMouse.OG.muscle.oma.sig,
                                              annotation.muscle.oma$mouse.gene.name,
                                              annotation.muscle.oma)
fly.corr.muscle.oma.sig <- preprocess.edges(expMatFly.OG.muscle.oma.sig,
                                            annotation.muscle.oma$fly.gene.name,
                                            annotation.muscle.oma)
worm.corr.muscle.oma.sig <- preprocess.edges(expMatWorm.OG.muscle.oma.sig,
                                             annotation.muscle.oma$worm.gene.name,
                                             annotation.muscle.oma)

# hippocampus
human.corr.hippo.oma.sig <- preprocess.edges(expMatHuman.OG.hippo.oma.sig,
                                             annotation.hippo.oma$human.gene.name,
                                             annotation.hippo.oma)
mouse.corr.hippo.oma.sig <- preprocess.edges(expMatMouse.OG.hippo.oma.sig,
                                             annotation.hippo.oma$mouse.gene.name,
                                             annotation.hippo.oma)
fly.corr.hippo.oma.sig <- preprocess.edges(expMatFly.OG.hippo.oma.sig,
                                           annotation.hippo.oma$fly.gene.name,
                                           annotation.hippo.oma)
worm.corr.hippo.oma.sig <- preprocess.edges(expMatWorm.OG.hippo.oma.sig,
                                            annotation.hippo.oma$worm.gene.name,
                                            annotation.hippo.oma)


# number of the
n.muscle.eog.oma.sig <- nrow(expMatHuman.OG.muscle.oma.sig)
n.muscle.eog.oma.sig # 2010
```

```
## [1] 2010
```

```r
n.hippo.eog.oma.sig <- nrow(expMatHuman.OG.hippo.oma.sig)
n.hippo.eog.oma.sig # 2075
```

```
## [1] 2075
```

Real data network data integration.

```r
## integration of co-expression links across species

muscle.coexpres.oma.sig <- data.integration(human.corr.muscle.oma.sig, mouse.corr.muscle.oma.sig,
                                            fly.corr.muscle.oma.sig, worm.corr.muscle.oma.sig,
                                            n.muscle.eog.oma.sig)
```

```
## Number of gene pairs: 2019045 2019045 2019045 2019045
```

```r
#  (2019045 links - pearson correlation method)
hippo.coexpres.oma.sig <- data.integration(human.corr.hippo.oma.sig, mouse.corr.hippo.oma.sig,
                                           fly.corr.hippo.oma.sig, worm.corr.hippo.oma.sig,
                                           n.hippo.eog.oma.sig)
```

```
## Number of gene pairs: 2151775 2151775 2151775 2151775
```

```r
# (2151775 links - pearson correlation method)
```

Significance threshold defined for the co-expression links consideration.

```r
# defining the threshold for significant co-expression links
sig.level <- 0.001
# muscle
aggregated.signif.muscle.oma.sig <-
  muscle.coexpres.oma.sig$aggregation[which(muscle.coexpres.oma.sig$aggregation$Score <= sig.level), ]
nrow(aggregated.signif.muscle.oma.sig) #  2887 significant co-expression links
```

```
## [1] 2887
```

```r
# hippocampus
aggregated.signif.hippo.oma.sig <-
  hippo.coexpres.oma.sig$aggregation[which(hippo.coexpres.oma.sig$aggregation$Score <= sig.level), ]
nrow(aggregated.signif.hippo.oma.sig) # 3353 significant co-expression links
```

```
## [1] 3353
```

Finding the giant component of the network for further analysis.

```r
# defining the edges
edge.list.muscle.oma.sig <- do.call(rbind, strsplit(rownames(aggregated.signif.muscle.oma.sig), ':'))
edge.list.hippo.oma.sig <- do.call(rbind, strsplit(rownames(aggregated.signif.hippo.oma.sig), ':'))


# 1142 genes
net.muscle.oma.sig <- graph_from_data_frame(d = edge.list.muscle.oma.sig, directed = FALSE)
# 1098 genes
net.hippo.oma.sig <- graph_from_data_frame(d = edge.list.hippo.oma.sig, directed = FALSE)

## weighting the edges for multilevel community
E(net.muscle.oma.sig)$weight <- -log10(aggregated.signif.muscle.oma.sig$Score)
E(net.hippo.oma.sig)$weight <- -log10(aggregated.signif.hippo.oma.sig$Score)


# decompose graph in order to remove singletones
comps.muscle.sig <- decompose.graph(net.muscle.oma.sig)
table(sapply(comps.muscle.sig, vcount))
```

```
##
##    2    3    4 1050
##   32    8    1    1
```

```r
# giant component
# 2    3    4 1050
#  32    8    1    1

comps.hippo.sig <- decompose.graph(net.hippo.oma.sig)
table(sapply(comps.hippo.sig, vcount))
```

```
##
##    2    3 1067
##   14    1    1
```

```r
# giant compononet
# 2    3 1067
```

```
#    14     1     1
```

Performing the greedy search algorithm (multilevel) on the networks in order to find modules, then calculating the sizes of the modules.

```
### networks
net.muscle.gc.sig <- decompose.graph(net.muscle.oma.sig)[[1]]
net.hippo.gc.sig <- decompose.graph(net.hippo.oma.sig)[[1]]

### searcing for cross-species modules
ml.com.muscle.sig <- multilevel.community(net.muscle.gc.sig)
ml.com.hippo.sig <- multilevel.community(net.hippo.gc.sig)

# muscle
multilevel.sizesComm.muscle.sig <- sizes(ml.com.muscle.sig)
multilevel.numComm.muscle.sig <- length(multilevel.sizesComm.muscle.sig )


# hippocampus
multilevel.sizesComm.hippo.sig <- sizes(ml.com.hippo.sig)
multilevel.numComm.hippo.sig <- length(multilevel.sizesComm.hippo.sig )
```

For skeletal muscle, first detected 20 modules, filtered to 10 modules (focus on the modules with more than 10 genes).

```
## building the network
net.muscle.tryout <- net.muscle.gc.sig
V(net.muscle.tryout)$ModuleMemb <- ml.com.muscle.sig$membership
module.names <- sort(unique(V(net.muscle.tryout)$ModuleMemb))


# replacing group of vertices with single meta-vertices
net.muscle.coarsen <- igraph::contract.vertices(net.muscle.tryout,
                                                ml.com.muscle.sig$membership)
E(net.muscle.coarsen)$weight <- 1


# selecting the modules bigger than size of 10
subg.muscle.sig <- list()
for(g in unique(membership(ml.com.muscle.sig))){
      subg.muscle.sig[[g]] <-
        induced.subgraph(net.muscle.gc.sig,
                    which((membership(ml.com.muscle.sig)==g)
                      & ( sizes(ml.com.muscle.sig)[[g]] >= 10)))
}
# nameing the modules
V(net.muscle.coarsen)$name <- paste0("M", 1:20)

# defines which modules are empty
empty.module <- which(sapply(subg.muscle.sig, vcount) == 0)
## this removes those small modules
net.muscle.coarsen <- net.muscle.coarsen - c(paste0("M", empty.module))
net.muscle.coarsen <- igraph::simplify(net.muscle.coarsen)

### the size of the nodes
```

```
l.cicr <- layout_in_circle(net.muscle.coarsen)
sizes <- sapply(subg.muscle.sig, vcount)[-empty.module]
node.size <- as.vector(sizes)


# pdf("~/Project1/manuscript_GSEA/results/Figure4A_SkeletalMuscle_coarsen_net_2.pdf",
# useDingbats = FALSE)
plot(net.muscle.coarsen,
          vertex.size= sqrt(node.size)*2,
          vertex.label= paste0("M",1:12),
                    vertex.label.color="black",
          margin=.5,
          vertex.color = c("#268bd2", "white", "white", "white","white", "#268bd2",
                           "#268bd2", "#268bd2", "white", "white", "#268bd2", "#268bd2"),
                    edge.width=sqrt(E(net.muscle.coarsen)$weight),
                    edge.arrow.size=0, layout = l.cicr, vertex.label.dist = 1,
                    main = "Skeletal Muscle")
```

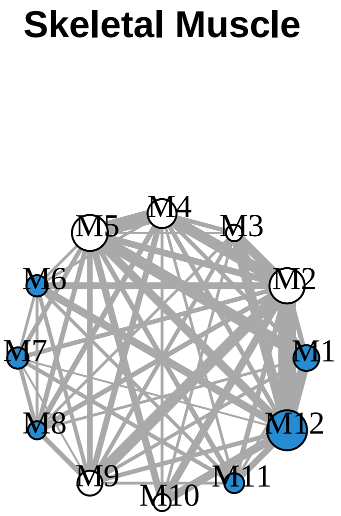


```
# dev.off()
```

For hippocampus, first detected 14 modules, filtered to 12 modules (focus on the modules with more than 10 genes).

```
###### hippocampus network across distant species
net.hippo.tryout <- net.hippo.gc.sig
V(net.hippo.tryout)$ModuleMemb <- ml.com.hippo.sig$membership
module.names.hippo <- sort(unique(V(net.hippo.tryout)$ModuleMemb))

# replacing group of vertices with single meta-verices
net.hippo.coarsen <- igraph::contract.vertices(net.hippo.tryout, ml.com.hippo.sig$membership)
E(net.hippo.coarsen)$weight <- 1


# selecting the modules bigger than size of 10
subg.hippo.sig <- list()
for(g in unique(membership(ml.com.hippo.sig))){
    subg.hippo.sig[[g]] <-
      induced.subgraph(net.hippo.gc.sig, which((membership(ml.com.hippo.sig)==g)
```

```
                                              & ( sizes(ml.com.hippo.sig)[[g]] >= 10)))
}

V(net.hippo.coarsen)$name <- paste0("M", 1:14)

empty.module.hippo <- which(sapply(subg.hippo.sig, vcount) == 0)

net.hippo.coarsen <- net.hippo.coarsen - c(paste0("M", empty.module.hippo))
net.hippo.coarsen <- igraph::simplify(net.hippo.coarsen)

### the size of the nodes
l.cicr.hippo<- layout_in_circle(net.hippo.coarsen)
sizes.hippo <- sapply(subg.hippo.sig, vcount)[-empty.module.hippo]
node.size.hippo <- as.vector(sizes.hippo)

# pdf("~/Project1/manuscript_GSEA/results/Figure4B_Hippocampus_coarsen_net_2.pdf",
# useDingbats = FALSE)
plot(net.hippo.coarsen,
          vertex.size= sqrt(node.size.hippo)*2,
          vertex.label= paste0("M",1:12),
                    vertex.label.color="black",
          margin=.5,
          vertex.color = c("white", "#268bd2", "#268bd2", "#268bd2",  "#268bd2",
                           "white", "white",  "white", "white", "white", "white", "#268bd2"),
                    edge.width=sqrt(E(net.hippo.coarsen)$weight),
                    edge.arrow.size=0, layout = l.cicr.hippo, vertex.label.dist = 1,
                    main = "Hippocampus")
```

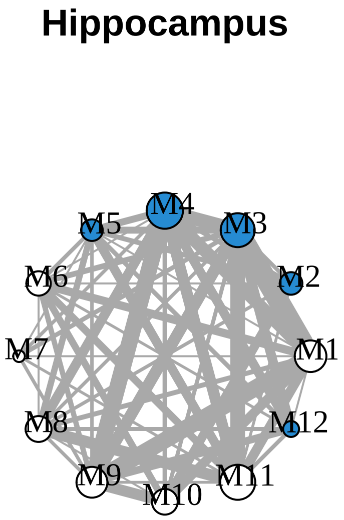


```
# dev.off()
```

```
## mapping to annotation
remap.human.subgraphs.muscle.ml.sig <-
  lapply(subg.muscle.sig, function(x)
    annotation.muscle.oma$human.gene.name[match(V(x)$name, rownames(annotation.muscle.oma))])
remap.human.subgraphs.hippo.ml.sig <-
  lapply(subg.hippo.sig, function(x)
    annotation.hippo.oma$human.gene.name[match(V(x)$name, rownames(annotation.hippo.oma))])
```

```
### significant ones
# muscle
remapping.human.muscle.ml.sig <-
  remap.human.subgraphs.muscle.ml.sig[sapply(remap.human.subgraphs.muscle.ml.sig, length) > 0]
subgraphs.muscle.sig <-
  subg.muscle.sig[sapply(remap.human.subgraphs.muscle.ml.sig, length) > 0]

## hippocampus
remapping.human.hippo.ml.sig <-
  remap.human.subgraphs.hippo.ml.sig[sapply(remap.human.subgraphs.hippo.ml.sig, length) > 0]
subgraphs.hippo.sig <-
  subg.hippo.sig[sapply(remap.human.subgraphs.hippo.ml.sig, length) > 0]


# final number of the modules
length(remapping.human.muscle.ml.sig)
```

```
## [1] 12
```

```
length(remapping.human.hippo.ml.sig)
```

```
## [1] 12
```

```
## loading the human annotation
ensembl.human <- useMart(biomart="ENSEMBL_MART_ENSEMBL", host="www.ensembl.org",
                         path="/biomart/martservice",dataset="hsapiens_gene_ensembl",
                         version = "Ensembl Genes 91")
#
all.ensembl.gene.id.names <- getBM(attributes = c("ensembl_gene_id", "external_gene_name",
                                                  "gene_biotype"),
                                                   values = "*", mart = ensembl.human)
# # 22285 / 22375
all.ensembl.gene.id.names <-
  all.ensembl.gene.id.names[all.ensembl.gene.id.names$gene_biotype == "protein_coding",]


# finding the hubs in the module
hubs.muscle <-
  lapply(subgraphs.muscle.sig, function(x)
    checking.hubs(x, all.ensembl.gene.id.names, annotation.muscle.oma))
hubs.hippo <-
  lapply(subgraphs.hippo.sig, function(x)
    checking.hubs(x, all.ensembl.gene.id.names, annotation.hippo.oma))
```

Perform GO enrichment of the modules. This takes a long time, so the data is loaded automatically, but the code how it was ran is below.

```
#### GO enrichment - on all ontologies, removes redundancies better than goseq.
top.go.enrichment <- function(targets, universe, genome){
  require(topGO)
  all.genes <- factor(as.integer (universe[,1] %in% targets))
    names(all.genes) <- universe[,1]

  ont <- c("BP", "MF", "CC")
```

```r
  ont.res <- list()
  for(i in 1:length(ont)){
        # nodsize is 5
    GOdata <- new("topGOdata", ontology = ont[i], allGenes = all.genes,
                      geneSel = function(p) p < 1e-2, description = "Test",
                      annot = annFUN.org, mapping=genome, ID="Ensembl")
    resultFisher <- runTest(GOdata, algorithm = "elim", statistic = "fisher")
    res <- GenTable(GOdata, elimFisher = resultFisher, topNodes = 40)
    corrected <- p.adjust(as.numeric(res$elimFisher), method="fdr")
    res$FDR <- corrected
    ont.res[[i]] <- res

  }

  names(ont.res) <- ont

  return(ont.res)

}



# ------------
## GO enrichment of the modules in skeletal muscle

# muscle.results.ml.sig <-
# lapply(remapping.human.muscle.ml.sig,
#        function(x) top.go.enrichment(x, all.ensembl.gene.id.names, "org.Hs.eg.db"))
# saveRDS(muscle.results.ml.sig,
# file = "~/Project1//manuscript_GSEA/data_preprocessing/modules_go/muscle_go_enrichment_conservedmodul

# hippo.results.ml.sig <-
# lapply(remapping.human.hippo.ml.sig,
# function(x) top.go.enrichment(x, all.ensembl.gene.id.names, "org.Hs.eg.db")
# saveRDS(hippo.results.ml.sig,
# file = "~/Project1//manuscript_GSEA/data_preprocessing/modules_go/hippo_go_enrichment_conservedmodule

path.go <- "~/Project1/manuscript_GSEA/data_preprocessing/modules_go/"
muscle.modules <- readRDS(paste0(path.go, "muscle_go_enrichment_conservedmodules.rds"))
hippo.modules <- readRDS(paste0(path.go, "hippo_go_enrichment_conservedmodules.rds"))



# --------------------------------------------
### proteostasis-linked modules
library(plyr)
## skeletal muscle - M1, M2, M3, M8, M9
muscle.proteo.modules <- c(2,3,4,6,8)
muscle.modules.go <- muscle.modules[muscle.proteo.modules]
muscle.proteo.module.go <- lapply(muscle.modules.go, '[[', 1)
names(muscle.proteo.module.go) <- paste0("M", c(2,3,4,6,8))
df.muscle.proteo <- ldply(muscle.proteo.module.go, data.frame, .id = "Cluster.Name")
```

```r
## hippocampus - M2, M4, M5, M8, M9
hippo.proteo.modules <- c(2,3,4,12,13)
hippo.modules.go <- hippo.modules[hippo.proteo.modules]
hippo.proteo.module.go <- lapply(hippo.modules.go, '[[', 1)
names(hippo.proteo.module.go) <- paste0("M", c(2,3,4,12,13))
df.hippo.proteo <- ldply(hippo.proteo.module.go, data.frame, .id = "Cluster.Name")
```

Enrichment of the module genes in the GWAS diseases.

Selecting the interesting modules according to proteostasis-linked processes and involvement in GWAS studies.

Skeletal muscle.

```r
### proteasome complex
go.human <- go.gsets("Human")
```

```
## Gene ID type for 'human' is: 'EG'
```

```r
go.human.sets <- go.human$go.sets
go.human.subs <- go.human$go.subs
gobpsets.human <- go.human.sets[go.human.subs$BP]
goccsets.human <- go.human.sets[go.human.subs$CC]

proteasome.complex.human <-
  goccsets.human[which(names(goccsets.human) == "GO:0000502 proteasome complex")]
proteasome.complex.human.symbol <- mapIds(org.Hs.eg.db,
                                          keys=proteasome.complex.human[[1]],
                                          column="SYMBOL",
                                          keytype="ENTREZID",
                                          multiVals="first")
```

```
## 'select()' returned 1:1 mapping between keys and columns
```

```r
# -------------------------
# skeletal muscle module 1 - because of the
module1.human.muscle <- clean.modules(subgraphs.muscle.sig[[1]],
                                      annotation.muscle.oma,
                                      proteasome.complex.human.symbol)
```

```
## 'select()' returned 1:many mapping between keys and columns
```

```
## 'select()' returned 1:1 mapping between keys and columns
```

```r
muscle.community.human.module1 <- module1.human.muscle$module
hubs.of.module1 <- module1.human.muscle$hubs
vertex.of.interest.module1 <-
  V(muscle.community.human.module1)[V(muscle.community.human.module1)$label == names(hubs.of.module1)[3
neigh.nodes.module1 <-
  neighbors(muscle.community.human.module1, vertex.of.interest.module1)$label
interested.genes.module1 <-
  c(as.vector(module1.human.muscle$process), neigh.nodes.module1, names(hubs.of.module1))

# pdf("~/Project1/manuscript_GSEA/results/Figure5A_module1_SM.pdf", 10, 7,
# useDingbats = FALSE)
plot.modules(interested.genes.module1, muscle.community.human.module1,
             neigh.nodes.module1, hubs.of.module1,
```

```
            "Skeletal muscle M1\n(GO:0031146 SCF-dependent proteasomal
            ubiquitin-dependent protein catabolic process")
```

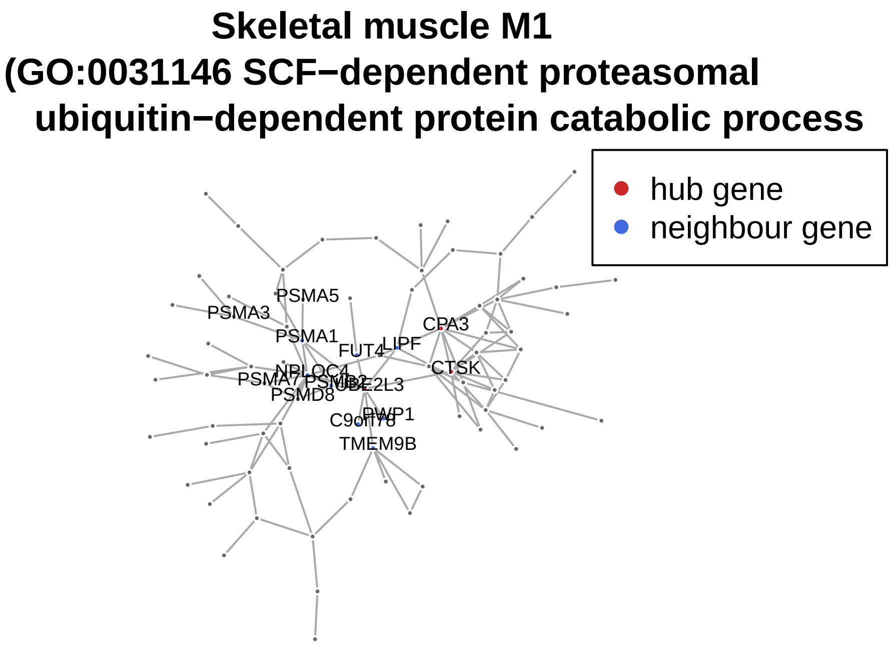


```
# dev.off()



#### ----------------
clean.module3.hippo <- function(subgraph.module,
                                annotation.oma,
                                process.human.symbol){
  tissue.community.module <- subgraph.module
  V(tissue.community.module)$size <- 3
  V(tissue.community.module)$frame.color <- "white"

  ### remap OGs to gene names
  remap.tissue.module <-
                    annotation.oma$human.gene.name[na.omit(match(V(tissue.community.module)$name,
                    rownames(annotation.oma)))]
  names(remap.tissue.module) <- mapIds(org.Hs.eg.db,
                                       keys=as.vector(remap.tissue.module),
                                       column="SYMBOL",
                                       keytype="ENSEMBL",
                                       multiVals="first")

  names(remap.tissue.module)[is.na(names(remap.tissue.module))] <- "AC068775.1"
  genes.process <-
    process.human.symbol[na.omit(match(names(remap.tissue.module),
                                   process.human.symbol))]

  V(tissue.community.module)$label <- names(remap.tissue.module)
  hub.score.module.tissue <- hub.score(tissue.community.module)$vector
  ordered.hubs.module.tissue <-
```

```
    hub.score.module.tissue[order(hub.score.module.tissue, decreasing = TRUE)][1:3]
  matched.hubs.tissue <-
    annotation.oma$human.gene.name[na.omit(match(names(ordered.hubs.module.tissue),
                                                 rownames(annotation.oma)))]

  names(matched.hubs.tissue) <- mapIds(org.Hs.eg.db,
                                       keys = as.vector(matched.hubs.tissue),
                                       column = "SYMBOL",
                                       keytype = "ENSEMBL",
                                       multiVals = "first")

  return(list(module = tissue.community.module, hubs = matched.hubs.tissue, process = genes.process))

}


module3.human.hippo <- clean.module3.hippo(subgraphs.hippo.sig[[3]],
                                           annotation.hippo.oma, proteasome.complex.human.symbol )
```

```
## 'select()' returned 1:many mapping between keys and columns
## 'select()' returned 1:1 mapping between keys and columns
```

```
hippo.community.human.module3 <- module3.human.hippo$module
hubs.of.module3 <- module3.human.hippo$hubs
vertex.of.interest.module3 <-
  V(hippo.community.human.module3)[V(hippo.community.human.module3)$label == names(hubs.of.module3)[1]]
neigh.nodes.module3 <-
  neighbors(hippo.community.human.module3, vertex.of.interest.module3)$label
interested.genes.module3 <-
  c(as.vector(module3.human.hippo$process), neigh.nodes.module3, names(hubs.of.module3))

# pdf("~/Project1/manuscript_GSEA/results/Figure5B_module3_hippo.pdf", 10, 7, useDingbats = FALSE)
plot.modules(interested.genes.module3, hippo.community.human.module3,
             neigh.nodes.module3, hubs.of.module3,
             "Hippocampus M3\n(GO:0000209 protein polyubiquitination)")
```

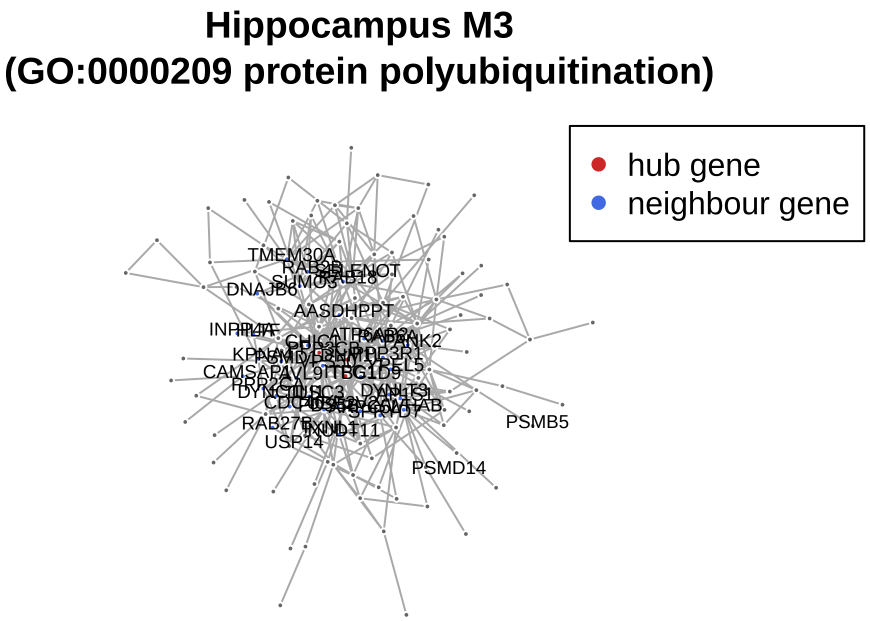


```r
# dev.off()


# ---------------------------
# Figure S10.
# --------------------------


# ---------------------------------
# Skeletal muscle additional module
module12.human.muscle <-
  clean.modules(subgraphs.muscle.sig[[12]], annotation.muscle.oma, proteasome.complex.human.symbol )
```

```
## 'select()' returned 1:1 mapping between keys and columns
## 'select()' returned 1:1 mapping between keys and columns
```

```r
muscle.community.human.module12 <- module12.human.muscle$module
hubs.of.module12 <- module12.human.muscle$hubs
vertex.of.interest.module12 <-
  V(muscle.community.human.module12)[V(muscle.community.human.module12)$label == names(hubs.of.module12
neigh.nodes.module12 <-
  neighbors(muscle.community.human.module12, vertex.of.interest.module12)$label
interested.genes.module12 <-
  c(as.vector(module12.human.muscle$process), neigh.nodes.module12, names(hubs.of.module12))

# pdf("~/Project1/manuscript_GSEA/results/FigureS9_module12_SM.pdf", 10, 7, useDingbats = FALSE)
plot.modules(interested.genes.module12, muscle.community.human.module12,
             neigh.nodes.module12, hubs.of.module12, "Skeletal muscle M12\n(GO:000209 protein polyubiqu
```

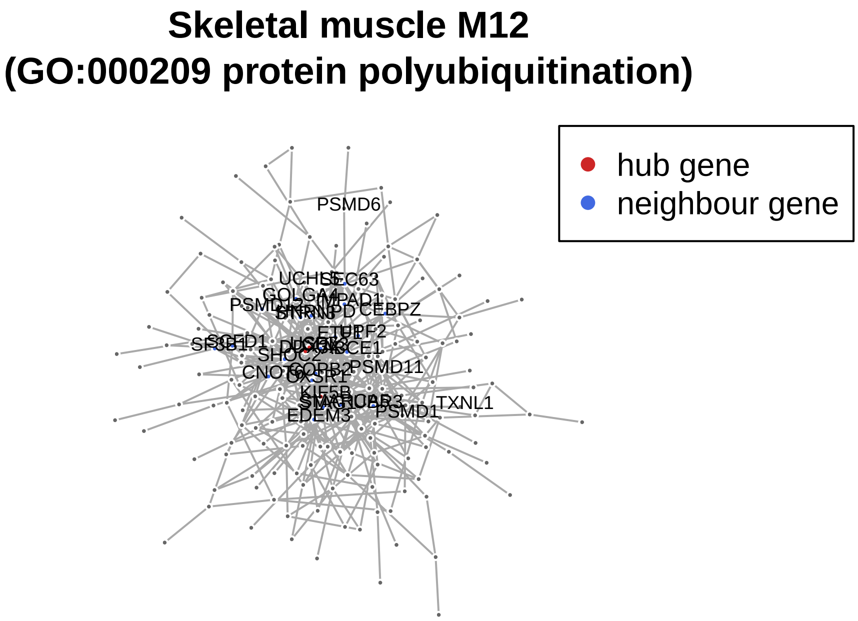


```r
# dev.off()





# ------------------------------------
### Hippocampus additional module
clean.module4.hippo <- function(subgraph.module, annotation.oma, process.human.symbol){
  tissue.community.module <- subgraph.module
  V(tissue.community.module)$size <- 3
  V(tissue.community.module)$frame.color <- "white"

  ### remap OGs to gene names
  remap.tissue.module <-
                    annotation.oma$human.gene.name[na.omit(match(V(tissue.community.module)$name,
                    rownames(annotation.oma)))]
  names(remap.tissue.module) <- mapIds(org.Hs.eg.db,
                                        keys=as.vector(remap.tissue.module),
                                        column="SYMBOL",
                                        keytype="ENSEMBL",
                                        multiVals="first")

  names(remap.tissue.module)[is.na(names(remap.tissue.module))] <- "POLR2A"
  genes.process <-
    process.human.symbol[na.omit(match(names(remap.tissue.module), process.human.symbol))]

  V(tissue.community.module)$label <- names(remap.tissue.module)
  hub.score.module.tissue <- hub.score(tissue.community.module)$vector
  ordered.hubs.module.tissue <-
    hub.score.module.tissue[order(hub.score.module.tissue, decreasing = TRUE)][1:3]
  matched.hubs.tissue <-
    annotation.oma$human.gene.name[na.omit(match(names(ordered.hubs.module.tissue), rownames(annotation
```

```
  names(matched.hubs.tissue) <- mapIds(org.Hs.eg.db,
                                       keys = as.vector(matched.hubs.tissue),
                                       column = "SYMBOL",
                                       keytype = "ENSEMBL",
                                       multiVals = "first")

  return(list(module = tissue.community.module, hubs = matched.hubs.tissue, process = genes.process))

}


# ---------------------------
# hippocampus module 4 - in proteostasis and coronary artery
module4.human.hippo <- clean.module4.hippo(subgraphs.hippo.sig[[4]], annotation.hippo.oma, proteasome.c
```

```
## 'select()' returned 1:1 mapping between keys and columns
## 'select()' returned 1:1 mapping between keys and columns
```

```
hippo.community.human.module4 <- module4.human.hippo$module
hubs.of.module4 <- module4.human.hippo$hubs
vertex.of.interest.module4 <- V(hippo.community.human.module4)[V(hippo.community.human.module4)$label ==
neigh.nodes.module4 <- neighbors(hippo.community.human.module4, vertex.of.interest.module4)$label
interested.genes.module4 <- c(as.vector(module4.human.hippo$process), neigh.nodes.module4, names(hubs.o

# pdf("~/Project1/manuscript_GSEA/results/FigureS9_module4_hippo.pdf", 10, 7, useDingbats = FALSE)
plot.modules(interested.genes.module4, hippo.community.human.module4,
             neigh.nodes.module4, hubs.of.module4,
             "Hippocampus M4\n(GO:1904874 positive regulation
             of telomerase RNA localization to Calaj body)")
```

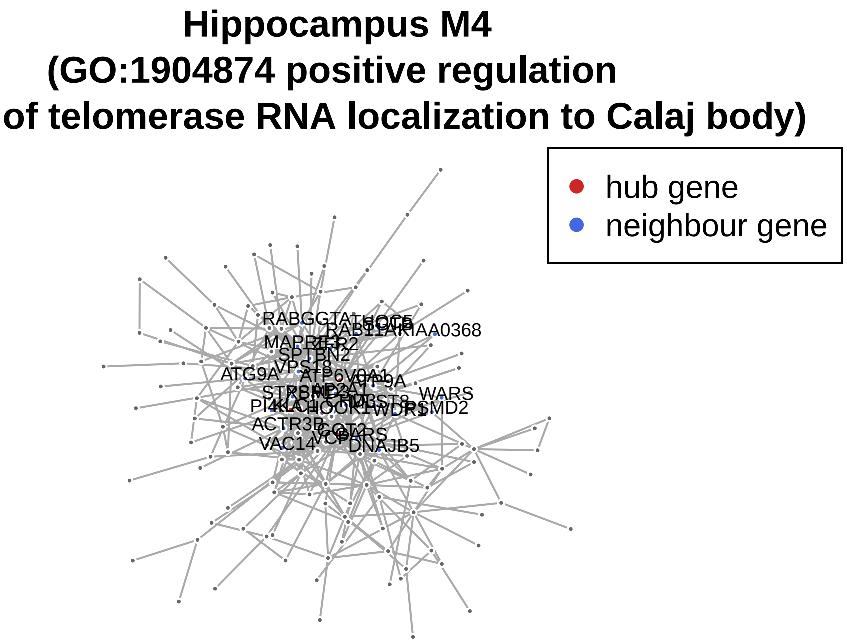


```
# dev.off()
```

Figure 5C. GWAS heatmap.

```
# mapping to GWAS studies - based on gene-level p-values from PASCAL
# downloaded from - http://regulatorycircuits.org/download.html -
# Supplementary data and code: GWAS_gene_scores - 12MB  (University of Lausanne)

gwas.directory <- "~/Project1/manuscript_GSEA/data_preprocessing/GWAS_gene_scores_v1"
gwas.files <- list.files(path = gwas.directory, full.names=TRUE)
gwas.files <- gwas.files[-1]
ldf <- lapply(gwas.files, function(x) read.table(x, header = TRUE))
names(ldf) <- sub("\\.txt", "", list.files(path = gwas.directory))[-1]

# selecting the genes that are having higher of association with GWAS
high.top.ranked.ldf <- lapply(ldf, function(x) x[x$pvalue < 0.1,])


interested.genes <- c( names(hubs.of.module3), as.vector(module3.human.hippo$process),
                       names(hubs.of.module4), as.vector(module4.human.hippo$process),
                       names(hubs.of.module1), as.vector(module1.human.muscle$process),
                       names(hubs.of.module12), as.vector(module12.human.muscle$process) )



interested.genes <- unique(interested.genes)

modules.gwas <-
  lapply(high.top.ranked.ldf, function(x) x[na.omit(match(interested.genes, x$gene_id)),])

# select the gwas that is age related
gwas.age.related <- c("11_rheumatoid_arthritis", "12_multiple_sclerosis", "15_alzheimers",
                      "17_parkinsons_disease", "18_hdl_cholesterol","19_ldl_cholesterol",
                      "20_total_cholesterol", "21_triglycerides", "22_blood_pressure_systolic",
                      "23_coronary_artery_disease", "24_fasting_glucose", "25_glycated_hemoglobin",
                      "26_type_2_diabetes", "27_2hr_glucose", "28_fasting_proinsulin",
                      "29_insulin_secretion", "30_insulin_resistance", "31_beta-cell_function",
                      "32_fasting_insulin", "35_macular_degeneration_neovascular", "36_macular_degenera
                      "37_osteoporosis")

df.modules <- ldply(modules.gwas[gwas.age.related], data.frame)

df.modules$seq <- with(df.modules, ave(pvalue, .id, gene_symbol, FUN = seq_along))
gwas.modules <- reshape2::dcast(.id + seq ~ gene_symbol, data = df.modules, value.var = "pvalue")

gg.mod <- gwas.modules
gg.mod <- gg.mod[,-1]
# missing values are replaced
gg.mod[is.na(gg.mod)] <- 0.99
gg.mod <- -log10(gg.mod)
gg.mod <- gg.mod[,-1]


names.gwas.module <- gwas.modules[,1]
# renaming for the labels on the heatmap
names.gwas.modules <- c("Rheumatoid arthritis", "Multiple sclerosis", "Alzheimer's disease",
                        "Parkinson's disease", "HDL cholesterol", "LDL cholesterol",
```

```
                              "Total cholesterol", "Triglycerides", "Blood pressure systolic",
                              "Coronary artery disease", "Fasting glucose", "Glycated hemoglobin",
                              "Type 2 diabetes", "2hr glucose", "Fasting proinsulin",
                              "Insulin secretion", "Insulin resistance", "Beta-cell function",
                              "Fasting insulin", "Macular degeneration neovasc." ,
                              "Macular degeneration dry", "Osteoporosis")


annotation.mat.gwas <- HeatmapAnnotation(text = anno_text(names.gwas.modules,
                                        rot = 45, just = "left", offset = unit(2, "mm")),
                                        gap = unit(c(2, 4), "mm"),
                                        annotation_height = unit(c(0.5, 0.5, 1.5, 1.5), "cm"))


# pdf("~/Project1/manuscript_GSEA/results/Figure5C_GWAS_heatmap_2.pdf", 11, 6, useDingbats = FALSE)
Heatmap(t(as.matrix(gg.mod)),
        heatmap_legend_param = list(color_bar = "discrete"),
        name =  "GWAS gene score\n(-log10(p-value))",
        cluster_columns = FALSE,
        show_column_dend = FALSE,
        top_annotation = annotation.mat.gwas,
        col = colorRamp2(seq(0,5,1), brewer.pal(6,"Paired")))
```

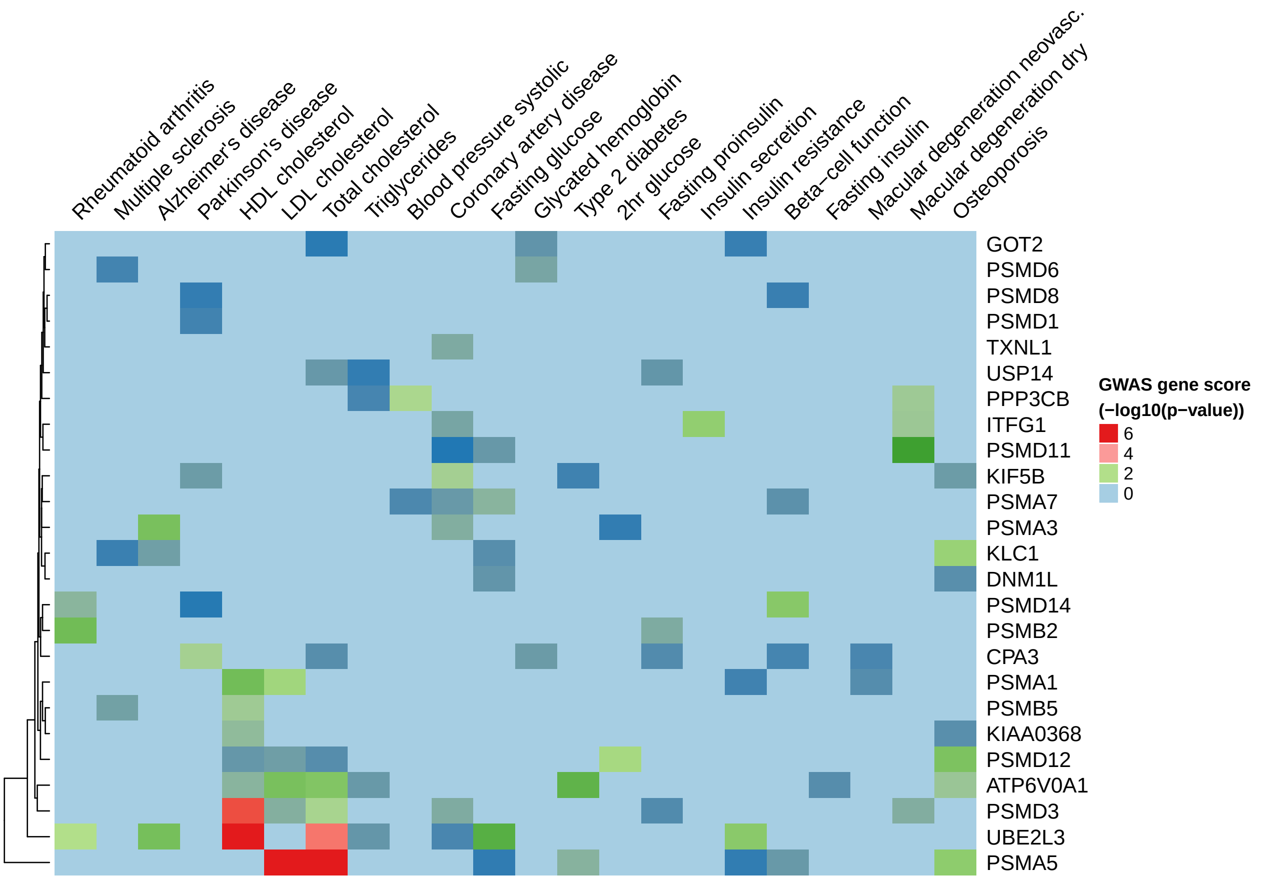


```
# dev.off()

## ggmod should go to the Suplement table
```

Generating the dietary restriction volcano plot for Figure 5D.

```
#### ---------------------------------------

### VOLCANO PLOTS for caloric restriction
path.cr <- "~/Project1/manuscript_GSEA/data_preprocessing/hsapiens/caloric_restriction_data/"
deg.human.dr <- readRDS(paste0(path.cr, "differential_expression_human_dietary_restriction.rds"))

## select genes from GWAS heatmap above
colnames.gwas <- mapIds(org.Hs.eg.db,
                        keys=colnames(gg.mod),
                        column="ENSEMBL",
                        keytype="SYMBOL",
                        multiVals="first")
```

```
## 'select()' returned 1:1 mapping between keys and columns
```

```
deg.human.muscle.dr.candidate.genes <-
  deg.human.dr[na.omit(match(as.vector(colnames.gwas), rownames(deg.human.dr))),]

deg.human.muscle.dr.candidate.genes$Gene <- mapIds(org.Hs.eg.db,
                                                    keys=rownames(deg.human.muscle.dr.candidate.genes),
                                                    column="SYMBOL",
                                                    keytype="ENSEMBL",
                                                    multiVals="first")
```

```
## 'select()' returned 1:1 mapping between keys and columns
```

```
deg.human.muscle.dr.candidate.genes$Significant <-
  ifelse(deg.human.muscle.dr.candidate.genes$adj.P.Val < 0.05, "FDR < 0.05", "FDR > 0.05")

# candidate genes in human dietary restriciton analysis
library(ggrepel)
p.candidate.genes <- ggplot(deg.human.muscle.dr.candidate.genes, aes(x = logFC, y = -log10(P.Value))) +
  geom_point(aes(color = Significant)) +
  scale_color_manual(values = c("red", "grey")) +
  theme_bw(base_size = 12) + theme(legend.position = "bottom") +
  ggtitle("H. sapiens - Caloric restriction (skeletal muscle)") +
  xlab(expression(log[2]~fold~change~(CR/Control))) + ylab(expression(-log[10]~pvalue)) + xlim(-2,4) +
  geom_vline(xintercept = 0) +
  geom_text_repel(
    data = subset(deg.human.muscle.dr.candidate.genes, adj.P.Val < 0.05),
    aes(label = Gene),
    size = 5,
    box.padding = unit(0.35, "lines"),
    point.padding = unit(0.3, "lines")
  )

# pdf("~/Project1/manuscript_GSEA/results/Figure5D_volcano_plots_on_caloric_restriction.pdf",
# 7, 5, useDingbats = FALSE)
plot(p.candidate.genes)
```

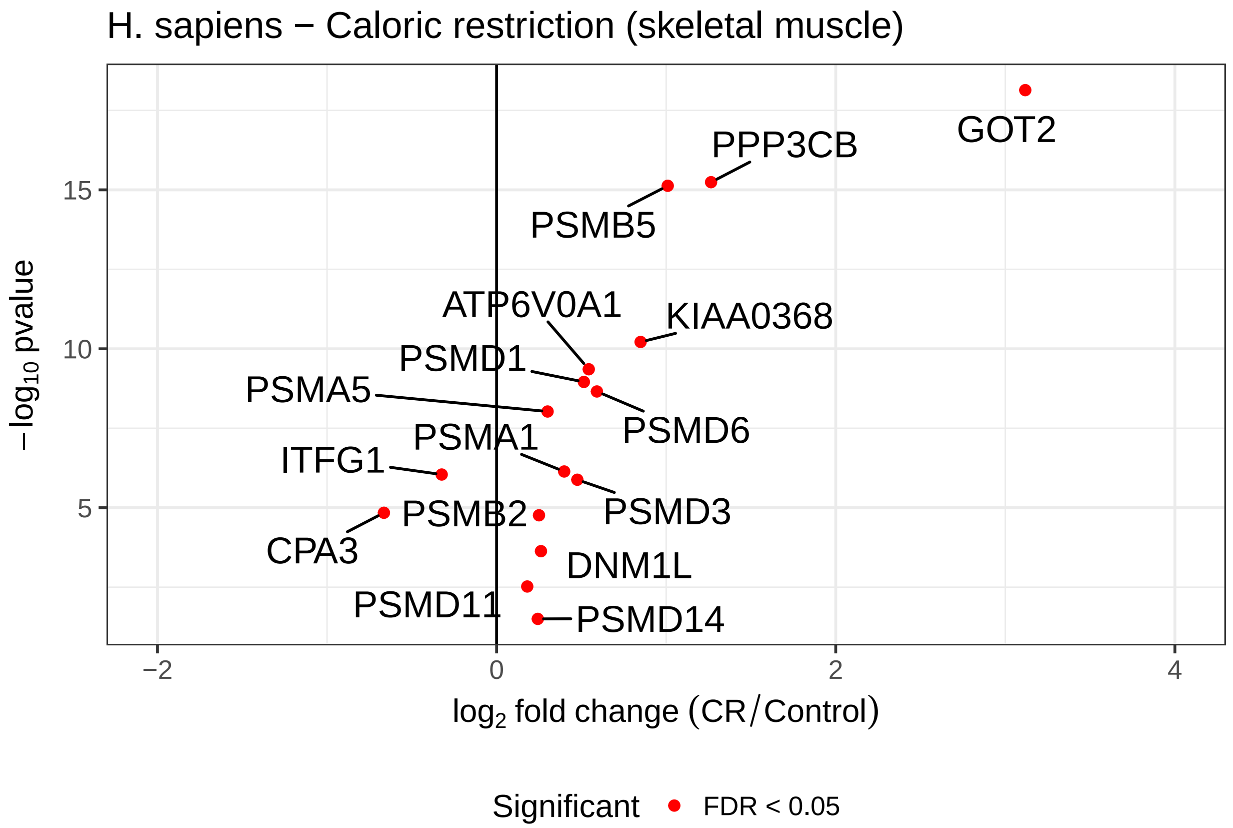


```
# dev.off()
```

Supplemental material for this part. Table S8 and Table S9.

```
# --------------------------------
### Table S8.
# --------------------------------

### proteostasis-linked modules

## skeletal muscle - M1, M6, M7, M8, M11, M12
muscle.proteo.modules <- c(1,6,7,8,11,12)
muscle.modules.go <- muscle.modules[muscle.proteo.modules]
muscle.proteo.module.go <- lapply(muscle.modules.go, '[[', 1)
names(muscle.proteo.module.go) <- paste0("M", c(1,6,7,8,11,12))
df.muscle.proteo <- ldply(muscle.proteo.module.go, data.frame, .id = "Cluster.Name")


# GWAS linked modules - M3, M4, M5, M12
muscle.disease.modules <- c(3,4,5,12)
muscle.modules.gwas <- muscle.gwas.sig[muscle.disease.modules]
names(muscle.modules.gwas) <- paste0("M", c(3,4,5,12))
df.muscle.proteo.gwas <- ldply(muscle.modules.gwas, data.frame, .id = "Cluster.Name")


# proteostasis-linked
## hippocampus - M2, M3, M4, M5, M12
hippo.proteo.modules <- c(2,3,4,5,12)
```

```r
hippo.modules.go <- hippo.modules[hippo.proteo.modules]
hippo.proteo.module.go <- lapply(hippo.modules.go, '[[', 1)
names(hippo.proteo.module.go) <- paste0("M", c(2,3,4,5,12))
df.hippo.proteo <- ldply(hippo.proteo.module.go, data.frame, .id = "Cluster.Name")


# GWAS linked modules
hippo.disease.modules <- c(4,6)
hippo.modules.gwas <- hippo.gwas.sig[hippo.disease.modules]
names(hippo.modules.gwas) <- paste0("M", 4)
df.hippo.proteo.gwas <- ldply(hippo.modules.gwas, data.frame, .id = "Cluster.Name")


df.module.sizes <- data.frame(module.names = paste0("M", 1:12),
                              muscle.modules.gene.size = sapply(subgraphs.muscle.sig, vcount),
                              hippo.modules.gene.size = sapply(subgraphs.hippo.sig, vcount))

df.modules.summary.stats <-
  list(Module.sizes = df.module.sizes,
             Muscle.GO.proteo.enrich = df.muscle.proteo,
             Hippo.GO.proteo.enrich = df.hippo.proteo,
             Muscle.GWAS.proteo.enrich = df.muscle.proteo.gwas,
             Hippo.GWAS.proteo.enrich = df.hippo.proteo.gwas)



#library(WriteXLS)
# WriteXLS(df.modules.summary.stats,
#     ExcelFileName = "~/Project1/manuscript_GSEA/supplementary_data/Table_S8.xlsx", SheetNames = NULL)



# -----------------------------
# Table S9.
# ----------------------------

# matrix with gene scores associated to GWAS diseases
gg.mod.supplement <- gwas.modules[,-2]
gwas.diseases.genes <- gg.mod.supplement


# WriteXLS(gwas.diseases.genes,
#    ExcelFileName = "~/Project1/manuscript_GSEA/supplementary_data/Table_S9.xlsx", SheetNames = NULL)



# -----------------------------
# Figure S9.
# ----------------------------

# Plotting random networks.

########
```

```r
# for p-value calculations
# https://www.ncbi.nlm.nih.gov/pubmed/21044043
# calculate significance for the random networks
calculate.p.val <- function(observed, randomized){
  # here is mean to summarize 100 randomizations
  random <- mean(randomized)
  return( (sum(abs(random) > abs(observed)) + 1) / (length(randomized) + 1) )
}



setwd("~/Project1/manuscript_GSEA/results/")
# hippo_random_nets <- readRDS("random_nets_hippo100.rds")
hippo_random_nets <- readRDS("random_nets_hippo100.rds")
summary.hippo <- as.matrix(sapply(hippo_random_nets, '['))


# muscle_random_nets <- readRDS("random_nets_muscle100.rds")
muscle_random_nets <- readRDS("random_nets_muscle100.rds")
summary.muscle <- as.matrix(sapply(muscle_random_nets, '['))


## number of connections
obs.no.connections.muscle <- 2887
obs.no.connections.hippo <- 3353


#pdf("SupplementS9_randomnets100_2.pdf", 7,7)
par(mfrow = c(2,1))
# for number of connections
hist(as.numeric(summary.muscle[,1]), xlim = c(2700, max(as.numeric(summary.muscle[,1]))),
  xlab = "Number of connections\n(skeletal muscle)", main = "Skeletal muscle network")
abline(v = obs.no.connections.muscle, col = "red", lwd = 2)
legend(3000,30, "Observed",
lty=1,
lwd=2.5, col= "red")

hist(as.numeric(summary.hippo[,1]), xlim = c(3200, max(as.numeric(summary.hippo[,1]))),
  xlab = "Number of connections\n(hippocampus)", main = "Hippocampus network")
abline(v = obs.no.connections.hippo, col = "red", lwd = 2)
```

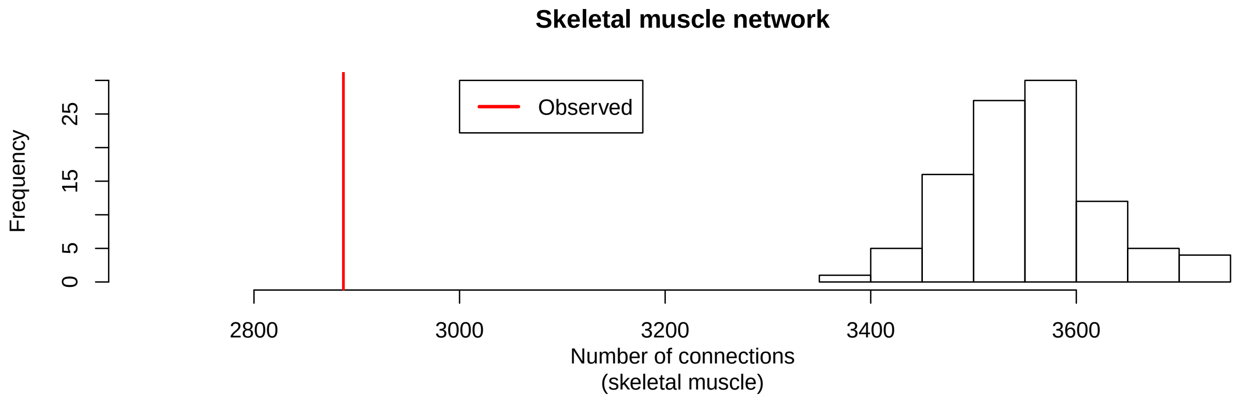


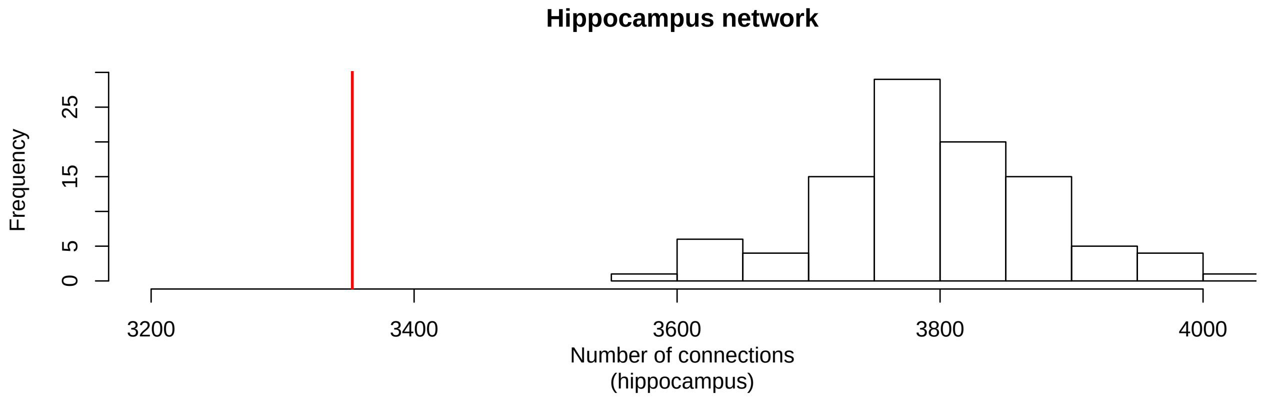


```r
# dev.off()
```

```r
sessionInfo()
```

```
## R version 3.4.3 (2017-11-30)
## Platform: x86_64-apple-darwin15.6.0 (64-bit)
## Running under: macOS Sierra 10.12.1
##
## Matrix products: default
## BLAS: /Library/Frameworks/R.framework/Versions/3.4/Resources/lib/libRblas.0.dylib
## LAPACK: /Library/Frameworks/R.framework/Versions/3.4/Resources/lib/libRlapack.dylib
##
## locale:
## [1] en_US.UTF-8/en_US.UTF-8/en_US.UTF-8/C/en_US.UTF-8/en_US.UTF-8
##
## attached base packages:
##  [1] parallel  stats4    grid      stats     graphics  grDevices utils
##  [8] datasets  methods   base
##
## other attached packages:
##  [1] GO.db_3.5.0            dplyr_0.7.4            data.table_1.10.4-3
##  [4] ggrepel_0.7.0          cowplot_0.9.2          graphite_1.24.1
##  [7] gage_2.28.2            org.Ce.eg.db_3.5.0     org.Dm.eg.db_3.5.0
## [10] org.Mm.eg.db_3.5.0     org.Hs.eg.db_3.5.0     AnnotationDbi_1.40.0
## [13] IRanges_2.12.0         S4Vectors_0.16.0       Biobase_2.38.0
## [16] BiocGenerics_0.24.0    igraph_1.1.2           circlize_0.4.3
## [19] ComplexHeatmap_1.17.1  WriteXLS_4.0.0         plyr_1.8.4
```

```
## [22] goseq_1.30.0           geneLenDataBase_1.14.0 BiasedUrn_1.07
## [25] biomaRt_2.34.2         RobustRankAggreg_1.1   Hmisc_4.1-1
## [28] ggplot2_2.2.1          Formula_1.2-2          survival_2.41-3
## [31] lattice_0.20-35        RColorBrewer_1.1-2
##
## loaded via a namespace (and not attached):
##  [1] nlme_3.1-131.1             bitops_1.0-6
##  [3] matrixStats_0.53.1         bit64_0.9-7
##  [5] progress_1.1.2             httr_1.3.1
##  [7] rprojroot_1.3-2            GenomeInfoDb_1.14.0
##  [9] tools_3.4.3                backports_1.1.2
## [11] R6_2.2.2                   rpart_4.1-13
## [13] DBI_0.8                    lazyeval_0.2.1
## [15] mgcv_1.8-23                colorspace_1.3-2
## [17] nnet_7.3-12                GetoptLong_0.1.6
## [19] gridExtra_2.3              prettyunits_1.0.2
## [21] RMySQL_0.10.14             curl_3.1
## [23] bit_1.1-12                 compiler_3.4.3
## [25] graph_1.56.0               htmlTable_1.11.2
## [27] DelayedArray_0.4.1         labeling_0.3
## [29] rtracklayer_1.38.3         scales_0.5.0
## [31] checkmate_1.8.5            rappdirs_0.3.1
## [33] stringr_1.3.0              digest_0.6.15
## [35] Rsamtools_1.30.0           foreign_0.8-69
## [37] rmarkdown_1.9              XVector_0.18.0
## [39] base64enc_0.1-3            pkgconfig_2.0.1
## [41] htmltools_0.3.6            htmlwidgets_1.0
## [43] rlang_0.2.0                GlobalOptions_0.0.12
## [45] rstudioapi_0.7             RSQLite_2.0
## [47] bindr_0.1                  shape_1.4.4
## [49] BiocParallel_1.12.0        acepack_1.4.1
## [51] RCurl_1.95-4.10            magrittr_1.5
## [53] GenomeInfoDbData_1.0.0     Matrix_1.2-12
## [55] Rcpp_0.12.15               munsell_0.4.3
## [57] stringi_1.1.6              yaml_2.1.17
## [59] SummarizedExperiment_1.8.1 zlibbioc_1.24.0
## [61] blob_1.1.0                 Biostrings_2.46.0
## [63] splines_3.4.3              GenomicFeatures_1.30.3
## [65] KEGGREST_1.18.1            knitr_1.20
## [67] pillar_1.2.1               GenomicRanges_1.30.3
## [69] rjson_0.2.15               reshape2_1.4.3
## [71] glue_1.2.0                 XML_3.98-1.10
## [73] evaluate_0.10.1            latticeExtra_0.6-28
## [75] png_0.1-7                  gtable_0.2.0
## [77] assertthat_0.2.0           tibble_1.4.2
## [79] GenomicAlignments_1.14.1   memoise_1.1.0
## [81] bindrcpp_0.2               cluster_2.0.6
```

### Main Figures + Supplement (Network part)

*Andrea Komljenović*

*3/8/2018*

**Step III**

Loading the packages.

Functions for the networks and module search.

```
## ----------------
# FUNCTIONS for constructing networks
## ----------------


### CORRELATION / Co-expression MATRIX
significant.edges <- function(expression.matrix){
    require(Hmisc)
  siglevel <- 0.05
    corrmatr <- rcorr(t(expression.matrix),type="pearson")
    corrmat <- corrmatr$r
    diag(corrmat) <- 0
    corrmat[corrmatr$P > siglevel] <- 0
    corrmat[corrmat < 0 ] <- 0
    # flatten correlation matrix for plotting
    corrmat.flat <- flattenCorrMatrix(corrmat)
  return( corrmat.flat)

}

flattenCorrMatrix <- function(cormat) {
  ut <- upper.tri(cormat, diag = FALSE)
  return(data.frame(
    row = rownames(cormat)[row(cormat)[ut]],
    column = rownames(cormat)[col(cormat)[ut]],
    cor  =(cormat)[ut]
    ))
}



preprocess.edges <- function(expression.matrix, annotation.species, annotation){

    mm <- match(rownames(expression.matrix), annotation.species)
    annot <- annotation[mm,]

    rownames(expression.matrix) <- rownames(annot)
    corrmat <- significant.edges(expression.matrix)
    return(corrmat)
}
```

```r
# data integration of co-expression networks per species / per tissue
data.integration <- function(coexp.sp1, coexp.sp2, coexp.sp3, coexp.sp4, n.genes.tissue){
  # require(RobustRankAggreg)
  correlations <- list(human = coexp.sp1, mouse = coexp.sp2, fly = coexp.sp3, worm = coexp.sp4)
  cat("Number of gene pairs:", sapply(correlations, nrow), "\n")

  coexpr.mat <- lapply(correlations, function(corrs) {
                              rownames(corrs) <- paste(corrs$row, corrs$column , sep = ":")
                              return(corrs)
                       })
  # order per species
  sorted.coexpr.mat <- lapply(coexpr.mat, function(x) {
                            x <- x[order(x$cor, na.last = TRUE, decreasing = TRUE), ]
                            return(x) })
  sorted.correlations.rownames <- lapply(sorted.coexpr.mat, function(s) rownames(s))
  rank.matrix <- rankMatrix(sorted.correlations.rownames, N = n.genes.tissue, full = TRUE)
  aggregated.ranks <- aggregateRanks(rmat = rank.matrix, N = n.genes.tissue, method = "RRA")
  return(list(aggregation = aggregated.ranks, ranks.per.species = rank.matrix))
}


clean.modules <- function(subgraph.module, annotation.oma, process.human.symbol){
  tissue.community.module <- subgraph.module
  V(tissue.community.module)$size <- 3
  V(tissue.community.module)$frame.color <- "white"

  ### remap OGs to gene names
  remap.tissue.module <-
                  annotation.oma$human.gene.name[match(V(tissue.community.module)$name,
                  rownames(annotation.oma))]
  names(remap.tissue.module) <- mapIds(org.Hs.eg.db,
                                        keys=as.vector(remap.tissue.module),
                                        column="SYMBOL",
                                        keytype="ENSEMBL",
                                        multiVals="first")


  genes.process <- process.human.symbol[na.omit(match(names(remap.tissue.module),
                                                  process.human.symbol))]

  V(tissue.community.module)$label <- names(remap.tissue.module)
  hub.score.module.tissue <- hub.score(tissue.community.module)$vector
  ordered.hubs.module.tissue <- hub.score.module.tissue[order(hub.score.module.tissue,
                                                        decreasing = TRUE)][1:3]
  matched.hubs.tissue <- annotation.oma$human.gene.name[match(names(ordered.hubs.module.tissue),
                                                        rownames(annotation.oma))]

  names(matched.hubs.tissue) <- mapIds(org.Hs.eg.db,
                                      keys = as.vector(matched.hubs.tissue),
                                      column = "SYMBOL",
                                      keytype = "ENSEMBL",
                                      multiVals = "first")
```

```r
  return(list(module = tissue.community.module, hubs = matched.hubs.tissue, process = genes.process))

}



## plotting the modules
plot.modules <- function(genes.of.interest, community.module.of.interest,
                         neighbourhood.nodes, hubs.genes, plot.title){

  idx <- na.omit(match(genes.of.interest, V(community.module.of.interest)$label))
  V(community.module.of.interest)$label[idx] <- genes.of.interest
  V(community.module.of.interest)$label[-idx] <- ""
  # neighbouring genes
  V(community.module.of.interest)$color <- "grey40"
  V(community.module.of.interest)$color[match(neighbourhood.nodes,
                                              V(community.module.of.interest)$label)] <- "royalblue"
  # hub genes
  V(community.module.of.interest)$color[match(names(hubs.genes),
                                              V(community.module.of.interest)$label)] <- "firebrick3"

  tissue.layout <- layout.kamada.kawai(community.module.of.interest)
  plot(community.module.of.interest,
         layout = tissue.layout,
         main = plot.title,
         vertex.label.color = "black",
         vertex.label.cex = 0.6,
         vertex.size = 2.5,
         vertex.label.dist = 0.4,

         vertex.label.family = "Helvetica")

         legend("topright", c("hub gene", "neighbour gene"), pch = 16,
       col = c("firebrick3", "royalblue") )


}


checking.hubs <- function(module, ensembl.genes.protein.coding, annotation.oma){

  tissue.community.module <- module

  remap.tissue.module <-
                annotation.oma$human.gene.name[match(V(tissue.community.module)$name,
                rownames(annotation.oma))]

  gene.name.module <-
    ensembl.genes.protein.coding[match(remap.tissue.module,
            ensembl.genes.protein.coding$ensembl_gene_id),]
  V(tissue.community.module)$label <- gene.name.module$external_gene_name
  V(tissue.community.module)$name <- gene.name.module$external_gene_name
```

```
  hub.score.module <- hub.score(tissue.community.module)$vector
  ordered.hubs.module <- hub.score.module[order(hub.score.module, decreasing = TRUE)][1:20]
  ensembl.module.hubs <-
    ensembl.genes.protein.coding[match(names(ordered.hubs.module [1:20]),
    ensembl.genes.protein.coding$external_gene_name),]
  return(ensembl.module.hubs)

}
```

Loading the expression matrices for networks and annotation of the orthogroups.

```
path.oma.exp <- "~/Project1/manuscript_GSEA/data_preprocessing/expressionmat_for_networks/"

# ------------------------
# skeletal muscle
# human
expMatHuman.OG.muscle.oma.sig <- readRDS(paste0(path.oma.exp, "expmat_hsapiens_muscle_oma_sig.rds"))

# mouse
expMatMouse.OG.muscle.oma.sig <- readRDS(paste0(path.oma.exp,"expmat_mmusculus_muscle_oma_sig.rds"))
## add sample names
colnames(expMatMouse.OG.muscle.oma.sig) <- c(rep("young", 4), rep("old",5))

# fly
expMatFly.OG.muscle.oma.sig <- readRDS(paste0(path.oma.exp, "expmat_dmelanogaster_muscle_oma_sig.rds"))

# worm
expMatWorm.OG.muscle.oma.sig <- readRDS(paste0(path.oma.exp, "expmat_celegans_muscle_oma_sig.rds"))
# remove too young samples
expMatWorm.OG.muscle.oma.sig <- expMatWorm.OG.muscle.oma.sig[, -c(1:3)]


# ----------------------
# hippocampus significant
# human
expMatHuman.OG.hippo.oma.sig <- readRDS(paste0(path.oma.exp, "expmat_hsapiens_hippo_oma_sig.rds"))

# mouse
expMatMouse.OG.hippo.oma.sig <- readRDS(paste0(path.oma.exp,"expmat_mmusculus_hippo_oma_sig.rds"))

# fly
expMatFly.OG.hippo.oma.sig <- readRDS(paste0(path.oma.exp, "expmat_dmelanogaster_hippo_oma_sig.rds"))

# worm
expMatWorm.OG.hippo.oma.sig <- readRDS(paste0(path.oma.exp, "expmat_celegans_hippo_oma_sig.rds"))
# remove too young samples
expMatWorm.OG.hippo.oma.sig <- expMatWorm.OG.hippo.oma.sig[, -c(1:3)]



# --------------------
# annotation
annotation.muscle.oma <- readRDS("~/Project1/manuscript_GSEA/results/annotation_muscle_oma.rds")
annotation.hippo.oma <- readRDS("~/Project1/manuscript_GSEA/results/annotation_hippocampus_oma.rds")
```

```r
# skeletal muscle
human.corr.muscle.oma.sig <- preprocess.edges(expMatHuman.OG.muscle.oma.sig,
                                              annotation.muscle.oma$human.gene.name,
                                              annotation.muscle.oma)
mouse.corr.muscle.oma.sig <- preprocess.edges(expMatMouse.OG.muscle.oma.sig,
                                              annotation.muscle.oma$mouse.gene.name,
                                              annotation.muscle.oma)
fly.corr.muscle.oma.sig <- preprocess.edges(expMatFly.OG.muscle.oma.sig,
                                            annotation.muscle.oma$fly.gene.name,
                                            annotation.muscle.oma)
worm.corr.muscle.oma.sig <- preprocess.edges(expMatWorm.OG.muscle.oma.sig,
                                             annotation.muscle.oma$worm.gene.name,
                                             annotation.muscle.oma)

# hippocampus
human.corr.hippo.oma.sig <- preprocess.edges(expMatHuman.OG.hippo.oma.sig,
                                             annotation.hippo.oma$human.gene.name,
                                             annotation.hippo.oma)
mouse.corr.hippo.oma.sig <- preprocess.edges(expMatMouse.OG.hippo.oma.sig,
                                             annotation.hippo.oma$mouse.gene.name,
                                             annotation.hippo.oma)
fly.corr.hippo.oma.sig <- preprocess.edges(expMatFly.OG.hippo.oma.sig,
                                           annotation.hippo.oma$fly.gene.name,
                                           annotation.hippo.oma)
worm.corr.hippo.oma.sig <- preprocess.edges(expMatWorm.OG.hippo.oma.sig,
                                            annotation.hippo.oma$worm.gene.name,
                                            annotation.hippo.oma)


# number of the
n.muscle.eog.oma.sig <- nrow(expMatHuman.OG.muscle.oma.sig)
n.muscle.eog.oma.sig # 2010
```

```
## [1] 2010
```

```r
n.hippo.eog.oma.sig <- nrow(expMatHuman.OG.hippo.oma.sig)
n.hippo.eog.oma.sig # 2075
```

```
## [1] 2075
```

Real data network data integration.

```r
## integration of co-expression links across species

muscle.coexpres.oma.sig <- data.integration(human.corr.muscle.oma.sig, mouse.corr.muscle.oma.sig,
                                            fly.corr.muscle.oma.sig, worm.corr.muscle.oma.sig,
                                            n.muscle.eog.oma.sig)
```

```
## Number of gene pairs: 2019045 2019045 2019045 2019045
```

```r
#  (2019045 links - pearson correlation method)
hippo.coexpres.oma.sig <- data.integration(human.corr.hippo.oma.sig, mouse.corr.hippo.oma.sig,
                                           fly.corr.hippo.oma.sig, worm.corr.hippo.oma.sig,
                                           n.hippo.eog.oma.sig)
```

```
## Number of gene pairs: 2151775 2151775 2151775 2151775
```

```r
# (2151775 links - pearson correlation method)
```

Significance threshold defined for the co-expression links consideration.

```r
# defining the threshold for significant co-expression links
sig.level <- 0.001
# muscle
aggregated.signif.muscle.oma.sig <-
  muscle.coexpres.oma.sig$aggregation[which(muscle.coexpres.oma.sig$aggregation$Score <= sig.level), ]
nrow(aggregated.signif.muscle.oma.sig) #  2887 significant co-expression links
```

```
## [1] 2887
```

```r
# hippocampus
aggregated.signif.hippo.oma.sig <-
  hippo.coexpres.oma.sig$aggregation[which(hippo.coexpres.oma.sig$aggregation$Score <= sig.level), ]
nrow(aggregated.signif.hippo.oma.sig) # 3353 significant co-expression links
```

```
## [1] 3353
```

Finding the giant component of the network for further analysis.

```r
# defining the edges
edge.list.muscle.oma.sig <- do.call(rbind, strsplit(rownames(aggregated.signif.muscle.oma.sig), ':'))
edge.list.hippo.oma.sig <- do.call(rbind, strsplit(rownames(aggregated.signif.hippo.oma.sig), ':'))


# 1142 genes
net.muscle.oma.sig <- graph_from_data_frame(d = edge.list.muscle.oma.sig, directed = FALSE)
# 1098 genes
net.hippo.oma.sig <- graph_from_data_frame(d = edge.list.hippo.oma.sig, directed = FALSE)

## weighting the edges for multilevel community
E(net.muscle.oma.sig)$weight <- -log10(aggregated.signif.muscle.oma.sig$Score)
E(net.hippo.oma.sig)$weight <- -log10(aggregated.signif.hippo.oma.sig$Score)


# decompose graph in order to remove singletones
comps.muscle.sig <- decompose.graph(net.muscle.oma.sig)
table(sapply(comps.muscle.sig, vcount))
```

```
##
##    2    3    4 1050
##   32    8    1    1
```

```r
# giant component
# 2    3    4 1050
#  32    8    1    1

comps.hippo.sig <- decompose.graph(net.hippo.oma.sig)
table(sapply(comps.hippo.sig, vcount))
```

```
##
##    2    3 1067
##   14    1    1
```

```r
# giant compononet
# 2    3 1067
```

```
#   14    1    1
```

Performing the greedy search algorithm (multilevel) on the networks in order to find modules, then calculating the sizes of the modules.

```
### networks
net.muscle.gc.sig <- decompose.graph(net.muscle.oma.sig)[[1]]
net.hippo.gc.sig <- decompose.graph(net.hippo.oma.sig)[[1]]

### searcing for cross-species modules
ml.com.muscle.sig <- multilevel.community(net.muscle.gc.sig)
ml.com.hippo.sig <- multilevel.community(net.hippo.gc.sig)

# muscle
multilevel.sizesComm.muscle.sig <- sizes(ml.com.muscle.sig)
multilevel.numComm.muscle.sig <- length(multilevel.sizesComm.muscle.sig )


# hippocampus
multilevel.sizesComm.hippo.sig <- sizes(ml.com.hippo.sig)
multilevel.numComm.hippo.sig <- length(multilevel.sizesComm.hippo.sig )
```

For skeletal muscle, first detected 20 modules, filtered to 10 modules (focus on the modules with more than 10 genes).

```
## building the network
net.muscle.tryout <- net.muscle.gc.sig
V(net.muscle.tryout)$ModuleMemb <- ml.com.muscle.sig$membership
module.names <- sort(unique(V(net.muscle.tryout)$ModuleMemb))


# replacing group of vertices with single meta-vertices
net.muscle.coarsen <- igraph::contract.vertices(net.muscle.tryout,
                                                ml.com.muscle.sig$membership)
E(net.muscle.coarsen)$weight <- 1


# selecting the modules bigger than size of 10
subg.muscle.sig <- list()
for(g in unique(membership(ml.com.muscle.sig))){
      subg.muscle.sig[[g]] <-
        induced.subgraph(net.muscle.gc.sig,
                    which((membership(ml.com.muscle.sig)==g)
                      & ( sizes(ml.com.muscle.sig)[[g]] >= 10)))
}
# nameing the modules
V(net.muscle.coarsen)$name <- paste0("M", 1:20)

# defines which modules are empty
empty.module <- which(sapply(subg.muscle.sig, vcount) == 0)
## this removes those small modules
net.muscle.coarsen <- net.muscle.coarsen - c(paste0("M", empty.module))
net.muscle.coarsen <- igraph::simplify(net.muscle.coarsen)

### the size of the nodes
```

```
l.cicr <- layout_in_circle(net.muscle.coarsen)
sizes <- sapply(subg.muscle.sig, vcount)[-empty.module]
node.size <- as.vector(sizes)


# pdf("~/Project1/manuscript_GSEA/results/Figure4A_SkeletalMuscle_coarsen_net_2.pdf",
# useDingbats = FALSE)
plot(net.muscle.coarsen,
          vertex.size= sqrt(node.size)*2,
          vertex.label= paste0("M",1:12),
                    vertex.label.color="black",
          margin=.5,
          vertex.color = c("#268bd2", "white", "white", "white","white", "#268bd2",
                           "#268bd2", "#268bd2", "white", "white", "#268bd2", "#268bd2"),
                    edge.width=sqrt(E(net.muscle.coarsen)$weight),
                    edge.arrow.size=0, layout = l.cicr, vertex.label.dist = 1,
                    main = "Skeletal Muscle")
```

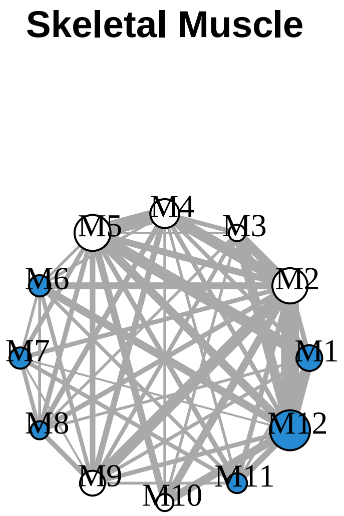


```
# dev.off()
```

For hippocampus, first detected 14 modules, filtered to 12 modules (focus on the modules with more than 10 genes).

```
###### hippocampus network across distant species
net.hippo.tryout <- net.hippo.gc.sig
V(net.hippo.tryout)$ModuleMemb <- ml.com.hippo.sig$membership
module.names.hippo <- sort(unique(V(net.hippo.tryout)$ModuleMemb))

# replacing group of vertices with single meta-verices
net.hippo.coarsen <- igraph::contract.vertices(net.hippo.tryout, ml.com.hippo.sig$membership)
E(net.hippo.coarsen)$weight <- 1


# selecting the modules bigger than size of 10
subg.hippo.sig <- list()
for(g in unique(membership(ml.com.hippo.sig))){
    subg.hippo.sig[[g]] <-
      induced.subgraph(net.hippo.gc.sig, which((membership(ml.com.hippo.sig)==g)
```

```
                                                                & ( sizes(ml.com.hippo.sig)[[g]] >= 10)))
}

V(net.hippo.coarsen)$name <- paste0("M", 1:14)

empty.module.hippo <- which(sapply(subg.hippo.sig, vcount) == 0)

net.hippo.coarsen <- net.hippo.coarsen - c(paste0("M", empty.module.hippo))
net.hippo.coarsen <- igraph::simplify(net.hippo.coarsen)

### the size of the nodes
l.cicr.hippo<- layout_in_circle(net.hippo.coarsen)
sizes.hippo <- sapply(subg.hippo.sig, vcount)[-empty.module.hippo]
node.size.hippo <- as.vector(sizes.hippo)

# pdf("~/Project1/manuscript_GSEA/results/Figure4B_Hippocampus_coarsen_net_2.pdf",
# useDingbats = FALSE)
plot(net.hippo.coarsen,
          vertex.size= sqrt(node.size.hippo)*2,
          vertex.label= paste0("M",1:12),
                    vertex.label.color="black",
          margin=.5,
          vertex.color = c("white", "#268bd2", "#268bd2", "#268bd2",  "#268bd2",
                           "white", "white",  "white", "white", "white", "white", "#268bd2"),
                    edge.width=sqrt(E(net.hippo.coarsen)$weight),
                    edge.arrow.size=0, layout = l.cicr.hippo, vertex.label.dist = 1,
                    main = "Hippocampus")
```

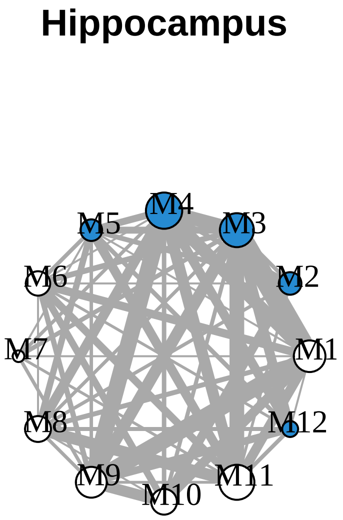


```
# dev.off()
```

```
## mapping to annotation
remap.human.subgraphs.muscle.ml.sig <-
  lapply(subg.muscle.sig, function(x)
    annotation.muscle.oma$human.gene.name[match(V(x)$name, rownames(annotation.muscle.oma))])
remap.human.subgraphs.hippo.ml.sig <-
  lapply(subg.hippo.sig, function(x)
    annotation.hippo.oma$human.gene.name[match(V(x)$name, rownames(annotation.hippo.oma))])
```

```r
### significant ones
# muscle
remapping.human.muscle.ml.sig <-
  remap.human.subgraphs.muscle.ml.sig[sapply(remap.human.subgraphs.muscle.ml.sig, length) > 0]
subgraphs.muscle.sig <-
  subg.muscle.sig[sapply(remap.human.subgraphs.muscle.ml.sig, length) > 0]

## hippocampus
remapping.human.hippo.ml.sig <-
  remap.human.subgraphs.hippo.ml.sig[sapply(remap.human.subgraphs.hippo.ml.sig, length) > 0]
subgraphs.hippo.sig <-
  subg.hippo.sig[sapply(remap.human.subgraphs.hippo.ml.sig, length) > 0]


# final number of the modules
length(remapping.human.muscle.ml.sig)
```

```
## [1] 12
```

```r
length(remapping.human.hippo.ml.sig)
```

```
## [1] 12
```

```r
## loading the human annotation
ensembl.human <- useMart(biomart="ENSEMBL_MART_ENSEMBL", host="www.ensembl.org",
                         path="/biomart/martservice",dataset="hsapiens_gene_ensembl",
                         version = "Ensembl Genes 91")
#
all.ensembl.gene.id.names <- getBM(attributes = c("ensembl_gene_id", "external_gene_name",
                                                  "gene_biotype"),
                                                   values = "*", mart = ensembl.human)
# # 22285 / 22375
all.ensembl.gene.id.names <-
  all.ensembl.gene.id.names[all.ensembl.gene.id.names$gene_biotype == "protein_coding",]


# finding the hubs in the module
hubs.muscle <-
  lapply(subgraphs.muscle.sig, function(x)
    checking.hubs(x, all.ensembl.gene.id.names, annotation.muscle.oma))
hubs.hippo <-
  lapply(subgraphs.hippo.sig, function(x)
    checking.hubs(x, all.ensembl.gene.id.names, annotation.hippo.oma))
```

Perform GO enrichment of the modules. This takes a long time, so the data is loaded automatically, but the code how it was ran is below.

```r
#### GO enrichment - on all ontologies, removes redundancies better than goseq.
top.go.enrichment <- function(targets, universe, genome){
  require(topGO)
  all.genes <- factor(as.integer (universe[,1] %in% targets))
    names(all.genes) <- universe[,1]

  ont <- c("BP", "MF", "CC")
```

```r
  ont.res <- list()
  for(i in 1:length(ont)){
        # nodsize is 5
    GOdata <- new("topGOdata", ontology = ont[i], allGenes = all.genes,
                       geneSel = function(p) p < 1e-2, description = "Test",
                       annot = annFUN.org, mapping=genome, ID="Ensembl")
    resultFisher <- runTest(GOdata, algorithm = "elim", statistic = "fisher")
    res <- GenTable(GOdata, elimFisher = resultFisher, topNodes = 40)
    corrected <- p.adjust(as.numeric(res$elimFisher), method="fdr")
    res$FDR <- corrected
    ont.res[[i]] <- res

  }

  names(ont.res) <- ont

  return(ont.res)

}



# ------------
## GO enrichment of the modules in skeletal muscle

# muscle.results.ml.sig <-
# lapply(remapping.human.muscle.ml.sig,
#        function(x) top.go.enrichment(x, all.ensembl.gene.id.names, "org.Hs.eg.db"))
# saveRDS(muscle.results.ml.sig,
# file = "~/Project1//manuscript_GSEA/data_preprocessing/modules_go/muscle_go_enrichment_conservedmodul

# hippo.results.ml.sig <-
# lapply(remapping.human.hippo.ml.sig,
# function(x) top.go.enrichment(x, all.ensembl.gene.id.names, "org.Hs.eg.db")
# saveRDS(hippo.results.ml.sig,
# file = "~/Project1//manuscript_GSEA/data_preprocessing/modules_go/hippo_go_enrichment_conservedmodule

path.go <- "~/Project1/manuscript_GSEA/data_preprocessing/modules_go/"
muscle.modules <- readRDS(paste0(path.go, "muscle_go_enrichment_conservedmodules.rds"))
hippo.modules <- readRDS(paste0(path.go, "hippo_go_enrichment_conservedmodules.rds"))



# --------------------------------------------
### proteostasis-linked modules
library(plyr)
## skeletal muscle - M1, M2, M3, M8, M9
muscle.proteo.modules <- c(2,3,4,6,8)
muscle.modules.go <- muscle.modules[muscle.proteo.modules]
muscle.proteo.module.go <- lapply(muscle.modules.go, '[[', 1)
names(muscle.proteo.module.go) <- paste0("M", c(2,3,4,6,8))
df.muscle.proteo <- ldply(muscle.proteo.module.go, data.frame, .id = "Cluster.Name")
```

```
## hippocampus - M2, M4, M5, M8, M9
hippo.proteo.modules <- c(2,3,4,12,13)
hippo.modules.go <- hippo.modules[hippo.proteo.modules]
hippo.proteo.module.go <- lapply(hippo.modules.go, '[[', 1)
names(hippo.proteo.module.go) <- paste0("M", c(2,3,4,12,13))
df.hippo.proteo <- ldply(hippo.proteo.module.go, data.frame, .id = "Cluster.Name")
```

Enrichment of the module genes in the GWAS diseases.

Selecting the interesting modules according to proteostasis-linked processes and involvement in GWAS studies.

Skeletal muscle.

```
### proteasome complex
go.human <- go.gsets("Human")
```

```
## Gene ID type for 'human' is: 'EG'
```

```
go.human.sets <- go.human$go.sets
go.human.subs <- go.human$go.subs
gobpsets.human <- go.human.sets[go.human.subs$BP]
goccsets.human <- go.human.sets[go.human.subs$CC]

proteasome.complex.human <-
  goccsets.human[which(names(goccsets.human) == "GO:0000502 proteasome complex")]
proteasome.complex.human.symbol <- mapIds(org.Hs.eg.db,
                                          keys=proteasome.complex.human[[1]],
                                          column="SYMBOL",
                                          keytype="ENTREZID",
                                          multiVals="first")
```

```
## 'select()' returned 1:1 mapping between keys and columns
```

```
# -------------------------
# skeletal muscle module 1 - because of the
module1.human.muscle <- clean.modules(subgraphs.muscle.sig[[1]],
                                      annotation.muscle.oma,
                                      proteasome.complex.human.symbol)
```

```
## 'select()' returned 1:many mapping between keys and columns
```

```
## 'select()' returned 1:1 mapping between keys and columns
```

```
muscle.community.human.module1 <- module1.human.muscle$module
hubs.of.module1 <- module1.human.muscle$hubs
vertex.of.interest.module1 <-
  V(muscle.community.human.module1)[V(muscle.community.human.module1)$label == names(hubs.of.module1)[3
neigh.nodes.module1 <-
  neighbors(muscle.community.human.module1, vertex.of.interest.module1)$label
interested.genes.module1 <-
  c(as.vector(module1.human.muscle$process), neigh.nodes.module1, names(hubs.of.module1))

# pdf("~/Project1/manuscript_GSEA/results/Figure5A_module1_SM.pdf", 10, 7,
# useDingbats = FALSE)
plot.modules(interested.genes.module1, muscle.community.human.module1,
             neigh.nodes.module1, hubs.of.module1,
```

```
            "Skeletal muscle M1\n(GO:0031146 SCF-dependent proteasomal
            ubiquitin-dependent protein catabolic process")
```

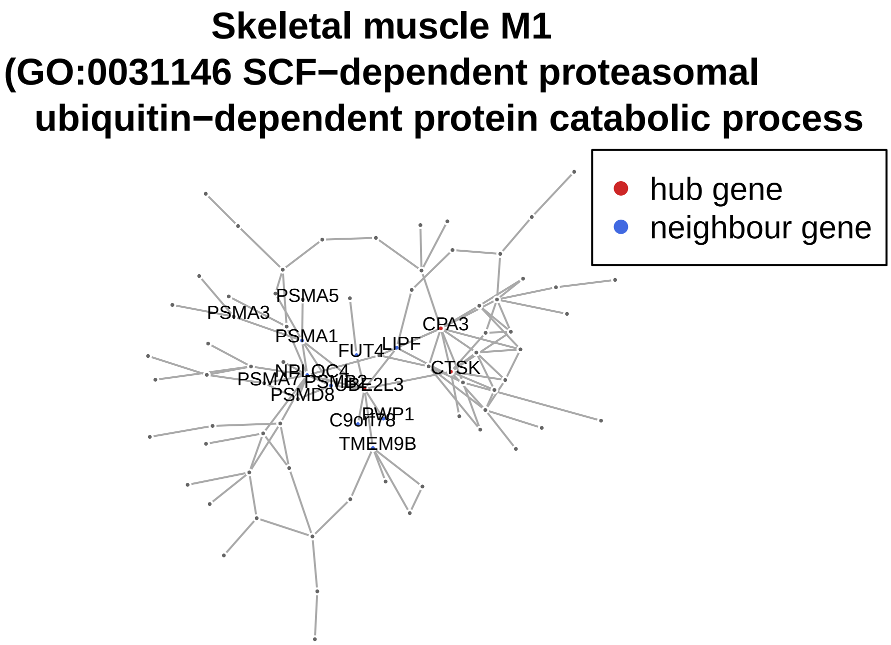


```
# dev.off()



#### ----------------
clean.module3.hippo <- function(subgraph.module,
                                annotation.oma,
                                process.human.symbol){
  tissue.community.module <- subgraph.module
  V(tissue.community.module)$size <- 3
  V(tissue.community.module)$frame.color <- "white"

  ### remap OGs to gene names
  remap.tissue.module <-
                    annotation.oma$human.gene.name[na.omit(match(V(tissue.community.module)$name,
                    rownames(annotation.oma)))]
  names(remap.tissue.module) <- mapIds(org.Hs.eg.db,
                                       keys=as.vector(remap.tissue.module),
                                       column="SYMBOL",
                                       keytype="ENSEMBL",
                                       multiVals="first")

  names(remap.tissue.module)[is.na(names(remap.tissue.module))] <- "AC068775.1"
  genes.process <-
    process.human.symbol[na.omit(match(names(remap.tissue.module),
                                    process.human.symbol))]

  V(tissue.community.module)$label <- names(remap.tissue.module)
  hub.score.module.tissue <- hub.score(tissue.community.module)$vector
  ordered.hubs.module.tissue <-
```

```r
    hub.score.module.tissue[order(hub.score.module.tissue, decreasing = TRUE)][1:3]
  matched.hubs.tissue <-
    annotation.oma$human.gene.name[na.omit(match(names(ordered.hubs.module.tissue),
                                              rownames(annotation.oma)))]

  names(matched.hubs.tissue) <- mapIds(org.Hs.eg.db,
                                       keys = as.vector(matched.hubs.tissue),
                                       column = "SYMBOL",
                                       keytype = "ENSEMBL",
                                       multiVals = "first")

  return(list(module = tissue.community.module, hubs = matched.hubs.tissue, process = genes.process))

}


module3.human.hippo <- clean.module3.hippo(subgraphs.hippo.sig[[3]],
                                           annotation.hippo.oma, proteasome.complex.human.symbol )
```

```
## 'select()' returned 1:many mapping between keys and columns
## 'select()' returned 1:1 mapping between keys and columns
```

```r
hippo.community.human.module3 <- module3.human.hippo$module
hubs.of.module3 <- module3.human.hippo$hubs
vertex.of.interest.module3 <-
  V(hippo.community.human.module3)[V(hippo.community.human.module3)$label == names(hubs.of.module3)[1]]
neigh.nodes.module3 <-
  neighbors(hippo.community.human.module3, vertex.of.interest.module3)$label
interested.genes.module3 <-
  c(as.vector(module3.human.hippo$process), neigh.nodes.module3, names(hubs.of.module3))

# pdf("~/Project1/manuscript_GSEA/results/Figure5B_module3_hippo.pdf", 10, 7, useDingbats = FALSE)
plot.modules(interested.genes.module3, hippo.community.human.module3,
             neigh.nodes.module3, hubs.of.module3,
             "Hippocampus M3\n(GO:0000209 protein polyubiquitination)")
```

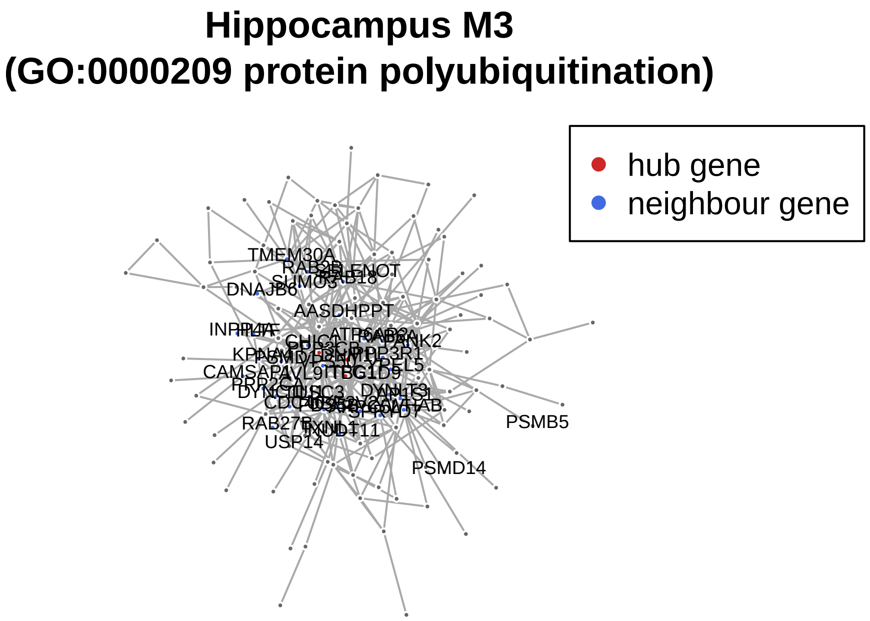


```r
# dev.off()


# ---------------------------
# Figure S10.
# --------------------------



# ---------------------------------
# Skeletal muscle additional module
module12.human.muscle <-
  clean.modules(subgraphs.muscle.sig[[12]], annotation.muscle.oma, proteasome.complex.human.symbol )
```

```
## 'select()' returned 1:1 mapping between keys and columns
## 'select()' returned 1:1 mapping between keys and columns
```

```r
muscle.community.human.module12 <- module12.human.muscle$module
hubs.of.module12 <- module12.human.muscle$hubs
vertex.of.interest.module12 <-
  V(muscle.community.human.module12)[V(muscle.community.human.module12)$label == names(hubs.of.module12)
neigh.nodes.module12 <-
  neighbors(muscle.community.human.module12, vertex.of.interest.module12)$label
interested.genes.module12 <-
  c(as.vector(module12.human.muscle$process), neigh.nodes.module12, names(hubs.of.module12))

# pdf("~/Project1/manuscript_GSEA/results/FigureS9_module12_SM.pdf", 10, 7, useDingbats = FALSE)
plot.modules(interested.genes.module12, muscle.community.human.module12,
             neigh.nodes.module12, hubs.of.module12, "Skeletal muscle M12\n(GO:000209 protein polyubiqu
```

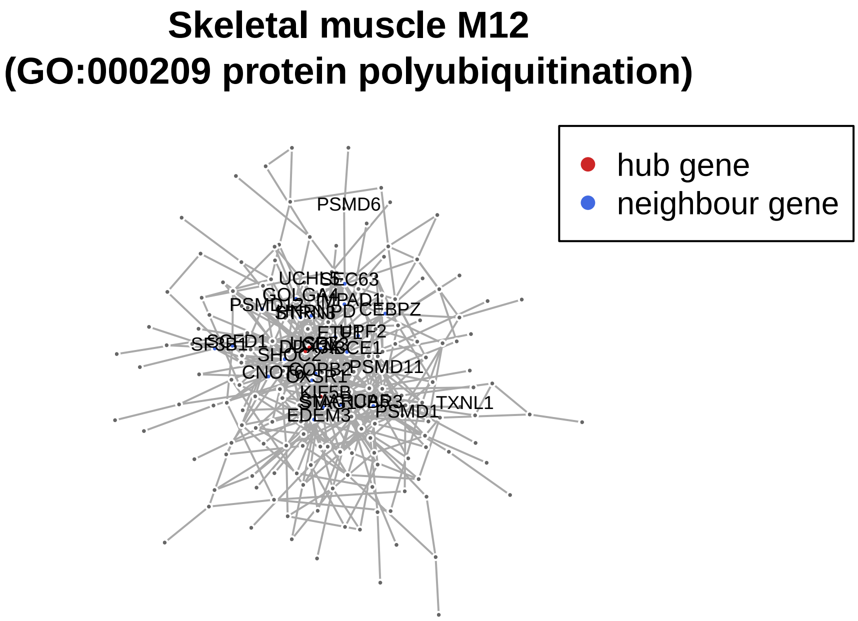


```
# dev.off()





# ------------------------------------
### Hippocampus additional module
clean.module4.hippo <- function(subgraph.module, annotation.oma, process.human.symbol){
  tissue.community.module <- subgraph.module
  V(tissue.community.module)$size <- 3
  V(tissue.community.module)$frame.color <- "white"

  ### remap OGs to gene names
  remap.tissue.module <-
                    annotation.oma$human.gene.name[na.omit(match(V(tissue.community.module)$name,
                    rownames(annotation.oma)))]
  names(remap.tissue.module) <- mapIds(org.Hs.eg.db,
                                          keys=as.vector(remap.tissue.module),
                                          column="SYMBOL",
                                          keytype="ENSEMBL",
                                          multiVals="first")

  names(remap.tissue.module)[is.na(names(remap.tissue.module))] <- "POLR2A"
  genes.process <-
    process.human.symbol[na.omit(match(names(remap.tissue.module), process.human.symbol))]

  V(tissue.community.module)$label <- names(remap.tissue.module)
  hub.score.module.tissue <- hub.score(tissue.community.module)$vector
  ordered.hubs.module.tissue <-
    hub.score.module.tissue[order(hub.score.module.tissue, decreasing = TRUE)][1:3]
  matched.hubs.tissue <-
    annotation.oma$human.gene.name[na.omit(match(names(ordered.hubs.module.tissue), rownames(annotation
```

```
  names(matched.hubs.tissue) <- mapIds(org.Hs.eg.db,
                                       keys = as.vector(matched.hubs.tissue),
                                       column = "SYMBOL",
                                       keytype = "ENSEMBL",
                                       multiVals = "first")

  return(list(module = tissue.community.module, hubs = matched.hubs.tissue, process = genes.process))

}


# --------------------------
# hippocampus module 4 - in proteostasis and coronary artery
module4.human.hippo <- clean.module4.hippo(subgraphs.hippo.sig[[4]], annotation.hippo.oma, proteasome.c
```

```
## 'select()' returned 1:1 mapping between keys and columns
## 'select()' returned 1:1 mapping between keys and columns
```

```
hippo.community.human.module4 <- module4.human.hippo$module
hubs.of.module4 <- module4.human.hippo$hubs
vertex.of.interest.module4 <- V(hippo.community.human.module4)[V(hippo.community.human.module4)$label ==
neigh.nodes.module4 <- neighbors(hippo.community.human.module4, vertex.of.interest.module4)$label
interested.genes.module4 <- c(as.vector(module4.human.hippo$process), neigh.nodes.module4, names(hubs.o

# pdf("~/Project1/manuscript_GSEA/results/FigureS9_module4_hippo.pdf", 10, 7, useDingbats = FALSE)
plot.modules(interested.genes.module4, hippo.community.human.module4,
             neigh.nodes.module4, hubs.of.module4,
             "Hippocampus M4\n(GO:1904874 positive regulation
             of telomerase RNA localization to Calaj body)")
```

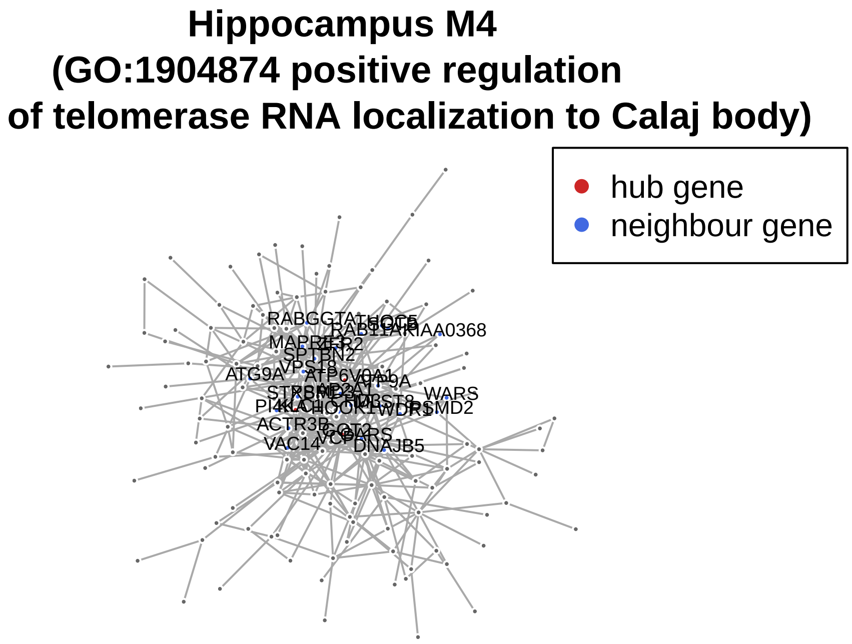


```
# dev.off()
```

Figure 5C. GWAS heatmap.

```r
# mapping to GWAS studies - based on gene-level p-values from PASCAL
# downloaded from - http://regulatorycircuits.org/download.html -
# Supplementary data and code: GWAS_gene_scores - 12MB  (University of Lausanne)

gwas.directory <- "~/Project1/manuscript_GSEA/data_preprocessing/GWAS_gene_scores_v1"
gwas.files <- list.files(path = gwas.directory, full.names=TRUE)
gwas.files <- gwas.files[-1]
ldf <- lapply(gwas.files, function(x) read.table(x, header = TRUE))
names(ldf) <- sub("\\.txt", "", list.files(path = gwas.directory))[-1]

# selecting the genes that are having higher of association with GWAS
high.top.ranked.ldf <- lapply(ldf, function(x) x[x$pvalue < 0.1,])


interested.genes <- c( names(hubs.of.module3), as.vector(module3.human.hippo$process),
                       names(hubs.of.module4), as.vector(module4.human.hippo$process),
                       names(hubs.of.module1), as.vector(module1.human.muscle$process),
                       names(hubs.of.module12), as.vector(module12.human.muscle$process) )



interested.genes <- unique(interested.genes)

modules.gwas <-
  lapply(high.top.ranked.ldf, function(x) x[na.omit(match(interested.genes, x$gene_id)),])

# select the gwas that is age related
gwas.age.related <- c("11_rheumatoid_arthritis", "12_multiple_sclerosis", "15_alzheimers",
                      "17_parkinsons_disease", "18_hdl_cholesterol","19_ldl_cholesterol",
                      "20_total_cholesterol", "21_triglycerides", "22_blood_pressure_systolic",
                      "23_coronary_artery_disease", "24_fasting_glucose", "25_glycated_hemoglobin",
                      "26_type_2_diabetes", "27_2hr_glucose", "28_fasting_proinsulin",
                      "29_insulin_secretion", "30_insulin_resistance", "31_beta-cell_function",
                      "32_fasting_insulin", "35_macular_degeneration_neovascular", "36_macular_degenera
                      "37_osteoporosis")

df.modules <- ldply(modules.gwas[gwas.age.related], data.frame)

df.modules$seq <- with(df.modules, ave(pvalue, .id, gene_symbol, FUN = seq_along))
gwas.modules <- reshape2::dcast(.id + seq ~ gene_symbol, data = df.modules, value.var = "pvalue")

gg.mod <- gwas.modules
gg.mod <- gg.mod[,-1]
# missing values are replaced
gg.mod[is.na(gg.mod)] <- 0.99
gg.mod <- -log10(gg.mod)
gg.mod <- gg.mod[,-1]


names.gwas.module <- gwas.modules[,1]
# renaming for the labels on the heatmap
names.gwas.modules <- c("Rheumatoid arthritis", "Multiple sclerosis", "Alzheimer's disease",
                        "Parkinson's disease", "HDL cholesterol", "LDL cholesterol",
```

```
                            "Total cholesterol", "Triglycerides", "Blood pressure systolic",
                            "Coronary artery disease", "Fasting glucose", "Glycated hemoglobin",
                            "Type 2 diabetes", "2hr glucose", "Fasting proinsulin",
                            "Insulin secretion", "Insulin resistance", "Beta-cell function",
                            "Fasting insulin", "Macular degeneration neovasc." ,
                            "Macular degeneration dry", "Osteoporosis")


annotation.mat.gwas <- HeatmapAnnotation(text = anno_text(names.gwas.modules,
                                     rot = 45, just = "left", offset = unit(2, "mm")),
                                     gap = unit(c(2, 4), "mm"),
                                     annotation_height = unit(c(0.5, 0.5, 1.5, 1.5), "cm"))


# pdf("~/Project1/manuscript_GSEA/results/Figure5C_GWAS_heatmap_2.pdf", 11, 6, useDingbats = FALSE)
Heatmap(t(as.matrix(gg.mod)),
        heatmap_legend_param = list(color_bar = "discrete"),
        name =  "GWAS gene score\n(-log10(p-value))",
        cluster_columns = FALSE,
        show_column_dend = FALSE,
        top_annotation = annotation.mat.gwas,
        col = colorRamp2(seq(0,5,1), brewer.pal(6,"Paired")))
```

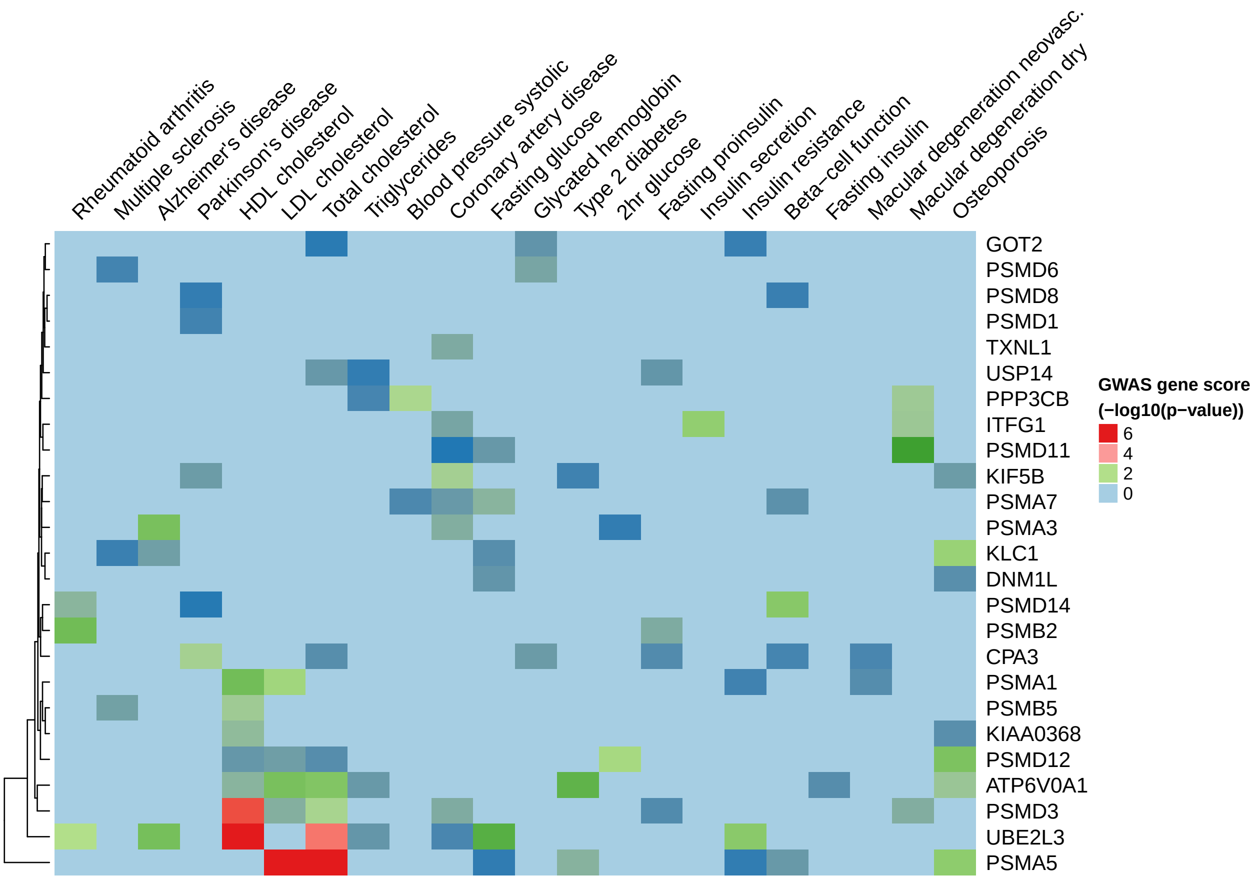


```
# dev.off()

## ggmod should go to the Suplement table
```

Generating the dietary restriction volcano plot for Figure 5D.

```
#### ---------------------------------------

### VOLCANO PLOTS for caloric restriction
path.cr <- "~/Project1/manuscript_GSEA/data_preprocessing/hsapiens/caloric_restriction_data/"
deg.human.dr <- readRDS(paste0(path.cr, "differential_expression_human_dietary_restriction.rds"))

## select genes from GWAS heatmap above
colnames.gwas <- mapIds(org.Hs.eg.db,
                        keys=colnames(gg.mod),
                        column="ENSEMBL",
                        keytype="SYMBOL",
                        multiVals="first")
```

```
## 'select()' returned 1:1 mapping between keys and columns
```

```
deg.human.muscle.dr.candidate.genes <-
  deg.human.dr[na.omit(match(as.vector(colnames.gwas), rownames(deg.human.dr))),]

deg.human.muscle.dr.candidate.genes$Gene <- mapIds(org.Hs.eg.db,
                                                   keys=rownames(deg.human.muscle.dr.candidate.genes),
                                                   column="SYMBOL",
                                                   keytype="ENSEMBL",
                                                   multiVals="first")
```

```
## 'select()' returned 1:1 mapping between keys and columns
```

```
deg.human.muscle.dr.candidate.genes$Significant <-
  ifelse(deg.human.muscle.dr.candidate.genes$adj.P.Val < 0.05, "FDR < 0.05", "FDR > 0.05")

# candidate genes in human dietary restriciton analysis
library(ggrepel)
p.candidate.genes <- ggplot(deg.human.muscle.dr.candidate.genes, aes(x = logFC, y = -log10(P.Value))) +
  geom_point(aes(color = Significant)) +
  scale_color_manual(values = c("red", "grey")) +
  theme_bw(base_size = 12) + theme(legend.position = "bottom") +
  ggtitle("H. sapiens - Caloric restriction (skeletal muscle)") +
  xlab(expression(log[2]~fold~change~(CR/Control))) + ylab(expression(-log[10]~pvalue)) + xlim(-2,4) +
  geom_vline(xintercept = 0) +
  geom_text_repel(
    data = subset(deg.human.muscle.dr.candidate.genes, adj.P.Val < 0.05),
    aes(label = Gene),
    size = 5,
    box.padding = unit(0.35, "lines"),
    point.padding = unit(0.3, "lines")
  )

# pdf("~/Project1/manuscript_GSEA/results/Figure5D_volcano_plots_on_caloric_restriction.pdf",
# 7, 5, useDingbats = FALSE)
plot(p.candidate.genes)
```

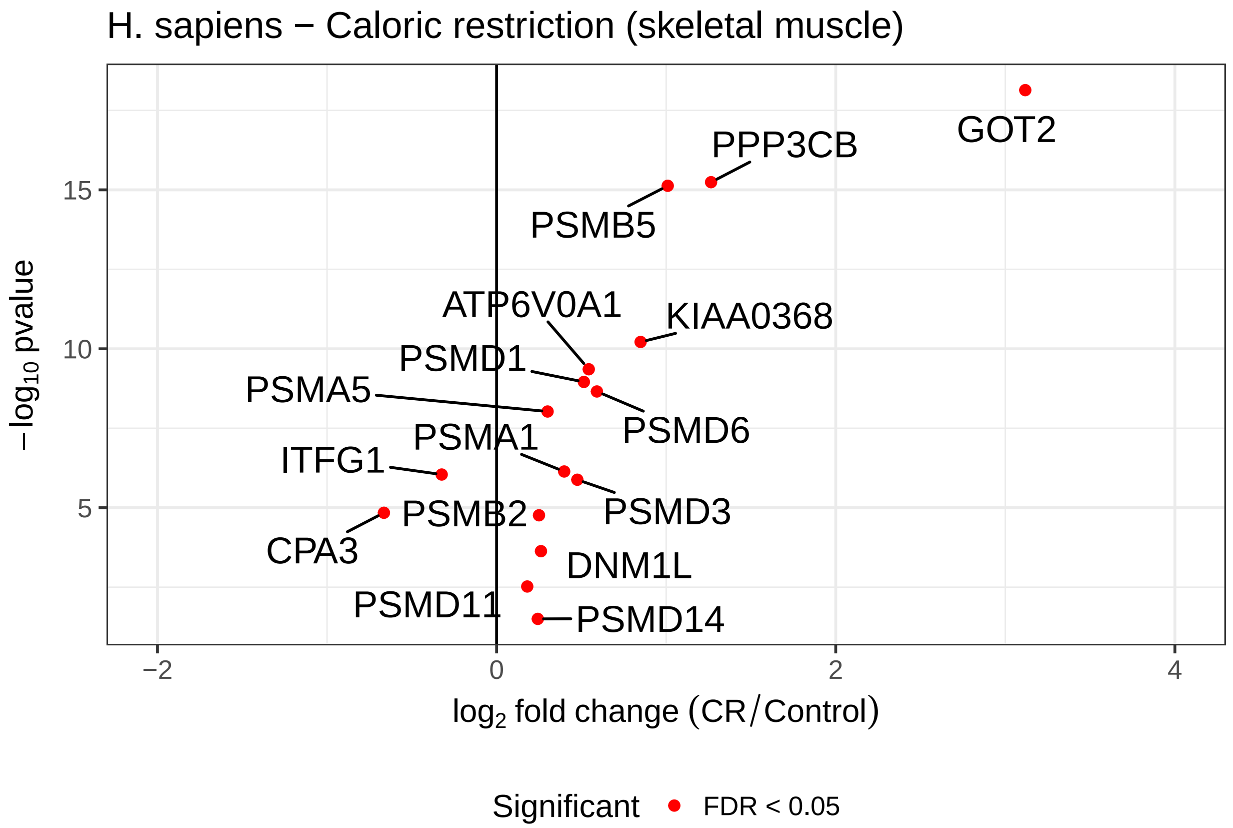


```
# dev.off()
```

Supplemental material for this part. Table S8 and Table S9.

```
# -------------------------------
### Table S8.
# -------------------------------

### proteostasis-linked modules

## skeletal muscle - M1, M6, M7, M8, M11, M12
muscle.proteo.modules <- c(1,6,7,8,11,12)
muscle.modules.go <- muscle.modules[muscle.proteo.modules]
muscle.proteo.module.go <- lapply(muscle.modules.go, '[[', 1)
names(muscle.proteo.module.go) <- paste0("M", c(1,6,7,8,11,12))
df.muscle.proteo <- ldply(muscle.proteo.module.go, data.frame, .id = "Cluster.Name")


# GWAS linked modules - M3, M4, M5, M12
muscle.disease.modules <- c(3,4,5,12)
muscle.modules.gwas <- muscle.gwas.sig[muscle.disease.modules]
names(muscle.modules.gwas) <- paste0("M", c(3,4,5,12))
df.muscle.proteo.gwas <- ldply(muscle.modules.gwas, data.frame, .id = "Cluster.Name")


# proteostasis-linked
## hippocampus - M2, M3, M4, M5, M12
hippo.proteo.modules <- c(2,3,4,5,12)
```

```r
hippo.modules.go <- hippo.modules[hippo.proteo.modules]
hippo.proteo.module.go <- lapply(hippo.modules.go, '[[', 1)
names(hippo.proteo.module.go) <- paste0("M", c(2,3,4,5,12))
df.hippo.proteo <- ldply(hippo.proteo.module.go, data.frame, .id = "Cluster.Name")


# GWAS linked modules
hippo.disease.modules <- c(4,6)
hippo.modules.gwas <- hippo.gwas.sig[hippo.disease.modules]
names(hippo.modules.gwas) <- paste0("M", 4)
df.hippo.proteo.gwas <- ldply(hippo.modules.gwas, data.frame, .id = "Cluster.Name")


df.module.sizes <- data.frame(module.names = paste0("M", 1:12),
                              muscle.modules.gene.size = sapply(subgraphs.muscle.sig, vcount),
                              hippo.modules.gene.size = sapply(subgraphs.hippo.sig, vcount))

df.modules.summary.stats <-
  list(Module.sizes = df.module.sizes,
             Muscle.GO.proteo.enrich = df.muscle.proteo,
             Hippo.GO.proteo.enrich = df.hippo.proteo,
             Muscle.GWAS.proteo.enrich = df.muscle.proteo.gwas,
             Hippo.GWAS.proteo.enrich = df.hippo.proteo.gwas)



#library(WriteXLS)
# WriteXLS(df.modules.summary.stats,
#     ExcelFileName = "~/Project1/manuscript_GSEA/supplementary_data/Table_S8.xlsx", SheetNames = NULL)



# ----------------------------
# Table S9.
# ---------------------------

# matrix with gene scores associated to GWAS diseases
gg.mod.supplement <- gwas.modules[,-2]
gwas.diseases.genes <- gg.mod.supplement


# WriteXLS(gwas.diseases.genes,
#    ExcelFileName = "~/Project1/manuscript_GSEA/supplementary_data/Table_S9.xlsx", SheetNames = NULL)



# ----------------------------
# Figure S9.
# ---------------------------

# Plotting random networks.

########
```

```r
# for p-value calculations
# https://www.ncbi.nlm.nih.gov/pubmed/21044043
# calculate significance for the random networks
calculate.p.val <- function(observed, randomized){
  # here is mean to summarize 100 randomizations
  random <- mean(randomized)
  return( (sum(abs(random) > abs(observed)) + 1) / (length(randomized) + 1) )
}



setwd("~/Project1/manuscript_GSEA/results/")
# hippo_random_nets <- readRDS("random_nets_hippo100.rds")
hippo_random_nets <- readRDS("random_nets_hippo100.rds")
summary.hippo <- as.matrix(sapply(hippo_random_nets, '['))


# muscle_random_nets <- readRDS("random_nets_muscle100.rds")
muscle_random_nets <- readRDS("random_nets_muscle100.rds")
summary.muscle <- as.matrix(sapply(muscle_random_nets, '['))


## number of connections
obs.no.connections.muscle <- 2887
obs.no.connections.hippo <- 3353


#pdf("SupplementS9_randomnets100_2.pdf", 7,7)
par(mfrow = c(2,1))
# for number of connections
hist(as.numeric(summary.muscle[,1]), xlim = c(2700, max(as.numeric(summary.muscle[,1]))),
  xlab = "Number of connections\n(skeletal muscle)", main = "Skeletal muscle network")
abline(v = obs.no.connections.muscle, col = "red", lwd = 2)
legend(3000,30, "Observed",
lty=1,
lwd=2.5, col= "red")

hist(as.numeric(summary.hippo[,1]), xlim = c(3200, max(as.numeric(summary.hippo[,1]))),
  xlab = "Number of connections\n(hippocampus)", main = "Hippocampus network")
abline(v = obs.no.connections.hippo, col = "red", lwd = 2)
```

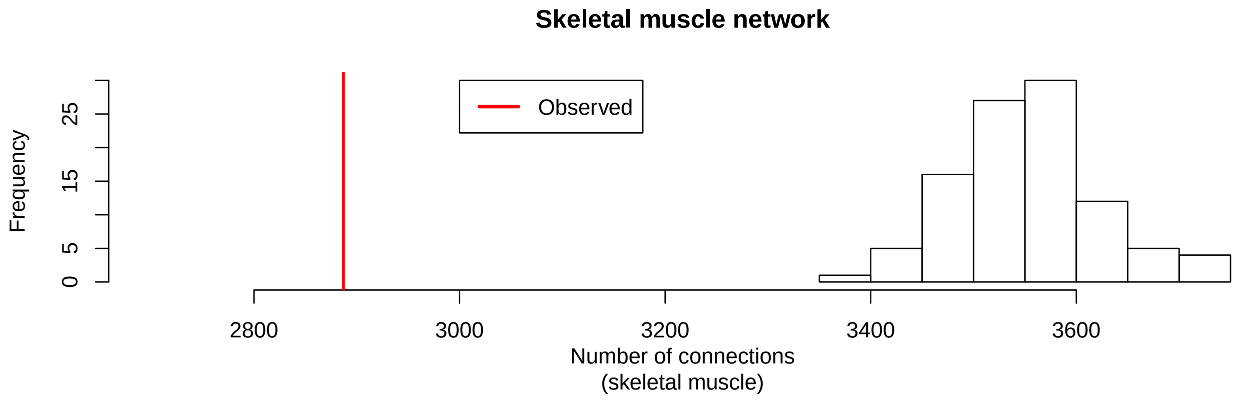


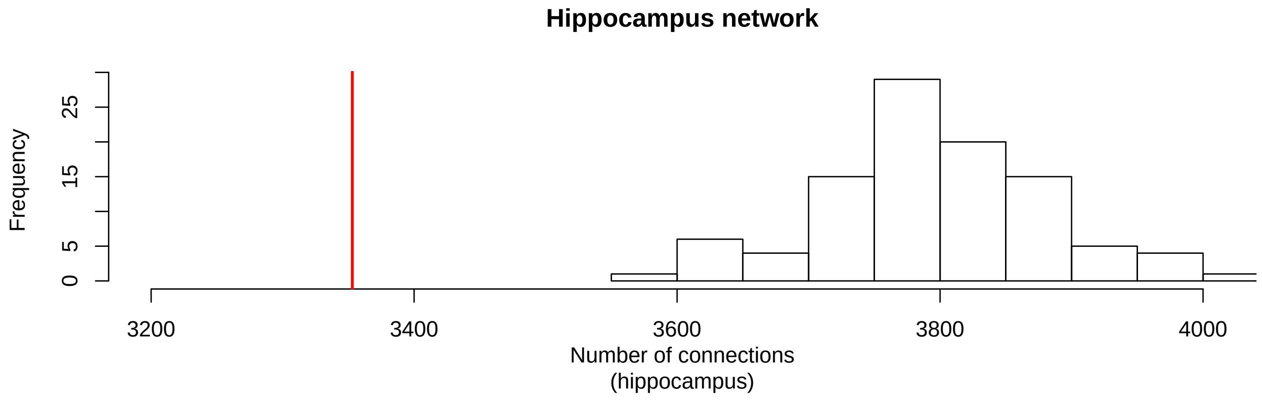


```
# dev.off()
```

```
sessionInfo()
```

```
## R version 3.4.3 (2017-11-30)
## Platform: x86_64-apple-darwin15.6.0 (64-bit)
## Running under: macOS Sierra 10.12.1
##
## Matrix products: default
## BLAS: /Library/Frameworks/R.framework/Versions/3.4/Resources/lib/libRblas.0.dylib
## LAPACK: /Library/Frameworks/R.framework/Versions/3.4/Resources/lib/libRlapack.dylib
##
## locale:
## [1] en_US.UTF-8/en_US.UTF-8/en_US.UTF-8/C/en_US.UTF-8/en_US.UTF-8
##
## attached base packages:
##  [1] parallel  stats4    grid      stats     graphics  grDevices utils
##  [8] datasets  methods   base
##
## other attached packages:
##  [1] GO.db_3.5.0            dplyr_0.7.4            data.table_1.10.4-3
##  [4] ggrepel_0.7.0          cowplot_0.9.2          graphite_1.24.1
##  [7] gage_2.28.2            org.Ce.eg.db_3.5.0     org.Dm.eg.db_3.5.0
## [10] org.Mm.eg.db_3.5.0     org.Hs.eg.db_3.5.0     AnnotationDbi_1.40.0
## [13] IRanges_2.12.0         S4Vectors_0.16.0       Biobase_2.38.0
## [16] BiocGenerics_0.24.0    igraph_1.1.2           circlize_0.4.3
## [19] ComplexHeatmap_1.17.1  WriteXLS_4.0.0         plyr_1.8.4
```

```
## [22] goseq_1.30.0           geneLenDataBase_1.14.0 BiasedUrn_1.07
## [25] biomaRt_2.34.2         RobustRankAggreg_1.1   Hmisc_4.1-1
## [28] ggplot2_2.2.1          Formula_1.2-2          survival_2.41-3
## [31] lattice_0.20-35        RColorBrewer_1.1-2
##
## loaded via a namespace (and not attached):
##  [1] nlme_3.1-131.1             bitops_1.0-6
##  [3] matrixStats_0.53.1         bit64_0.9-7
##  [5] progress_1.1.2             httr_1.3.1
##  [7] rprojroot_1.3-2            GenomeInfoDb_1.14.0
##  [9] tools_3.4.3                backports_1.1.2
## [11] R6_2.2.2                   rpart_4.1-13
## [13] DBI_0.8                    lazyeval_0.2.1
## [15] mgcv_1.8-23                colorspace_1.3-2
## [17] nnet_7.3-12                GetoptLong_0.1.6
## [19] gridExtra_2.3              prettyunits_1.0.2
## [21] RMySQL_0.10.14             curl_3.1
## [23] bit_1.1-12                 compiler_3.4.3
## [25] graph_1.56.0               htmlTable_1.11.2
## [27] DelayedArray_0.4.1         labeling_0.3
## [29] rtracklayer_1.38.3         scales_0.5.0
## [31] checkmate_1.8.5            rappdirs_0.3.1
## [33] stringr_1.3.0              digest_0.6.15
## [35] Rsamtools_1.30.0           foreign_0.8-69
## [37] rmarkdown_1.9              XVector_0.18.0
## [39] base64enc_0.1-3            pkgconfig_2.0.1
## [41] htmltools_0.3.6            htmlwidgets_1.0
## [43] rlang_0.2.0                GlobalOptions_0.0.12
## [45] rstudioapi_0.7             RSQLite_2.0
## [47] bindr_0.1                  shape_1.4.4
## [49] BiocParallel_1.12.0        acepack_1.4.1
## [51] RCurl_1.95-4.10            magrittr_1.5
## [53] GenomeInfoDbData_1.0.0     Matrix_1.2-12
## [55] Rcpp_0.12.15               munsell_0.4.3
## [57] stringi_1.1.6              yaml_2.1.17
## [59] SummarizedExperiment_1.8.1 zlibbioc_1.24.0
## [61] blob_1.1.0                 Biostrings_2.46.0
## [63] splines_3.4.3              GenomicFeatures_1.30.3
## [65] KEGGREST_1.18.1            knitr_1.20
## [67] pillar_1.2.1               GenomicRanges_1.30.3
## [69] rjson_0.2.15               reshape2_1.4.3
## [71] glue_1.2.0                 XML_3.98-1.10
## [73] evaluate_0.10.1            latticeExtra_0.6-28
## [75] png_0.1-7                  gtable_0.2.0
## [77] assertthat_0.2.0           tibble_1.4.2
## [79] GenomicAlignments_1.14.1   memoise_1.1.0
## [81] bindrcpp_0.2               cluster_2.0.6
```
